# Identification of a proteolysis regulator for an essential enzyme in *Mycobacterium tuberculosis*

**DOI:** 10.1101/2024.03.29.587195

**Authors:** Shoshanna C. Kahne, Jin Hee Yoo, James Chen, Kehilwe Nakedi, Lakshminarayan M. Iyer, Gregory Putzel, Nora M. Samhadaneh, Alejandro Pironti, L. Aravind, Damian C. Ekiert, Gira Bhabha, Kyu Y. Rhee, K. Heran Darwin

## Abstract

In *Mycobacterium tuberculosis* proteins that are post-translationally modified with Pup, a prokaryotic ubiquitin-like protein, can be degraded by proteasomes. While pupylation is reversible, mechanisms regulating substrate specificity have not been identified. Here, we identify the first depupylation regulators: CoaX, a pseudokinase, and pantothenate, an essential, central metabolite. In a Δ*coaX* mutant, pantothenate synthesis enzymes were more abundant, including PanB, a substrate of the Pup-proteasome system. Media supplementation with pantothenate decreased PanB levels in a *coaX* and Pup-proteasome-dependent manner. *In vitro*, CoaX accelerated depupylation of Pup∼PanB, while addition of pantothenate inhibited this reaction. Collectively, we propose CoaX contributes to proteasomal degradation of PanB by modulating depupylation of Pup∼PanB in response to pantothenate levels.

**One Sentence Summary:** A pseudo-pantothenate kinase regulates proteasomal degradation of a pantothenate synthesis enzyme in *M. tuberculosis*.

## Main Text

*Mycobacterium tuberculosis* (*Mtb*), the causative agent of tuberculosis (TB), is responsible for an ancient and enduring global pandemic. While about 85% of TB cases are curable, therapies are burdensome, typically requiring 4-6 months of antibiotic treatment for drug susceptible strains (*1*). Barriers of access to treatment, among other factors, contributed to an estimated 1.6 million deaths from TB in 2021 (*1*). Without new therapeutic options, increasing incidence of multidrug- and extensively drug-resistant strains forebode a dire future. Increased mechanistic understanding of *Mtb* biology contributing to virulence may reveal new approaches for tackling this devastating pandemic.

The Pup-proteasome system (PPS), an ATP-dependent proteolytic pathway in *Mtb*, contributes to virulence in mouse infection models (*2–6*). Like the eukaryotic ubiquitin-proteasome system, the PPS degrades numerous proteins when they are post-translationally modified with Pup (*7, 8*). Like ubiquitin, Pup must be activated prior to substrate ligation. However, in contrast to eukaryotes that encode hundreds of ubiquitin ligases and deubiquitinases to confer specificity and regulate post-translational modification, the PPS activates, ligates, and removes Pup from substrates using only two enzymes.

Pup is translated with a C-terminal glutamine that must be deamidated by the enzyme <u>d</u>eamidase <u>o</u>f <u>P</u>up (Dop) before Pup can be ligated to a surface exposed lysine by the sole Pup ligase, <u>p</u>roteasome <u>a</u>ccessory <u>f</u>actor <u>A</u> (PafA) (*9, 10*). Dop can also reverse this pupylation reaction (*10, 11*). Together these enzymes define the “pupylome” (*4*). Pupylated proteins can only be degraded by the proteasome when they are recruited and unfolded by the mycobacterial proteasome <u>A</u>TPase (Mpa) (*8, 12, 13*). While enzymatic mechanisms of PPS enzymes have been extensively characterized, little is known about how their activities are regulated (*14*).

The steady state abundance of any protein that can be pupylated is a function of its pupylation, depupylation, and degradation rates. Global changes in the pupylome in response to growth conditions have been described in both *Mtb* (*15*) and *Mycobacterium smegmatis* (*Msm*) (*16*), but the molecular mechanisms underlying these phenotypes are unknown. Motifs strongly defining pupylation targets, including discrimination among potential surface exposed lysines on a single substrate, have been elusive. Verified PPS substrates are pupylated with variable efficiency *in vitro* and amino acid charges proximal to the target lysine in the three-dimensional conformation of substrates only partially explain this variation (*17*). The lack of a specific sequence or structural feature strongly predicting lysine pupylation has led to the hypothesis pupylation-specificity is low and the pupylated status of a protein is determined to a greater extent by depupylation. Depupylation also varies *in vitro* and this has been hypothesized to be conferred by differential steric hinderance associated with the positioning of Pup in the Dop active site (*18, 19*). This steric hinderance may be affected by substrate interaction with the Dop loop, a conserved stretch of approximately 40 amino acids that is inferred to be disordered and proximal to the active site (*20*). Deletion of the Dop loop in *Msm* accelerates depupylation *in vitro* (*19*) and yields a diminished pupylome *in vivo* in *Msm* and *Mtb* (*19, 21*). Taken together, these data reveal intrinsic factors affecting specificity and activity of Dop and PafA but fail to illuminate how pupylation of an individual substrate could be differentially regulated in response to cellular needs or environmental conditions.

In this study, we found CoaX (Rv3600c) regulated depupylation of pupylated PanB (Pup∼PanB). PanB is an essential enzyme that catalyzes an early committed step in pantothenate (vitamin B5) synthesis in mycobacteria. In the presence of excess pantothenate, the ultimate product of the PanB pathway, depupylation by Dop was inhibited in a CoaX-dependent manner, and proteasomes degraded Pup∼PanB. Thus, we propose PanB levels are regulated by a negative feedback loop mediated by CoaX and the PPS in *Mtb*.

## Results

To identify potential depupylation regulators, we immunoprecipitated carboxyl (C)-terminally epitope-tagged Dop and PafA (referred to as “Dop_TAP_” and “PafA_TAP_”, respectively) from *Mtb* lysates (see **Supplemental table S1** for strains used). We observed by silver stain a 25-37 kD protein that co-purified with Dop but not PafA (**Fig. 1A**). Mass spectrometry (MS) identified this protein as CoaX (Rv3600c), a predicted type-III pantothenate kinase (PanK) (**Supplemental table S2**). Quantitative MS of all proteins that immunoprecipitated with Dop and PafA revealed CoaX was the fourth most abundant protein in Dop samples and undetectable in the PafA samples (**Supplemental table S3**). CoaX has never been identified as a member of the pupylome, and immunoblotting with a monoclonal antibody to Pup showed CoaX association with Dop could not be explained by pupylation (**Fig. 1B**).

**Fig. 1:**
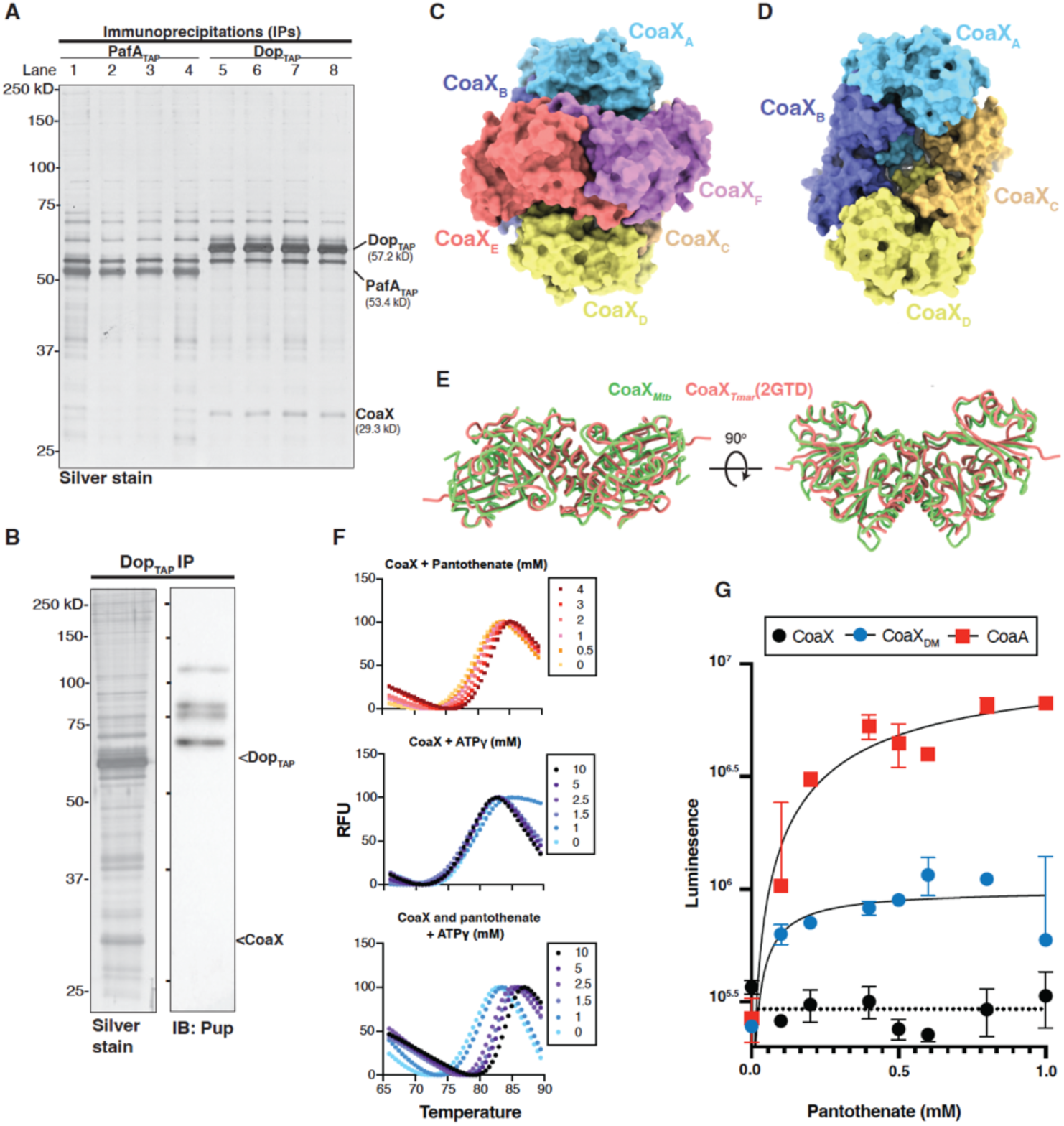
Dop interacts with CoaX, a pseudo-pantothenate kinase. **(A)** Elutions from PafA_TAP_ (lanes 1-4) and Dop_TAP_ (lanes 5-8) immunoprecipitations (IPs) from *Mtb* were visualized by silver stain. Dop_TAP_, PafA_TAP_, and CoaX, a Dop-specific binding partner identified by mass spectrometry, are indicated. **(B)** Elution from a Dop_TAP_ IP from *Mtb* (left, silver stain) was probed with monoclonal antibodies to Pup (right, immunoblot). (**C**) Structure of CoaX*_Mtb_* hexamer and (**D**) tetramer rendered as a molecular surface. Dimer 1 includes CoaX_A_ (light blue) and CoaX_B_ (slate); dimer 2, CoaX_C_ (gold) and CoaX_D_ (yellow); dimer 3, CoaX_E_ (light red) and CoaX_F_ (light purple). (**E**) CoaX*_Mtb_* and CoaX*_Tmar_* dimer (PDB 2GTD) were compared (RMSD 1.39 Å over 419 atoms). **(F)** Thermal stability of CoaX with pantothenate (top), ATPγS (middle), or ATPγS in the presence of 4 mM pantothenate (bottom) was measured by differential scanning fluorimetry. **(G)** CoaA (red squares), CoaX (black circles), or CoaX_DM_ (double mutant CoaX*_Mtb_* R8G and H226G, blue circles) activity was determined by ADP-Glo Luminescent assays (n=2). Luminescence is proportional to ADP formed in each reaction. Empirically determined average baseline fluorescence of reactions containing no enzyme is shown as a dotted line. Michaelis-Menten curves for CoaX_DM_ and CoaA are shown.

PanK enzymes phosphorylate pantothenate in the first step of coenzyme A (coA) synthesis (*22*). CoA, and its derivatives, function as acyl carriers and participate as cofactors or substrates in hundreds of chemical reactions supporting basic metabolism and bacterial survival (*23*). While *Mtb* can use extracellular pantothenate to survive *in vitro* and *in vivo* through an unknown uptake mechanism, imported pantothenate is insufficient to support full virulence in a mouse infection model (*24*). In the absence of supplementation, all genes encoding pantothenate synthesis enzymes are essential in *Mtb* (*25–27*).

PanK enzymes are classified into three types. Type I PanK enzymes (herein referred to as CoaA) belong to the P-loop kinase family (*28, 29*), while type II and type III enzymes belong to the acetate and sugar kinases/Hsc70/actin superfamily (type III PanK enzymes herein referred to as CoaX) (*30*). Many bacteria, including *Mtb*, are predicted to encode more than one PanK enzyme; however, the biological consequences associated with encoding multiple PanK enzymes has not been well studied in many species. CoaA and CoaX from *Bacillus subtilis* are both active *in vitro* and either is individually sufficient to support life (*31*). In *Bacillus anthracis* CoaX is active *in vitro* and essential; however, a gene with homology to a eukaryotic type II PanK enzyme did not show activity *in vitro* (*32, 33*).

In *Mtb*, CoaA (CoaA*_Mtb_*), but not CoaX (CoaX*_Mtb_*), is essential and active *in vitro* (*34*). The gene encoding CoaX (*coaX*) is found across mycobacteria, including in *Mycobacterium leprae* (*Mlp*), the causative agent of leprosy, which has a markedly reduced and decayed genome compared to *Mtb* (*35*). Thus, genes conserved in *Mlp*, including those encoding the PPS and pantothenate synthesis enzymes, are thought to represent a core genome of minimal functions required for pathogenesis and survival.

To structurally characterize the Dop and CoaX interaction, we co-expressed *dop_Msm_* and *coaX_Mtb_* in *E. coli* and attempted to purify the complex for single particle cryo-electron microscopy (cryo-EM) (**Supplemental Fig, S1**). While we were unable to identify complexes containing Dop*_Msm_* and CoaX*_Mtb_*, our dataset revealed two predominant oligomerization states of CoaX_Mtb_, a hexamer (**Fig. 1C**) and a tetramer (**Fig. 1D**). Similar to other CoaX structures published to date (*30, 33, 36, 37*), CoaX*_Mtb_* formed dimers with the presumed active sites at the dimer interface (**Supplemental Fig. S2 and Table S4**). Dimers of CoaX assembled to form either tetramers or hexamers, for which we solved structures to average resolutions of 2.81Å and 2.59Å, respectively. CoaX*_Mtb_* dimers showed overall structural similarity to the crystal structure of CoaX dimers from *Thermotoga maritima* (CoaX*_Tmar_*) (**Fig. 1E**) which were also resolved as a trimer of dimers (*32*).

To test CoaX*_Mtb_* binding of pantothenate and ATP, we performed thermal shift differential scanning fluorimetry (DSF). CoaX thermostability was unaffected by increasing concentrations of ATP alone (**Fig. 1F**, top). However, increasing pantothenate concentration had a subtle effect on CoaX thermostability (**Fig. 1F**, middle), while increasing concentrations of ATP in the presence of pantothenate had a larger effect (**Fig. 1F**, bottom), suggesting CoaX*_Mtb_* bound both ATP and pantothenate. We next tested if CoaX*_Mtb_* could hydrolyze ATP in the presence of pantothenate. In agreement with a previous study (*30*), we detected no activity for CoaX*_Mtb_* while CoaA*_Mtb_* had robust ATPase activity (**Fig. 1G**). Comparison of CoaX homologs revealed many conserved residues across diverse bacterial species. However, two amino acids (R8 and H229 in CoaX*_Mtb_*) showed a striking pattern of difference, mostly restricted to PPS- encoding actinobacteria (**Supplemental Fig. S3**). To test if these residues contribute to the lack of observed CoaX activity, we mutated both to glycine (“CoaX_DM_”), the most common amino acid found at this position in non-PPS-containing species (**Supplemental Fig. S3**). We found that CoaX_DM_ had measurable activity, albeit lower than CoaA (**Fig. 1G**).

In *Mtb*, *coaX* is the final gene in a presumed operon that includes three pantothenate synthesis genes: *panG* (Rv3603c), *panC* (Rv3602c), and *panD* (Rv3601c, **Fig. 2A**). *panB* (Rv2225) is encoded elsewhere, adjacent to a gene of unknown function, Rv2226, and *coaA* (Rv1092c) at a third locus (**Fig. 2A**). We deleted most of the coding sequence of *coaX* from *Mtb* and replaced it with a hygromycin resistance cassette (“Δ*coaX*”), maintaining the start codon and 198 bp of sequence at the 3’ end to minimize potential polar effects on downstream genes (**Fig. 2B**). Loss of *coaX* had no effect on growth (**Supplemental Figure S4 A**) or bacterial survival in the lungs or spleens of infected mice (**Supplemental Figure S4 B, C**).

**Fig. 2:**
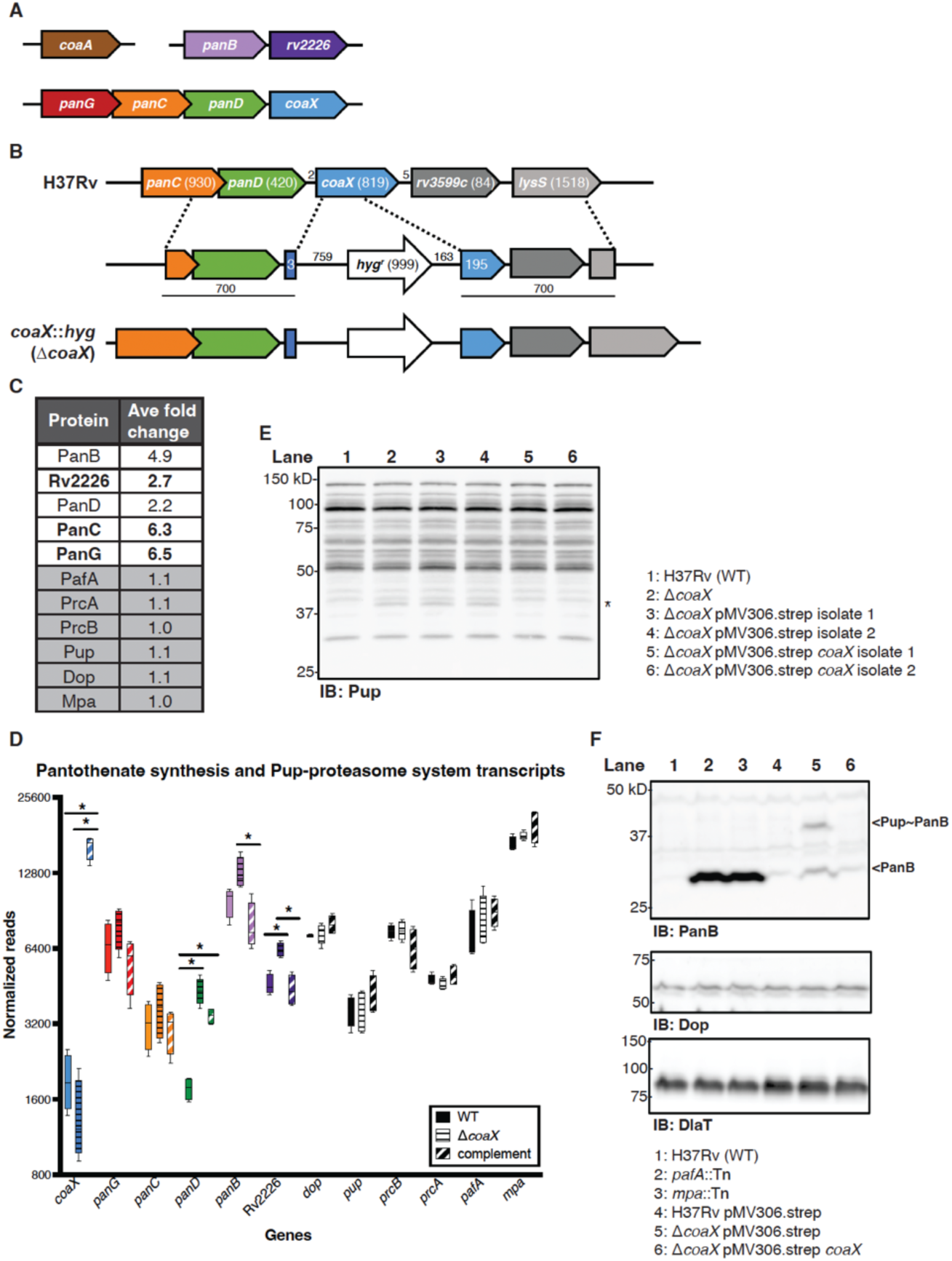
Loss of *coaX* leads to accumulation of Pup∼PanB. **(A)** Organization of three loci encoding pantothenate synthesis genes. **(B)** *coaX* was disrupted and replaced with a hygromycin selectable marker as described in the Materials and Methods (see **Supplementary Table S1** for plasmids and primers). **(C)** Average fold change of pantothenate synthesis (white boxes) and Pup-proteasome system (PPS, grey boxes) enzymes detected by data-independent acquisition mass spectrometry from WT and Δ*coaX* cell lysates. Proteins indicated in bold were significantly more abundant in the mutant than in WT bacteria (*P*<0.05). **(D)** Normalized reads associated with genes encoding pantothenate synthesis and PPS enzymes in RNA extracted from WT (solid bars), Δ*coaX* (horizontal black-striped bars), or a complemented strain (diagonal white striped bars). Statistically significant differences (*P*<0.1) are indicated with an asterisk. Pantothenate synthesis enzymes are colored corresponding to panel A. **(E)** Immunoblot against lysates extracted from *Mtb* strains indicated using monoclonal antibodies to Pup. A unique pupylated protein in the *coaX* mutant strains is indicated with an asterisk. **(F)** Immunoblot against lysates extracted from the indicated *Mtb* strains using polyclonal antibodies to PanB, Dop, or DlaT (dihydrolipoamide acyltransferase, loading control). Arrowheads indicate Pup∼PanB and PanB.

Data-independent mass spectrometry (DIA-MS) of Δ*coaX* and WT *Mtb* strains revealed all pantothenate synthesis enzymes and Rv2226 were increased more than two-fold in the mutant (**Fig. 2C**, **Supplemental Table S5**). No change in abundance of PPS enzymes was observed (**Fig. 2C**, **Supplemental Table S5**). RNA-sequencing (RNA-seq) analysis revealed loss of *coaX* led to increased transcripts associated with pantothenate synthesis genes and Rv2226, with no changes observed for PPS genes (**Fig. 2D**, **Supplemental Table S6**). Additionally, no proteins or gene expression changes were observed for coA synthesis enzymes (**Supplemental Tables S5 and S6**, respectively). Taken together, we concluded that CoaX contributes to the regulation of pantothenate synthesis enzymes.

Given the association between CoaX and Dop, we hypothesized CoaX might be functionally related to depupylation of one or more substrates. To test this hypothesis, we looked at the pupylome in lysates of WT, Δ*coaX*, and complemented strains. We observed a unique pupylated species in the Δ*coaX* strain that was not present in WT or complemented strains (**Fig. 2E)**. Among the proteins that were more abundant in mutant compared to WT bacteria as assessed by DIA-MS, PanB is the only confirmed member of the pupylome (*3, 8*). We raised polyclonal rabbit antiserum to *Mtb* PanB and detected robust accumulation of endogenous PanB in *pafA* and *mpa* mutants (**Fig. 2F, lanes 2 and 3, respectively**). Consistent with the proteomics data, the Δ*coaX* strain displayed a distinct accumulation of PanB compared to WT bacteria (**Fig. 2F, lane 1 compared to 5**). Strikingly, in addition to full length PanB, a higher molecular weight species was observed in the Δ*coaX* strain and lost upon complementation with *coaX* in single copy (**Fig. 2F, lane 5 compared to 6**). The size of this larger, anti-PanB-reactive species in the Δ*coaX* strain was consistent with the size of Pup∼PanB (*8, 38*).

Given the apparent ability of CoaX to bind pantothenate, we next tested if pantothenate might have an effect on PanB abundance in a CoaX-dependent manner. As a positive control, we also tested PanB levels in an *mpa* mutant, which accumulates PanB due to a lack of PPS- dependent degradation (*3*). Pantothenate supplementation of the cultures reduced PanB levels in WT and complemented strains (**Fig. 3A, lanes 1-3, 7-9, 10-12, 16-18**), but not in Δ*coaX* or *mpa*::*Tn* strains (**Fig. 3A, lanes 4-6 and 13-15, respectively**). FabD, another PPS substrate, and Dop levels were unaffected by pantothenate supplementation (**Fig. 3A**).

**Fig. 3.**
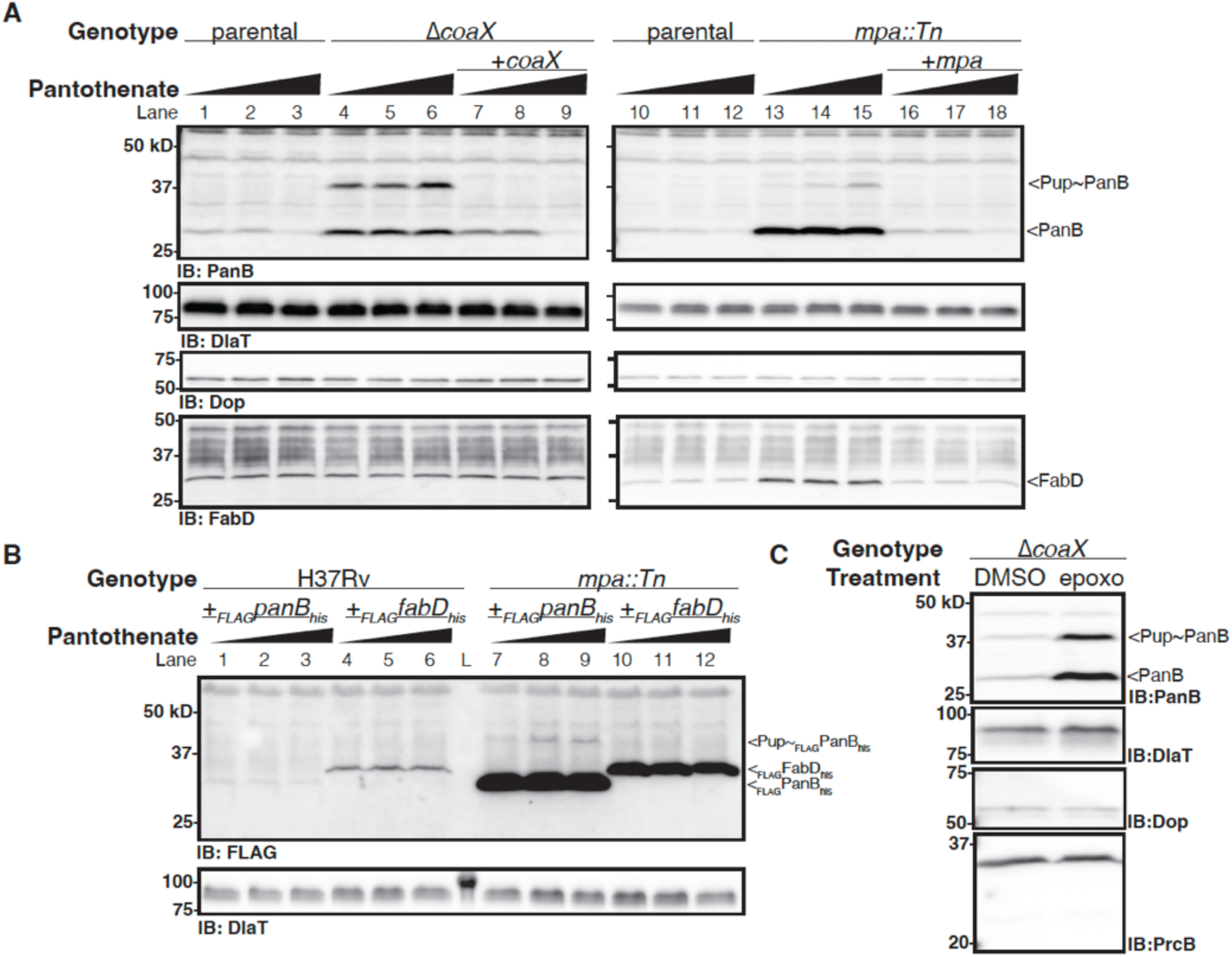
Pup∼PanB levels are regulated by pantothenate and the proteasome in *Mtb*. **(A)** Immunoblots (IB) of lysates extracted from Δ*coaX*, *mpa*::Tn, complemented strains, and parental controls supplemented with 0, 10, or 100 µM pantothenate are shown. Parental controls were transformed with empty vectors used for complementation constructs (see **Table S1** for more information). IBs using DlaT or PanB antisera were performed on the same membrane. Endogenous Pup∼PanB and PanB are indicated by arrowheads. **(B)** IBs against lysates of H37Rv (WT) or *mpa*::Tn strains ectopically producing _FLAG_PanB_his_ or _FLAG_FabD_his_ are shown. IBs for DlaT were used as loading controls (bottom). FLAG-tagged constructs are indicated. “L” indicates a lane containing protein MW standards. **(C)** IBs against lysates of the Δ*coaX* strain treated for 24 hours with DMSO (vehicle control) or 50 µM epoxomicin (“epoxo”), an irreversible proteasome inhibitor, are shown. IBs using DlaT or PanB antisera were performed on the same membrane. IBs using Dop or PrcB antisera were performed on the same membrane.

We observed no other changes in the pupylome with pantothenate supplementation (**Supplemental Figure S5**). To distinguish between transcriptional and post-translational regulation of PanB levels in response to pantothenate, we tested the effects of pantothenate supplementation on strains expressing _FLAG_PanB_his_ or _FLAG_FabD_his_ ectopically and under the control of a non-native (*hsp60*) promoter (*3*). While _FLAG_PanB_his_ levels were almost undetectable in WT *Mtb*, Pup∼ _FLAG_PanB_his_ accumulated in the *mpa* strain in a dose-dependent manner with pantothenate supplementation (**Fig. 3B, lanes 1-3, 7-9**). Pup∼_FLAG_FabD_his_ was undetectable under any condition, as observed for endogenous FabD, which was unchanged in response to pantothenate (**Fig 3B, lanes 4-6, 10-12**). While pupylation alone is insufficient for rapid proteasomal turnover of all substrates (*7*), treatment of the Δ*coaX* strain with the proteasome inhibitor epoxomicin demonstrated a marked accumulation of endogenous PanB and Pup∼PanB (**Fig. 3C**) indicating that Pup∼PanB was turned over by proteasomes in the Δ*coaX* strain. Levels of Dop and PrcB, the catalytic subunit of the proteasome, were unaffected by epoxomicin treatment. Taken together, our data suggest PanB levels are post-translationally regulated by the PPS in response to pantothenate levels.

To test if CoaX and pantothenate are sufficient for modulating depupylation of PanB, we reconstituted depupylation *in vitro*. As shown previously, Dop robustly depupylated Pup∼PanB *in vitro* (**Fig. 4a**) (*18–20, 39, 40*). We found addition of CoaX significantly accelerated depupylation of Pup∼PanB but not Pup∼FabD (**Fig. 4A, Supplemental Fig. S6A, C**). Addition of pantothenate to depupylation reactions containing CoaX significantly inhibited depupylation of Pup∼PanB but not Pup∼FabD (**Fig. 4B, and Supplemental Fig. S6B, D**). Pantothenate supplementation in the absence of CoaX or Dop had no effect on Pup∼PanB levels (**Supplemental Fig. S6E**). Collectively, our data support a model where CoaX specifically facilitates depupylation of Pup∼PanB, a process that is inhibited by pantothenate.

**Fig. 4:**
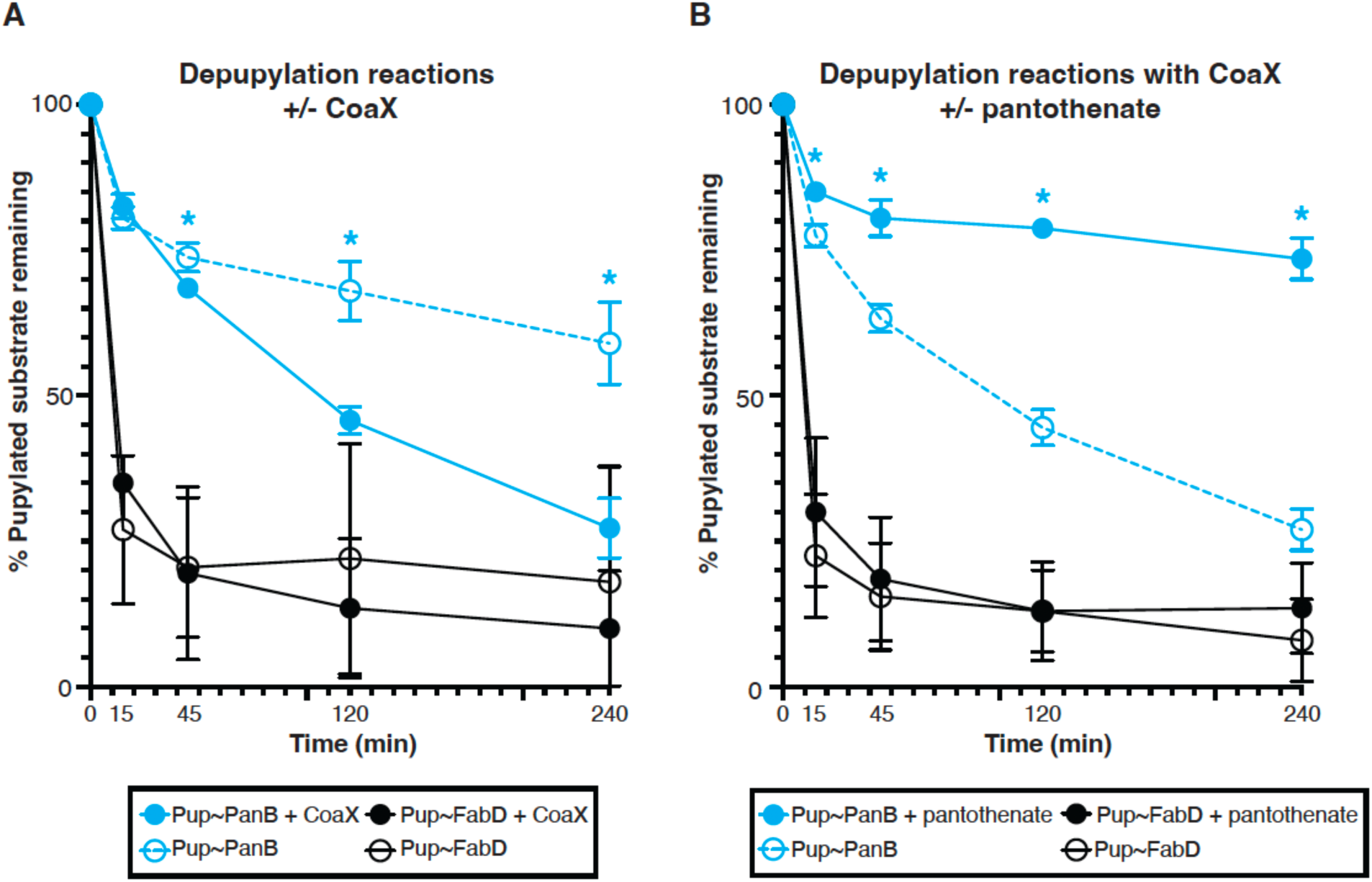
Pup∼PanB depupylation is accelerated by CoaX and inhibited by pantothenate in a CoaX-dependent manner. **(A)** Percent Pup∼PanB (cyan) or Pup∼FabD (black) remaining after 0, 15, 45, 120, and 240 min of depupylation in the presence (filled circles) or absence (open circles) of CoaX are graphed. **(B)** Percent Pup∼PanB (cyan) or Pup∼FabD (black) remaining after 0, 15, 45, 120, and 240 min of depupylation reactions containing CoaX in the presence (filled triangles) or absence (open triangles) of 1 mM pantothenate are graphed. Statistically significant time points (*P*<0.05) are indicated with asterisks.

## Discussion

In this work, we show that CoaX and pantothenate work together to regulate the abundance of PanB, an essential enzyme that catalyzes the first step of pantothenate biosynthesis (**Fig. 5A-C**). This work is the first report of a protein or a metabolite that regulates the pupylation status of a specific PPS substrate.

**Fig. 5.**
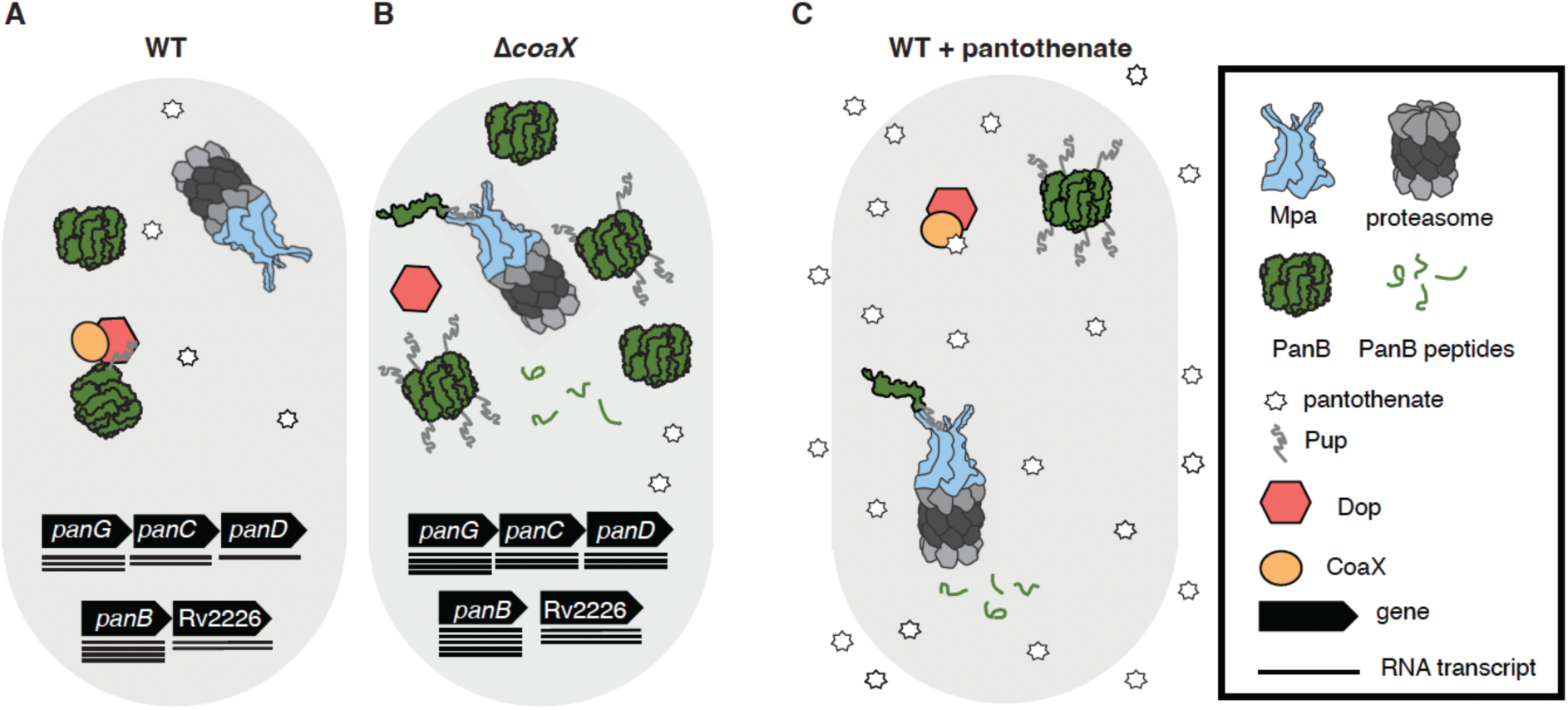
Model of PanB regulation by CoaX, the Pup-proteasome system, and pantothenate. (A) Under standard culture conditions in WT *Mtb*, PanB (green, homodecamers) levels are low, and depupylated by Dop (red) along with CoaX (orange). Functional PanB decamers synthesize pantothenate (white stars). **(B)** Loss of CoaX regulated depupylation in the Δ*coaX Mtb* leads to increased pupylated (grey lines) PanB and turnover by Mpa (light blue, homohexamers) and the proteasome (grey scale, hetero-28-mers; proteasome processed PanB peptides shown as green lines). Additionally, RNA transcripts (black lines) associated with pantothenate synthesis genes (black arrows), including *panB*, are elevated in this strain compared to WT bacteria by an unknown mechanism. **(C)** Upon pantothenate supplementation, PanB depupylation decreases, Pup∼PanB increases, and more PanB is degraded by Mpa and the proteasome leading to lower intracellular PanB levels.

Our data support a model in which CoaX accelerates depupylation of Pup∼PanB in the absence of supplemental pantothenate. In the absence of CoaX, less PanB would be depupylated, leading to more turnover by the proteasome. While we expected that increased PanB degradation would be lethal, RNA-seq analysis showed transcripts associated with pantothenate synthesis increased in a Δ*coaX* strain (**Figure 2D**). To our knowledge, nothing is known about how the *pan* loci are transcriptionally regulated in mycobacteria. As there is no evidence to suggest CoaX plays a direct role in *pan* gene transcription or RNA half-life, other regulators likely exist. Regardless of mechanism, we hypothesize this change in transcript abundance leads to higher PanB protein levels to compensate for increased proteasomal turnover of PanB and maintain pantothenate synthesis levels required for survival.

Given the importance of coA and its derivatives in numerous cellular processes, the existence of multiple regulatory mechanisms to control pantothenate synthesis is perhaps unsurprising. Relatedly, previous metabolomics analyses performed on an *mpa* mutant showed pantothenate and other related metabolite levels are unchanged in the absence of PPS-dependent degradation of PanB (*41*). Thus, our model has some parallels to the ubiquitin-proteasome system-dependent regulation of cholesterol metabolism by 3-hydroxy-3-methylglutaryl coenzyme A (HMGcoA) reductase in eukaryotes, where this enzyme is ubiquitylated and degraded in response to sterol levels (*42*).

Many outstanding molecular and biological questions remain. Structurally, a greater understanding of CoaX, Dop, PanB, and pantothenate interactions would help clarify how CoaX and pantothenate affect PanB depupylation. PanB is active as a homodecamer of two, stacked, pentameric rings held together by interactions between the C-termini of monomers in each ring (*43*). *In vitro*, PanB is pupylated at a single surface exposed lysine that would extend from the top or bottom of the decameric complex (*38*). How CoaX and Dop interact with the pupylated surfaces of the PanB decamer could yield insights into how Dop activity might be modulated by other regulators. Additionally, understanding the stoichiometry of proteins involved could resolve outstanding questions regarding the relevance of higher order CoaX complexes that have been structurally observed.

Physiologically, it is unknown when over the course of infection pantothenate might be exogenously available, how it is acquired by *Mtb*, or when synthesis is most important for survival and virulence. Rapid proteasomal turnover of PanB at steady state may underlie a significant cost to pantothenate production. The existence of an apparently specific depupylation regulator may indicate that rapid induction of pantothenate synthesis is key in some contexts. Furthermore, the differential conservation of CoaX residues in proteasome containing species compared to proteasome lacking species may indicate regulation of PanB by CoaX, Dop, pantothenate, and the proteasome is a widespread phenomenon.

Both pantothenate synthesis and PPS enzymes have previously been investigated as potential drug targets [reviewed recently in (*44*) and (*45*)]. Focus on pantothenate biosynthesis has been motivated by the fact that humans do not make pantothenate, and pantothenate synthesis enzymes are essential in *Mtb* under most laboratory conditions (*24*). Similarly, loss of any of the PPS enzymes yields dramatic virulence defects in mice infected with *Mtb* (*2–6*). A greater understanding of how pantothenate synthesis is regulated, including by the PPS, may offer new approaches for the successful development of antimicrobials targeting this pathway. Additionally, identification of PPS regulators, and general principles for the regulation of individual substrates, has the potential to reveal new targets and pathways for drug development.

## Acknowledgements

We thank A. Darwin for reading a draft version of this manuscript and S. Zhang for making the TAP-tagged plasmids. We thank J. Ilmain for assistance with SEC and negative stain EM. We thank the Office of Science & Research High-Containment Laboratories at NYU Grossman School of Medicine for their support in the completion of this research. We thank W. Rice, B. Wang, H. Juang for assistance with cryo-EM grid screening and microscope operation. EM data processing used computing resources at the HPC Facility at NYU, and we thank the members of the HPC team for high-performance computing support.

## Funding

KHD is funded by NIH grant AI088075. SCK was supported in part by a Public Health Service Institutional Research Training Award T32 AI007180. JC is a fellow of The Jane Coffin Childs Memorial Fund for Medical Research. This investigation has been aided by a grant from The Jane Coffin Childs Memorial Fund for Medical Research. GB and DE were funded by NIH grant AI174646. LA and LMI are supported by the National Library of Medicine Intramural Research Program at the National Institutes of Health, USA. Some of the work was performed at NYU Langone Health’s Cryo-Electron Microscopy Laboratory (RRID: SCR_019202), which is partially supported by the Laura and Isaac Perlmutter Cancer Center Support Grant NIH/NCI P30CA016087. KYR is supported by NIAID P01 AI143575 and BMGF INV-204250.

## Author contributions

SCK and KHD conceived of the study. SCK made and complemented the Δ*coaX* strain, prepared samples for DIA-MS and RNA-seq, and performed mouse infections, pantothenate supplementation experiments, growth curves, and depupylation assays. JHY and SCK produced protein for PanB anti-sera development. JHY performed Dop and PafA immunoprecipitations for mass spectrometry. JHY purified CoaX, JHY and JC prepared grids for cryo-EM and JC, GB, and DCE conducted cryo-EM analysis. KN conducted DSF and kinase activity assays. SCK, LMI, and LA performed evolutionary analysis of PanK sequences. LMI and LA modeled CoaX*_Mtb_* in complex with pantothenate and ADP. SCK performed RNA-Seq analysis with work-flow developed and customized by GP, NMS, and AP. SCK, JHY, JC, KN, LMI, LA, GB, DCE, KYR, and HD contributed to data analysis. SCK, JC, GB, DCE, and HD wrote the manuscript. All authors read and approved of the manuscript.

## Competing interests

The authors declare no competing interests.

## Data and materials availability

Correspondence and requests for materials should be addressed to Heran Darwin. The cryo-EM maps have been deposited in the Electron Microscopy Data Bank with accession codes: CoaX*_Mtb_* hexamer (EMD-44303), and CoaX*_Mtb_* tetramer (EMD-44304). The coordinates of the atomic models have been deposited in the Protein Data Bank under accession codes: CoaX*_Mtb_* hexamer (PDB 9B78), and CoaX*_Mtb_* tetramer (PDB 9B79).

## Materials and Methods

Except where otherwise indicated, all reagents were purchased from ThermoFisher Scientific.

### Cloning and bacterial transformation

All constructs in this study were cloned by polymerase chain reaction (PCR) and restriction digestion and inserts were verified by Sanger sequencing (GENEWIZ, Inc). All primers were purchased from Integrated DNA Technologies. Restriction enzymes, quick calf intestinal phosphatase, and T4 DNA ligase were purchased from New England Biolabs (NEB). PCR was performed with Taq (Qiagen) or Phusion (NEB) polymerases according to manufacturer’s instructions. *Mtb* DNA extraction was performed as previously described (*46*).

As necessary, QIAquick Gel extraction or PCR purification Kits (Qiagen) were used. Verified plasmids were extracted from *Escherichia coli* (*E. coli*) using the QIAprep Spin Miniprep Kit or HiSpeed Plasmid Midi Kit (Qiagen) and dialyzed (0.025µM MCE Membrane, Millipore) prior to transformation. Chemically competent DH5α and ER2566 *E. coli* were prepared and transformed as described previously (*47*). Chemically competent *Msm* and *Mtb* were prepared and transformed as described previously (*48*). For a list of all strains used in this work see Supplementary Table S1.

To generate pET24b(+)-*coaX*, primers Rv3600c_H3_R and NdeI_Rv3600cF were used to amplify *coaX* from *Mtb* gDNA. This insert was digested with NdeI and HindIII for ligation into pET24b(+). pET24b(+)-*coaX* served as the template for PCR with Rv3600c_H3_R and KpnI_Rv3600c_F to generate an insert for pETDUET-_his6_*pup*_91_-*coaX*-*dop_Msm_*_C438S_. The *coaX* insert was digested with HindIII and partially digested with KpnI before ligation into pETDUET- _his6_*pup*_91_-*dop_Msm_*_C438S_ digested with the same enzymes. For inducible protein production, pET24b(+)-*coaX* was transformed into ER2566 *E. coli* and pETDUET-_his6_*pup*_91_-*coaX*- *dop_Msm_*_C438S_ was transformed into ER2566 with p*groESL* to improve Dop solubility as in (*49*).

pET28a-*coaA_his_* and pET28a-*coaX_his_* were made by cloning *coaA* or *coaX*, codon optimized for expression in *E. coli*, into pET28a-His6 using NdeI and BamHI by GenScript (NJ, USA). pET28a-*coaX*_DM_his_ was synthesized similarly by Twist Bioscience (CA, USA) however codons corresponding to residues 8 and 229 were altered to be translated as glycines. For inducible protein production, all pET28a based plasmids were transformed into BL21(DE3) *E. coli*.

To generate pET24b(+)-*_FLAG_panB_his_*, *_FLAG_panB_his_* was amplified from pMN402-*_FLAG_panB_his_* using primers FLAG-panB-Nde-f2 and panB-Not-r2. The insert was digested with NotI and NdeI and inserted into pET24b(+) digested with the same enzymes. To generate pOLYG-*_Myc_pup*-*_FLAG_panB_his_*, pET24b(+)-*_FLAG_panB_his_* was amplified with primers pET24bF-PstI and pet24bR-SpeI, digested with SpeI and PstI and ligated into pOLYG-*_Myc_pup_E_-_FLAG_fabD_his_* digested with the same enzymes to replace _FLAG_*fabD_his_* with *_FLAG_panB_his_*. pET24b(+)-*_FLAG_panB_his_* was transformed into ER2566 for inducible protein production and pOLYG-*_Myc_pup*-*_FLAG_panB_his_* was transformed into MsHD336 to generate MsHD369.

*dop_TAP_ and pafA_TAP_* were amplified from *Mtb* gDNA using primer pairs HindIIIdopfor/BamHIdopTAPrev, and HindIIIpafAfor/BamHIpafATAPr, respectively. Inserts were cloned into pOLYG using HindIII and BamHI. pOLYG-*dop_TAP_* and pOLYG-*pafA_TAP_* were transformed into MHD58 to generate MHD1097 and 1187, respectively.

Deletion-disruption of *coaX* with a hygromycin resistance cassette was performed by allelic exchange as previously described (*50*). To generate an allelic exchange vector, 700 bp upstream of *coaX* (the “upstream flank”), including the start codon, was amplified *Mtb* gDNA using XbaI_CoaXupstreamF and StuI_CoaXupstreamR. The upstream flank was digested with XbaI and StuI and cloned into pYUB854. To minimize disruption of Rv3599c, which begins 5bp after *coaX*, a 700 bp fragment beginning 200bp upstream of the Rv3599c start codon was amplified from *Mtb* gDNA using BstBI_CoaXdownstreamF and HindIII_CoaXdownstreamR (the “downstream flank”). The downstream flank was digested with BstBI and HindIII-HF and cloned into the pYUB854 vector already containing the upstream flank to generate pYUB854- *coaX*::*hyg*. Successful allelic exchange thus preserves the *coaX* start codon followed by 1921bp of intervening sequence, including a hygromycin resistance cassette, followed by 198 bp of sequence upstream of the *coaX* stop codon, which may contain regulatory sequences for Rv3599c. pYUB854-*coaX*::*hyg* was transformed into MHD1 to generate MHD1824. Deletion and replacement of the *coaX* locus with a hygromycin resistance cassette in MHD1824 was verified by DNA extraction and PCR using coaX_Hyg_externalF and coaX_Hyg_externalR.

Because *coaX* appeared to be transcribed in an operon beginning with *panG*, based on read pileups generated from previous RNA-seq experiments in our lab (*51*), we designed a complementation vector for the Δ*coaX* strain using 200bp upstream of *panG* as the promoter sequence. The promoter was amplified from *Mtb* gDNA using Rv3603cp_R and Rv3603cp_F, *coaX* was amplified with Rv3600c_H3_R and coaXcodon2_F, and the two fragments fused together by overlap extension (*52*) PCR using Rv3603cp_F and Rv3600c_H3_R primers. This insert was cloned into pMV306.strep by HindIII and NheI digest to generate pMV306.strep- *coaX*. pMV306.strep-*coaX* was transformed into MHD1824 to complement the loss of *coaX* and generate MHD1846. MHD1824 was also transformed with pMV306.strep as a control strain against which complementation alone could be isolated from other effects of plasmid transformation.

### Bacterial culture conditions

*Mtb* and *Msm* were grown in Middlebrook 7H9 liquid broth (BD Difco) supplemented with 0.2% glycerol (Research Products International), 0.05% Tween-80, 0.5% bovine serum albumin fraction V (“BSA”, MilliporeSigma), 0.2% dextrose (VWR Chemicals BDH), and 0.085% sodium chloride (“7H9c”). For growth on agar, we used Middlebrook 7H11 (BD Difco) supplemented with 0.5% glycerol and 10% BBL Middlebrook Oleic Albumin Dextrose Catalase Growth Supplement (BD Difco). For growth curves in minimal media *Mtb* were grown to mid log, washed 3x in 1x DPBS (Corning) with 0.05% Tween-80 (“PBS-T”), and resuspended at OD 0.05 in Sauton’s (3.7 mM potassium phosphate, monobasic; 2.4 mM magnesium sulfate heptahydrate; 30 mM L-(+)-asparagine monohydrate; 3.5 mM zinc sulfate; 9.5 mM citric acid trisodium salt dihyrate (Acros Organics); 6.0% glycerol; 0.005% ferric ammonium citrate (MP Biochemicals); 0.05% Tween-80) or Proscauer-Beck (0.5% potassium phosphate monobasic, 0.06% magnesium sulfate heptahydrate, 1.5% glycerol, 0.25% magnesium citrate, dibasic anhydrous (Chem-Impex International), 0.05% Tween 80, and 10 mM L-(+)-asparagine monohydrate) media. Antibiotics were used as follows for mycobacteria: kanamycin sulfate 50 µg/ml, hygromycin B (Corning) 50 µg/ml, or streptomycin sulfate (Acros) 25 µg/ml. For pantothenate supplementation, D-Calcium pantothenate was dissolved at 100 µM in water, syringe filter sterilized (Corning, 0.45 µM, nylon) and diluted into 7H9c.

All *Mtb* cultures were grown static at 37°C and OD measured at 580 nm. Bacteria harvested for immunoblotting or mass spectrometry were washed with PBS-T prior to lysis.

*E. coli* was cultured in Luria-Bertani broth (BD Difco) or on Luria-Bertani agar (Research Products International). For *E. coli*, selection was performed with: 100 µg/ml kanamycin, 150 µg/ml hygromycin, 50 µg/ml streptomycin sulfate, or 200 µg/ml ampicillin sodium salt. *E. coli* was grown at 37°C with shaking and growth monitored by absorbance at 600 nm. For collection of pellets for preparative purification, the *dop_Msm_* expression strain was grown to mid log (OD 0.4-1.0), induced with 0.6 mM Isopropyl ß-D-1-thiogalactopyranoside (IPTG, IBI Scientific), and grown for another 4-6 hours. The *coaX* expression strain was grown and collected similarly but induced with 1 mM IPTG.

### Lysate extraction

*Mtb* lysates were extracted in 50 mM Tris-Cl pH8, 1 mM EDTA pH8 by bead beating, as previously described (*21*). Lysates extracted for immunoprecipitation were syringe filtered with 0.45 µM nylon filters. Lysate extracted for mass spectrometry was filtered with SpinX Centrifugal tube filters (costar). *E. coli* lysates were extracted by sonication (1 second on, 1 second off, 30 seconds total, 3 min on ice in between).

All samples analyzed by Sodium dodecyl-sulfate polyacrylamide gel electrophoresis (“SDS-PAGE”) were mixed with 4X Sample Buffer (“4X SB”: 250 mM Tris pH 6.8, 2% Dodecyl sulfate sodium salt (Acros Organics), 20% β-mercaptoethanol, 40% glycerol, 1% bromophenol blue (G-Biosciences)). Polyacrylamide gels were made as previously described (*53*).

### Immunoblotting

Lysates were analyzed by SDS-PAGE, soaked in transfer buffer (25 mM Tris-Base, 192 mM glycine, 20% methanol) for 1 hour, and transferred to Amersham^TM^ Protran 0.2µM nitrocellulose (GE Healthcare) by semi-dry transfer (15V, 15 min). PonceauS solution (Sigma-Aldrich) was used to ensure equal loading and transfer of all blots. Immunoblots were performed with monoclonal antibodies to Pup (*4, 21*) or polyclonal antibodies to Dop (*11*), DlaT (*54*), PrcB (*15*), or FabD (*7*) as previously described. Monoclonal ANTI-FLAG M2 antibody (Sigma Aldrich) was used according to manufacturer’s instructions. Blocking and antibody solutions were prepared in 25 mM Tris-Cl, 125 mM NaCl, and 0.05% Tween-20 (“1X TBS-T”). All blots were blocked rocking at RT for at least an hour, incubated in primary antibody overnight rocking at 4C, washed three times with TBS-T, incubated in secondary antibody rocking for at least one hour at RT, and washed again three times with TBS-T. With the exception of Pup and FLAG antibodies, all blocking and antibody solutions contained 3% BSA in 1X TSB-T. All Pup IBs solutions contained 2% milk and were diluted in 0.1X TBS-T. FLAG IB solutions contained 3% milk in 1X TBS-T. Secondary horseradish peroxidase (HRP)-coupled anti-mouse or anti-rabbit antibodies were used according to manufacturer’s instructions (Pierce). Immunoblots were developed using SuperSignal West Pico PLUS and imaged on ChemiDoc Touch Imaging system (Bio-Rad).

For polyclonal antibody production, _FLAG_PanB_his_ was purified from *E. coli* under native conditions as described above except with a different sonication protocol: 10 seconds on and 10 seconds off (65% amplitude), for 1 min. Lysate was collected and NiNTA purification performed as described above. Elutions were pooled and dialyzed overnight at 4°C in Dulbecco’s Phosphate Buffered Saline (PBS, Corning) using a 3500 MWCO Slide-A-Lyzer® Dialysis Cassette. Protein was concentrated with Amicon® Ultracel ® 3K filters (Millipore) for immunization of rabbits by Covance (Denver, PA).

### Protein purification and purification for depupylation assays

Dop purification for depupylation assays was performed as described previously (*49*). In brief, N-terminally 6xhis tagged N-terminally truncated Pup (“_his6_Pup_91_”) was co-produced in *E. coli* with Dop*_Msm_* and GroESL*_Ecoli_*. Untagged Dop_Msm_ was purified by interaction with _his_Pup_91_ using Ni-NTA Agarose (Qiagen) as previously published (*49*) and further purified by size exclusion chromatography (SEC) over a Superose^TM^ 6 10/300 GL column using an ÄKTA pure protein purification system (Citivia). As untagged CoaX has an intrinsic affinity for Ni-NTA Agarose, it was purified using NiNTA agarose and SEC in the same manner. SEC fractions of each protein purification were concentrated and buffer exchanged with Amicon Ultra 10 kDa MWCO Centrifugal Filters (MilliporeSigma) according to manufacturer’s instructions.

_Myc_Pup∼_FLAG_PanB_his_, and _Myc_Pup∼_FLAG_FabD_his_ were purified from proteasome-lacking *Msm* for depupylation assay as previously described (*55*). _Myc_Pup∼_FLAG_PanB_his_ was purified in complex with _FLAG_PanB_his_ due to apparent heterogeneous pupylation of the PanB decamer (*43*).

All proteins purified for use in depupylation assays were stored in 50mM NaPO4 pH 8, 10mM MgCl2, 150mM NaCl, and 20% glycerol and flash frozen with an ethanol-dry ice bath for storage at −80°C. Protein concentrations were quantified by Bradford assay using Bio-Rad Protein Assay Dye Reagent Concentrate (Bio-Rad) according to manufacturer’s instructions.

CoaA_his_, CoaX_his_, and CoaX_DM_his_ were produced for purification by transforming corresponding pET28a based plasmids into BL21(DE3) and selected for kanamycin resistance overnight. Single colonies were inoculated into 10mL LB media supplemented with kanamycin, grown ON, and subcultured into 1L LB broth the following morning. Upon reaching OD_600_ 0.5- 0.7, they were cold shocked and induced with 1mM IPTG ON at 18°C. The cell pellet was obtained by centrifugation for 45 min @ 10,000 x g and resuspended in 50ml lysis buffer (50mM Tris/HCl, pH 8.0, 300mM NaCl, 10% glycerol, 25mM Imidazole). Cells were lysed by French Press Cell Disruptor, with icing between cycles. Soluble lysate was loaded onto a Ni-NTA column, equilibrated with lysis buffer. Flow through was collected and reloaded on the column before washing. Proteins were eluted with 300mM Imidazole. Gravity PD-10 Desalting column (Cytiva) was used to clean up purified proteins according to manufacturer’s instructions.

### Cryo-EM sample preparation

ER2566 containing *pgroESL* transformed with pETDUET-*_his_pup_91_*-*coaX*-*dop_Msm_*_C438S_ was grown to OD ∼0.5 and production of recombinant proteins induced with 0.6 mM IPTG for 4 hours shaking at 37°C. Cells were collected by centrifugation and frozen at −20°C. Lysate was extracted by sonication and NiNTA purification was performed as published previously for Dop*_Msm_*_C438S_ and _his6_Pup_91_ (*49*) followed by SEC as described for proteins used in depupylation assays above. Fractions containing the most protein, as determined by chromatogram, were collected for imaging.

SEC fractions were screened by negative-stain electron microscopy to assess sample quality and homogeneity. To prepare grids for negative stain electron microscopy, protein sample was applied on a freshly glow discharged carbon coated 400 mesh copper grid (Ted Pella Inc., cat. #01754-F) and subsequently blotted off on filter paper. Immediately after blotting off the sample, a 0.75% uranyl formate solution was applied for staining and followed by blotting. Application and blotting of stain was repeated five times. Grids were dried before imaging on a Talos L120C TEM (FEI) equipped with a 4K x 4K OneView camera (Gatan).

Fractions of interest concentrated to ∼0.01 mg/mL in cryo-EM buffer (50 mM Tris-HCl pH 8.0, 150 mM NaCl, and 1 mM DTT). Continuous carbon grids (Quantifoil R 2/2 on Cu 300 mesh grids + 2 nm Carbon, Quantifoil Micro Tools C2-C16nCu30-01) were glow-discharged for 5 sec in an easiGlow Glow Discharge Cleaning System (Ted Pella Inc.). 3.5 µL sample was added to the glow-discharged grid. Using a Vitrobot Mark IV, grids were blotted for 3 seconds at 22 °C with 100% chamber humidity and plunge-frozen into liquid ethane. Grids were clipped for screening.

### Cryo-EM screening and data collection

Clipped cryo-EM grids were screened at NYU Cryo-EM Laboratory on a Talos Arctica operated at 200 kV equipped with a K3 camera (Gatan). Images of the grids were collected using Leginon v3.6 (*56*) at a nominal magnification of 36,000x (corresponding to a pixel size of 1.096 Å) with total dose of ∼49 e- per Å^2^, over a defocus range of −2.0 to −3.0 µm. Grids were selected for data collection based on ice quality and particle distribution. Selected cryo-EM grids were imaged at NYU Cryo-EM Laboratory on a Krios-3Gi operated at 300 kV with a K3 camera (Gatan) and an energy filter slit width of 20 eV. Super-resolution movies were collected using Leginon v3.6 at a nominal magnification of 105,000x, corresponding to a super-resolution pixel size of 0.41275 Å (or a nominal pixel size of 0.8255 Å after binning by 2). Movies were collected at a dose rate of 26.95 e-/Å^2^/s with a total exposure of 1.80 seconds, for an accumulated dose of 48.51 e-/Å^2^. Intermediate frames were recorded every 0.05 seconds for a total of 40 frames per micrograph. A total of 7913 images were collected at a nominal defocus range of 0.8 – 2.5 µm.

### Cryo-EM data processing

The Krios dataset was split into 8 batches of 1,000 movies and processed in cryoSPARC v4.31 (*57*), as described in Supplemental Figure S1 to maximize the use of computational resources. Dose-fractionated movies were gain-normalized, drift-corrected, summed, and dose-weighted using the cryoSPARC Patch Motion module (Max resolution alignment: 3 Å, Output F-crop factor: 1/2). The contrast transfer function was estimated for each summed image using cryoSPARC Patch CTF (Amplitude Contrast: 0.07, Maximum resolution: 1.64 Å, Maximum search defocus: 50,000 Å).

Micrographs from each batch were picked using cryoSPARC Blob Picker (Min particle diameter: 100 Å, Max particle diameter: 200 Å, Max # of local maxima to consider: 1000) and extracted using Extract From Micrographs module (Extraction box size: 256 px, Fourier crop to box size: 64 px). Extracted particles were sorted using 2D Classification (N= # of classes = 50, Remove duplicates particles). Particles from ‘junk’ classes where selected and processed using cryoSPARC Ab initio Reconstruction (Ab-initio Reconstruction (N = 3, # of particles to use: 100,000) to generate three decoy models (Decoy 1, Decoy 2, and Decoy 3) for downstream particle curation using cryoSPARC Heterogenous Refinement. Particles from well-defined classes were selected from each batch, combined (1,742,997 particles total) and sorted by another round of 2D Classification (N = 50). Particles from well-resolved 2D classes (1,194,966 particles) were selected and re-extracted using Extract From Micrographs module (Extraction box size: 256 px, no binning).

Extracted particles were processed using cryoSPARC Ab initio Reconstruction (N = 3, # of particles to use: 100,000, Max res: 7 Å, Initial resolution: 9 Å) to generate initial 3D models (Model 1 (22,953 particles), Model 2 (19,586 particles), Model 3 (57,461 particles)). Extracted particles were further curated with five rounds of Heterogeneous Refinement (N = 6, templates = (1) Model 1, (2) Model 2, (3) Model 3, (4) Decoy 1, (5) Decoy 2, (6) Decoy 3), in which particles sorted into Models 1, 2, and 3 were used as input for the next round. After multiple rounds of Heterogeneous refinement (round 1: 990,417 particles, round 2: 899,837 particles, round 3: 838,692 particles, round 4: 798,094 particles, round 5: 774,783 particles), curated particles were classified using Heterogenous Refinement (N = 3, templates = (1) Model 3, (2) Model 3, (3) Model 3), revealing two distinct classes of Mtb CoaX: hexamer (Ref 1) and tetramer (Ref 2), which were used as references for downstream particle curation.

To sample more views of the hexamer and tetramer, particles from Models 1, 2, and 3 were each sorted by 2D classification (N = 50) and particles from 2D classes containing unique views selected. Particles from the 5 different views were separately trained within the Topaz Train module (*58*) in cryoSPARC (Expected # of particles: 1000, Model architecture: ResNet16).

After training, particles were picked using the trained Topaz model and extracted (Extraction box size: 256 px, Fourier crop to box size: 64 px). Particles from the five separate Topaz pickings were aligned using 2D Classification (N = 50), combined, and duplicate particles picks were removed. Particles were re-extracted using Extract From Micrographs module (Extraction box size: 256 px, no binning), resulting in 8,929,726 particles.

Extracted particles were curated by running four rounds of Heterogeneous Refinement (N = 5, templates = (1) Ref 1, (2) Ref 2, (3) Decoy 1, (4) Decoy 2, (5) Decoy 3), in which particles sorted into Ref 1 and Ref 2 were used as input for the next round. After multiple rounds of Heterogeneous refinement (round 1: 4,653,022 particles, round 2: 3,412,231 particles, round 3: 2,347,502 particles, round 4: 2,154,528 particles) and removing remaining duplicates (alignment3D), the 1,285,898 curated particles in the hexamer class (Ref 1) and the 771,380 curated particles in the tetramer class (Ref 2) were refined using cryoSPARC Non-Uniform Refinement (*59*) (Initial lowpass resolution: 15 Å) generating density maps with average resolutions of 2.59 Å-resolution and 2.81 Å, respectively.

### Model building and refinement

For the initial model of CoaX*_Mtb_*, we used AlphaFold2 (*60*) to generate a monomer of CoaX through ColabFold (*61*). The predicted CoaX monomers were manually fit as rigid bodies into the cryo-EM maps of the CoaX hexamer and tetramer using ChimeraX (*62*). CoaX hexamer and tetramer models were further refined using real-space refinement in PHENIX v1.20.19 (*63*). The refined models were manually inspected in COOT v0.8.9.2 (*64*) to assess the overall fit for the Ca backbone and side chains of each CoaX protomer into the maps. These models were iteratively inspected, manually rebuilt in COOT, and refined in PHENIX until completion. Final model refinements were performed using PHENIX with global minimization, Ramachandran restraints, secondary structure restraints, and nonbond weight set at 200.0. The final model for the CoaX hexamer is nearly complete apart from the C-terminal regions of each CoaX promoter (residues 261-272). The final model for the CoaX tetramer is also nearly complete, aside from chain A (residues 165-172), chain A (residues 166-172), and the C-terminal regions of each CoaX promoter (residues 261-272). These residues were not modeled since they were not well-resolved in the cryo-EM maps.

Quantification and statistical analyses for model refinement and validation on deposited models were extracted from the results of the real_space_refine algorithm in PHENIX (*63*) as well as MolProbity (*65*) and EMringer (*66*). Structural alignments and associated RMSD values were calculated using PyMOL (Schröodinger, LLC). The local resolution of the cryo-EM maps was estimated using cryoSPARC Local Resolution (*57*). FSCs that were calculated in cryoSPARC were plotted in Prism v10 (GraphPad). Directional 3DFSCs were calculated using 3DFSC2 (*67*). Figures were generated with PyMOL (Schröodinger, LLC) and ChimeraX (*62*).

### Mass spectrometry to identify Dop binding partners and characterize the proteome of WT and Δ*coaX* strains

C-terminally TAP-tagged Dop was immunoprecipitated from *Mtb* (120 OD equivalents) in 50mM NaH_2_PO_4_ pH 7.4, 100mM NaCl by batch purification with EZview^TM^ Red ANTI-FLAG M2 Affinity gel (Sigma) and eluted with 3XFLAG peptide (Sigma) according to manufacturer’s instructions. Elution was examined by SDS-PAGE on a 10% acrylamide gel and Coomassie stained. The dominant binding partner was excised from the gel and processed by the Proteomics Laboratory at NYU Langone. In brief, the gel was destained with a 1:1 (v/v) solution of methanol and 100 mM ammonium bicarbonate. Gel pieces were dehydrated with an acetonitrile rinse and use of a SpeedVac concentrator. Samples were reduced with DTT at 57°C for 1 hour and alkylated with iodoacetamide at RT in the dark for 45 min. Samples were dehydrated again as before and digested with sequencing grade modified trypsin (Promega) shaking overnight at RT. Peptides were extracted from dried gel pieces using a 1:2 (v/v) 5% formic acid/ acetonitrile extraction buffer at 37°C shaking for 15 min. Peptide solution was dehydrated by SpeedVac and reconstituted with 0.5% acetic acid acidified to pH pH≤2 with 10% TFA. Samples were rinsed three times with 0.1% TFA on an equilibrated microspin Harvard apparatus (Millipore) using a microcentrifuge and eluted with 40% acetonitrile in 0.5% acetic acid followed by the addition of 80% acetonitrile in 0.5% acetic acid. Organic solvent was removed by SpeedVac and samples were reconstituted in 0.5% acetic acid. LC separation was performed online with the autosampler of a EASY-nLC 1000 and gradient eluted directly into the Q Exactive mass spectrometer (1 hr gradient). High resolution full MS spectra were acquired with a resolution of 70,000, an AGC target of 1e6, with a maximum ion time of 120 ms, and scan range of 400 to 1500 m/z. Following each full MS twenty data-dependent high resolution HCD MS/MS spectra were acquired (resolution of 17,500, AGC target of 5e4, maximum ion time of 120 ms, one microscan, 2 m/z isolation window, fixed first mass of 150 m/z, and NCE of 27).

MS/MS spectra were searched against a Uniprot *Mycobacterium tuberculosis* database using Sequest within Proteome Discoverer 1.4. Results were filtered to exclude contaminant (non *Mtb*) proteins. CoaX was identified as the Dop_TAP_ binding partner as it was the protein with the greatest number of peptide spectral matches (PSMs).

Mass spectrometry to globally identify Dop and PafA binding partners was performed as described above with a few changes to the sample preparation. C-terminally tagged Dop_TAP_ and PafA_TAP_ were immunoprecipitated from quadruplicate *Mtb* cultures (51 OD equivalents) using anti-FLAG resin as described above. Following digestion, peptides were rinsed thrice with 0.1% TFA using C18 Spin Columns (Pierce) using a microcentrifuge and eluted with 80% acetonitrile in 0.5% acetic acid.

Total proteomics was performed on lysates extracted from quadruplicate cultures (15 OD equivalents) of WT and Δ*coaX* strains. Lysates were prepared in 100 mM Tris pH 8.0 1mM EDTA and analysis performed as described above with a few changes. Samples were supplemented with SDS for efficient denaturation. Proteins were cleaned and digested on SP3 magnetic beads. Peptides were analyzed in data-independent (DIA) mode on Q-Exactive HFX using Evosep One LC 88 min gradient separation. Significantly differentially detected proteins were determined by students T-test with a multiple hypothesis correction threshold of false discovery rate (FDR) of less than 5%.

### Pantothenate kinase activity assays

CoaA*_Mtb_*, CoaX*_Mtb_*, and CoaA*_Mtb_DM_* enzymatic activity was measured using ADP-Glo Luminescent assay (Promega) according to manufacturer’s instructions. In brief, the ADP-Glo™ Kinase Assay measures the formation of ADP by converting it to ATP used to generate light in a luciferase reaction. Recombinant proteins were incubated with increasing concentrations of pantothenate for 30 min in a kinase buffer (50mM HEPES, 25mM KCl, 20mM MgCl_2_, 0.1M EDTA, 2mM DTT, 0.002% Brij-35, 5mM ATP) as previously described (*34*). Reaction was quenched by addition and incubation with ADP Glo Reagent. Kinase Detection Reagent was then added, and luminescence measured with SpectraMax iD5 Microplate reader. Graphs were generated with Prism and represent results from duplicate experiments.

### Differential Fluorimetry Scanning (DSF)

Experimental conditions for all differential scanning fluorimetry (DSF) experiments were standardized as follows: 10 µM of the protein of purified CoaX was mixed with 5 µL 1X SYPRO Orange (Thermo Fisher Scientific) and various concentrations of substrates (1-10 mM ATPγ or 0-5 mM pantothenate) in 50 mM Tris-HCl, 300 mM NaCl, and 10% glycerol. For experiments containing pantothenate and ATPγ, reactions contained 4 mM pantothenate and concentrations of 0-10 mM ATPγ were tested. Reactions were conducted in a 96-well microplate, with a final reaction volume of 25 µL. Reactions were monitored using a BioRad CFX 96 qPCR instrument equipped with a FRET channel.

### Evolutionary Analysis of CoaX sequences

Homologs of CoaX*_Mtb_* were retrieved from sequence profiles searches using the CoaX*_Mtb_* amino acid sequence as query in a PSI-BLAST search (*68*) against the non-redundant protein database from National Center for Biotechnology Information (NCBI) that was clustered down to 50% identity (nr50). A multiple sequence alignment (MSA) of retrieved sequences was generated with FAMSA (*69*), adjusted based on structural data. Visualization and manipulation of 3D structures were performed with the MOL* viewer package (*70*). Taxid and CoaX accession number associated with each species in the MSA: *Mycobacterium tuberculosis H37Rv* (83332, CCP46423.1), *Mycobacterium marinum M* (216594, ACC43507.1), *Mycobacterium leprae Br4923* (561304, CAR70325.1), *Mycolicibacterium smegmatis* (1772, AIU24341.1), *Rhodococcus erythropolis* (1833, AKD95918.1), *Rhodococcus opacus* (37919, WP_105419480.1), *Prescottella agglutinans* (1644129, WP_280762943.1), *Aldersonia kunmingensis* (408066, WP_068280339.1), *Nocardia nova SH22a* (1415166, AHH15251.1), *Nocardia brasiliensis* ATCC 700358 (1133849, AFT98415.1), *Nocardia seriolae* (37332, APA94408.1), *Nocardia mangyaensis* (2213200, APE32922.1), *Umezawaea tangerina* (84725, WP_106185318.1), *Streptomyces coelicolor A3* (*2*) (100226, CAB40880.1), *Streptosporangium roseum* DSM 43021 (479432, ACZ91776.1), *Streptomyces albus* (1888, AJE83257.1), *Parafrankia* (2994362, WP_091279711.1), *Thermobifida fusca YX* (269800, AAZ56915.1), *Thermobispora bispora* DSM 43833 (469371, ADG90113.1), *Clostridium acetobutylicum* DSM 1731 (991791, AEI34108.1), *Bacillus subtilis subsp. subtilis* (135461, ASB68048.1), *Deinococcus radiodurans R1* = ATCC 13939 = DSM 20539 (243230, AAF10040.1), *Thermotoga maritima MSB8* (243274, 3BEX), *Syntrophobacter fumaroxidans MPOB* (335543, ABK16248.1), *Verrucomicrobia bacterium 13_1_20CM_4_54_11* (1805408, OLD86718.1), *Methylacidiphilum infernorum V4* (481448, ACD83758.1), *Candidatus Xiphinematobacter sp. Idaho Grape* (1704307, ALJ56590.1), *Spirochaeta africana* DSM 8902 (889378, AFG38852.1), *Prosthecochloris sp*. CIB 2401 (1868325, ANT64476.1), *Helicobacter pylori* (210, WRC04122.1), *Pseudomonas aeruginosa* (PAO1208964, AAG07667.1).

To determine the phyletic patterns of Type I and Type II PanK enzymes in species shown in the CoaX multiple sequence alignment, we utilized PSI-BLAST searches on the nr50 database using CoaA*_Mtb_* (CCP43845.1) and *Staphylococcus aureus* PanK (NP_646871.1) as queries, respectively. The NCBI taxids of species in the CoaX alignment were employed to filter the searches and assess the presence of Type I and Type II Pantothenate kinases in those species.

### RNAseq

RNA was extracted from 10 OD equivalents of quadruplicate biological cultures of MHD761 (H37Rv pMV306.strep), MHD1845 (Δ*coaX*::*hyg* pMV306.strep), and MHD1846 (Δ*coaX*::*hyg* pMV306.strep-*coaX*) as previously described (*71*). Library preparation (using Ribozero plus) and paired end 50 bp sequencing (on an SP 100 Cycle Flow Cell v1.5 using an Illumina NovaSeq 6000) was conducted by the NYU Langone Genome Technology Center (available for download at). Reads were mapped to the H37Rv genome (RefSeq identifier GCF_000195955.2 with *socAB* annotation added as previously described (*51*)) using Bowtie2 v2.4.1 (*72*) and sorted using samtools v1.13 (*73*). Aligned read counts for each gene were extracted using the featureCounts command in Subread v2.0.1 (*74*). Normalization and differential expression analysis was performed using DESeq2 v1.40.2 (*75*) defaults using R (*76*), RStudio v2023.9.0.463, and tidyverse v2.0.0 (*77*). Genes with Wald test p-values adjusted for multiple hypothesis testing (Bejamini and Hochberg) less than 0.1 were considered significant. Normalized read counts for pantothenate synthesis enzymes and PPS enzymes were plotted with Prism v10.

### Mouse infections

Duplicate infections of six to eight-week old female mice (C57BL6/J, Jackson Laboratories) with WT, Δ*coaX*, and the complemented strain of *Mtb* were performed as previously described (*5*) with the approval of the New York University Institutional Animal Care and Use Committee. Mice were humanely sacrificed and organs harvested at days 1 (n =6), 22 (n = 8), 56 (n = 8), and 196 days (n = 8-10) following infection. Lungs were harvested at all time points and spleens were harvested at all time points except day 1.

### Depupylation assays

Purified pupylated substrates (_Myc_Pup∼_FLAG_PanB*_Mtb_*__his_ or _Myc_Pup∼_FLAG_FabD*_Mtb_*__his_) were mixed with Dop*_Msm_* at a 4:1 ratio (substrate: Dop) in 50 mM NaPO_4_, 10 mM MgCl_2_, 150 mM NaCl, 1 mM DTT, 5 mM ATP, and 10% glycerol (total volume of 100 µL) on ice. _Myc_Pup∼_FLAG_PanB*_Mtb_*__his_ depupylation reactions contained 2 µg of purified substrate; _Myc_Pup∼_FLAG_FabD*_Mtb_*__his_ reactions contained 0.9µg. When included, CoaX was mixed at a ratio of 2:1 with Dop, prior to the addition of substrate. When included, 1 mM pantothenate was pre-mixed with reactions prior to the addition of substrate.

A sample was collected at start, and reactions incubated at 30°C in an ThermoMixer C (Eppendorf). Additional samples were collected at 15, 45, 120, and 240 min, mixed with 4X SB and boiled at 100°C. Pupylated protein in each sample was visualized by SDS-PAGE. Depupylation of _Myc_Pup∼_FLAG_PanB*_Mtb_*__his_ was analyzed by staining with ProtoStain Blue (National Diagnostics) according to manufacturer’s instructions. Depupylation of _Myc_Pup∼_FLAG_FabD*_Mtb_*__his_ was analyzed using Pierce^TM^ Silver Stain for Mass Spectrometry according to manufacturer’s instructions using Glacial Acetic Acid and 190 proof Ethanol (Decon Labs).

Gels were scanned and band density was measured with Fiji (Image J). Pupylated protein remaining at each time point was calculated as a ratio of the proportion of pupylated protein at each time point relative to the starting proportion of pupylated protein. Results were graphed and Mann-Whitney tests performed in Prism v10.

---

**Fig. S1.**
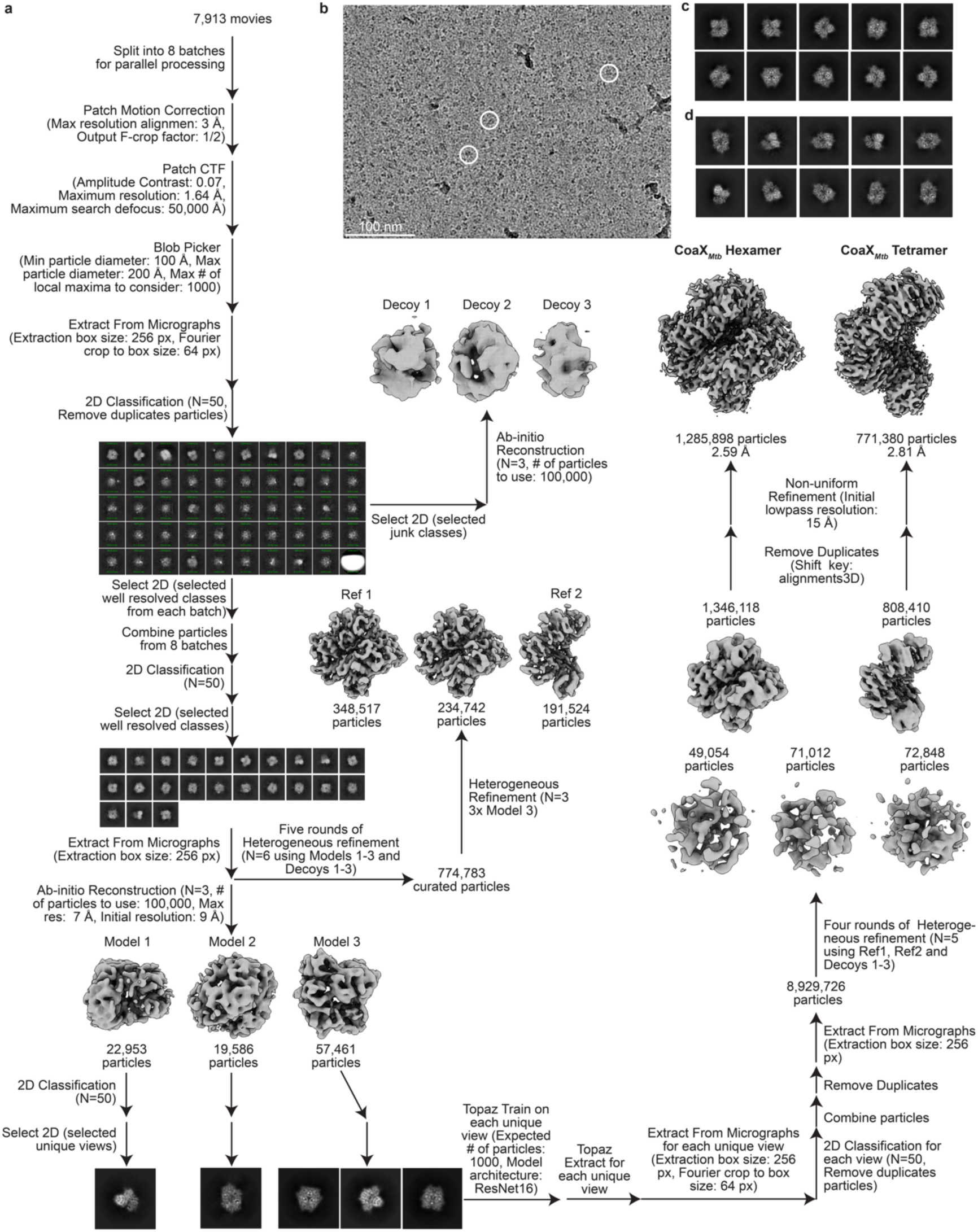
Cryo-EM data processing workflow. **(A)** Cryo-EM data processing pipeline. **(B)** Representative cryo-EM micrograph. Particles of interest are circled in white. Scale bar (100 nm) is indicated on the bottom left of the micrograph. **(C-D)** Representative 2D class averages showing different views of the particles were generated in cryoSPARC (*57*) using the final set of particles extracted with box size of 256 pixels and no binning for CoaX*_Mtb_* hexamer complex (**c**) and CoaX*_Mtb_* tetramer complex (**d**).

**Fig. S2.**
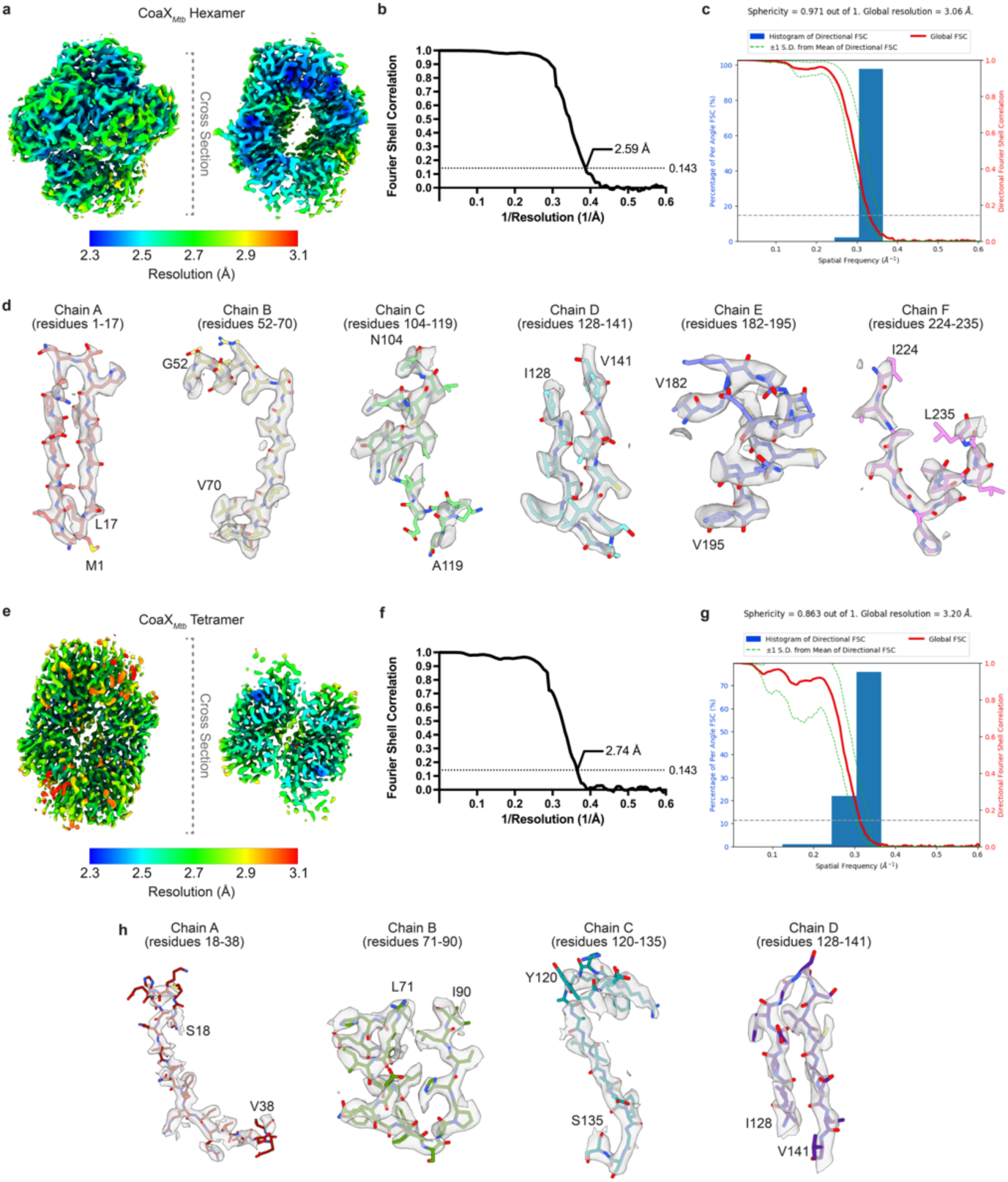
Cryo-EM validation and representative densities. **(A)** CoaX*_Mtb_* hexamer map (contour level: 0.25) colored by local resolution that was estimated in cryoSPARC (*1*). (left) Whole structure view. (right) Cross-sectional view. **(B)** Gold-standard FSC curve calculated in cryoSPARC for raw maps of CoaX*_Mtb_* hexamer. The dotted line represents the 0.143 FSC cutoff. **(C)** Directional 3DFSC calculated for the CoaX*_Mtb_* hexamer map. **(D)** Examples of cryo-EM densities (transparent gray surface) with corresponding areas of the model (sticks) are shown from each subunit for the CoaX*_Mtb_* hexamer map. Protein densities rendered using ChimeraX^3^ ‘volume zone’ with 2.0 Å distance cutoff around the indicated protein residues with the following contour levels: ChainA (0.6); ChainB (0.3), ChainC (0.4), ChainD (0.6), ChainE (0.3), ChainF (0.4). **(E)** CoaX*_Mtb_* tetramer map (contour level: 0.3) colored by local resolution that was estimated in cryoSPARC. (left) Whole structure view. (right) Cross-sectional view. **(F)** Gold-standard FSC curve calculated in cryoSPARC for raw maps of CoaX*_Mtb_* tetramer. The dotted line represents the 0.143 FSC cutoff.**(G)** Directional 3DFSC calculated for the CoaX*_Mtb_* tetramer map. **(H)** Examples of cryo-EM densities (transparent gray surface) with corresponding areas of the model (sticks) are shown from each subunit for the CoaX*_Mtb_* tetramer map. Protein densities rendered using ChimeraX (*2*) ‘volume zone’ with 2.0 Å distance cutoff around the indicated protein residues with the following contour levels: ChainA (0.25); ChainB (0.3), ChainC (0.3), ChainD (0.6).

**Fig. S3.**
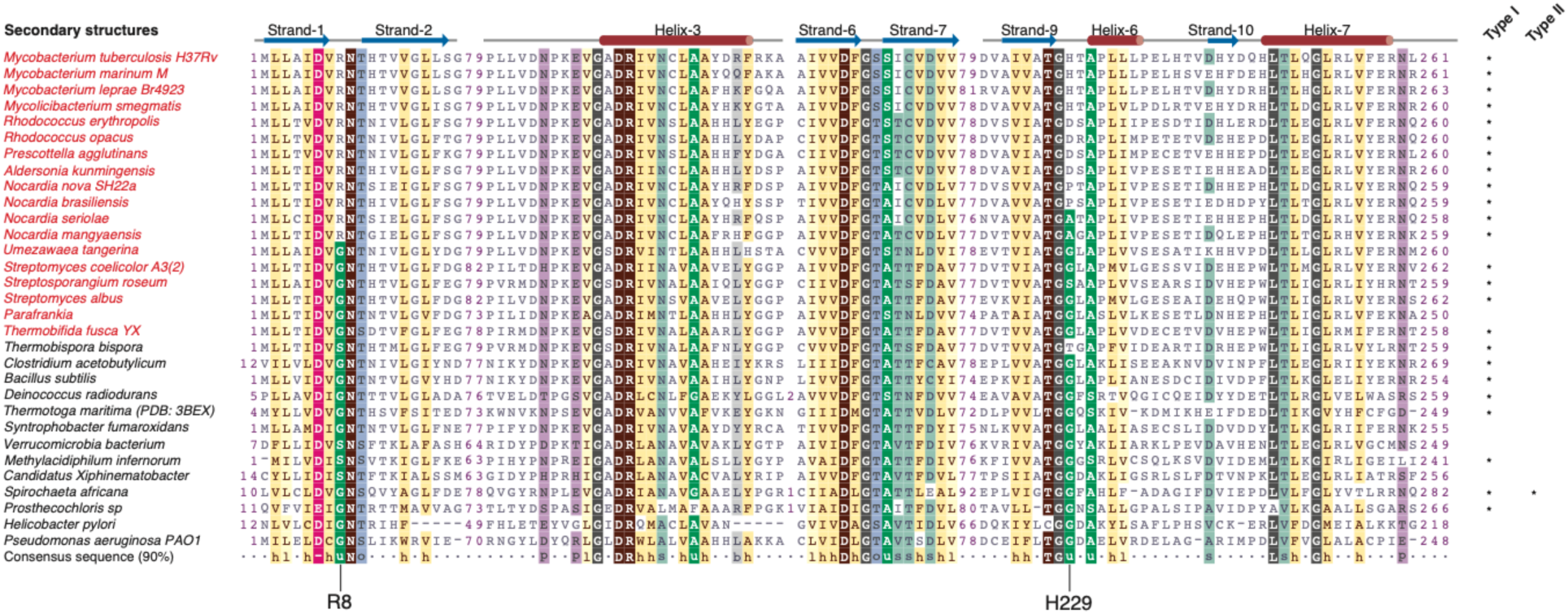
Multiple sequence alignment of CoaX homologs from proteasome containing and absent species. Proteasome containing species are indicated in red. Secondary structural features are indicated above. Species containing additional Type I and Type II PanK enzymes are indicated with an asterisk. For TaxIDs and CoaX accession numbers see Materials and Methods.

**Fig. S4.**
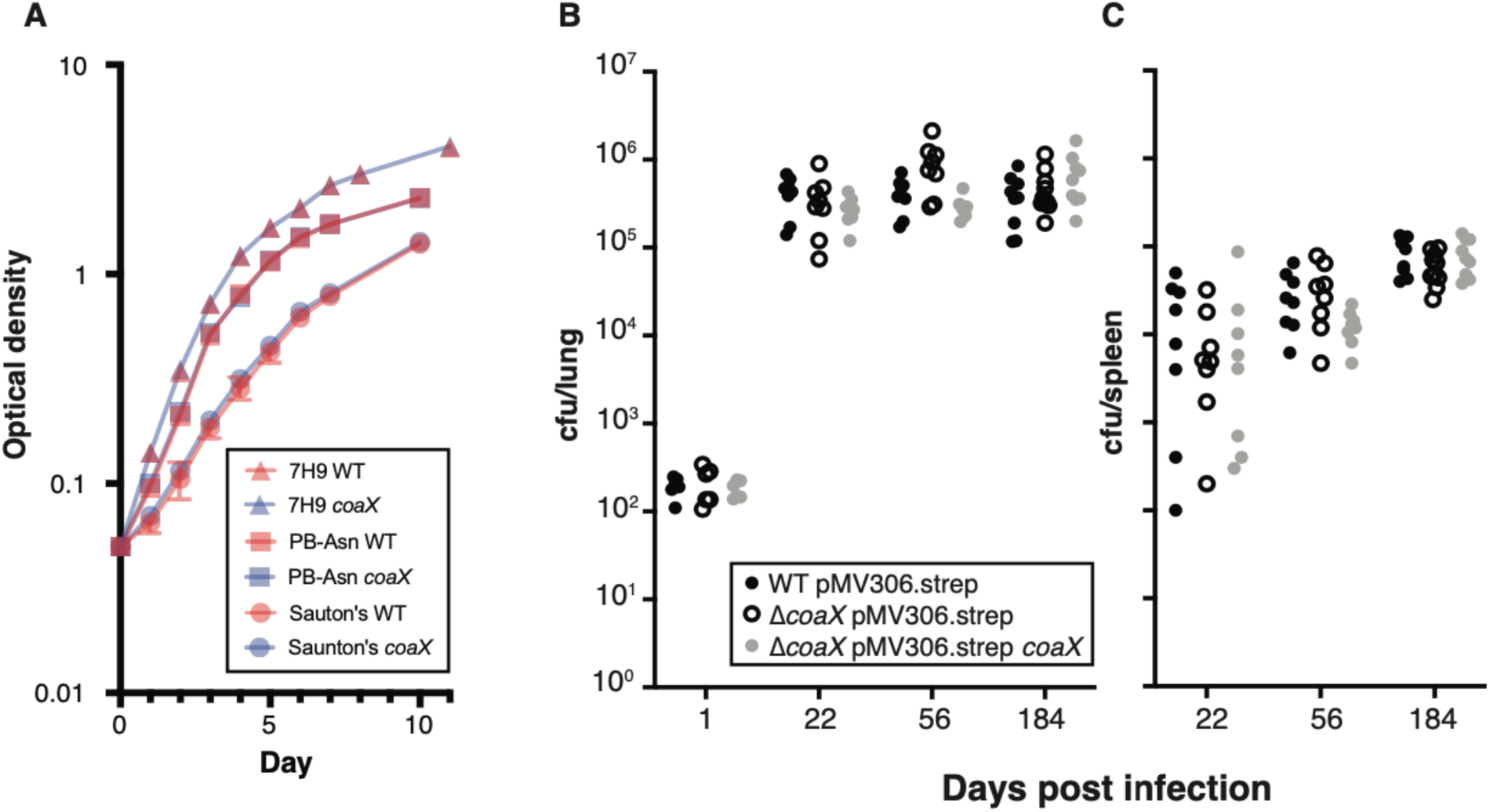
Loss of *coaX* had no effect on growth *in vitro* or virulence in C57BL6J mice. **(A)** Growth curves comparing the Δ*coaX* mutant (blue) to WT (red) in 7H9 (triangles) Sauton’s media (circles), or Proskauer-Beck media (squares). Growth curves were performed in duplicate. **(B)** Colony forming units (cfus) from the lungs or spleen of C57BL/6J mice infected with WT (black filled circles), Δ*coaX* (black open circles), or a complemented strain (grey, filled circles) are plotted following low-dose aerosol infection. WT and Δ*coaX* infection strains were transformed with an empty complementation vector a control.

**Fig. S5.**
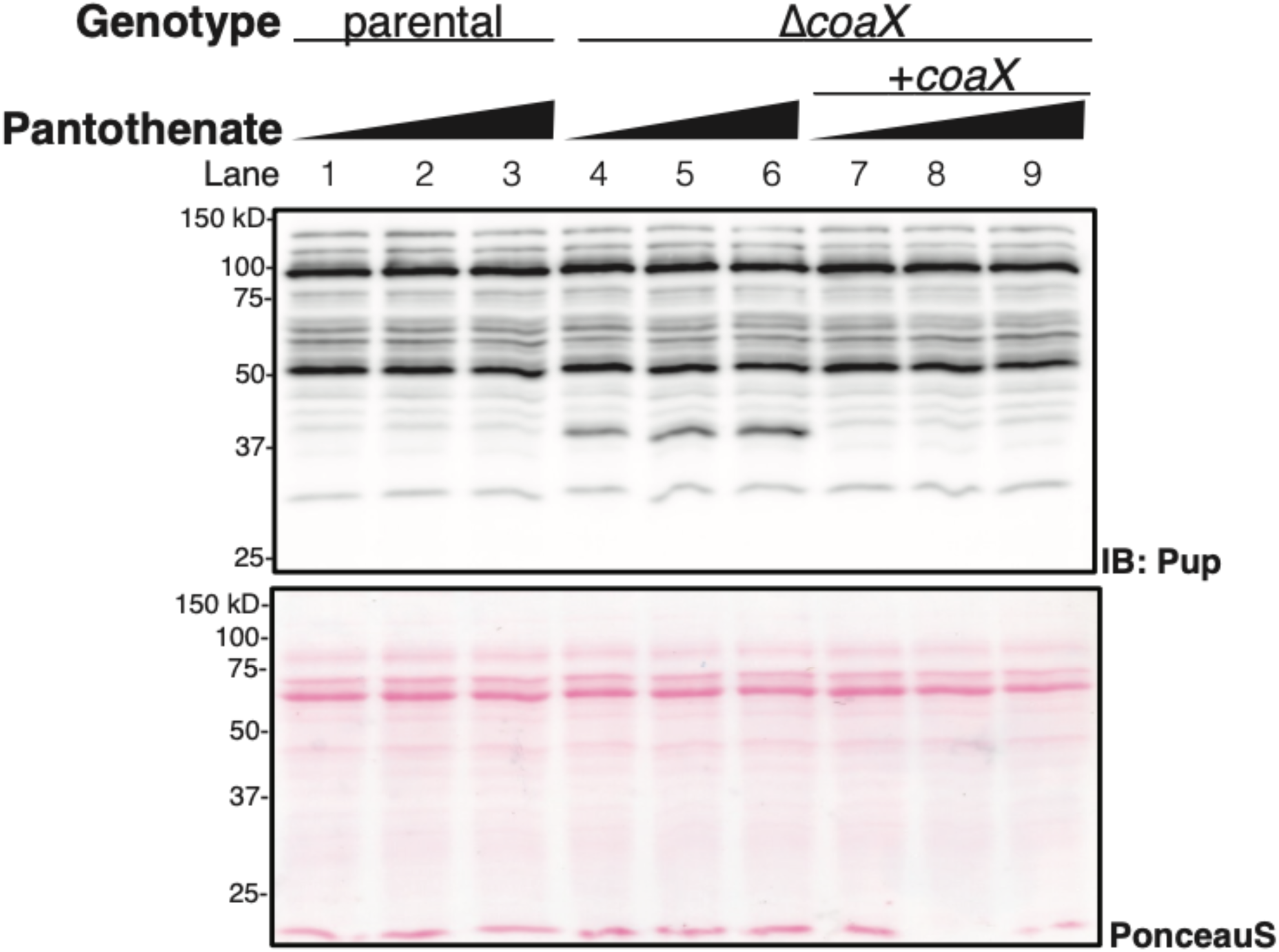
Global pupylome unchanged by pantothenate supplementation in Δ*coaX*, WT, or complemented strain. Immunoblot (IB) against lysates extracted from Δ*coaX*, *mpa*::Tn, complemented strains, and parental controls supplemented with 0, 10, or 100 µM pantothenate are shown. Parental controls were transformed with empty complementation vectors. IB was performed with monoclonal antibodies against Pup; PonceauS shown as a loading control.

**Fig. S6.**
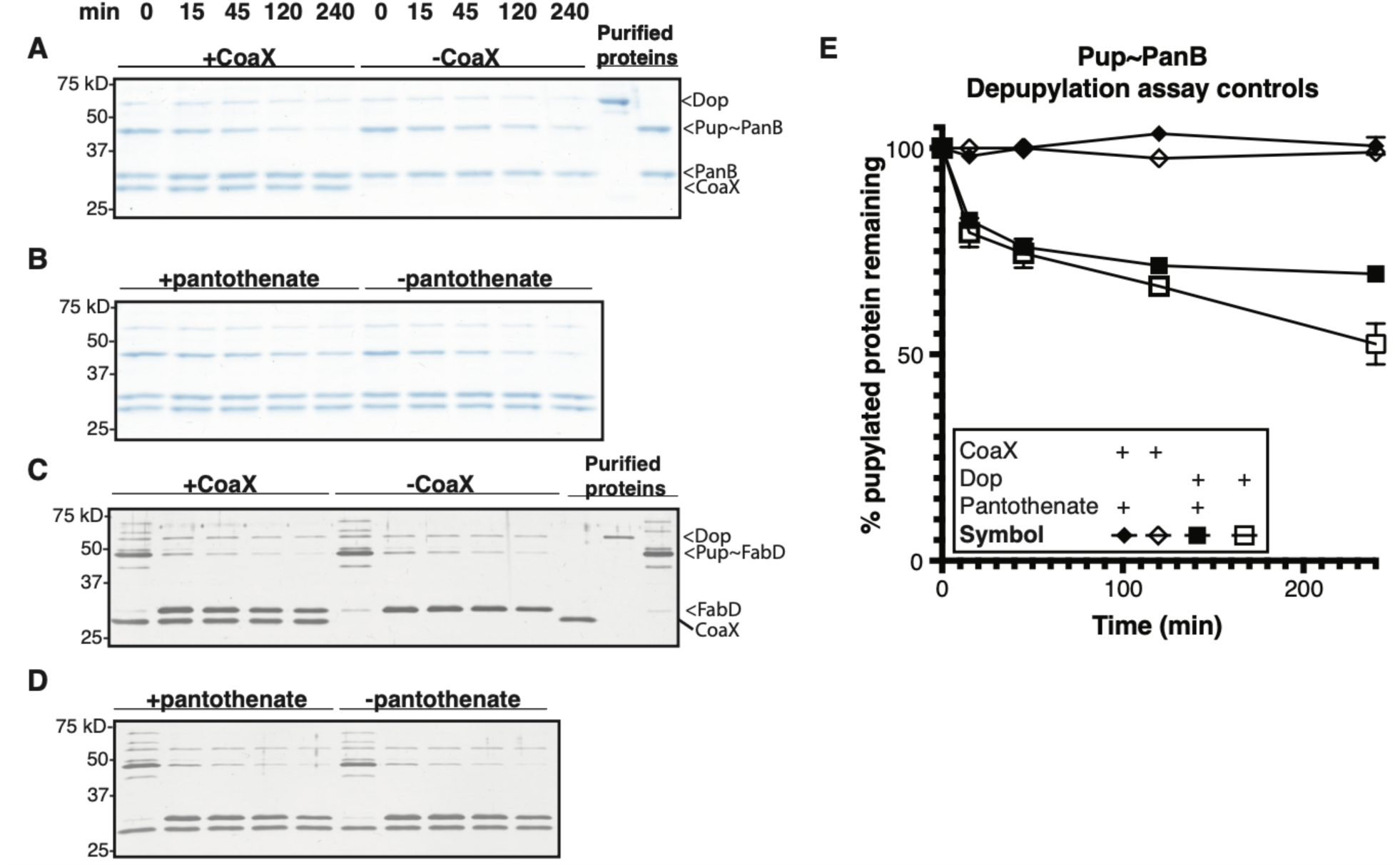
Representative depupylation reaction gels used for quantification, additional controls. Representative Coomassie-stained gels containing Pup∼PanB depupylation samples used to quantify reactions, **a,** with and without CoaX or, **b,** reactions containing CoaX and with or without 1 mM pantothenate. Representative silver-stained gels containing Pup∼FabD depupylation samples used to quantify reactions, **a,** with and without CoaX or, **b,** reactions containing CoaX and with or without 1 mM pantothenate. Purified proteins are indicated. **e,** Quantification of Pup∼PanB depupylation with (closed square) and without (open square) pantothenate in the absence of CoaX. Quantification of Pup∼PanB with (closed diamond) and without (open diamond) pantothenate in the absence of Dop.

**Table S1.**
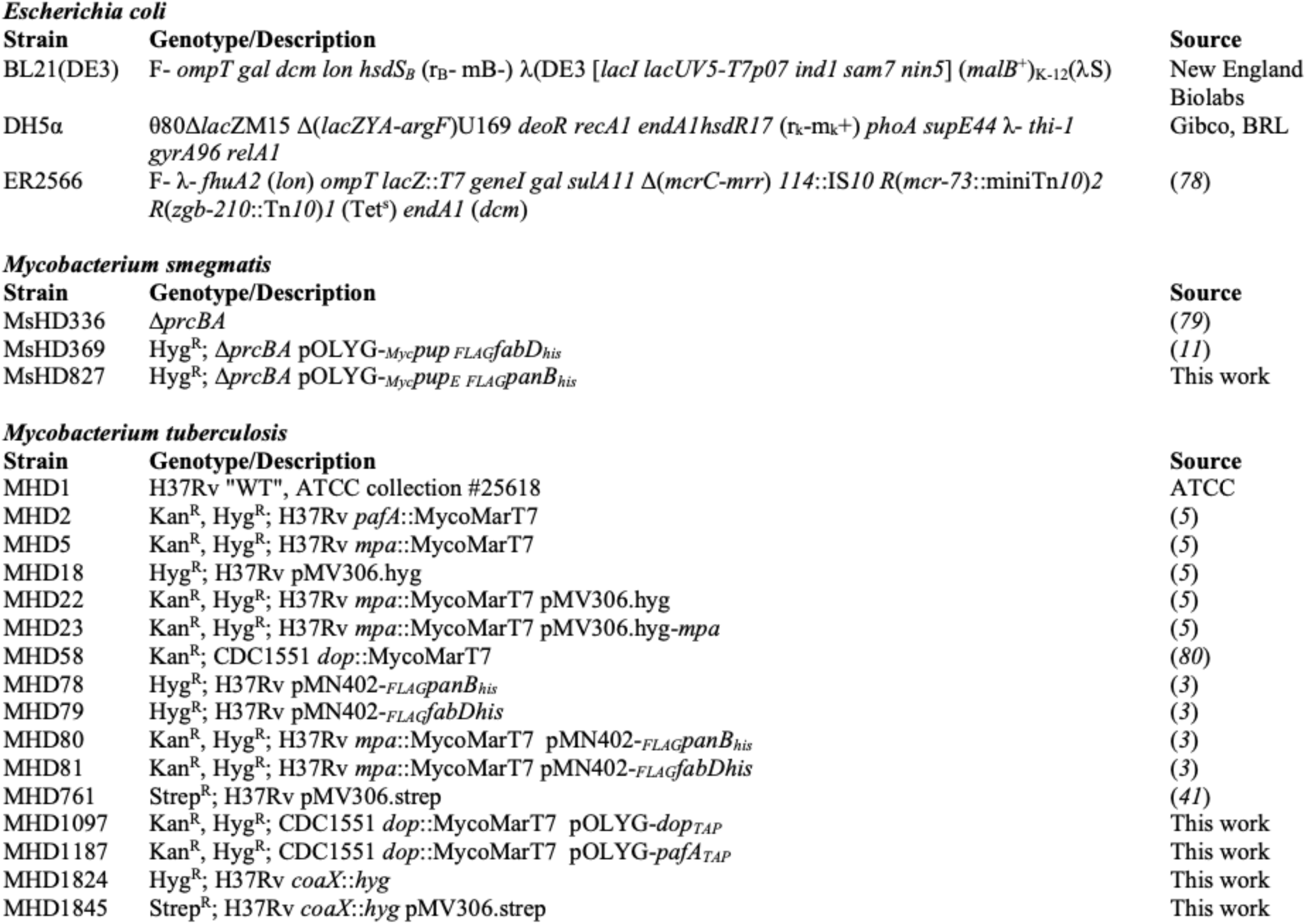
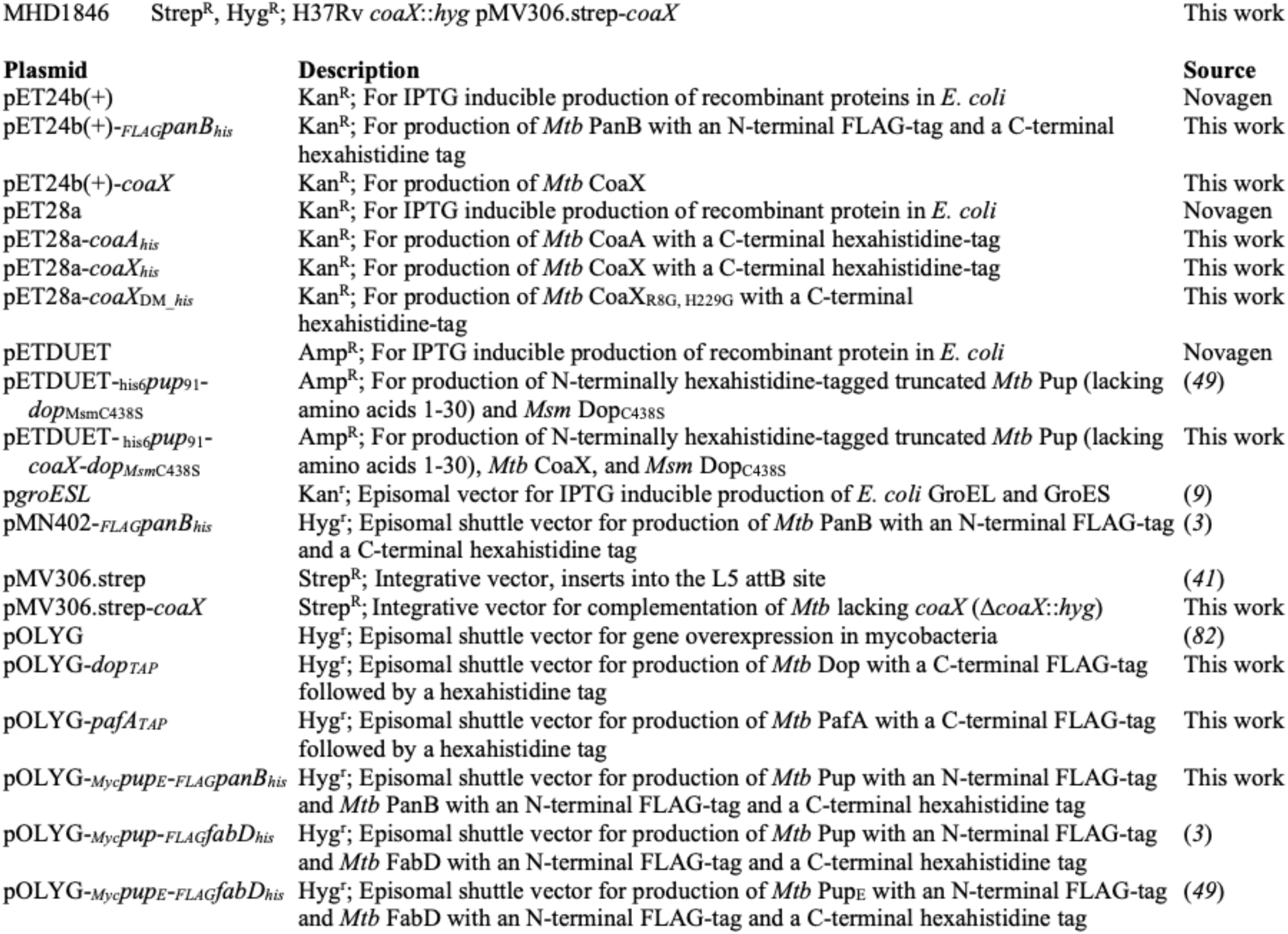
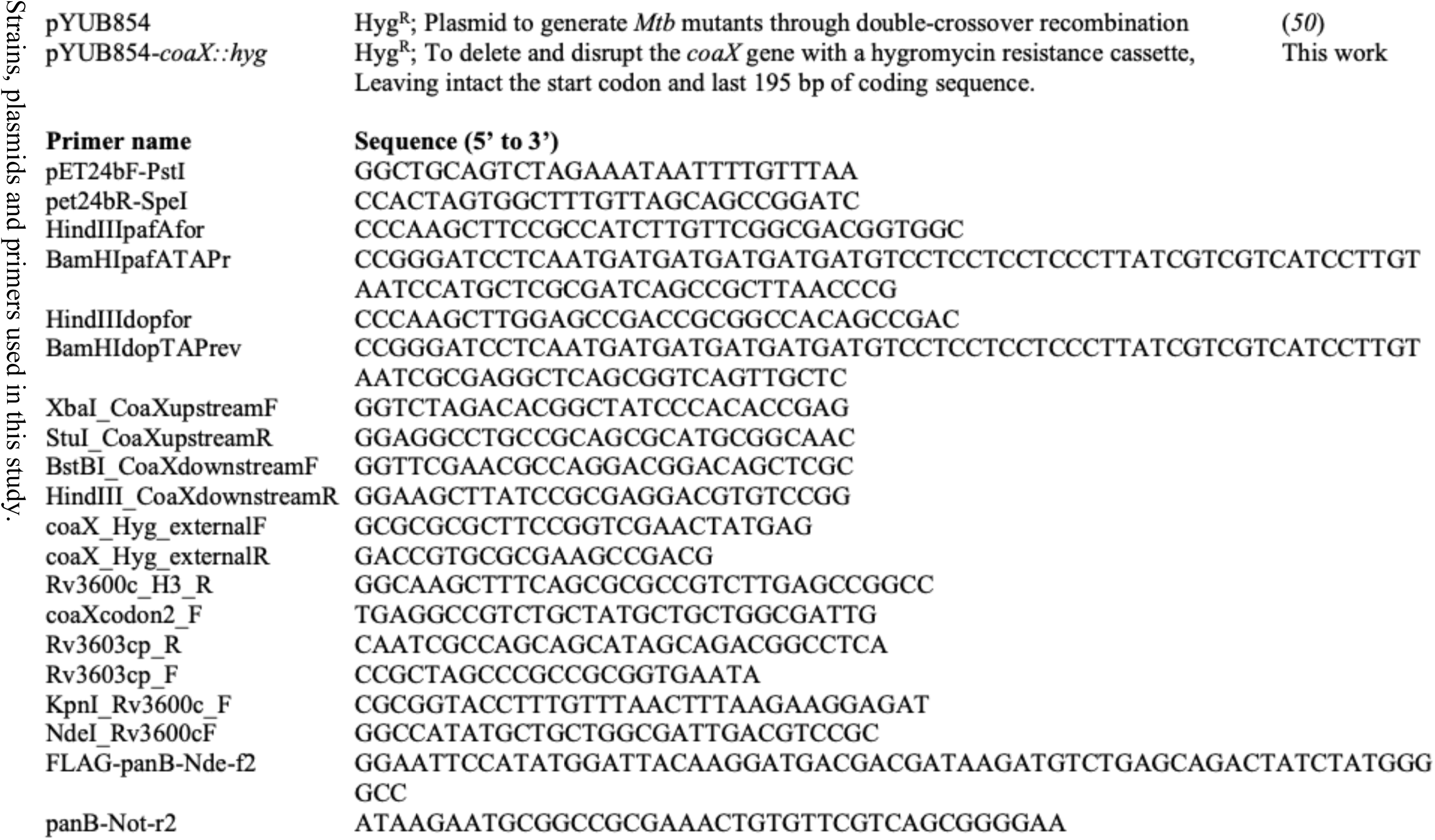

**Table S2.**

| Accession # | Protein | Score | Coverage | # Peptides | # PSMs | # AAs | MW [kDa] |
| --- | --- | --- | --- | --- | --- | --- | --- |
| **P9WPA1** | **coaX** | **82.26** | **28.68** | **9** | **36** | **272** | **29.3** |
| P9WNU9 | dop | 32.69 | 21.78 | 13 | 24 | 505 | 55.1 |
| P9WFD3 | Rv3134c | 34.29 | 21.27 | 4 | 17 | 268 | 28.0 |
| I6W XK4 | ptpB | 18.69 | 33.33 | 8 | 16 | 276 | 30.1 |
| P9WPE7 | groEL2 | 26.09 | 16.48 | 11 | 13 | 540 | 56.7 |
| P9WJC7 | espG3 | 18.53 | 18.64 | 6 | 10 | 295 | 31.5 |
| P9WIV5 | nuoE | 7.56 | 19.05 | 4 | 10 | 252 | 27.2 |

Identification of a Dop binding partner, CoaX. Proteins purified by immunoprecipitation of Dop<sub>TAP</sub> from *Mtb* were separated by SDS-PAGE. A dominant protein uniquely copurifying with Dop<sub>TAP</sub> compared to PafA<sub>TAP</sub>, running between 25 and 37 kD, was excised and the proteins extracted and analyzed by mass spectrometry. Proteins with 10 or more peptide spectral matches (PSMs) detected are shown. Based on the number of PSMs and size of proteins detected, the band was identified as CoaX. See Materials and Methods for more details.

Table S3.

Global identification of Dop and PafA binding partners by purification and Mass spectrometry. Dop_TAP_ and PafA_TAP_ were purified from *Mtb* lysates. CoaX had the fourth greatest number of PSMs of any protein detected in the Dop purifications and was undetected in all PafA samples. Only proteins with 10 or more average PSMs in Dop_TAP_ and/or PafA_TAP_ samples are shown. See Materials and Methods for more details.

**Table S4.**
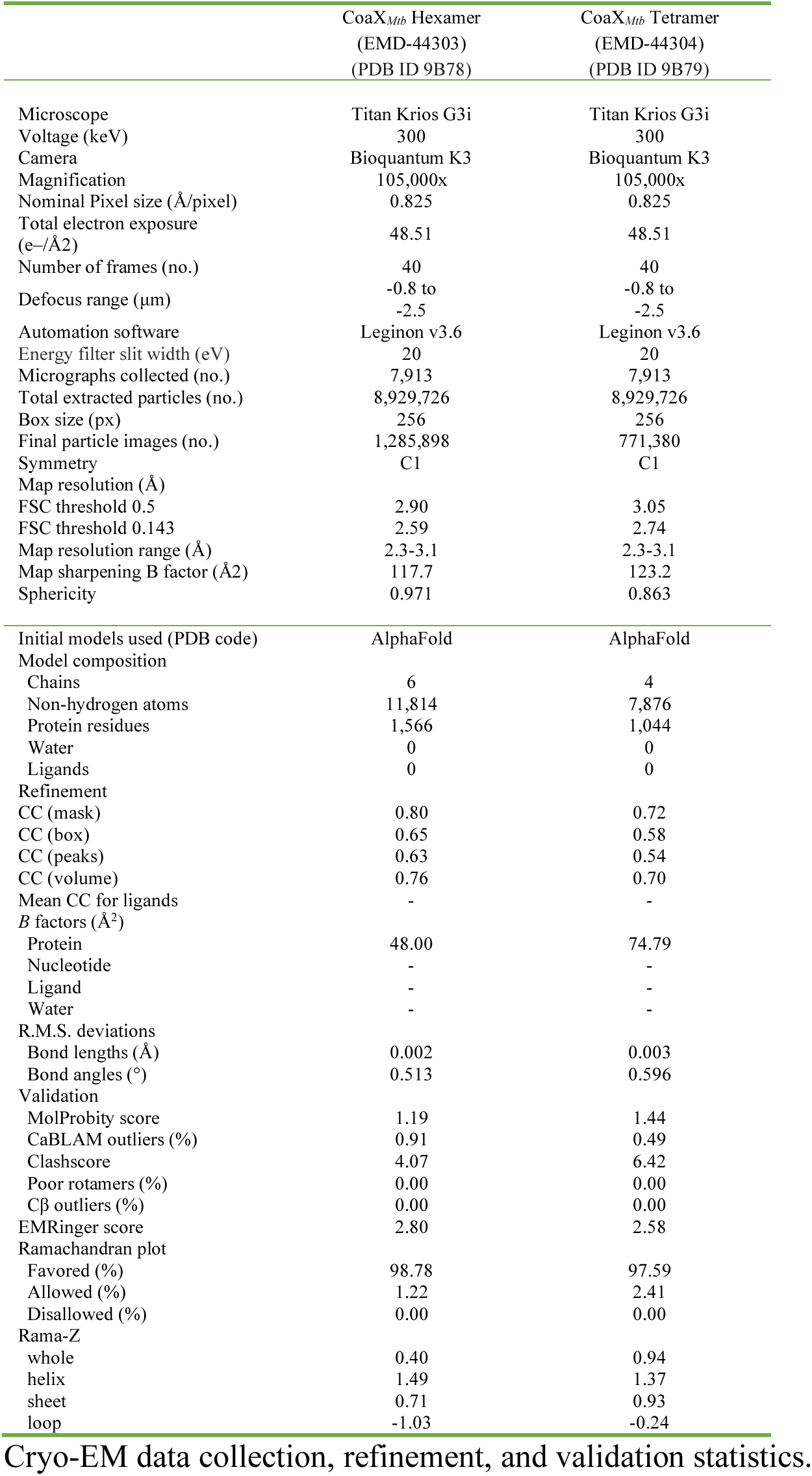

|  | CoaX<sub>Mtb</sub> Hexamer<br>(EMD-44303)<br>(PDB ID 9B78) | CoaX<sub>Mtb</sub> Tetramer<br>(EMD-44304)<br>(PDB ID 9B79) |
| --- | --- | --- |
| Microscope | Titan Krios G3i | Titan Krios G3i |
| Voltage (keV) | 300 | 300 |
| Camera | Bioquantum K3 | Bioquantum K3 |
| Magnification | 105,000x | 105,000x |
| Nominal Pixel size (Å/pixel) | 0.825 | 0.825 |
| Total electron exposure<br>(e-/Å<sup>2</sup>) | 48.51 | 48.51 |
| Number of frames (no.) | 40 | 40 |
| Defocus range (µm) | -0.8 to<br>-2.5 | -0.8 to<br>-2.5 |
| Automation software | Leginon v3.6 | Leginon v3.6 |
| Energy filter slit width (eV) | 20 | 20 |
| Micrographs collected (no.) | 7,913 | 7,913 |
| Total extracted particles (no.) | 8,929,726 | 8,929,726 |
| Box size (px) | 256 | 256 |
| Final particle images (no.) | 1,285,898 | 771,380 |
| Symmetry | C1 | C1 |
| Map resolution (Å) |  |  |
| FSC threshold 0.5 | 2.90 | 3.05 |
| FSC threshold 0.143 | 2.59 | 2.74 |
| Map resolution range (Å) | 2.3-3.1 | 2.3-3.1 |
| Map sharpening B factor (Å<sup>2</sup>) | 117.7 | 123.2 |
| Sphericity | 0.971 | 0.863 |
| Initial models used (PDB code) | AlphaFold | AlphaFold |
| Model composition |  |  |
| Chains | 6 | 4 |
| Non-hydrogen atoms | 11,814 | 7,876 |
| Protein residues | 1,566 | 1,044 |
| Water | 0 | 0 |
| Ligands | 0 | 0 |
| Refinement |  |  |
| CC (mask) | 0.80 | 0.72 |
| CC (box) | 0.65 | 0.58 |
| CC (peaks) | 0.63 | 0.54 |
| CC (volume) | 0.76 | 0.70 |
| Mean CC for ligands | - | - |
| B factors (Å<sup>2</sup>) |  |  |
| Protein | 48.00 | 74.79 |
| Nucleotide | - | - |
| Ligand | - | - |
| Water | - | - |
| R.M.S. deviations |  |  |
| Bond lengths (Å) | 0.002 | 0.003 |
| Bond angles (°) | 0.513 | 0.596 |
| Validation |  |  |
| MolProbity score | 1.19 | 1.44 |
| CaBLAM outliers (%) | 0.91 | 0.49 |
| Clashscore | 4.07 | 6.42 |
| Poor rotamers (%) | 0.00 | 0.00 |
| Cβ outliers (%) | 0.00 | 0.00 |
| EMRinger score | 2.80 | 2.58 |
| Ramachandran plot |  |  |
| Favored (%) | 98.78 | 97.59 |
| Allowed (%) | 1.22 | 2.41 |
| Disallowed (%) | 0.00 | 0.00 |
| Rama-Z |  |  |
| whole | 0.40 | 0.94 |
| helix | 1.49 | 1.37 |
| sheet | 0.71 | 0.93 |
| loop | -1.03 | -0.24 |

Cryo-EM data collection, refinement, and validation statistics.

**Table S5.**

Proteomic comparison of WT (H37Rv) and Δ*coaX Mtb* strains. Quadruplicate lysates extracted from WT and Δ*coaX* strains were analyzed by data-independent acquisition mass spectrometry. Proteins with FC greater than or equal to 1.1, or less than or equal to 0.9, are shown. Proteins with a q-value < 0.05 were considered significant. See Materials and Methods for more details.

**Table S6.**

Statistical analysis of RNA-seq for all *Mtb* genes. RNA was extracted from four independent cultures of MHD761 (H37Rv pMV306.strep; “WT”), MHD1845 (Δ*coaX*::*hyg* pMV306.strep; “Δ*coaX*”), and MHD1846 (Δ*coaX*::*hyg* pMV306.strep-*coaX*; “complement”) and sequenced on an Illumina NovaSeq 6000. 50 bp paired reads were mapped to the H37Rv reference genome and aligned read counts extracted and normalized as described in the materials and methods.

## Notes

### Competing Interest Statement

The authors have declared no competing interest.

