## Supplementary material for "Identification of a proteolysis regulator for an essential enzyme in *Mycobacterium tuberculosis*": Table S6

| | WT v Δ*coaX* | | Δ*coaX* v complement | | WT v complement | |
|---|---|---|---|---|---|---|
| **GeneID** | **log2[FC]** | **padj** | **log2[FC]** | **padj** | **log2[FC]** | **padj** |
| dnaA | 0.149 | 0.999 | -0.101 | 0.940 | -0.250 | 0.837 |
| dnaN | 0.145 | 0.999 | 0.011 | 0.981 | -0.135 | 0.819 |
| recF | 0.046 | 0.999 | 0.013 | 0.993 | -0.034 | 0.983 |
| Rv0004 | -0.033 | 0.999 | 0.096 | 0.895 | 0.129 | 0.846 |
| gyrB | 0.154 | 0.999 | -0.120 | 0.931 | -0.274 | 0.835 |
| gyrA | 0.202 | 0.999 | -0.002 | 0.996 | -0.204 | 0.816 |
| Rv0007 | 0.038 | 0.999 | 0.235 | 0.831 | 0.197 | 0.816 |
| Rv0008c | -0.195 | 0.999 | 0.307 | 0.848 | 0.501 | 0.816 |
| ppiA | -0.078 | 0.999 | -0.058 | 0.927 | 0.020 | 0.967 |
| Rv0010c | 0.016 | 0.999 | -0.018 | 0.970 | -0.034 | 0.932 |
| Rv0011c | -0.111 | 0.999 | -0.210 | 0.852 | -0.098 | 0.896 |
| Rv0012 | 0.070 | 0.999 | 0.114 | 0.895 | 0.044 | 0.951 |
| trpG | 0.051 | 0.999 | 0.007 | 0.990 | -0.045 | 0.942 |
| pknB | -0.029 | 0.999 | -0.116 | 0.859 | -0.087 | 0.862 |
| pknA | 0.046 | 0.999 | -0.286 | 0.848 | -0.333 | 0.816 |
| pbpA | 0.086 | 0.999 | 0.047 | 0.948 | -0.039 | 0.945 |
| rodA | -0.046 | 0.999 | -0.118 | 0.848 | -0.072 | 0.868 |
| pstP | -0.004 | 0.999 | -0.076 | 0.859 | -0.072 | 0.837 |
| fhaB | 0.057 | 0.999 | -0.100 | 0.870 | -0.157 | 0.816 |
| fhaA | -0.019 | 0.999 | -0.038 | 0.906 | -0.019 | 0.942 |
| Rv0021c | 0.003 | 0.999 | 0.023 | 0.974 | 0.020 | 0.977 |
| whiB5 | 0.140 | NA | 0.239 | 0.879 | 0.098 | 0.937 |
| Rv0023 | 0.096 | 0.999 | 0.261 | 0.826 | 0.166 | 0.825 |
| Rv0024 | 0.049 | 0.999 | 0.465 | 0.660 | 0.416 | 0.816 |
| Rv0025 | -0.038 | 0.999 | -0.066 | 0.922 | -0.028 | 0.958 |
| Rv0026 | -0.006 | 0.999 | -0.187 | 0.878 | -0.181 | 0.846 |
| Rv0027 | -0.021 | 0.999 | -0.159 | 0.856 | -0.138 | 0.838 |
| Rv0028 | -0.141 | NA | -0.088 | 0.942 | 0.053 | 0.955 |
| Rv0029 | -0.087 | 0.999 | 0.157 | 0.831 | 0.244 | 0.816 |
| Rv0030 | 0.048 | 0.999 | 0.159 | 0.931 | 0.111 | 0.942 |
| bioF2 | 0.063 | 0.999 | -0.435 | 0.859 | -0.498 | 0.819 |
| acpA | -0.374 | NA | -0.875 | 0.830 | -0.502 | 0.837 |
| Rv0034 | -0.342 | NA | -0.255 | 0.879 | 0.088 | 0.947 |
| fadD34 | 0.020 | 0.999 | -0.882 | 0.820 | -0.902 | 0.816 |
| Rv0036c | 0.048 | 0.999 | 0.281 | 0.832 | 0.233 | 0.817 |
| Rv0037c | -0.039 | 0.999 | 0.353 | 0.832 | 0.392 | 0.816 |
| Rv0038 | 0.027 | 0.999 | 0.383 | 0.820 | 0.356 | 0.816 |
| Rv0039c | -0.029 | 0.999 | 0.758 | 0.787 | 0.787 | 0.816 |
| mtc28 | 0.073 | 0.999 | 0.014 | 0.982 | -0.059 | 0.930 |
| leuS | 0.027 | 0.999 | -0.128 | 0.831 | -0.155 | 0.816 |
| Rv0042c | 0.017 | 0.999 | 0.196 | 0.831 | 0.178 | 0.816 |
| Rv0043c | -0.066 | 0.999 | -0.253 | 0.848 | -0.187 | 0.840 |
| Rv0044c | -0.113 | 0.999 | 0.107 | 0.886 | 0.221 | 0.816 |
| Rv0045c | 0.001 | 0.999 | 0.143 | 0.879 | 0.141 | 0.865 |
| ino1 | 0.024 | 0.999 | -0.278 | 0.826 | -0.301 | 0.816 |
| Rv0047c | 0.143 | 0.999 | -0.115 | 0.879 | -0.258 | 0.816 |
| Rv0048c | 0.106 | 0.999 | 0.186 | 0.832 | 0.080 | 0.879 |
| Rv0049 | -0.101 | 0.999 | 0.254 | 0.831 | 0.355 | 0.816 |
| ponA1 | 0.051 | 0.999 | -0.037 | 0.961 | -0.088 | 0.882 |
| Rv0051 | -0.012 | 0.999 | 0.062 | 0.936 | 0.073 | 0.904 |
| Rv0052 | -0.059 | 0.999 | -0.018 | 0.977 | 0.041 | 0.950 |
| rpsF | -0.057 | 0.999 | -0.148 | 0.883 | -0.091 | 0.910 |
| ssb | -0.022 | 0.999 | 0.064 | 0.948 | 0.086 | 0.909 |
| rpsR1 | 0.276 | 0.999 | 0.116 | 0.879 | -0.161 | 0.819 |
| rplI | -0.088 | 0.999 | -0.295 | 0.826 | -0.207 | 0.816 |
| Rv0057 | 0.069 | 0.999 | -0.389 | 0.820 | -0.458 | 0.816 |
| dnaB | -0.006 | 0.999 | -0.234 | 0.660 | -0.228 | 0.816 |
| Rv0059 | 0.117 | 0.999 | -0.464 | 0.843 | -0.581 | 0.816 |
| Rv0060 | 0.008 | 0.999 | -0.342 | 0.826 | -0.350 | 0.816 |
| Rv0061c | 0.005 | 0.999 | 0.170 | 0.848 | 0.165 | 0.817 |
| celA1 | 0.009 | 0.999 | 0.233 | 0.856 | 0.224 | 0.835 |
| Rv0063 | -0.006 | 0.999 | 0.292 | 0.826 | 0.299 | 0.816 |
| Rv0063a | 0.060 | 0.999 | 0.448 | 0.826 | 0.388 | 0.816 |
| Rv0064 | -0.010 | 0.999 | -0.225 | 0.826 | -0.216 | 0.816 |
| vapB1 | -0.056 | NA | -0.626 | 0.820 | -0.570 | 0.816 |
| vapC1 | 0.098 | 0.999 | -0.675 | 0.695 | -0.773 | 0.816 |
| icd2 | 0.062 | 0.999 | -0.135 | 0.856 | -0.196 | 0.816 |
| Rv0067c | 0.095 | 0.999 | -0.085 | 0.890 | -0.180 | 0.816 |
| Rv0068 | -0.053 | NA | -0.131 | 0.879 | -0.078 | 0.899 |
| sdaA | 0.170 | 0.999 | 0.013 | 0.975 | -0.156 | 0.816 |
| glyA2 | 0.220 | 0.999 | 0.097 | 0.879 | -0.122 | 0.837 |
| Rv0071 | -0.087 | NA | -0.284 | 0.852 | -0.197 | 0.862 |
| Rv0072 | 0.184 | 0.999 | 0.017 | 0.970 | -0.168 | 0.816 |
| Rv0073 | 0.135 | 0.999 | -0.025 | 0.970 | -0.160 | 0.821 |
| Rv0074 | 0.186 | 0.999 | -0.014 | 0.982 | -0.200 | 0.816 |
| Rv0075 | 0.066 | 0.999 | 0.031 | 0.948 | -0.035 | 0.927 |
| Rv0076c | -0.032 | NA | -0.395 | 0.826 | -0.363 | 0.816 |
| Rv0077c | -0.455 | NA | 0.140 | 0.915 | 0.596 | 0.816 |
| Rv0078 | -0.275 | NA | -0.363 | 0.879 | -0.088 | 0.958 |
| Rv0078A | -0.003 | 0.999 | -0.207 | 0.832 | -0.204 | 0.816 |
| Rv0078B | -0.108 | 0.999 | -0.494 | 0.854 | -0.386 | 0.852 |
| Rv0079 | -0.603 | 0.999 | -0.046 | 0.974 | 0.557 | 0.816 |
| Rv0080 | -0.525 | 0.303 | -0.551 | 0.114 | -0.025 | 0.965 |
| Rv0081 | -0.420 | 0.999 | 0.037 | 0.978 | 0.457 | 0.817 |
| Rv0082 | -0.254 | 0.999 | -0.153 | 0.879 | 0.101 | 0.897 |
| Rv0083 | -0.356 | 0.999 | -0.219 | 0.879 | 0.137 | 0.900 |
| hycD | -0.315 | 0.999 | -0.030 | 0.977 | 0.286 | 0.831 |
| hycP | -0.222 | 0.999 | -0.088 | 0.958 | 0.134 | 0.921 |
| hycQ | -0.386 | 0.999 | -0.264 | 0.854 | 0.123 | 0.899 |
| hycE | -0.475 | 0.298 | -0.538 | 0.050 | -0.062 | 0.904 |
| Rv0088 | -0.275 | 0.999 | -0.387 | 0.377 | -0.112 | 0.838 |
| Rv0089 | -0.138 | 0.999 | -0.373 | 0.820 | -0.235 | 0.817 |
| Rv0090 | -0.396 | NA | -0.525 | 0.826 | -0.128 | 0.921 |
| mtn | 0.101 | NA | 0.139 | 0.848 | 0.038 | 0.940 |
| ctpA | -0.138 | 0.999 | 0.055 | 0.935 | 0.193 | 0.816 |
| Rv0093c | -0.045 | 0.999 | -0.048 | 0.940 | -0.003 | 0.996 |
| Rv0094c | -0.053 | 0.999 | -0.786 | 0.859 | -0.732 | 0.838 |
| Rv0095c | -0.469 | NA | -0.702 | 0.826 | -0.232 | 0.891 |
| PPE1 | -0.148 | NA | -1.395 | NA | -1.247 | NA |
| Rv0097 | 0.146 | 0.999 | -0.573 | 0.837 | -0.720 | 0.816 |
| fcoT | 0.234 | NA | -0.484 | 0.816 | -0.719 | 0.816 |
| fadD10 | 0.033 | 0.999 | -0.260 | 0.825 | -0.293 | 0.816 |
| Rv0100 | 0.095 | NA | -0.261 | 0.885 | -0.356 | 0.836 |
| nrp | 0.106 | 0.999 | -0.019 | 0.965 | -0.125 | 0.816 |
| Rv0102 | -0.108 | 0.999 | 0.079 | 0.925 | 0.187 | 0.819 |
| ctpB | 0.117 | 0.999 | 0.131 | 0.859 | 0.014 | 0.977 |
| Rv0104 | 0.601 | 0.999 | 0.305 | 0.848 | -0.296 | 0.822 |
| rpmB1 | -0.090 | NA | -0.513 | 0.895 | -0.423 | 0.896 |
| Rv0106 | -0.094 | 0.999 | -0.058 | 0.927 | 0.036 | 0.942 |
| ctpI | 0.133 | 0.999 | 0.239 | 0.834 | 0.106 | 0.884 |
| Rv0108c | 0.212 | 0.999 | 0.315 | 0.826 | 0.104 | 0.881 |
| PE_PGRS1 | -0.074 | NA | -0.543 | 0.831 | -0.469 | 0.816 |
| Rv0110 | -0.100 | 0.999 | 0.004 | 0.995 | 0.104 | 0.896 |
| Rv0111 | -0.001 | 0.999 | 0.052 | 0.933 | 0.053 | 0.917 |
| gca | 0.305 | 0.999 | -0.152 | 0.866 | -0.456 | 0.816 |
| gmhA | 0.425 | 0.389 | 0.063 | 0.922 | -0.362 | 0.816 |
| gmhB | 0.441 | 0.999 | 0.147 | 0.935 | -0.294 | 0.854 |
| hddA | 0.164 | 0.999 | -0.006 | 0.987 | -0.170 | 0.816 |
| Rv0115a | 0.128 | 0.999 | -0.084 | 0.885 | -0.212 | 0.816 |
| ldtA | -0.147 | 0.999 | -0.210 | 0.826 | -0.063 | 0.896 |
| oxyS | -0.058 | 0.999 | -0.248 | 0.826 | -0.190 | 0.816 |
| oxcA | 0.214 | 0.999 | 0.369 | 0.359 | 0.156 | 0.816 |
| fadD7 | 0.246 | 0.999 | 0.497 | 0.289 | 0.250 | 0.816 |
| fusA2 | 0.053 | 0.999 | 0.103 | 0.879 | 0.049 | 0.917 |
| Rv0121c | -0.050 | 0.999 | -0.013 | 0.975 | 0.037 | 0.936 |
| Rv0122 | -0.180 | NA | 0.162 | 0.902 | 0.342 | 0.816 |
| Rv0123 | 0.165 | NA | 0.247 | 0.854 | 0.082 | 0.932 |
| PE_PGRS2 | -0.205 | NA | -0.693 | 0.834 | -0.487 | 0.837 |
| pepA | 0.067 | 0.999 | 0.304 | 0.831 | 0.237 | 0.817 |
| treS | 0.070 | 0.999 | -0.114 | 0.848 | -0.183 | 0.816 |
| mak | 0.007 | 0.999 | 0.267 | 0.848 | 0.260 | 0.819 |
| Rv0128 | 0.074 | 0.999 | 0.075 | 0.897 | 0.001 | 0.998 |
| fbpC | 0.108 | 0.999 | 0.195 | 0.848 | 0.086 | 0.896 |
| htdZ | -0.036 | NA | -0.237 | 0.848 | -0.201 | 0.836 |
| fadE1 | 0.094 | 0.999 | -0.145 | 0.846 | -0.239 | 0.816 |
| fgd2 | 0.019 | NA | -0.071 | 0.922 | -0.090 | 0.881 |
| Rv0133 | -0.054 | 0.999 | 0.234 | 0.832 | 0.288 | 0.816 |
| ephF | 0.066 | 0.999 | 0.363 | 0.820 | 0.297 | 0.816 |
| Rv0135c | -0.189 | 0.999 | -0.092 | 0.941 | 0.097 | 0.925 |
| cyp138 | -0.207 | 0.999 | -0.073 | 0.911 | 0.134 | 0.829 |
| msrA | 0.064 | 0.999 | -0.309 | 0.831 | -0.374 | 0.816 |
| Rv0138 | 0.094 | 0.999 | -0.229 | 0.830 | -0.322 | 0.816 |
| Rv0139 | 0.057 | 0.999 | 0.134 | 0.895 | 0.076 | 0.930 |
| Rv0140 | -0.109 | 0.999 | -0.497 | 0.795 | -0.388 | 0.816 |
| Rv0141c | 0.067 | 0.999 | 0.088 | 0.879 | 0.021 | 0.961 |
| Rv0142 | -0.046 | 0.999 | 0.719 | 0.659 | 0.765 | 0.816 |
| Rv0143c | -0.058 | 0.999 | 0.307 | 0.831 | 0.365 | 0.816 |
| Rv0144 | 0.013 | 0.999 | 0.093 | 0.895 | 0.081 | 0.894 |
| Rv0145 | -0.042 | 0.999 | 0.170 | 0.832 | 0.212 | 0.816 |
| Rv0146 | -0.173 | 0.999 | -0.183 | 0.826 | -0.010 | 0.985 |
| Rv0147 | -0.006 | 0.999 | -0.107 | 0.879 | -0.101 | 0.862 |
| Rv0148 | -0.162 | 0.999 | -0.141 | 0.938 | 0.020 | 0.989 |
| Rv0149 | -0.101 | 0.999 | -0.178 | 0.820 | -0.076 | 0.843 |
| Rv0150c | -0.212 | NA | -0.595 | 0.834 | -0.383 | 0.846 |
| PE1 | -0.023 | 0.999 | -0.487 | 0.848 | -0.464 | 0.819 |
| PE2 | -0.081 | 0.999 | -0.196 | 0.885 | -0.115 | 0.920 |
| ptbB | -0.052 | 0.999 | -0.199 | 0.826 | -0.147 | 0.816 |
| fadE2 | 0.103 | 0.999 | -0.030 | 0.973 | -0.133 | 0.868 |
| pntAa | 0.097 | 0.999 | 0.144 | 0.831 | 0.047 | 0.900 |
| pntAb | 0.059 | 0.999 | -0.148 | 0.894 | -0.207 | 0.837 |
| pntB | 0.077 | 0.999 | 0.113 | 0.879 | 0.035 | 0.943 |
| Rv0157A | -0.168 | 0.999 | 0.079 | 0.949 | 0.247 | 0.835 |
| Rv0158 | 0.048 | 0.999 | 0.073 | 0.885 | 0.025 | 0.951 |
| PE3 | 0.105 | NA | -0.329 | 0.870 | -0.434 | 0.817 |
| PE4 | -0.032 | NA | -0.341 | 0.881 | -0.309 | 0.868 |
| Rv0161 | -0.062 | 0.999 | 0.126 | 0.922 | 0.188 | 0.866 |
| adhE1 | -0.221 | 0.999 | 0.810 | 0.820 | 1.031 | 0.816 |
| Rv0163 | 0.105 | NA | 0.003 | 0.995 | -0.102 | 0.896 |
| TB18.5 | -0.021 | 0.999 | -0.221 | 0.832 | -0.200 | 0.816 |
| mce1R | 0.342 | 0.221 | 0.443 | 0.006 | 0.101 | 0.822 |
| fadD5 | 0.372 | 0.999 | 0.107 | 0.879 | -0.264 | 0.816 |
| yrbE1A | 0.266 | 0.999 | 0.342 | 0.831 | 0.076 | 0.932 |
| yrbE1B | 0.180 | 0.999 | 0.193 | 0.879 | 0.012 | 0.989 |

| | WT v Δ*coaX* | | Δ*coaX* v complement | | WT v complement | |
|---|---|---|---|---|---|---|
| **GeneID** | **log2[FC]** | **padj** | **log2[FC]** | **padj** | **log2[FC]** | **padj** |
| Rv0804 | 0.040 | 0.999 | 0.275 | 0.831 | 0.235 | 0.816 |
| Rv0805 | -0.147 | 0.999 | -0.126 | 0.843 | 0.020 | 0.957 |
| cpsY | 0.090 | 0.999 | -0.101 | 0.910 | -0.191 | 0.822 |
| Rv0807 | 0.030 | 0.999 | 0.109 | 0.918 | 0.079 | 0.925 |
| purF | -0.066 | 0.999 | 0.007 | 0.989 | 0.073 | 0.896 |
| purM | 0.083 | 0.999 | 0.098 | 0.848 | 0.015 | 0.964 |
| Rv0810c | -0.147 | 0.999 | 0.042 | 0.948 | 0.189 | 0.816 |
| Rv0811c | -0.032 | 0.999 | 0.253 | 0.826 | 0.285 | 0.816 |
| Rv0812 | -0.244 | 0.999 | 0.143 | 0.890 | 0.387 | 0.816 |
| Rv0813c | -0.039 | 0.999 | 0.347 | 0.831 | 0.386 | 0.816 |
| Rv0814c | 0.121 | 0.999 | 0.660 | 0.805 | 0.539 | 0.816 |
| cysA2 | 0.003 | 0.999 | -0.171 | 0.826 | -0.174 | 0.816 |
| thiX | 0.176 | NA | -0.097 | 0.895 | -0.273 | 0.816 |
| Rv0817c | 0.019 | 0.999 | -0.142 | 0.890 | -0.161 | 0.862 |
| Rv0818 | -0.038 | 0.999 | 0.198 | 0.848 | 0.236 | 0.816 |
| mshD | 0.142 | 0.999 | -0.066 | 0.925 | -0.208 | 0.816 |
| phoT | 0.044 | 0.999 | -0.314 | 0.766 | -0.358 | 0.816 |
| phoY2 | -0.096 | 0.999 | -0.233 | 0.826 | -0.137 | 0.819 |
| Rv0822c | 0.000 | 0.999 | -0.167 | 0.843 | -0.168 | 0.816 |
| Rv0823c | -0.051 | 0.999 | -0.166 | 0.848 | -0.115 | 0.852 |
| desA1 | 0.087 | 0.999 | -0.257 | 0.831 | -0.344 | 0.816 |
| Rv0825c | 0.212 | 0.999 | 0.082 | 0.929 | -0.129 | 0.868 |
| Rv0826 | -0.207 | NA | -0.314 | 0.848 | -0.107 | 0.924 |
| kmtR | -0.083 | 0.999 | -0.040 | 0.959 | 0.043 | 0.942 |
| Rv0828c | -0.010 | NA | -0.036 | 0.971 | -0.026 | 0.976 |
| Rv0830 | -0.102 | 0.999 | -0.052 | 0.941 | 0.050 | 0.931 |
| Rv0831c | 0.184 | 0.999 | -0.102 | 0.931 | -0.286 | 0.817 |
| PE_PGRS12 | 0.069 | 0.999 | 0.519 | 0.826 | 0.450 | 0.816 |
| PE_PGRS13 | -0.200 | 0.999 | 0.068 | 0.971 | 0.268 | 0.871 |
| PE_PGRS14 | -0.253 | 0.999 | 0.371 | 0.843 | 0.625 | 0.816 |
| lpqQ | 0.150 | 0.999 | -0.090 | 0.938 | -0.240 | 0.831 |
| Rv0836c | -0.353 | NA | -0.716 | 0.846 | -0.363 | 0.877 |
| Rv0837c | 0.105 | NA | -0.519 | 0.859 | -0.624 | 0.817 |
| lpqR | -0.046 | 0.999 | 0.017 | 0.974 | 0.063 | 0.899 |
| Rv0839 | -0.054 | 0.999 | 0.212 | 0.832 | 0.266 | 0.816 |
| pip | -0.043 | 0.999 | -0.218 | 0.859 | -0.175 | 0.855 |
| Rv0841 | -0.114 | NA | 0.007 | 0.995 | 0.121 | 0.947 |
| Rv0842 | -0.115 | 0.999 | -0.006 | 0.994 | 0.109 | 0.888 |
| Rv0843 | -0.043 | NA | 0.004 | 0.995 | 0.047 | 0.942 |
| narL | 0.008 | 0.999 | 0.231 | 0.879 | 0.223 | 0.859 |
| Rv0845 | -0.088 | 0.999 | 0.078 | 0.927 | 0.165 | 0.835 |
| Rv0846c | -0.572 | 0.999 | -0.505 | 0.804 | 0.068 | 0.942 |
| lpqS | -0.571 | 0.587 | -0.590 | 0.293 | -0.019 | 0.979 |
| cysK2 | -0.649 | 0.010 | -0.747 | 0.000 | -0.098 | 0.863 |
| Rv0849 | -0.692 | 0.002 | -0.809 | 0.000 | -0.117 | 0.837 |
| Rv0851c | 0.059 | 0.999 | 0.234 | 0.829 | 0.175 | 0.817 |
| Rv0852 | 0.030 | 0.999 | 0.202 | 0.858 | 0.172 | 0.842 |
| pdc | 0.135 | 0.999 | 0.045 | 0.926 | -0.090 | 0.837 |
| Rv0854 | 0.181 | 0.999 | -0.097 | 0.962 | -0.278 | 0.868 |
| far | 0.097 | 0.999 | 0.025 | 0.973 | -0.072 | 0.909 |
| Rv0856 | -0.005 | 0.999 | -0.217 | 0.879 | -0.213 | 0.855 |
| Rv0857 | 0.075 | 0.999 | 0.077 | 0.903 | 0.002 | 0.997 |
| dapC | 0.115 | 0.999 | 0.169 | 0.826 | 0.054 | 0.894 |
| Rv0859 | 0.133 | 0.999 | 0.266 | 0.442 | 0.133 | 0.816 |
| fadB | 0.050 | 0.999 | 0.138 | 0.859 | 0.088 | 0.871 |
| ercc3 | 0.052 | 0.999 | -0.114 | 0.859 | -0.166 | 0.816 |
| Rv0862c | 0.056 | 0.999 | 0.190 | 0.848 | 0.134 | 0.846 |
| Rv0863 | -0.064 | 0.999 | -0.470 | 0.510 | -0.406 | 0.816 |
| moaC2 | 0.068 | 0.999 | -0.245 | 0.834 | -0.313 | 0.816 |
| mog | -0.156 | 0.999 | 0.086 | 0.951 | 0.242 | 0.849 |
| moaE2 | -0.069 | 0.999 | 0.250 | 0.820 | 0.318 | 0.816 |
| rpfA | -0.141 | 0.999 | -0.080 | 0.912 | 0.061 | 0.917 |
| moaD2 | -0.023 | 0.999 | -0.374 | 0.834 | -0.351 | 0.816 |
| moaA2 | -0.009 | 0.999 | 0.132 | 0.895 | 0.141 | 0.868 |
| Rv0870c | 0.008 | 0.999 | -0.441 | 0.588 | -0.449 | 0.816 |
| cspB | -0.099 | 0.999 | 0.327 | 0.832 | 0.426 | 0.816 |
| PE_PGRS15 | -0.166 | 0.999 | -0.263 | 0.859 | -0.097 | 0.930 |
| fadE10 | 0.053 | 0.999 | -0.036 | 0.927 | -0.089 | 0.819 |
| Rv0874c | -0.213 | 0.999 | 0.091 | 0.879 | 0.304 | 0.816 |
| Rv0875c | 0.033 | 0.999 | -0.180 | 0.855 | -0.213 | 0.816 |
| Rv0876c | -0.047 | 0.999 | 0.116 | 0.848 | 0.163 | 0.816 |
| Rv0877 | 0.052 | 0.999 | 0.202 | 0.859 | 0.150 | 0.862 |
| PPE13 | 0.039 | 0.999 | -0.162 | 0.924 | -0.200 | 0.884 |
| Rv0879c | 0.025 | 0.999 | 0.222 | 0.848 | 0.197 | 0.835 |
| Rv0880 | -0.131 | 0.999 | 0.183 | 0.879 | 0.314 | 0.816 |
| Rv0881 | 0.016 | NA | -0.071 | 0.930 | -0.088 | 0.894 |
| Rv0882 | 0.240 | NA | 0.554 | 0.510 | 0.314 | 0.816 |
| Rv0883c | 0.134 | 0.999 | 0.232 | 0.826 | 0.097 | 0.862 |
| serC | -0.005 | 0.999 | -0.131 | 0.834 | -0.126 | 0.816 |
| Rv0885 | 0.004 | 0.999 | -0.150 | 0.848 | -0.154 | 0.817 |
| fprB | -0.143 | 0.999 | 0.331 | 0.831 | 0.475 | 0.816 |
| Rv0887c | 0.009 | NA | -0.104 | 0.958 | -0.112 | 0.942 |
| Rv0888 | 0.290 | 0.999 | -0.126 | 0.892 | -0.416 | 0.816 |
| citA | -0.104 | 0.999 | 0.198 | 0.826 | 0.302 | 0.816 |
| Rv0890c | 0.070 | 0.999 | 0.235 | 0.522 | 0.165 | 0.816 |
| Rv0891c | 0.093 | 0.999 | -0.254 | 0.859 | -0.347 | 0.816 |
| Rv0892 | -0.006 | 0.999 | -0.155 | 0.879 | -0.149 | 0.855 |
| Rv0893c | -0.099 | 0.999 | -0.085 | 0.927 | 0.014 | 0.986 |
| Rv0894 | -0.260 | NA | -0.789 | 0.831 | -0.529 | 0.835 |
| Rv0895 | -0.215 | NA | -0.157 | 0.912 | 0.059 | 0.956 |
| gltA2 | 0.006 | 0.999 | -0.136 | 0.832 | -0.142 | 0.816 |
| Rv0897c | -0.054 | 0.999 | 0.026 | 0.931 | 0.080 | 0.816 |
| Rv0898c | -0.002 | 0.999 | -0.123 | 0.941 | -0.121 | 0.930 |
| arfA | 0.062 | 0.999 | 0.073 | 0.935 | 0.011 | 0.988 |
| arfB | -0.022 | NA | -0.335 | 0.843 | -0.313 | 0.817 |
| arfC | 0.095 | 0.999 | -0.014 | 0.979 | -0.109 | 0.862 |
| prrB | 0.040 | 0.999 | -0.209 | 0.825 | -0.249 | 0.816 |
| prrA | 0.003 | 0.999 | 0.083 | 0.936 | 0.080 | 0.925 |
| accD3 | 0.003 | 0.999 | 0.248 | 0.829 | 0.245 | 0.816 |
| echA6 | -0.211 | 0.999 | -0.444 | 0.688 | -0.233 | 0.816 |
| Rv0906 | -0.268 | 0.999 | -0.020 | 0.985 | 0.248 | 0.837 |
| Rv0907 | -0.163 | 0.999 | -0.236 | 0.825 | -0.073 | 0.883 |
| ctpE | -0.051 | 0.999 | -0.157 | 0.831 | -0.106 | 0.835 |
| Rv0909 | -0.067 | 0.999 | -0.379 | 0.857 | -0.312 | 0.846 |
| Rv0910 | 0.045 | 0.999 | -0.095 | 0.927 | -0.140 | 0.868 |
| Rv0911 | -0.022 | 0.999 | 0.046 | 0.970 | 0.068 | 0.947 |
| Rv0912 | -0.097 | 0.999 | 0.304 | 0.820 | 0.401 | 0.816 |
| Rv0913c | 0.098 | 0.999 | -0.040 | 0.970 | -0.138 | 0.882 |
| Rv0914c | -0.005 | 0.999 | 0.194 | 0.801 | 0.199 | 0.816 |
| PPE14 | 0.030 | NA | -0.122 | 0.892 | -0.152 | 0.852 |
| PE7 | -0.145 | NA | -0.384 | 0.826 | -0.239 | 0.819 |
| betP | -0.004 | 0.999 | -0.160 | 0.853 | -0.156 | 0.828 |
| Rv0918 | 0.090 | NA | -0.091 | 0.922 | -0.181 | 0.835 |
| Rv0919 | -0.014 | NA | -0.293 | 0.859 | -0.279 | 0.836 |
| Rv0920c | 0.106 | 0.999 | -0.123 | 0.948 | -0.229 | 0.879 |
| Rv0921 | -0.184 | 0.999 | -0.295 | 0.912 | -0.111 | 0.955 |
| Rv0922 | -0.013 | 0.999 | 0.026 | 0.956 | 0.039 | 0.918 |
| Rv0923c | 0.028 | 0.999 | 0.004 | 0.994 | -0.025 | 0.955 |
| mntH | 0.056 | NA | 0.125 | 0.895 | 0.068 | 0.930 |
| Rv0925c | 0.066 | 0.999 | -0.091 | 0.862 | -0.156 | 0.816 |
| Rv0926c | 0.103 | 0.999 | 0.103 | 0.863 | 0.001 | 0.999 |
| Rv0927c | 0.006 | 0.999 | -0.044 | 0.936 | -0.050 | 0.908 |
| pstS3 | 0.045 | 0.999 | -0.136 | 0.922 | -0.180 | 0.868 |
| pstC2 | 0.142 | 0.999 | 0.068 | 0.938 | -0.074 | 0.917 |
| pstA1 | 0.037 | 0.999 | -0.038 | 0.962 | -0.076 | 0.904 |
| pknD | -0.064 | 0.999 | -0.113 | 0.859 | -0.048 | 0.919 |
| pstS2 | -0.030 | 0.999 | -0.201 | 0.826 | -0.171 | 0.816 |
| pstB | 0.129 | 0.999 | -0.061 | 0.940 | -0.190 | 0.817 |
| pstS1 | 0.121 | 0.999 | -0.054 | 0.927 | -0.175 | 0.816 |
| pstC1 | 0.232 | 0.999 | 0.194 | 0.854 | -0.038 | 0.956 |
| pstA2 | -0.002 | 0.999 | 0.055 | 0.933 | 0.057 | 0.916 |
| mku | -0.010 | 0.999 | -0.175 | 0.897 | -0.164 | 0.887 |
| ligD | 0.091 | 0.999 | 0.175 | 0.855 | 0.084 | 0.897 |
| Rv0939 | -0.027 | 0.999 | 0.182 | 0.831 | 0.209 | 0.816 |
| Rv0940c | -0.040 | 0.999 | -0.157 | 0.859 | -0.117 | 0.865 |
| Rv0941c | -0.093 | 0.999 | -0.082 | 0.906 | 0.011 | 0.986 |
| Rv0942 | 0.099 | NA | 0.091 | 0.962 | -0.008 | 0.996 |
| Rv0943c | -0.297 | NA | -0.424 | 0.858 | -0.127 | 0.939 |
| Rv0944 | 0.030 | NA | 0.652 | 0.831 | 0.621 | 0.816 |
| Rv0945 | -0.002 | 0.999 | 0.125 | 0.831 | 0.127 | 0.816 |
| pgi | -0.020 | 0.999 | -0.152 | 0.831 | -0.132 | 0.816 |
| Rv0948c | -0.058 | 0.999 | 0.391 | 0.826 | 0.449 | 0.816 |
| uvrD1 | 0.095 | 0.999 | 0.052 | 0.895 | -0.043 | 0.897 |
| Rv0950c | -0.047 | 0.999 | -0.125 | 0.879 | -0.079 | 0.892 |
| sucC | -0.061 | 0.999 | -0.305 | 0.767 | -0.245 | 0.816 |
| sucD | -0.109 | 0.999 | -0.143 | 0.831 | -0.033 | 0.930 |
| Rv0953c | 0.035 | NA | -0.036 | 0.975 | -0.071 | 0.954 |
| Rv0954 | 0.140 | 0.999 | 0.185 | 0.883 | 0.045 | 0.962 |
| Rv0955 | 0.065 | 0.999 | 0.337 | 0.831 | 0.272 | 0.817 |
| purN | 0.081 | 0.999 | -0.248 | 0.826 | -0.329 | 0.816 |
| purH | 0.074 | 0.999 | 0.144 | 0.826 | 0.069 | 0.852 |
| Rv0958 | 0.069 | 0.999 | 0.205 | 0.832 | 0.135 | 0.837 |
| Rv0959 | 0.108 | 0.999 | 0.114 | 0.879 | 0.006 | 0.993 |
| vapB9 | -0.236 | NA | 0.116 | 0.931 | 0.352 | 0.816 |
| vapC9 | 0.012 | 0.999 | -0.219 | 0.879 | -0.231 | 0.838 |
| Rv0961 | -0.037 | 0.999 | 0.097 | 0.879 | 0.134 | 0.831 |
| lprP | -0.390 | NA | -1.642 | NA | -1.251 | NA |
| Rv0963c | -0.150 | NA | -0.532 | 0.826 | -0.382 | 0.816 |
| Rv0964c | 0.130 | 0.999 | -0.178 | 0.848 | -0.308 | 0.816 |
| Rv0965c | -0.017 | NA | 0.140 | 0.893 | 0.157 | 0.862 |
| Rv0966c | -0.091 | 0.999 | 0.187 | 0.843 | 0.278 | 0.816 |
| csoR | -0.337 | 0.999 | -0.777 | 0.716 | -0.440 | 0.816 |
| Rv0968 | -0.310 | 0.999 | -0.626 | 0.001 | -0.316 | 0.816 |
| ctpV | -0.366 | 0.999 | -0.183 | 0.888 | 0.183 | 0.868 |
| Rv0970 | -0.296 | 0.999 | -0.242 | 0.826 | 0.054 | 0.925 |
| echA7 | 0.153 | NA | 0.566 | 0.608 | 0.414 | 0.816 |
| fadE12 | 0.021 | 0.999 | -0.181 | 0.879 | -0.202 | 0.852 |
| accA2 | -0.215 | 0.999 | -0.082 | 0.899 | 0.133 | 0.835 |
| accD2 | -0.089 | 0.999 | -0.177 | 0.848 | -0.089 | 0.888 |
| fadE13 | -0.161 | NA | 0.023 | 0.973 | 0.184 | 0.816 |
| Rv0976c | -0.147 | 0.999 | 0.120 | 0.879 | 0.268 | 0.816 |
| PE_PGRS16 | 0.086 | 0.999 | -0.254 | 0.879 | -0.340 | 0.831 |
| PE_PGRS17 | -0.165 | NA | -0.283 | 0.907 | -0.118 | 0.950 |

| | WT v Δ*coaX* | | Δ*coaX* v complement | | WT v complement | |
|---|---|---|---|---|---|---|
| **GeneID** | **log2[FC]** | **padj** | **log2[FC]** | **padj** | **log2[FC]** | **padj** |
| tesB1 | 0.057 | 0.999 | 0.281 | 0.879 | 0.224 | 0.879 |
| Rv1619 | 0.061 | 0.999 | 0.281 | 0.848 | 0.220 | 0.836 |
| cydC | -0.102 | 0.999 | 0.147 | 0.868 | 0.249 | 0.816 |
| cydD | -0.208 | 0.999 | 0.109 | 0.946 | 0.318 | 0.831 |
| cydB | -0.221 | 0.999 | -0.321 | 0.825 | -0.100 | 0.882 |
| cydA | -0.072 | 0.999 | -0.434 | 0.657 | -0.362 | 0.816 |
| Rv1624c | -0.045 | 0.999 | 0.077 | 0.948 | 0.122 | 0.895 |
| cya | -0.043 | 0.999 | -0.011 | 0.973 | 0.032 | 0.910 |
| Rv1626 | 0.045 | 0.999 | -0.045 | 0.948 | -0.090 | 0.868 |
| Rv1627c | 0.023 | 0.999 | 0.269 | 0.826 | 0.246 | 0.816 |
| Rv1628c | -0.086 | 0.999 | 0.077 | 0.936 | 0.163 | 0.852 |
| polA | 0.031 | 0.999 | 0.033 | 0.922 | 0.003 | 0.994 |
| rpsA | 0.072 | 0.999 | -0.037 | 0.948 | -0.108 | 0.836 |
| coaE | 0.117 | 0.999 | 0.011 | 0.984 | -0.106 | 0.865 |
| Rv1632c | 0.083 | 0.999 | 0.476 | 0.826 | 0.394 | 0.816 |
| uvrB | -0.002 | 0.999 | -0.123 | 0.831 | -0.121 | 0.816 |
| Rv1634 | -0.008 | 0.999 | 0.321 | 0.831 | 0.328 | 0.816 |
| Rv1635c | -0.077 | 0.999 | 0.042 | 0.938 | 0.119 | 0.819 |
| TB15.3 | -0.236 | 0.999 | -0.406 | 0.803 | -0.170 | 0.838 |
| Rv1637c | 0.016 | 0.999 | -0.060 | 0.922 | -0.076 | 0.877 |
| uvrA | -0.033 | 0.999 | -0.084 | 0.901 | -0.051 | 0.930 |
| Rv1638A | 0.107 | 0.999 | 0.076 | 0.936 | -0.031 | 0.967 |
| Rv1639c | 0.068 | 0.999 | 0.150 | 0.879 | 0.082 | 0.918 |
| lysX | 0.201 | 0.999 | -0.082 | 0.905 | -0.283 | 0.816 |
| infC | 0.012 | 0.999 | -0.112 | 0.948 | -0.124 | 0.932 |
| rpmI | 0.138 | 0.999 | -0.290 | 0.843 | -0.428 | 0.816 |
| rplT | 0.052 | 0.999 | -0.466 | 0.795 | -0.518 | 0.816 |
| tsnR | -0.011 | 0.999 | 0.153 | 0.885 | 0.164 | 0.862 |
| Rv1645c | 0.091 | 0.999 | -0.151 | 0.886 | -0.242 | 0.819 |
| PE17 | -0.026 | 0.999 | 0.460 | 0.820 | 0.486 | 0.816 |
| Rv1647 | -0.141 | 0.999 | 0.130 | 0.918 | 0.270 | 0.819 |
| Rv1648 | 0.051 | 0.999 | 0.090 | 0.905 | 0.038 | 0.947 |
| pheS | 0.004 | 0.999 | -0.116 | 0.879 | -0.119 | 0.838 |
| pheT | 0.018 | 0.999 | 0.305 | 0.848 | 0.287 | 0.819 |
| PE_PGRS30 | -0.011 | 0.999 | 0.060 | 0.962 | 0.071 | 0.942 |
| argC | -0.055 | 0.999 | -0.055 | 0.946 | 0.000 | 0.999 |
| argJ | -0.216 | 0.999 | -0.475 | 0.054 | -0.259 | 0.816 |
| argB | -0.187 | 0.999 | -0.030 | 0.975 | 0.157 | 0.884 |
| argD | -0.421 | 0.303 | -0.433 | 0.144 | -0.011 | 0.983 |
| argF | -0.412 | 0.999 | -0.254 | 0.859 | 0.157 | 0.879 |
| argR | -0.351 | NA | 0.121 | 0.948 | 0.472 | 0.817 |
| argG | -0.102 | 0.999 | -0.135 | 0.832 | -0.033 | 0.936 |
| argH | -0.116 | 0.999 | 0.361 | 0.834 | 0.477 | 0.816 |
| pks10 | -0.115 | 0.999 | -0.031 | 0.959 | 0.084 | 0.865 |
| pks7 | -0.019 | 0.999 | -0.319 | 0.879 | -0.300 | 0.860 |
| pks8 | -0.134 | 0.999 | -0.456 | 0.883 | -0.322 | 0.896 |
| pks17 | -0.162 | 0.999 | -0.620 | 0.832 | -0.458 | 0.831 |
| pks9 | 0.116 | 0.999 | -0.414 | 0.832 | -0.530 | 0.816 |
| pks11 | 0.054 | 0.999 | 0.131 | 0.895 | 0.077 | 0.925 |
| cyp139 | 0.137 | 0.999 | 0.021 | 0.970 | -0.115 | 0.830 |
| Rv1669 | -0.340 | NA | -1.674 | NA | -1.334 | NA |
| Rv1670 | 0.002 | NA | -0.220 | 0.879 | -0.222 | 0.852 |
| Rv1671 | -0.035 | NA | -0.384 | 0.831 | -0.349 | 0.816 |
| Rv1672c | -0.030 | 0.999 | 0.472 | 0.772 | 0.502 | 0.816 |
| Rv1673c | -0.081 | NA | -0.283 | 0.832 | -0.202 | 0.835 |
| Rv1674c | -0.105 | NA | -0.490 | 0.832 | -0.384 | 0.822 |
| cmr | -0.061 | NA | -0.207 | 0.859 | -0.146 | 0.868 |
| Rv1676 | 0.130 | 0.999 | -0.143 | 0.886 | -0.273 | 0.816 |
| dsbF | -0.035 | 0.999 | 0.214 | 0.853 | 0.249 | 0.816 |
| Rv1678 | 0.041 | 0.999 | 0.382 | 0.831 | 0.340 | 0.816 |
| fadE16 | -0.029 | 0.999 | 0.062 | 0.936 | 0.091 | 0.884 |
| Rv1680 | 0.190 | 0.999 | -0.019 | 0.975 | -0.209 | 0.816 |
| moeX | -0.043 | 0.999 | 0.170 | 0.848 | 0.212 | 0.816 |
| Rv1682 | -0.162 | 0.999 | -0.370 | 0.820 | -0.208 | 0.819 |
| Rv1683 | -0.087 | 0.999 | -0.146 | 0.845 | -0.059 | 0.896 |
| Rv1684 | -0.040 | 0.999 | -0.335 | 0.767 | -0.296 | 0.816 |
| Rv1685c | -0.313 | NA | -0.480 | 0.826 | -0.167 | 0.881 |
| Rv1686c | -0.130 | NA | -0.482 | 0.767 | -0.352 | 0.816 |
| Rv1687c | -0.215 | 0.999 | -0.137 | 0.936 | 0.078 | 0.954 |
| mpg | -0.249 | NA | 0.194 | 0.922 | 0.443 | 0.819 |
| tyrS | 0.080 | 0.999 | 0.041 | 0.947 | -0.039 | 0.936 |
| lprJ | 0.146 | 0.999 | 0.165 | 0.862 | 0.019 | 0.978 |
| Rv1691 | -0.075 | 0.999 | 0.249 | 0.826 | 0.324 | 0.816 |
| Rv1692 | -0.022 | 0.999 | 0.261 | 0.859 | 0.283 | 0.825 |
| Rv1693 | -0.119 | 0.999 | 0.029 | 0.977 | 0.147 | 0.894 |
| tlyA | 0.159 | 0.999 | 0.410 | 0.396 | 0.251 | 0.816 |
| ppnK | 0.021 | 0.999 | 0.086 | 0.856 | 0.065 | 0.858 |
| recN | 0.027 | 0.999 | 0.148 | 0.879 | 0.120 | 0.868 |
| Rv1697 | 0.165 | 0.999 | 0.053 | 0.930 | -0.112 | 0.837 |
| mctB | 0.187 | 0.999 | 0.129 | 0.831 | -0.058 | 0.868 |
| pyrG | 0.033 | 0.999 | -0.150 | 0.832 | -0.183 | 0.816 |
| Rv1700 | -0.001 | 0.999 | -0.008 | 0.987 | -0.007 | 0.988 |
| Rv1701 | 0.017 | 0.999 | 0.136 | 0.895 | 0.119 | 0.889 |
| Rv1702c | 0.046 | 0.999 | -0.694 | 0.866 | -0.740 | 0.835 |
| Rv1703c | 0.052 | 0.999 | 0.032 | 0.940 | -0.020 | 0.951 |
| cycA | 0.139 | 0.999 | -0.015 | 0.974 | -0.155 | 0.816 |
| PPE22 | -0.073 | NA | -0.173 | 0.879 | -0.100 | 0.913 |
| Rv1706A | -0.290 | NA | -0.513 | 0.878 | -0.223 | 0.924 |
| PPE23 | -0.184 | NA | 0.189 | 0.879 | 0.373 | 0.816 |
| Rv1707 | -0.097 | 0.999 | -0.045 | 0.951 | 0.052 | 0.932 |
| Rv1708 | 0.161 | 0.999 | 0.133 | 0.826 | -0.029 | 0.932 |
| scpA | 0.292 | 0.999 | 0.232 | 0.831 | -0.060 | 0.926 |
| scpB | 0.076 | 0.999 | -0.062 | 0.950 | -0.138 | 0.868 |
| Rv1711 | 0.166 | 0.999 | 0.039 | 0.958 | -0.127 | 0.846 |
| cmk | 0.206 | 0.999 | 0.405 | 0.706 | 0.199 | 0.817 |
| engA | 0.099 | 0.999 | 0.326 | 0.380 | 0.227 | 0.816 |
| Rv1714 | -0.195 | 0.999 | 0.595 | 0.805 | 0.790 | 0.816 |
| fadB3 | 0.116 | 0.999 | 0.403 | 0.820 | 0.287 | 0.816 |
| Rv1716 | 0.168 | 0.999 | 0.033 | 0.962 | -0.135 | 0.837 |
| Rv1717 | 0.198 | NA | 0.097 | 0.948 | -0.101 | 0.932 |
| Rv1718 | 0.140 | 0.999 | 0.325 | 0.613 | 0.184 | 0.816 |
| Rv1719 | 0.130 | 0.999 | 0.103 | 0.927 | -0.027 | 0.977 |
| vapC12 | 0.130 | NA | -0.067 | 0.934 | -0.198 | 0.817 |
| vapB12 | -0.491 | NA | -0.196 | 0.922 | 0.294 | 0.857 |
| Rv1722 | 0.111 | 0.999 | -0.173 | 0.859 | -0.284 | 0.816 |
| Rv1723 | -0.016 | NA | 0.373 | 0.772 | 0.389 | 0.816 |
| Rv1724c | 0.156 | 0.999 | -0.254 | 0.853 | -0.409 | 0.816 |
| Rv1725c | -0.005 | 0.999 | -0.087 | 0.856 | -0.082 | 0.835 |
| Rv1726 | -0.129 | NA | -0.425 | 0.879 | -0.297 | 0.885 |
| Rv1727 | -0.504 | NA | 0.193 | 0.927 | 0.697 | 0.816 |
| Rv1728c | 0.050 | 0.999 | 0.051 | 0.914 | 0.002 | 0.997 |
| Rv1729c | 0.026 | 0.999 | -0.115 | 0.848 | -0.142 | 0.816 |
| Rv1730c | -0.087 | 0.999 | -0.216 | 0.831 | -0.129 | 0.838 |
| gabD2 | -0.120 | 0.999 | 0.014 | 0.977 | 0.134 | 0.835 |
| Rv1732c | -0.173 | 0.999 | 0.048 | 0.958 | 0.222 | 0.816 |
| Rv1733c | -0.544 | 0.398 | -0.752 | 0.008 | -0.208 | 0.816 |
| Rv1734c | -0.355 | NA | -0.785 | 0.172 | -0.430 | 0.816 |
| Rv1735a | -0.626 | 0.999 | -0.844 | 0.589 | -0.218 | 0.866 |
| Rv1735c | -0.368 | NA | -0.709 | 0.000 | -0.341 | 0.816 |
| narX | -0.469 | 0.999 | -0.545 | 0.659 | -0.076 | 0.930 |
| narK2 | -0.525 | 0.999 | -0.519 | 0.820 | 0.006 | 0.996 |
| Rv1738 | -0.521 | 0.374 | -0.811 | 0.001 | -0.290 | 0.816 |
| Rv1739c | 0.020 | 0.999 | -0.209 | 0.856 | -0.229 | 0.819 |
| vapB34 | -0.029 | NA | -0.304 | 0.848 | -0.274 | 0.831 |
| vapC34 | -0.134 | NA | -0.427 | 0.826 | -0.293 | 0.817 |
| Rv1742 | -0.106 | 0.999 | 0.095 | 0.895 | 0.200 | 0.816 |
| pknE | 0.020 | 0.999 | -0.045 | 0.879 | -0.065 | 0.819 |
| Rv1744c | -0.238 | NA | -0.632 | 0.831 | -0.394 | 0.837 |
| idi | -0.121 | NA | -0.264 | 0.848 | -0.143 | 0.868 |
| pknF | 0.081 | 0.999 | 0.070 | 0.895 | -0.012 | 0.979 |
| Rv1747 | -0.013 | 0.999 | 0.054 | 0.931 | 0.067 | 0.896 |
| Rv1748 | -0.051 | 0.999 | 0.205 | 0.834 | 0.256 | 0.816 |
| Rv1749c | -0.051 | 0.999 | 0.012 | 0.983 | 0.062 | 0.917 |
| fadD1 | -0.011 | 0.999 | -0.115 | 0.879 | -0.105 | 0.868 |
| Rv1751 | -0.004 | 0.999 | 0.059 | 0.883 | 0.062 | 0.859 |
| Rv1752 | 0.008 | 0.999 | 0.152 | 0.854 | 0.144 | 0.833 |
| PPE24 | 0.211 | 0.999 | -0.368 | 0.879 | -0.579 | 0.817 |
| Rv1754c | 0.149 | 0.999 | -0.043 | 0.973 | -0.193 | 0.868 |
| cut1 | -0.426 | NA | -0.942 | 0.832 | -0.516 | 0.862 |
| wag22 | -0.174 | 0.999 | -0.136 | 0.942 | 0.038 | 0.981 |
| Rv1760 | -0.040 | 0.999 | 0.003 | 0.995 | 0.043 | 0.942 |
| Rv1761c | -0.102 | NA | -0.449 | 0.820 | -0.347 | 0.816 |
| Rv1762c | 0.211 | 0.999 | -0.060 | 0.972 | -0.272 | 0.865 |
| Rv1765c | -0.055 | 0.999 | -0.248 | 0.832 | -0.193 | 0.819 |
| Rv1766 | 0.126 | 0.999 | -0.070 | 0.927 | -0.197 | 0.816 |
| Rv1767 | -0.009 | 0.999 | 0.152 | 0.879 | 0.161 | 0.852 |
| PE_PGRS31 | -0.013 | 0.999 | 0.230 | 0.879 | 0.242 | 0.852 |
| Rv1769 | 0.177 | 0.999 | 0.056 | 0.948 | -0.121 | 0.866 |
| Rv1770 | 0.150 | 0.999 | 0.195 | 0.879 | 0.045 | 0.962 |
| Rv1771 | 0.125 | 0.999 | -0.114 | 0.848 | -0.239 | 0.816 |
| Rv1772 | -0.243 | 0.413 | -0.026 | 0.946 | 0.217 | 0.816 |
| Rv1773c | -0.354 | 0.999 | -0.590 | 0.359 | -0.236 | 0.816 |
| Rv1774 | -0.162 | 0.999 | -0.182 | 0.795 | -0.021 | 0.950 |
| Rv1775 | -0.074 | 0.999 | -0.029 | 0.946 | 0.045 | 0.892 |
| Rv1776c | 0.058 | NA | -0.195 | 0.879 | -0.253 | 0.831 |
| cyp144 | -0.141 | NA | -0.174 | 0.895 | -0.033 | 0.976 |
| Rv1778c | -0.308 | 0.999 | -0.210 | 0.839 | 0.098 | 0.881 |
| Rv1779c | -0.384 | 0.999 | -0.197 | 0.859 | 0.187 | 0.836 |
| Rv1780 | 0.129 | 0.999 | 0.142 | 0.846 | 0.013 | 0.977 |
| malQ | -0.081 | 0.999 | 0.156 | 0.859 | 0.237 | 0.816 |
| eccB5 | 0.044 | 0.999 | -0.143 | 0.848 | -0.188 | 0.816 |
| eccC5 | -0.017 | 0.999 | -0.033 | 0.948 | -0.016 | 0.967 |
| cyp143 | 0.009 | 0.999 | -0.048 | 0.944 | -0.057 | 0.918 |
| Rv1786 | 0.061 | 0.999 | -0.130 | 0.885 | -0.191 | 0.829 |
| PPE25 | 0.020 | 0.999 | -0.065 | 0.940 | -0.086 | 0.898 |
| PE18 | 0.107 | NA | -0.073 | 0.946 | -0.180 | 0.846 |
| PPE26 | -0.034 | 0.999 | 0.264 | 0.826 | 0.298 | 0.816 |
| PPE27 | 0.023 | 0.999 | 0.089 | 0.918 | 0.066 | 0.924 |
| PE19 | 0.173 | 0.999 | -0.023 | 0.989 | -0.196 | 0.909 |
| esxN | 0.073 | 0.999 | -0.361 | 0.831 | -0.434 | 0.816 |
| Rv1794 | 0.030 | 0.999 | -0.132 | 0.895 | -0.162 | 0.862 |
| eccD5 | 0.072 | 0.999 | 0.118 | 0.895 | 0.045 | 0.950 |
| mycP5 | 0.014 | 0.999 | 0.038 | 0.970 | 0.024 | 0.977 |

| | WT v Δ*coaX* | | Δ*coaX* v complement | | WT v complement | |
|---|---|---|---|---|---|---|
| **GeneID** | **log2[FC]** | **padj** | **log2[FC]** | **padj** | **log2[FC]** | **padj** |
| Rv2435c | 0.031 | 0.999 | -0.256 | 0.858 | -0.286 | 0.819 |
| rbsK | -0.119 | 0.999 | 0.169 | 0.834 | 0.288 | 0.816 |
| Rv2437 | 0.122 | NA | -0.165 | 0.890 | -0.287 | 0.819 |
| Rv2438A | -0.912 | NA | -2.421 | 0.772 | -1.509 | 0.816 |
| nadE | 0.030 | 0.999 | -0.059 | 0.895 | -0.090 | 0.835 |
| proB | -0.040 | 0.999 | 0.441 | 0.826 | 0.481 | 0.816 |
| obg | 0.013 | 0.999 | 0.222 | 0.826 | 0.209 | 0.816 |
| rpmA | 0.018 | 0.999 | 0.319 | 0.939 | 0.301 | 0.930 |
| rplU | -0.146 | 0.999 | -0.243 | 0.935 | -0.097 | 0.967 |
| dctA | -0.014 | 0.999 | 0.198 | 0.869 | 0.212 | 0.835 |
| rne | 0.003 | 0.999 | 0.110 | 0.853 | 0.108 | 0.829 |
| ndkA | 0.087 | 0.999 | 0.145 | 0.879 | 0.058 | 0.936 |
| Rv2446c | 0.050 | NA | -0.179 | 0.879 | -0.229 | 0.819 |
| folC | -0.001 | 0.999 | 0.114 | 0.902 | 0.115 | 0.882 |
| valS | -0.106 | 0.999 | 0.072 | 0.931 | 0.178 | 0.829 |
| Rv2449c | 0.011 | 0.999 | 0.078 | 0.885 | 0.066 | 0.882 |
| rpfE | -0.631 | 0.641 | -0.319 | 0.831 | 0.312 | 0.816 |
| Rv2451 | -0.171 | NA | -0.356 | 0.879 | -0.185 | 0.922 |
| Rv2452c | -0.259 | 0.999 | -0.571 | 0.859 | -0.312 | 0.892 |
| mobA | -0.032 | 0.999 | -0.115 | 0.890 | -0.083 | 0.901 |
| Rv2454c | -0.090 | 0.999 | -0.286 | 0.555 | -0.196 | 0.816 |
| Rv2455c | -0.219 | 0.999 | -0.165 | 0.848 | 0.053 | 0.924 |
| Rv2456c | 0.128 | 0.999 | 0.053 | 0.895 | -0.075 | 0.844 |
| clpX | 0.003 | 0.999 | -0.142 | 0.836 | -0.146 | 0.816 |
| mmuM | -0.050 | 0.999 | 0.067 | 0.941 | 0.117 | 0.871 |
| Rv2459 | 0.120 | 0.999 | 0.115 | 0.855 | -0.005 | 0.992 |
| clpP2 | 0.021 | 0.999 | 0.007 | 0.988 | -0.014 | 0.978 |
| clpP1 | 0.001 | 0.999 | -0.142 | 0.856 | -0.143 | 0.829 |
| tig | 0.045 | 0.999 | -0.042 | 0.933 | -0.088 | 0.846 |
| lipP | -0.097 | 0.999 | 2.418 | 0.000 | 2.514 | 0.000 |
| Rv2464c | -0.070 | 0.999 | -0.159 | 0.879 | -0.089 | 0.917 |
| rpiB | -0.026 | 0.999 | -0.352 | 0.879 | -0.326 | 0.856 |
| Rv2466c | -0.258 | 0.999 | -0.305 | 0.832 | -0.047 | 0.954 |
| pepN | 0.085 | 0.999 | 0.028 | 0.970 | -0.057 | 0.930 |
| Rv2468A | 0.175 | 0.999 | -0.331 | 0.769 | -0.506 | 0.816 |
| Rv2468c | 0.086 | 0.999 | 0.301 | 0.848 | 0.215 | 0.855 |
| Rv2469c | 0.134 | 0.999 | 0.455 | 0.795 | 0.322 | 0.816 |
| glbO | 0.035 | 0.999 | -0.385 | 0.728 | -0.420 | 0.816 |
| aglA | -0.097 | 0.999 | 0.013 | 0.982 | 0.110 | 0.866 |
| Rv2472 | 0.085 | 0.999 | -0.558 | 0.848 | -0.643 | 0.816 |
| Rv2473 | 0.038 | 0.999 | -0.074 | 0.934 | -0.112 | 0.879 |
| Rv2474c | -0.065 | 0.999 | 0.274 | 0.831 | 0.339 | 0.816 |
| Rv2475c | 0.168 | 0.999 | 0.059 | 0.948 | -0.108 | 0.889 |
| gdh | -0.014 | 0.999 | -0.042 | 0.962 | -0.027 | 0.969 |
| Rv2477c | 0.102 | 0.999 | 0.090 | 0.883 | -0.012 | 0.983 |
| Rv2478c | -0.005 | 0.999 | 0.115 | 0.930 | 0.120 | 0.907 |
| Rv2481c | -0.080 | NA | 0.682 | 0.801 | 0.761 | 0.816 |
| plsB2 | 0.066 | 0.999 | 0.052 | 0.942 | -0.014 | 0.981 |
| plsC | -0.010 | 0.999 | 0.006 | 0.987 | 0.016 | 0.963 |
| Rv2484c | -0.005 | 0.999 | 0.198 | 0.848 | 0.203 | 0.816 |
| lipQ | 0.492 | 0.096 | 0.839 | 0.000 | 0.347 | 0.816 |
| echA14 | -0.141 | 0.999 | 0.049 | 0.971 | 0.190 | 0.875 |
| PE_PGRS42 | -0.153 | NA | -1.124 | 0.826 | -0.971 | 0.816 |
| Rv2488c | 0.087 | 0.999 | -0.011 | 0.975 | -0.098 | 0.835 |
| Rv2489c | -0.309 | NA | 0.019 | 0.986 | 0.328 | 0.817 |
| Rv2490a | -0.211 | NA | -0.647 | 0.846 | -0.436 | 0.852 |
| PE_PGRS43 | -0.268 | 0.999 | -1.386 | 0.831 | -1.119 | 0.816 |
| Rv2491 | 0.045 | NA | -0.464 | 0.848 | -0.510 | 0.816 |
| Rv2492 | 0.037 | NA | -0.739 | 0.732 | -0.776 | 0.816 |
| vapB38 | 0.238 | NA | 0.242 | 0.865 | 0.005 | 0.997 |
| vapC38 | 0.004 | 0.999 | -0.095 | 0.934 | -0.099 | 0.916 |
| bkdC | -0.107 | 0.999 | -0.026 | 0.973 | 0.081 | 0.904 |
| bkdB | -0.197 | 0.999 | -0.201 | 0.820 | -0.004 | 0.994 |
| bkdA | -0.206 | 0.999 | -0.341 | 0.848 | -0.135 | 0.902 |
| citE | -0.185 | 0.999 | 0.098 | 0.937 | 0.283 | 0.819 |
| Rv2499c | -0.069 | 0.999 | -0.474 | 0.498 | -0.404 | 0.816 |
| fadE19 | -0.110 | 0.999 | -0.273 | 0.879 | -0.163 | 0.897 |
| accA1 | -0.107 | 0.999 | -0.158 | 0.859 | -0.051 | 0.937 |
| accD1 | -0.004 | 0.999 | -0.123 | 0.848 | -0.119 | 0.819 |
| scoB | 0.029 | 0.999 | -0.321 | 0.820 | -0.349 | 0.816 |
| scoA | -0.052 | 0.999 | 0.040 | 0.961 | 0.092 | 0.886 |
| fadD35 | 0.161 | 0.999 | -0.070 | 0.918 | -0.231 | 0.816 |
| Rv2506 | -0.025 | 0.999 | -0.025 | 0.973 | 0.000 | 0.999 |
| Rv2507 | 0.158 | 0.999 | -0.046 | 0.970 | -0.203 | 0.858 |
| Rv2508c | 0.092 | 0.999 | 0.076 | 0.912 | -0.016 | 0.977 |
| Rv2509 | 0.021 | 0.999 | 0.170 | 0.848 | 0.149 | 0.832 |
| Rv2510c | 0.232 | 0.999 | 0.016 | 0.974 | -0.216 | 0.816 |
| orn | -0.073 | 0.999 | 0.257 | 0.853 | 0.330 | 0.816 |
| Rv2512a | 0.061 | 0.999 | -0.443 | 0.843 | -0.504 | 0.816 |
| Rv2512c | -0.130 | NA | -0.365 | 0.852 | -0.235 | 0.867 |
| Rv2513 | -0.034 | 0.999 | -0.226 | 0.879 | -0.192 | 0.868 |
| Rv2514c | -0.019 | NA | -0.531 | 0.826 | -0.511 | 0.816 |
| Rv2515c | -0.056 | NA | 0.041 | 0.970 | 0.097 | 0.905 |
| Rv2516c | 0.031 | 0.999 | -0.115 | 0.913 | -0.146 | 0.866 |
| Rv2517c | -0.276 | 0.999 | 0.057 | 0.975 | 0.333 | 0.865 |
| ldtB | -0.248 | 0.999 | -0.462 | 0.155 | -0.214 | 0.816 |
| PE26 | -0.091 | 0.999 | -0.001 | 0.996 | 0.090 | 0.819 |
| Rv2520c | -0.214 | NA | -0.461 | 0.859 | -0.247 | 0.896 |
| bcp | -0.026 | 0.999 | 0.044 | 0.936 | 0.070 | 0.878 |
| Rv2522c | -0.143 | 0.999 | -0.078 | 0.920 | 0.065 | 0.911 |
| acpS | -0.256 | 0.999 | 0.128 | 0.922 | 0.384 | 0.816 |
| fas | -0.322 | 0.999 | -0.284 | 0.848 | 0.038 | 0.967 |
| Rv2525c | 0.098 | 0.999 | 0.221 | 0.837 | 0.123 | 0.865 |
| vapB17 | 0.037 | 0.999 | -0.507 | 0.750 | -0.544 | 0.816 |
| vapC17 | 0.056 | 0.999 | -0.279 | 0.848 | -0.334 | 0.816 |
| mrr | -0.142 | NA | -0.333 | 0.873 | -0.192 | 0.896 |
| Rv2529 | 0.110 | NA | 0.000 | 1.000 | -0.110 | 0.936 |
| vapB39 | 0.022 | 0.999 | 0.142 | 0.948 | 0.120 | 0.942 |
| vapC39 | 0.158 | 0.999 | -0.049 | 0.952 | -0.207 | 0.817 |
| Rv2531c | 0.039 | 0.999 | -0.098 | 0.878 | -0.137 | 0.816 |
| Rv2532c | 0.134 | 0.999 | -0.207 | 0.859 | -0.341 | 0.816 |
| nusB | -0.056 | 0.999 | 0.041 | 0.970 | 0.097 | 0.909 |
| efp | 0.117 | 0.999 | 0.248 | 0.826 | 0.131 | 0.838 |
| pepQ | 0.042 | 0.999 | 0.297 | 0.684 | 0.254 | 0.816 |
| Rv2536 | 0.043 | 0.999 | 0.141 | 0.832 | 0.098 | 0.835 |
| aroD | 0.075 | 0.999 | -0.158 | 0.883 | -0.233 | 0.822 |
| aroB | -0.118 | 0.999 | 0.110 | 0.921 | 0.227 | 0.822 |
| aroK | 0.022 | 0.999 | 0.198 | 0.879 | 0.176 | 0.867 |
| aroF | -0.090 | 0.999 | 0.084 | 0.941 | 0.174 | 0.862 |
| Rv2541 | 0.264 | NA | 1.000 | 0.826 | 0.735 | 0.817 |
| Rv2542 | 0.150 | 0.999 | 0.241 | 0.820 | 0.091 | 0.862 |
| lppA | 0.371 | NA | -0.359 | 0.879 | -0.730 | 0.816 |
| lppB | -0.024 | NA | -0.706 | NA | -0.682 | NA |
| vapB18 | 0.299 | NA | -0.384 | 0.866 | -0.683 | 0.816 |
| vapC18 | 0.424 | NA | 0.076 | 0.962 | -0.348 | 0.831 |
| vapB19 | -0.119 | 0.999 | -0.181 | 0.879 | -0.062 | 0.947 |
| vapC19 | 0.025 | 0.999 | -0.173 | 0.879 | -0.198 | 0.835 |
| Rv2548A | -0.054 | 0.999 | -0.043 | 0.948 | 0.011 | 0.985 |
| vapC20 | 0.110 | NA | -0.234 | 0.843 | -0.344 | 0.816 |
| vapB20 | -0.141 | NA | 0.095 | 0.940 | 0.236 | 0.836 |
| Rv2551c | 0.001 | 0.999 | 0.329 | 0.826 | 0.328 | 0.816 |
| aroE | -0.009 | 0.999 | 0.105 | 0.879 | 0.114 | 0.852 |
| Rv2553c | 0.095 | 0.999 | -0.069 | 0.922 | -0.164 | 0.817 |
| Rv2554c | 0.040 | 0.999 | 0.340 | 0.843 | 0.300 | 0.819 |
| alaS | -0.007 | 0.999 | -0.026 | 0.970 | -0.018 | 0.975 |
| Rv2556c | -0.035 | 0.999 | -0.430 | 0.826 | -0.395 | 0.816 |
| Rv2557 | 0.147 | 0.999 | 0.048 | 0.959 | -0.100 | 0.894 |
| Rv2558 | -0.056 | 0.999 | 0.168 | 0.879 | 0.224 | 0.822 |
| Rv2559c | 0.113 | 0.999 | 0.313 | 0.820 | 0.200 | 0.816 |
| Rv2560 | 0.023 | NA | -0.047 | 0.973 | -0.070 | 0.954 |
| Rv2563 | 0.100 | 0.999 | -0.023 | 0.975 | -0.123 | 0.882 |
| glnQ | 0.121 | 0.999 | 0.315 | 0.820 | 0.194 | 0.816 |
| Rv2565 | 0.057 | 0.999 | -0.159 | 0.854 | -0.216 | 0.816 |
| Rv2566 | 0.111 | 0.999 | 0.411 | 0.659 | 0.300 | 0.816 |
| Rv2567 | 0.147 | 0.999 | 0.060 | 0.881 | -0.087 | 0.824 |
| Rv2568c | 0.051 | 0.999 | -0.170 | 0.843 | -0.221 | 0.816 |
| Rv2569c | 0.157 | 0.999 | 0.021 | 0.970 | -0.136 | 0.819 |
| Rv2570 | -0.273 | NA | 0.005 | 0.994 | 0.277 | 0.816 |
| Rv2571c | 0.053 | 0.999 | -0.048 | 0.959 | -0.100 | 0.892 |
| aspS | 0.104 | 0.999 | 0.165 | 0.832 | 0.061 | 0.896 |
| Rv2573 | 0.139 | NA | 0.635 | 0.527 | 0.495 | 0.816 |
| Rv2574 | 0.069 | 0.999 | 0.148 | 0.879 | 0.079 | 0.922 |
| Rv2575 | 0.143 | 0.999 | 0.024 | 0.967 | -0.119 | 0.831 |
| Rv2576c | -0.108 | 0.999 | 0.365 | 0.826 | 0.473 | 0.816 |
| Rv2577 | 0.138 | NA | -0.154 | 0.879 | -0.292 | 0.816 |
| Rv2578c | 0.175 | 0.999 | 0.033 | 0.958 | -0.142 | 0.817 |
| dhaA | 0.044 | 0.999 | -0.057 | 0.885 | -0.100 | 0.816 |
| hisS | 0.044 | 0.999 | -0.015 | 0.975 | -0.059 | 0.906 |
| Rv2581c | 0.088 | 0.999 | -0.326 | 0.732 | -0.414 | 0.816 |
| ppiB | 0.220 | 0.999 | -0.377 | 0.879 | -0.597 | 0.816 |
| relA | 0.029 | 0.999 | 0.054 | 0.922 | 0.025 | 0.951 |
| apt | -0.205 | 0.999 | 0.278 | 0.848 | 0.483 | 0.816 |
| Rv2585c | 0.023 | 0.999 | -0.073 | 0.948 | -0.096 | 0.911 |
| secF | 0.070 | 0.999 | 0.222 | 0.848 | 0.151 | 0.862 |
| secD | 0.131 | 0.999 | -0.083 | 0.879 | -0.213 | 0.816 |
| yajC | 0.019 | 0.999 | -0.052 | 0.940 | -0.071 | 0.896 |
| gabT | -0.010 | 0.999 | 0.090 | 0.886 | 0.100 | 0.862 |
| fadD9 | 0.029 | 0.999 | 0.024 | 0.973 | -0.005 | 0.995 |
| PE_PGRS44 | 0.032 | 0.999 | 0.071 | 0.901 | 0.039 | 0.936 |
| ruvB | 0.070 | 0.999 | 0.262 | 0.831 | 0.191 | 0.819 |
| ruvA | 0.107 | 0.999 | -0.042 | 0.962 | -0.148 | 0.854 |
| ruvC | -0.030 | 0.999 | -0.215 | 0.879 | -0.184 | 0.868 |
| vapB40 | -0.159 | 0.999 | 0.039 | 0.970 | 0.198 | 0.835 |
| vapC40 | 0.107 | 0.999 | -0.207 | 0.852 | -0.313 | 0.816 |
| Rv2597 | -0.066 | 0.999 | -0.045 | 0.941 | 0.021 | 0.964 |
| Rv2598 | -0.059 | NA | 0.101 | 0.885 | 0.160 | 0.819 |
| Rv2599 | 0.192 | NA | -0.024 | 0.974 | -0.216 | 0.817 |
| Rv2600 | 0.049 | 0.999 | 0.138 | 0.918 | 0.089 | 0.932 |
| speE | 0.187 | 0.999 | 0.226 | 0.829 | 0.039 | 0.942 |
| vapB41 | -0.061 | 0.999 | 0.115 | 0.895 | 0.177 | 0.836 |
| vapC41 | 0.000 | 0.999 | 0.014 | 0.982 | 0.014 | 0.984 |
| Rv2603c | -0.064 | 0.999 | 0.069 | 0.895 | 0.133 | 0.817 |
| snoP | 0.132 | 0.999 | 0.000 | 0.999 | -0.131 | 0.819 |

| | WT v Δ*coaX* | | Δ*coaX* v complement | | WT v complement | |
|---|---|---|---|---|---|---|
| **GeneID** | **log2[FC]** | **padj** | **log2[FC]** | **padj** | **log2[FC]** | **padj** |
| Rv3239c | -0.079 | 0.999 | 0.253 | 0.832 | 0.332 | 0.816 |
| secA1 | 0.040 | 0.999 | -0.130 | 0.848 | -0.169 | 0.816 |
| Rv3241c | 0.059 | 0.999 | -0.115 | 0.848 | -0.173 | 0.816 |
| Rv3242c | -0.023 | 0.999 | -0.140 | 0.831 | -0.117 | 0.816 |
| Rv3243c | 0.082 | 0.999 | 0.067 | 0.946 | -0.014 | 0.986 |
| lpqB | 0.025 | 0.999 | -0.084 | 0.922 | -0.109 | 0.868 |
| mtrB | 0.117 | 0.999 | -0.151 | 0.879 | -0.268 | 0.816 |
| mtrA | 0.168 | 0.999 | 0.286 | 0.826 | 0.118 | 0.866 |
| tmk | 0.025 | 0.999 | 0.235 | 0.843 | 0.210 | 0.817 |
| sahH | 0.039 | 0.999 | -0.191 | 0.843 | -0.230 | 0.816 |
| Rv3249c | -0.060 | 0.999 | -0.468 | 0.820 | -0.408 | 0.816 |
| rubB | 0.018 | 0.999 | -0.282 | 0.832 | -0.300 | 0.816 |
| rubA | -0.064 | 0.999 | 0.302 | 0.915 | 0.366 | 0.868 |
| alkB | -0.006 | 0.999 | -0.161 | 0.899 | -0.155 | 0.884 |
| Rv3253c | 0.090 | 0.999 | 0.343 | 0.826 | 0.253 | 0.816 |
| Rv3254 | -0.079 | 0.999 | -0.026 | 0.960 | 0.053 | 0.896 |
| manA | 0.032 | 0.999 | 0.123 | 0.859 | 0.091 | 0.865 |
| Rv3256c | 0.062 | 0.999 | 0.476 | 0.826 | 0.415 | 0.816 |
| pmmA | 0.044 | 0.999 | 0.179 | 0.832 | 0.135 | 0.831 |
| Rv3258c | 0.085 | 0.999 | 0.632 | 0.795 | 0.547 | 0.816 |
| Rv3259 | -0.021 | 0.999 | 0.100 | 0.878 | 0.121 | 0.824 |
| whiB2 | 0.114 | 0.999 | 0.159 | 0.843 | 0.045 | 0.930 |
| fbiA | 0.139 | 0.999 | 0.152 | 0.824 | 0.013 | 0.967 |
| fbiB | 0.076 | 0.999 | 0.263 | 0.825 | 0.186 | 0.816 |
| Rv3263 | -0.113 | 0.999 | -0.234 | 0.439 | -0.121 | 0.816 |
| manB | 0.055 | 0.999 | 0.090 | 0.922 | 0.035 | 0.958 |
| wbbL1 | 0.124 | 0.999 | 0.170 | 0.861 | 0.046 | 0.947 |
| rmlD | -0.030 | 0.999 | 0.227 | 0.883 | 0.257 | 0.854 |
| Rv3267 | 0.098 | 0.999 | -0.099 | 0.879 | -0.198 | 0.816 |
| Rv3268 | -0.128 | 0.999 | 0.309 | 0.832 | 0.438 | 0.816 |
| Rv3269 | -0.290 | 0.999 | -0.350 | 0.826 | -0.060 | 0.942 |
| ctpC | -0.185 | 0.999 | -0.158 | 0.922 | 0.028 | 0.984 |
| Rv3271c | -0.066 | 0.999 | 0.400 | 0.826 | 0.467 | 0.816 |
| Rv3272 | -0.023 | 0.999 | -0.111 | 0.862 | -0.087 | 0.862 |
| Rv3273 | 0.063 | 0.999 | -0.093 | 0.890 | -0.156 | 0.819 |
| fadE25 | 0.078 | 0.999 | 0.147 | 0.843 | 0.068 | 0.883 |
| purE | 0.106 | 0.999 | 0.459 | 0.820 | 0.353 | 0.816 |
| purK | 0.098 | 0.999 | 0.325 | 0.825 | 0.227 | 0.816 |
| Rv3277 | 0.090 | 0.999 | 0.013 | 0.975 | -0.077 | 0.871 |
| Rv3278c | 0.154 | NA | -0.056 | 0.927 | -0.210 | 0.816 |
| birA | 0.037 | 0.999 | 0.304 | 0.660 | 0.267 | 0.816 |
| accD5 | 0.115 | 0.999 | -0.109 | 0.879 | -0.224 | 0.816 |
| accE5 | 0.002 | 0.999 | -0.405 | 0.660 | -0.407 | 0.816 |
| Rv3282 | 0.187 | 0.999 | 0.365 | 0.826 | 0.178 | 0.844 |
| sseA | -0.003 | 0.999 | -0.078 | 0.885 | -0.075 | 0.868 |
| Rv3284 | 0.108 | 0.999 | -0.135 | 0.879 | -0.242 | 0.816 |
| accA3 | -0.163 | 0.999 | 0.220 | 0.848 | 0.383 | 0.816 |
| sigF | 0.010 | 0.999 | -0.051 | 0.936 | -0.060 | 0.904 |
| rsbW | -0.123 | 0.999 | 0.143 | 0.856 | 0.265 | 0.816 |
| usfY | -0.064 | 0.999 | -0.643 | 0.213 | -0.579 | 0.816 |
| Rv3289c | -0.107 | NA | -0.172 | 0.859 | -0.065 | 0.925 |
| lat | -0.111 | 0.999 | -0.107 | 0.859 | 0.004 | 0.994 |
| lrpA | 0.140 | 0.999 | 0.166 | 0.862 | 0.026 | 0.969 |
| Rv3292 | 0.060 | 0.999 | 0.163 | 0.832 | 0.103 | 0.846 |
| pcd | 0.090 | 0.999 | 0.238 | 0.678 | 0.148 | 0.816 |
| Rv3294c | -0.067 | NA | -1.285 | NA | -1.219 | NA |
| Rv3295 | 0.045 | 0.999 | 0.354 | 0.834 | 0.309 | 0.819 |
| lhr | 0.014 | 0.999 | 0.153 | 0.879 | 0.139 | 0.860 |
| nei | 0.053 | 0.999 | 0.024 | 0.973 | -0.029 | 0.961 |
| lpqC | -0.138 | 0.999 | 0.405 | 0.820 | 0.543 | 0.816 |
| atsB | 0.070 | 0.999 | 0.047 | 0.879 | -0.023 | 0.930 |
| Rv3300c | 0.078 | 0.999 | 0.329 | 0.820 | 0.251 | 0.816 |
| phoY1 | 0.129 | 0.999 | -0.038 | 0.970 | -0.168 | 0.855 |
| glpD2 | 0.098 | 0.999 | 0.056 | 0.886 | -0.042 | 0.896 |
| lpdA | 0.097 | 0.999 | 0.228 | 0.820 | 0.131 | 0.819 |
| Rv3304 | -0.053 | 0.999 | 0.335 | 0.831 | 0.388 | 0.816 |
| amiA1 | -0.027 | 0.999 | 0.248 | 0.859 | 0.274 | 0.822 |
| amiB1 | 0.012 | 0.999 | 0.136 | 0.859 | 0.124 | 0.838 |
| deoD | 0.019 | 0.999 | 0.464 | 0.831 | 0.445 | 0.816 |
| pmmB | 0.030 | 0.999 | 0.299 | 0.831 | 0.269 | 0.816 |
| upp | -0.016 | NA | -0.078 | 0.933 | -0.062 | 0.936 |
| sapM | 0.173 | 0.999 | 0.106 | 0.843 | -0.067 | 0.855 |
| Rv3311 | 0.034 | 0.999 | -0.113 | 0.848 | -0.147 | 0.816 |
| Rv3312A | 0.110 | 0.999 | 0.285 | 0.848 | 0.175 | 0.862 |
| Rv3312b | 0.061 | 0.999 | -0.033 | 0.958 | -0.094 | 0.860 |
| Rv3312c | 0.004 | 0.999 | 0.093 | 0.879 | 0.089 | 0.855 |
| add | 0.133 | 0.999 | 0.347 | 0.831 | 0.215 | 0.836 |
| deoA | 0.066 | 0.999 | 0.107 | 0.886 | 0.041 | 0.946 |
| cdd | 0.175 | NA | -0.067 | 0.930 | -0.242 | 0.816 |
| sdhC | 0.076 | 0.999 | -0.490 | 0.820 | -0.567 | 0.816 |
| sdhD | 0.220 | 0.999 | -0.281 | 0.832 | -0.502 | 0.816 |
| sdhA | 0.070 | 0.999 | 0.021 | 0.973 | -0.049 | 0.932 |
| sdhB | 0.078 | 0.999 | -0.072 | 0.901 | -0.150 | 0.816 |
| vapC44 | 0.142 | 0.999 | -0.589 | 0.769 | -0.731 | 0.816 |
| vapB44 | 0.033 | NA | 0.030 | 0.973 | -0.002 | 0.998 |
| Rv3322c | 0.289 | 0.999 | 0.368 | 0.801 | 0.079 | 0.906 |
| moaX | 0.115 | 0.999 | -0.153 | 0.859 | -0.269 | 0.816 |
| moaC3 | -0.068 | NA | -0.297 | 0.848 | -0.229 | 0.842 |
| Rv3327 | -0.090 | 0.999 | -0.514 | 0.826 | -0.425 | 0.816 |
| sigJ | -0.001 | 0.999 | 0.347 | 0.832 | 0.348 | 0.816 |
| Rv3329 | 0.057 | 0.999 | 0.160 | 0.848 | 0.103 | 0.866 |
| dacB1 | 0.044 | 0.999 | 0.398 | 0.826 | 0.354 | 0.816 |
| sugI | 0.074 | 0.999 | 0.594 | 0.820 | 0.520 | 0.816 |
| nagA | 0.162 | 0.999 | 0.405 | 0.709 | 0.243 | 0.816 |
| Rv3333c | -0.295 | NA | -0.261 | 0.886 | 0.034 | 0.984 |
| Rv3334 | -0.180 | 0.999 | -0.666 | 0.826 | -0.486 | 0.817 |
| Rv3335c | -0.027 | 0.999 | -0.155 | 0.832 | -0.129 | 0.819 |
| trpS | 0.016 | 0.999 | 0.042 | 0.927 | 0.026 | 0.942 |
| icd1 | 0.036 | 0.999 | -0.039 | 0.956 | -0.075 | 0.892 |
| metC | -0.086 | 0.999 | -0.115 | 0.848 | -0.029 | 0.939 |
| metA | 0.011 | 0.999 | 0.127 | 0.879 | 0.116 | 0.868 |
| Rv3342 | -0.088 | 0.999 | -0.085 | 0.907 | 0.003 | 0.997 |
| PPE54 | -0.010 | 0.999 | -0.678 | 0.848 | -0.669 | 0.819 |
| PE_PGRS49 | 0.106 | NA | -0.951 | 0.848 | -1.057 | 0.816 |
| PE_PGRS50 | -0.190 | NA | -1.250 | 0.826 | -1.060 | 0.816 |
| Rv3346c | -0.209 | NA | -0.231 | 0.895 | -0.022 | 0.988 |
| PPE55 | 0.009 | 0.999 | -0.270 | 0.862 | -0.279 | 0.835 |
| Rv3348 | -0.019 | 0.999 | -0.363 | 0.820 | -0.344 | 0.816 |
| Rv3349c | -0.080 | NA | -0.091 | 0.962 | -0.010 | 0.996 |
| PPE56 | 0.044 | 0.999 | -0.472 | 0.832 | -0.516 | 0.816 |
| Rv3351c | 0.129 | NA | -0.383 | 0.854 | -0.513 | 0.816 |
| Rv3352c | 0.051 | NA | 0.108 | 0.952 | 0.057 | 0.969 |
| Rv3353c | -0.028 | NA | 0.364 | 0.848 | 0.391 | 0.817 |
| Rv3354 | 0.126 | 0.999 | -0.214 | 0.848 | -0.341 | 0.816 |
| Rv3355c | 0.160 | NA | 0.242 | 0.895 | 0.082 | 0.956 |
| folD | 0.052 | 0.999 | 0.109 | 0.892 | 0.057 | 0.932 |
| relJ | 0.030 | NA | 0.178 | 0.902 | 0.147 | 0.899 |
| relK | 0.100 | NA | -0.300 | 0.859 | -0.400 | 0.816 |
| Rv3359 | -0.167 | NA | -0.023 | 0.975 | 0.144 | 0.866 |
| Rv3360 | -0.005 | NA | -0.375 | 0.826 | -0.370 | 0.816 |
| Rv3361c | 0.030 | 0.999 | -0.010 | 0.984 | -0.039 | 0.938 |
| Rv3362c | 0.080 | 0.999 | -0.025 | 0.973 | -0.105 | 0.870 |
| Rv3363c | 0.146 | 0.999 | 0.325 | 0.832 | 0.179 | 0.861 |
| Rv3364c | 0.162 | 0.999 | 0.053 | 0.959 | -0.108 | 0.896 |
| Rv3365c | 0.100 | 0.999 | 0.108 | 0.848 | 0.008 | 0.984 |
| spoU | -0.001 | 0.999 | 0.241 | 0.843 | 0.242 | 0.816 |
| PE_PGRS51 | -0.025 | 0.999 | 0.153 | 0.886 | 0.178 | 0.855 |
| Rv3368c | 0.035 | 0.999 | 0.384 | 0.832 | 0.349 | 0.816 |
| Rv3369 | -0.203 | 0.999 | 0.517 | 0.832 | 0.720 | 0.816 |
| dnaE2 | -0.255 | 0.999 | -0.235 | 0.859 | 0.020 | 0.985 |
| Rv3371 | -0.123 | 0.999 | -0.179 | 0.831 | -0.057 | 0.904 |
| otsB2 | -0.008 | 0.999 | 0.432 | 0.831 | 0.440 | 0.816 |
| echA18 | 0.043 | 0.999 | 0.144 | 0.859 | 0.101 | 0.868 |
| amiD | 0.080 | 0.999 | 0.001 | 0.995 | -0.078 | 0.862 |
| Rv3376 | 0.051 | 0.999 | -0.149 | 0.848 | -0.200 | 0.816 |
| Rv3377c | 0.136 | 0.999 | -0.443 | 0.826 | -0.580 | 0.816 |
| Rv3378c | 0.171 | 0.999 | -0.398 | 0.831 | -0.569 | 0.816 |
| dxs2 | -0.193 | 0.999 | -0.433 | 0.859 | -0.240 | 0.891 |
| Rv3382c | 0.170 | 0.999 | -0.056 | 0.922 | -0.226 | 0.816 |
| idsB | 0.045 | 0.999 | -0.086 | 0.951 | -0.132 | 0.909 |
| vapC46 | -0.056 | 0.999 | 0.224 | 0.834 | 0.279 | 0.816 |
| vapB46 | -0.087 | 0.999 | 0.440 | 0.832 | 0.527 | 0.816 |
| Rv3386 | 0.027 | NA | -0.076 | 0.922 | -0.104 | 0.868 |
| Rv3387 | 0.094 | NA | -0.631 | 0.826 | -0.725 | 0.816 |
| PE_PGRS52 | 0.025 | NA | -0.563 | 0.848 | -0.587 | 0.817 |
| htdY | 0.039 | 0.999 | 0.041 | 0.973 | 0.003 | 0.998 |
| lpqD | 0.181 | 0.999 | -0.258 | 0.879 | -0.439 | 0.816 |
| acrA1 | 0.185 | 0.999 | 0.445 | 0.118 | 0.260 | 0.816 |
| cmaA1 | 0.086 | 0.999 | -0.117 | 0.886 | -0.204 | 0.817 |
| iunH | -0.098 | 0.999 | 0.190 | 0.859 | 0.287 | 0.816 |
| Rv3394c | -0.131 | 0.999 | 0.383 | 0.834 | 0.514 | 0.816 |
| Rv3395A | -0.223 | NA | -0.336 | 0.831 | -0.113 | 0.896 |
| Rv3395c | -0.079 | 0.999 | 0.061 | 0.971 | 0.140 | 0.926 |
| guaA | -0.064 | 0.999 | 0.140 | 0.859 | 0.204 | 0.816 |
| phyA | -0.290 | NA | -0.115 | 0.922 | 0.175 | 0.862 |
| idsA1 | -0.008 | NA | -0.099 | 0.922 | -0.091 | 0.904 |
| Rv3399 | -0.138 | 0.999 | -0.240 | 0.860 | -0.102 | 0.921 |
| Rv3400 | -0.084 | 0.999 | 0.068 | 0.909 | 0.152 | 0.816 |
| Rv3401 | 0.012 | 0.999 | 0.010 | 0.981 | -0.002 | 0.996 |
| Rv3402c | 0.188 | 0.999 | -0.228 | 0.831 | -0.416 | 0.816 |
| Rv3403c | 0.139 | 0.999 | -0.189 | 0.820 | -0.328 | 0.816 |
| Rv3404c | 0.022 | 0.999 | 0.143 | 0.848 | 0.121 | 0.837 |
| Rv3405c | 0.083 | 0.999 | 0.350 | 0.498 | 0.267 | 0.816 |
| Rv3406 | -0.218 | 0.999 | -0.169 | 0.862 | 0.048 | 0.944 |
| vapB47 | 0.028 | 0.999 | -0.060 | 0.927 | -0.088 | 0.868 |
| vapC47 | -0.042 | 0.999 | -0.357 | 0.820 | -0.315 | 0.816 |
| choD | -0.024 | 0.999 | -0.081 | 0.879 | -0.057 | 0.886 |
| guaB3 | -0.148 | 0.999 | 0.018 | 0.985 | 0.166 | 0.873 |
| guaB2 | -0.155 | 0.999 | -0.004 | 0.992 | 0.151 | 0.816 |
| Rv3412 | -0.083 | 0.999 | 0.054 | 0.929 | 0.136 | 0.819 |
| Rv3413c | 0.006 | 0.999 | 0.355 | 0.831 | 0.349 | 0.816 |
| sigD | -0.034 | 0.999 | -0.121 | 0.885 | -0.087 | 0.896 |
| Rv3415c | -0.014 | NA | 0.548 | 0.732 | 0.562 | 0.816 |

| | | | | | | |
|---|---|---|---|---|---|---|
| mce1A | 0.200 | 0.999 | 0.191 | 0.831 | -0.009 | 0.986 |
| mce1B | 0.175 | 0.999 | -0.145 | 0.879 | -0.321 | 0.816 |
| mce1C | 0.236 | 0.836 | 0.205 | 0.659 | -0.031 | 0.924 |
| mce1D | 0.143 | 0.999 | 0.042 | 0.934 | -0.100 | 0.835 |
| lprK | 0.228 | 0.999 | -0.090 | 0.924 | -0.318 | 0.816 |
| mce1F | 0.117 | 0.999 | 0.125 | 0.855 | 0.008 | 0.987 |
| Rv0175 | 0.140 | 0.999 | 0.126 | 0.891 | -0.014 | 0.986 |
| Rv0176 | 0.087 | 0.999 | 0.002 | 0.995 | -0.085 | 0.868 |
| Rv0177 | 0.150 | 0.999 | -0.198 | 0.860 | -0.347 | 0.816 |
| Rv0178 | 0.100 | 0.999 | 0.018 | 0.985 | -0.082 | 0.936 |
| lprO | 0.086 | 0.999 | 0.112 | 0.826 | 0.025 | 0.925 |
| Rv0180c | -0.087 | 0.999 | -0.053 | 0.927 | 0.035 | 0.942 |
| Rv0181c | 0.052 | 0.999 | -0.321 | 0.767 | -0.373 | 0.816 |
| sigG | -0.040 | 0.999 | 0.014 | 0.970 | 0.053 | 0.868 |
| Rv0183 | 0.061 | 0.999 | 0.112 | 0.879 | 0.050 | 0.932 |
| Rv0184 | -0.041 | 0.999 | -0.103 | 0.919 | -0.062 | 0.937 |
| Rv0185 | 0.057 | 0.999 | -0.259 | 0.826 | -0.317 | 0.816 |
| bglS | -0.004 | 0.999 | 0.264 | 0.826 | 0.268 | 0.816 |
| mymT | -0.709 | 0.999 | -0.140 | 0.948 | 0.569 | 0.816 |
| Rv0187 | -0.101 | 0.999 | -0.150 | 0.843 | -0.049 | 0.918 |
| Rv0188 | -0.102 | 0.999 | 0.061 | 0.922 | 0.163 | 0.816 |
| ilvD | -0.006 | 0.999 | 0.097 | 0.844 | 0.103 | 0.816 |
| Rv0190 | -0.252 | 0.999 | -0.433 | 0.570 | -0.180 | 0.819 |
| Rv0191 | -0.218 | 0.999 | -0.102 | 0.922 | 0.116 | 0.891 |
| Rv0192 | -0.109 | 0.999 | 0.013 | 0.975 | 0.123 | 0.822 |
| Rv0193c | 0.301 | 0.999 | -0.404 | 0.826 | -0.705 | 0.816 |
| Rv0194 | 0.025 | NA | -0.074 | 0.962 | -0.099 | 0.942 |
| Rv0195 | -0.187 | NA | -0.727 | 0.831 | -0.540 | 0.819 |
| Rv0196 | -0.079 | NA | 0.333 | 0.859 | 0.412 | 0.817 |
| Rv0197 | -0.077 | 0.999 | 0.009 | 0.989 | 0.087 | 0.905 |
| zmp1 | 0.006 | 0.999 | -0.104 | 0.883 | -0.110 | 0.862 |
| Rv0199 | 0.060 | 0.999 | -0.266 | 0.848 | -0.326 | 0.816 |
| Rv0200 | -0.027 | 0.999 | 0.321 | 0.699 | 0.347 | 0.816 |
| Rv0201c | -0.033 | 0.999 | -0.270 | 0.820 | -0.237 | 0.816 |
| mmpL11 | -0.002 | 0.999 | 0.187 | 0.832 | 0.189 | 0.816 |
| Rv0203 | 0.191 | 0.999 | 0.299 | 0.826 | 0.108 | 0.879 |
| Rv0204c | -0.035 | 0.999 | 0.371 | 0.826 | 0.406 | 0.816 |
| Rv0205 | 0.074 | 0.999 | 0.228 | 0.831 | 0.154 | 0.832 |
| mmpL3 | 0.053 | 0.999 | -0.059 | 0.922 | -0.112 | 0.837 |
| Rv0207c | 0.035 | 0.999 | 0.137 | 0.848 | 0.103 | 0.837 |
| Rv0208c | 0.019 | 0.999 | -0.028 | 0.962 | -0.047 | 0.926 |
| Rv0209 | 0.101 | 0.999 | 0.350 | 0.831 | 0.250 | 0.819 |
| Rv0210 | 0.146 | 0.999 | 0.106 | 0.879 | -0.039 | 0.936 |
| pckA | -0.226 | 0.999 | -0.190 | 0.820 | 0.036 | 0.924 |
| nadR | -0.226 | 0.999 | -0.044 | 0.946 | 0.182 | 0.816 |
| Rv0213c | 0.049 | NA | -0.259 | 0.848 | -0.307 | 0.816 |
| fadD4 | -0.082 | 0.999 | -0.144 | 0.848 | -0.062 | 0.904 |
| Rv0216 | -0.085 | 0.999 | 0.120 | 0.879 | 0.206 | 0.816 |
| lipW | -0.101 | 0.999 | 0.101 | 0.926 | 0.202 | 0.837 |
| Rv0218 | -0.019 | NA | 0.115 | 0.901 | 0.134 | 0.867 |
| Rv0219 | 0.069 | NA | 0.286 | 0.848 | 0.216 | 0.846 |
| lipC | 0.024 | 0.999 | 0.267 | 0.826 | 0.244 | 0.816 |
| Rv0221 | 0.102 | 0.999 | -0.075 | 0.895 | -0.178 | 0.816 |
| echA1 | -0.076 | 0.999 | 0.229 | 0.848 | 0.305 | 0.816 |
| Rv0223c | 0.116 | 0.999 | 0.092 | 0.933 | -0.024 | 0.979 |
| Rv0224c | -0.041 | 0.999 | 0.621 | 0.826 | 0.662 | 0.816 |
| Rv0225 | -0.051 | 0.999 | 0.471 | 0.491 | 0.522 | 0.816 |
| Rv0226c | 0.103 | 0.999 | 0.445 | 0.701 | 0.341 | 0.816 |
| Rv0227c | 0.112 | 0.999 | -0.159 | 0.885 | -0.271 | 0.817 |
| Rv0228 | -0.082 | 0.999 | 0.417 | 0.831 | 0.499 | 0.816 |
| Rv0229c | 0.092 | 0.999 | 0.112 | 0.832 | 0.020 | 0.950 |
| php | 0.182 | 0.999 | -0.090 | 0.936 | -0.272 | 0.817 |
| fadE4 | 0.065 | 0.999 | -0.048 | 0.927 | -0.113 | 0.824 |
| Rv0232 | 0.261 | 0.999 | 0.246 | 0.848 | -0.015 | 0.986 |
| nrdB | 0.198 | 0.999 | -0.079 | 0.934 | -0.277 | 0.816 |
| gabD1 | -0.095 | 0.999 | 0.288 | 0.843 | 0.383 | 0.816 |
| Rv0235c | -0.006 | 0.999 | 0.019 | 0.977 | 0.024 | 0.973 |
| Rv0236A | 0.051 | 0.999 | 0.072 | 0.915 | 0.021 | 0.967 |
| aftD | -0.007 | 0.999 | 0.467 | 0.831 | 0.475 | 0.816 |
| lpqI | 0.022 | 0.999 | 0.428 | 0.829 | 0.406 | 0.816 |
| Rv0238 | 0.110 | 0.999 | -0.049 | 0.962 | -0.159 | 0.860 |
| vapB24 | 0.010 | 0.999 | -0.220 | 0.859 | -0.230 | 0.831 |
| vapC24 | -0.124 | 0.999 | -0.234 | 0.826 | -0.110 | 0.854 |
| htdX | 0.246 | 0.999 | 0.098 | 0.879 | -0.149 | 0.817 |
| fabG4 | 0.204 | 0.999 | 0.283 | 0.527 | 0.079 | 0.856 |
| fadA2 | 0.140 | 0.999 | 0.267 | 0.826 | 0.127 | 0.852 |
| fadE5 | 0.270 | 0.999 | 0.494 | 0.055 | 0.224 | 0.816 |
| Rv0245 | 0.001 | 0.999 | -0.012 | 0.975 | -0.012 | 0.975 |
| Rv0246 | 0.058 | 0.999 | -0.275 | 0.848 | -0.333 | 0.816 |
| Rv0247c | 0.257 | 0.999 | -0.247 | 0.879 | -0.504 | 0.816 |
| Rv0248c | 0.183 | 0.999 | -0.045 | 0.947 | -0.228 | 0.816 |
| Rv0249c | 0.240 | 0.999 | -0.054 | 0.946 | -0.294 | 0.816 |
| Rv0250c | 0.353 | 0.999 | 0.135 | 0.910 | -0.219 | 0.837 |
| hsp | -0.124 | 0.999 | 0.173 | 0.859 | 0.297 | 0.816 |
| nirB | -0.053 | 0.999 | -0.156 | 0.831 | -0.103 | 0.832 |
| nirD | 0.061 | NA | -0.173 | 0.859 | -0.234 | 0.816 |
| cobU | 0.073 | NA | 0.405 | 0.772 | 0.332 | 0.816 |
| cobQ1 | 0.066 | 0.999 | 0.179 | 0.826 | 0.113 | 0.831 |
| PPE2 | 0.125 | 0.999 | 0.031 | 0.974 | -0.094 | 0.924 |
| Rv0257 | 0.028 | 0.999 | 0.215 | 0.855 | 0.187 | 0.837 |
| Rv0258c | 0.056 | NA | 0.148 | 0.895 | 0.092 | 0.924 |
| Rv0259c | 0.059 | NA | -0.256 | 0.879 | -0.315 | 0.831 |
| Rv0260c | -0.058 | NA | -0.546 | 0.854 | -0.488 | 0.836 |
| narK3 | -0.067 | NA | 0.017 | 0.975 | 0.083 | 0.882 |
| aac | 0.041 | 0.999 | 0.343 | 0.820 | 0.302 | 0.816 |
| Rv0263c | -0.050 | 0.999 | 0.129 | 0.913 | 0.179 | 0.859 |
| Rv0264c | 0.192 | 0.999 | 0.136 | 0.879 | -0.056 | 0.932 |
| Rv0265c | 0.012 | 0.999 | 0.016 | 0.970 | 0.004 | 0.989 |
| oplA | 0.012 | 0.999 | 0.051 | 0.885 | 0.039 | 0.892 |
| narU | -0.091 | NA | -0.617 | 0.848 | -0.526 | 0.832 |
| Rv0268c | 0.110 | 0.999 | -0.022 | 0.974 | -0.132 | 0.856 |
| Rv0269c | 0.010 | 0.999 | 0.173 | 0.879 | 0.163 | 0.862 |
| fadD2 | 0.089 | 0.999 | 0.119 | 0.859 | 0.030 | 0.947 |
| fadE6 | -0.005 | 0.999 | -0.085 | 0.895 | -0.080 | 0.883 |
| Rv0272c | 0.032 | 0.999 | -0.120 | 0.912 | -0.152 | 0.866 |
| Rv0273c | -0.032 | 0.999 | -0.188 | 0.826 | -0.156 | 0.816 |
| Rv0274 | -0.067 | 0.999 | -0.112 | 0.886 | -0.045 | 0.942 |
| Rv0275c | -0.195 | 0.999 | 0.178 | 0.879 | 0.373 | 0.816 |
| Rv0276 | -0.082 | 0.999 | 0.246 | 0.832 | 0.328 | 0.816 |
| vapC25 | 0.101 | 0.999 | -0.169 | 0.857 | -0.270 | 0.816 |
| Rv0278c | -0.168 | NA | -0.151 | 0.914 | 0.017 | 0.988 |
| PE_PGRS4 | -0.037 | 0.999 | 0.013 | 0.991 | 0.050 | 0.967 |
| PPE3 | -0.037 | 0.999 | 0.098 | 0.951 | 0.135 | 0.920 |
| Rv0281 | -0.043 | 0.999 | -0.015 | 0.962 | 0.028 | 0.912 |
| eccA3 | 0.076 | 0.999 | -0.046 | 0.947 | -0.122 | 0.838 |
| eccB3 | 0.099 | 0.999 | 0.149 | 0.905 | 0.051 | 0.958 |
| eccC3 | 0.105 | 0.999 | 0.016 | 0.974 | -0.089 | 0.862 |
| PE5 | 0.142 | 0.999 | 0.426 | 0.825 | 0.284 | 0.816 |
| PPE4 | 0.107 | 0.999 | 0.048 | 0.934 | -0.059 | 0.897 |
| esxG | 0.196 | 0.999 | 0.532 | 0.826 | 0.336 | 0.817 |
| esxH | 0.129 | 0.999 | -0.208 | 0.854 | -0.337 | 0.816 |
| espG3 | 0.163 | 0.999 | 0.240 | 0.879 | 0.077 | 0.947 |
| eccD3 | 0.108 | 0.999 | 0.506 | 0.826 | 0.398 | 0.816 |
| mycP3 | 0.048 | 0.999 | 0.537 | 0.826 | 0.489 | 0.816 |
| eccE3 | 0.153 | 0.999 | 0.392 | 0.825 | 0.239 | 0.817 |
| Rv0293c | -0.028 | 0.999 | 0.165 | 0.878 | 0.192 | 0.831 |
| tam | 0.090 | NA | 0.241 | 0.831 | 0.152 | 0.836 |
| Rv0295c | 0.023 | 0.999 | 0.167 | 0.885 | 0.144 | 0.881 |
| Rv0296c | 0.088 | 0.999 | -0.047 | 0.944 | -0.135 | 0.831 |
| PE_PGRS5 | -0.150 | 0.999 | 0.178 | 0.879 | 0.328 | 0.816 |
| Rv0298 | -0.042 | 0.999 | -0.246 | 0.826 | -0.204 | 0.816 |
| Rv0299 | -0.067 | 0.999 | 0.104 | 0.918 | 0.171 | 0.845 |
| vapB2 | 0.054 | 0.999 | -0.386 | 0.831 | -0.440 | 0.816 |
| vapC2 | -0.111 | 0.999 | -0.428 | 0.660 | -0.317 | 0.816 |
| Rv0302 | -0.044 | 0.999 | -0.058 | 0.895 | -0.014 | 0.967 |
| Rv0303 | -0.114 | 0.999 | -0.108 | 0.875 | 0.007 | 0.989 |
| PPE5 | 0.093 | 0.999 | -0.753 | 0.848 | -0.845 | 0.816 |
| PPE6 | -0.012 | 0.999 | -0.394 | 0.866 | -0.383 | 0.838 |
| Rv0306 | -0.285 | NA | -0.699 | NA | -0.414 | NA |
| Rv0307c | 0.001 | 0.999 | 0.382 | 0.820 | 0.381 | 0.816 |
| Rv0308 | 0.082 | 0.999 | 0.178 | 0.831 | 0.096 | 0.855 |
| Rv0309 | 0.014 | 0.999 | 0.294 | 0.826 | 0.281 | 0.816 |
| Rv0310c | 0.044 | NA | 0.299 | 0.832 | 0.255 | 0.817 |
| Rv0311 | -0.224 | 0.999 | -0.264 | 0.826 | -0.040 | 0.950 |
| Rv0312 | 0.145 | 0.999 | 0.071 | 0.879 | -0.074 | 0.842 |
| Rv0313 | -0.228 | 0.999 | 0.047 | 0.948 | 0.275 | 0.816 |
| Rv0314c | -0.026 | 0.999 | 0.331 | 0.826 | 0.357 | 0.816 |
| Rv0315 | -0.042 | 0.999 | 0.391 | 0.826 | 0.433 | 0.816 |
| Rv0316 | 0.154 | 0.999 | 0.319 | 0.635 | 0.166 | 0.816 |
| glpQ2 | 0.011 | 0.999 | 0.142 | 0.848 | 0.131 | 0.831 |
| Rv0318c | -0.039 | NA | 0.236 | 0.855 | 0.274 | 0.817 |
| pcp | -0.142 | 0.999 | -0.017 | 0.985 | 0.124 | 0.896 |
| Rv0320 | 0.260 | 0.999 | 0.171 | 0.831 | -0.089 | 0.862 |
| dcd | 0.227 | 0.999 | 0.071 | 0.914 | -0.155 | 0.817 |
| udgA | -0.054 | 0.999 | -0.142 | 0.913 | -0.087 | 0.932 |
| Rv0323c | 0.039 | NA | -0.184 | 0.848 | -0.223 | 0.816 |
| Rv0324 | -0.105 | NA | -0.147 | 0.883 | -0.042 | 0.957 |
| Rv0325 | -0.194 | NA | -0.005 | 0.995 | 0.189 | 0.865 |
| Rv0326 | -0.174 | NA | -0.530 | 0.659 | -0.356 | 0.816 |
| cyp135A1 | 0.005 | NA | -0.781 | 0.831 | -0.786 | 0.816 |
| Rv0328 | -0.022 | 0.999 | 0.044 | 0.959 | 0.067 | 0.924 |
| Rv0329c | -0.005 | NA | 0.018 | 0.988 | 0.022 | 0.986 |
| Rv0330c | 0.035 | NA | 0.127 | 0.879 | 0.092 | 0.888 |
| Rv0331 | -0.166 | NA | -0.104 | 0.886 | 0.062 | 0.920 |
| Rv0332 | 0.005 | 0.999 | 0.357 | 0.831 | 0.353 | 0.816 |
| Rv0333 | 0.126 | 0.999 | 0.081 | 0.895 | -0.045 | 0.932 |
| rmlA | 0.116 | 0.999 | -0.211 | 0.866 | -0.328 | 0.816 |
| PE6 | -0.241 | NA | -0.418 | 0.865 | -0.177 | 0.924 |
| Rv0336 | -0.027 | 0.999 | -0.113 | 0.879 | -0.086 | 0.868 |
| aspC | 0.055 | 0.999 | 0.006 | 0.990 | -0.049 | 0.936 |
| Rv0338c | -0.008 | 0.999 | -0.111 | 0.848 | -0.102 | 0.819 |
| Rv0339c | -0.007 | 0.999 | 0.110 | 0.877 | 0.117 | 0.836 |
| Rv0340 | -0.139 | 0.999 | -0.221 | 0.826 | -0.082 | 0.882 |
| iniB | -0.449 | 0.999 | 0.109 | 0.895 | 0.558 | 0.816 |
| iniA | -0.306 | 0.999 | -0.287 | 0.820 | 0.018 | 0.975 |
| rpmF | -0.143 | 0.999 | 0.524 | 0.831 | 0.667 | 0.816 |
| Rv0979c | 0.125 | NA | -0.033 | 0.989 | -0.158 | 0.947 |
| PE_PGRS18 | -0.227 | NA | -0.545 | 0.843 | -0.319 | 0.862 |
| mprA | 0.059 | 0.999 | -0.100 | 0.879 | -0.159 | 0.817 |
| mprB | -0.038 | 0.999 | 0.007 | 0.983 | 0.045 | 0.900 |
| pepD | -0.188 | 0.999 | 0.214 | 0.859 | 0.403 | 0.816 |
| moaB2 | -0.072 | 0.999 | 0.443 | 0.826 | 0.515 | 0.816 |
| mscL | -0.038 | NA | -0.758 | NA | -0.720 | NA |
| Rv0986 | 0.007 | 0.999 | -0.351 | 0.832 | -0.358 | 0.816 |
| Rv0987 | -0.071 | 0.999 | -0.503 | 0.829 | -0.432 | 0.816 |
| Rv0988 | -0.009 | 0.999 | -0.232 | 0.897 | -0.222 | 0.884 |
| grcC2 | 0.081 | 0.999 | -0.145 | 0.948 | -0.226 | 0.897 |
| Rv0990c | -0.285 | NA | -0.202 | 0.906 | 0.083 | 0.950 |
| Rv0991c | -0.019 | 0.999 | -0.132 | 0.879 | -0.113 | 0.865 |
| Rv0992c | -0.111 | 0.999 | 0.449 | 0.826 | 0.559 | 0.816 |
| galU | 0.273 | 0.999 | 0.257 | 0.831 | -0.017 | 0.981 |
| moeA1 | 0.134 | 0.999 | 0.098 | 0.879 | -0.036 | 0.936 |
| rimJ | 0.225 | 0.999 | 0.044 | 0.962 | -0.181 | 0.836 |
| Rv0996 | 0.198 | 0.999 | 0.024 | 0.981 | -0.173 | 0.868 |
| Rv0997 | 0.239 | 0.999 | 0.041 | 0.948 | -0.198 | 0.816 |
| Rv0997a | 0.017 | 0.999 | 0.176 | 0.832 | 0.159 | 0.816 |
| Rv0998 | 0.064 | 0.999 | 0.268 | 0.826 | 0.203 | 0.816 |
| Rv0999 | -0.108 | 0.999 | 0.457 | 0.831 | 0.565 | 0.816 |
| Rv1000c | -0.074 | 0.999 | 0.316 | 0.859 | 0.391 | 0.817 |
| arcA | -0.007 | 0.999 | 0.063 | 0.922 | 0.070 | 0.892 |
| Rv1002c | 0.039 | 0.999 | 0.457 | 0.826 | 0.418 | 0.816 |
| Rv1003 | 0.144 | 0.999 | 0.358 | 0.826 | 0.214 | 0.835 |
| Rv1004c | -0.183 | 0.999 | 0.802 | 0.826 | 0.985 | 0.816 |
| Rv1005c | 0.029 | 0.999 | 0.280 | 0.832 | 0.251 | 0.816 |
| Rv1006 | 0.079 | 0.999 | 0.070 | 0.886 | -0.009 | 0.984 |
| metS | -0.014 | 0.999 | 0.152 | 0.858 | 0.166 | 0.819 |
| tatD | -0.035 | NA | 0.286 | 0.848 | 0.321 | 0.816 |
| rpfB | 0.200 | 0.999 | 0.307 | 0.857 | 0.107 | 0.930 |
| ksgA | 0.172 | 0.999 | 0.402 | 0.826 | 0.229 | 0.834 |
| ispE | 0.105 | 0.999 | 0.321 | 0.826 | 0.216 | 0.819 |
| Rv1012 | 0.033 | NA | -0.100 | 0.929 | -0.132 | 0.884 |
| pks16 | 0.010 | 0.999 | 0.146 | 0.879 | 0.135 | 0.862 |
| pth | -0.076 | 0.999 | 0.075 | 0.948 | 0.152 | 0.876 |
| rplY | -0.015 | 0.999 | -0.155 | 0.843 | -0.140 | 0.819 |
| lpqT | 0.057 | 0.999 | -0.140 | 0.879 | -0.197 | 0.817 |
| prsA | 0.051 | 0.999 | 0.053 | 0.927 | 0.002 | 0.997 |
| glmU | 0.097 | 0.999 | 0.108 | 0.848 | 0.011 | 0.977 |
| Rv1019 | -0.246 | 0.999 | -0.067 | 0.962 | 0.180 | 0.882 |
| mfd | 0.002 | 0.999 | 0.104 | 0.879 | 0.102 | 0.862 |
| Rv1021 | 0.081 | 0.999 | -0.050 | 0.945 | -0.131 | 0.837 |
| lpqU | 0.040 | 0.999 | 0.185 | 0.831 | 0.145 | 0.817 |
| eno | 0.040 | 0.999 | 0.090 | 0.886 | 0.049 | 0.925 |
| Rv1024 | 0.138 | 0.999 | -0.301 | 0.795 | -0.439 | 0.816 |
| Rv1025 | 0.023 | 0.999 | 0.108 | 0.859 | 0.085 | 0.856 |
| Rv1026 | 0.083 | 0.999 | 0.190 | 0.846 | 0.107 | 0.867 |
| kdpE | 0.015 | NA | 0.195 | 0.848 | 0.180 | 0.821 |
| kdpF | -0.102 | NA | -0.370 | 0.879 | -0.268 | 0.892 |
| kdpD | -0.102 | 0.999 | -0.222 | 0.826 | -0.121 | 0.842 |
| kdpA | 0.048 | NA | -0.283 | 0.826 | -0.331 | 0.816 |
| kdpB | -0.068 | 0.999 | -0.114 | 0.859 | -0.046 | 0.925 |
| kdpC | -0.072 | NA | 0.029 | 0.970 | 0.101 | 0.879 |
| trcS | 0.055 | 0.999 | -0.140 | 0.869 | -0.195 | 0.816 |
| trcR | -0.043 | 0.999 | -0.142 | 0.848 | -0.099 | 0.857 |
| esxI | -0.074 | NA | -0.133 | 0.885 | -0.059 | 0.939 |
| esxJ | -0.038 | 0.999 | -0.679 | 0.826 | -0.641 | 0.816 |
| PPE15 | -0.180 | NA | -0.654 | 0.848 | -0.475 | 0.852 |
| PE8 | -0.046 | NA | -0.464 | 0.859 | -0.418 | 0.846 |
| Rv1041c | -0.019 | 0.999 | -0.141 | 0.854 | -0.122 | 0.837 |
| Rv1042c | 0.055 | 0.999 | -0.210 | 0.914 | -0.265 | 0.867 |
| Rv1043c | 0.014 | 0.999 | 0.012 | 0.987 | -0.003 | 0.997 |
| Rv1044 | 0.022 | NA | -0.034 | 0.973 | -0.056 | 0.952 |
| Rv1045 | -0.036 | 0.999 | -0.156 | 0.848 | -0.120 | 0.842 |
| Rv1047 | 0.031 | NA | -0.430 | 0.859 | -0.462 | 0.829 |
| Rv1048c | -0.046 | 0.999 | -0.095 | 0.933 | -0.049 | 0.956 |
| Rv1049 | -0.040 | NA | -0.290 | 0.918 | -0.251 | 0.906 |
| Rv1050 | -0.442 | NA | -1.133 | 0.830 | -0.691 | 0.835 |
| Rv1051c | -0.301 | NA | -0.783 | 0.846 | -0.482 | 0.860 |
| Rv1052 | -0.087 | 0.999 | 0.134 | 0.879 | 0.221 | 0.816 |
| Rv1056 | 0.026 | 0.999 | -0.022 | 0.962 | -0.048 | 0.899 |
| Rv1057 | -0.523 | 0.591 | -0.577 | 0.180 | -0.055 | 0.936 |
| fadD14 | -0.178 | 0.999 | -0.411 | 0.660 | -0.233 | 0.816 |
| Rv1059 | -0.097 | 0.999 | 0.093 | 0.866 | 0.190 | 0.816 |
| Rv1060 | -0.026 | 0.999 | 0.034 | 0.970 | 0.060 | 0.941 |
| Rv1061 | 0.024 | 0.999 | -0.421 | 0.830 | -0.445 | 0.816 |
| Rv1062 | -0.036 | 0.999 | 0.385 | 0.831 | 0.421 | 0.816 |
| Rv1063c | -0.161 | 0.999 | 0.156 | 0.912 | 0.317 | 0.819 |
| lpqV | 0.320 | 0.999 | 0.166 | 0.914 | -0.154 | 0.896 |
| Rv1065 | -0.152 | 0.999 | 0.116 | 0.922 | 0.268 | 0.819 |
| Rv1066 | -0.115 | 0.999 | -0.579 | 0.832 | -0.694 | 0.816 |
| PE_PGRS19 | 0.104 | NA | -0.024 | 0.987 | -0.127 | 0.936 |
| PE_PGRS20 | -0.115 | NA | -0.371 | 0.846 | -0.255 | 0.846 |
| Rv1069c | 0.123 | 0.999 | 0.071 | 0.890 | -0.052 | 0.902 |
| echA8 | 0.104 | 0.999 | -0.096 | 0.858 | -0.200 | 0.816 |
| echA9 | 0.100 | 0.999 | 0.141 | 0.848 | 0.040 | 0.932 |
| Rv1072 | -0.056 | 0.999 | 0.078 | 0.909 | 0.134 | 0.835 |
| Rv1073 | -0.144 | 0.999 | -0.176 | 0.845 | -0.033 | 0.951 |
| fadA3 | 0.009 | 0.999 | -0.115 | 0.831 | -0.124 | 0.816 |
| Rv1075c | 0.103 | NA | -0.115 | 0.922 | -0.217 | 0.837 |
| lipU | 0.153 | 0.999 | -0.066 | 0.935 | -0.219 | 0.816 |
| cbs | -0.059 | 0.999 | 0.087 | 0.912 | 0.147 | 0.836 |
| Rv1078 | -0.163 | 0.999 | 0.198 | 0.879 | 0.360 | 0.816 |
| metB | -0.024 | 0.999 | 0.071 | 0.939 | 0.095 | 0.896 |
| greA | 0.034 | 0.999 | -0.051 | 0.941 | -0.086 | 0.879 |
| Rv1081c | -0.031 | 0.999 | 0.306 | 0.820 | 0.337 | 0.816 |
| mca | -0.028 | 0.999 | -0.180 | 0.826 | -0.153 | 0.816 |
| Rv1083 | -0.155 | 0.999 | -0.526 | 0.588 | -0.372 | 0.816 |
| Rv1084 | -0.158 | 0.999 | 0.271 | 0.859 | 0.429 | 0.816 |
| Rv1085c | 0.161 | 0.999 | 0.367 | 0.802 | 0.206 | 0.817 |
| Rv1086 | -0.071 | 0.999 | -0.076 | 0.906 | -0.005 | 0.994 |
| PE_PGRS21 | -0.131 | 0.999 | 0.175 | 0.915 | 0.306 | 0.836 |
| PE9 | 0.007 | NA | -0.310 | 0.895 | -0.317 | 0.874 |
| Rv1088a | 0.265 | NA | -0.752 | 0.843 | -1.017 | 0.816 |
| PE_PGRS22 | -0.145 | 0.999 | -0.552 | 0.848 | -0.407 | 0.846 |
| coaA | 0.051 | 0.999 | -0.145 | 0.832 | -0.195 | 0.816 |
| glyA1 | 0.072 | 0.999 | 0.322 | 0.826 | 0.250 | 0.816 |
| desA2 | -0.002 | 0.999 | -0.225 | 0.795 | -0.223 | 0.816 |
| phoH2 | -0.041 | 0.999 | -0.090 | 0.890 | -0.049 | 0.930 |
| Rv1096 | 0.030 | 0.999 | -0.164 | 0.843 | -0.194 | 0.816 |
| Rv1097c | 0.049 | 0.999 | 0.112 | 0.859 | 0.063 | 0.888 |
| fum | 0.036 | 0.999 | -0.072 | 0.927 | -0.108 | 0.868 |
| glpX | 0.089 | 0.999 | -0.158 | 0.927 | -0.248 | 0.866 |
| Rv1100 | -0.004 | 0.999 | -0.167 | 0.859 | -0.163 | 0.816 |
| Rv1101c | -0.039 | 0.999 | -0.106 | 0.911 | -0.066 | 0.930 |
| mazF3 | 0.121 | 0.999 | -0.444 | 0.826 | -0.565 | 0.816 |
| mazE3 | 0.037 | 0.999 | 0.177 | 0.859 | 0.141 | 0.854 |
| Rv1106c | 0.104 | 0.999 | 0.084 | 0.879 | -0.020 | 0.958 |
| xseB | 0.069 | 0.999 | -0.084 | 0.961 | -0.154 | 0.908 |
| xseA | 0.035 | 0.999 | -0.044 | 0.948 | -0.080 | 0.891 |
| Rv1109c | 0.126 | 0.999 | -0.133 | 0.879 | -0.259 | 0.816 |
| lytB2 | -0.012 | 0.999 | -0.008 | 0.981 | 0.005 | 0.988 |
| Rv1111c | 0.045 | 0.999 | 0.370 | 0.716 | 0.326 | 0.816 |
| Rv1112 | -0.047 | 0.999 | -0.068 | 0.958 | -0.021 | 0.985 |
| vapB32 | -0.193 | NA | -0.089 | 0.956 | 0.104 | 0.936 |
| vapC32 | 0.172 | NA | -0.373 | 0.826 | -0.545 | 0.816 |
| Rv1115 | -0.097 | NA | -0.094 | 0.917 | 0.003 | 0.997 |
| Rv1116 | 0.038 | NA | -0.385 | 0.826 | -0.423 | 0.816 |
| Rv1117 | -0.150 | 0.999 | -0.240 | 0.832 | -0.090 | 0.895 |
| Rv1118c | 0.154 | NA | 0.049 | 0.958 | -0.105 | 0.887 |
| Rv1119c | -0.792 | NA | -0.413 | 0.925 | 0.379 | 0.910 |
| Rv1120c | -0.164 | NA | -0.807 | 0.856 | -0.643 | 0.852 |
| zwf1 | -0.093 | 0.999 | 0.046 | 0.958 | 0.139 | 0.854 |
| gnd2 | -0.002 | 0.999 | -0.098 | 0.859 | -0.096 | 0.836 |
| bpoB | -0.089 | 0.999 | 0.176 | 0.837 | 0.265 | 0.816 |
| ephC | -0.127 | 0.999 | 0.710 | 0.510 | 0.836 | 0.816 |
| Rv1125 | -0.220 | NA | -0.034 | 0.973 | 0.187 | 0.854 |
| Rv1126c | 0.023 | 0.999 | -0.266 | 0.820 | -0.289 | 0.816 |
| ppdK | 0.024 | 0.999 | 0.195 | 0.826 | 0.170 | 0.816 |
| Rv1128c | -0.378 | 0.999 | -0.558 | 0.439 | -0.180 | 0.835 |
| Rv1129c | -0.520 | 0.137 | -0.727 | 0.001 | -0.207 | 0.816 |
| prpD | -0.508 | 0.013 | -0.633 | 0.000 | -0.126 | 0.817 |
| prpC | -0.554 | 0.999 | -0.709 | 0.202 | -0.155 | 0.862 |
| Rv1132 | -0.281 | 0.999 | -0.237 | 0.826 | 0.044 | 0.937 |
| metE | 0.103 | 0.999 | 0.112 | 0.896 | 0.009 | 0.989 |
| Rv1134 | 0.054 | NA | -0.355 | 0.848 | -0.302 | 0.829 |
| PPE16 | -0.060 | 0.999 | -0.424 | 0.859 | -0.364 | 0.846 |
| Rv1138a | -0.269 | NA | -0.434 | 0.831 | -0.165 | 0.883 |
| Rv1138c | 0.072 | NA | 0.034 | 0.959 | -0.037 | 0.942 |
| Rv1139c | -0.060 | 0.999 | 0.185 | 0.879 | 0.245 | 0.817 |
| Rv1140 | 0.084 | 0.999 | 0.126 | 0.879 | 0.041 | 0.942 |
| echA11 | 0.032 | 0.999 | 0.157 | 0.831 | 0.124 | 0.817 |
| echA10 | 0.033 | 0.999 | 0.328 | 0.826 | 0.296 | 0.816 |
| mcr | -0.059 | 0.999 | 0.021 | 0.970 | 0.080 | 0.866 |
| Rv1144 | -0.071 | 0.999 | 0.237 | 0.834 | 0.308 | 0.816 |
| Rv1145 | 0.002 | 0.999 | -0.418 | 0.829 | -0.420 | 0.816 |
| Rv1146 | -0.003 | NA | 0.148 | 0.879 | 0.151 | 0.856 |
| Rv1147 | 0.196 | NA | 0.406 | 0.843 | 0.211 | 0.869 |
| Rv1148c | -0.020 | 0.999 | -0.415 | 0.831 | -0.395 | 0.816 |
| Rv1149 | -0.099 | 0.999 | -0.169 | 0.831 | -0.070 | 0.882 |
| Rv1151c | 0.021 | 0.999 | 0.180 | 0.832 | 0.158 | 0.817 |
| Rv1152 | 0.069 | 0.999 | -0.040 | 0.975 | -0.109 | 0.936 |
| omt | 0.098 | 0.999 | 0.041 | 0.926 | -0.057 | 0.877 |
| Rv1154c | 0.181 | 0.999 | 0.020 | 0.978 | -0.161 | 0.854 |
| Rv1155 | 0.034 | 0.999 | -0.003 | 0.995 | -0.037 | 0.947 |
| Rv1155a | -0.071 | NA | 0.247 | 0.848 | 0.318 | 0.816 |
| Rv1156 | -0.007 | 0.999 | -0.229 | 0.701 | -0.222 | 0.816 |
| Rv1157c | 0.093 | 0.999 | 0.326 | 0.769 | 0.233 | 0.816 |
| Rv1158c | 0.219 | 0.999 | 0.264 | 0.856 | 0.045 | 0.962 |
| pimE | -0.048 | 0.999 | 0.147 | 0.879 | 0.195 | 0.831 |
| Rv1159A | -0.080 | 0.999 | -0.125 | 0.877 | -0.045 | 0.936 |
| mutT2 | 0.143 | 0.999 | 0.375 | 0.820 | 0.233 | 0.816 |
| eccE5 | 0.078 | 0.999 | 0.029 | 0.970 | -0.049 | 0.939 |
| eccA5 | 0.094 | 0.999 | -0.144 | 0.879 | -0.238 | 0.816 |
| lppT | 0.013 | 0.999 | 0.243 | 0.832 | 0.230 | 0.816 |
| PPE28 | -0.037 | 0.999 | 0.014 | 0.982 | 0.051 | 0.939 |
| PPE29 | -0.142 | NA | -0.364 | 0.877 | -0.222 | 0.892 |
| PPE30 | -0.174 | NA | -0.156 | 0.916 | 0.018 | 0.988 |
| PE_PGRS32 | -0.032 | 0.999 | -0.368 | 0.859 | -0.336 | 0.838 |
| Rv1804c | -0.079 | NA | -0.209 | 0.879 | -0.130 | 0.899 |
| Rv1805c | -0.093 | NA | -1.437 | 0.832 | -1.344 | 0.816 |
| PE20 | 0.021 | NA | -0.285 | 0.879 | -0.306 | 0.846 |
| PPE32 | -0.049 | 0.999 | 0.104 | 0.925 | 0.153 | 0.868 |
| PPE33 | -0.040 | 0.999 | -0.008 | 0.990 | 0.032 | 0.962 |
| Rv1810 | -0.157 | 0.999 | 0.017 | 0.977 | 0.174 | 0.819 |
| mgtC | -0.207 | 0.999 | -0.012 | 0.983 | 0.195 | 0.816 |
| Rv1812c | -0.592 | 0.999 | -0.730 | 0.413 | -0.138 | 0.888 |
| Rv1813c | -0.506 | 0.999 | -1.068 | 0.000 | -0.561 | 0.816 |
| erg3 | 0.011 | 0.999 | -0.237 | 0.588 | -0.247 | 0.816 |
| Rv1815 | 0.116 | 0.999 | 0.310 | 0.439 | 0.194 | 0.816 |
| Rv1816 | -0.067 | 0.999 | -0.053 | 0.951 | 0.014 | 0.985 |
| Rv1817 | -0.149 | 0.999 | -0.244 | 0.846 | -0.095 | 0.899 |
| PE_PGRS33 | -0.155 | 0.999 | 0.416 | 0.848 | 0.571 | 0.816 |
| bacA | 0.144 | 0.999 | -0.013 | 0.973 | -0.157 | 0.816 |
| ilvG | 0.075 | 0.999 | 0.168 | 0.834 | 0.093 | 0.862 |
| secA2 | 0.057 | 0.999 | -0.089 | 0.879 | -0.146 | 0.816 |
| pgsA2 | 0.060 | 0.999 | -0.044 | 0.940 | -0.104 | 0.838 |
| Rv1823 | 0.070 | 0.999 | -0.066 | 0.923 | -0.136 | 0.835 |
| Rv1824 | 0.056 | 0.999 | 0.314 | 0.832 | 0.258 | 0.817 |
| Rv1825 | 0.113 | 0.999 | -0.199 | 0.843 | -0.312 | 0.816 |
| gcvH | -0.009 | 0.999 | -0.077 | 0.879 | -0.068 | 0.868 |
| garA | -0.111 | 0.999 | 0.035 | 0.951 | 0.146 | 0.816 |
| Rv1828 | -0.039 | 0.999 | 0.361 | 0.832 | 0.400 | 0.816 |
| Rv1829 | 0.116 | 0.999 | -0.211 | 0.829 | -0.327 | 0.816 |
| Rv1830 | -0.025 | 0.999 | -0.037 | 0.951 | -0.012 | 0.983 |
| Rv1831 | -0.084 | 0.999 | -0.162 | 0.859 | -0.078 | 0.899 |
| gcvB | -0.033 | 0.999 | -0.307 | 0.592 | -0.274 | 0.816 |
| Rv1833c | -0.010 | 0.999 | -0.169 | 0.848 | -0.158 | 0.831 |
| lipZ | -0.088 | 0.999 | -0.002 | 0.997 | 0.086 | 0.904 |
| Rv1835c | 0.034 | 0.999 | 0.062 | 0.941 | 0.028 | 0.966 |
| Rv1836c | -0.057 | 0.999 | 0.133 | 0.859 | 0.190 | 0.816 |
| glcB | 0.019 | 0.999 | -0.068 | 0.912 | -0.086 | 0.865 |
| vapC13 | 0.036 | 0.999 | -0.050 | 0.966 | -0.086 | 0.930 |
| vapB13 | 0.010 | NA | -0.233 | 0.859 | -0.243 | 0.835 |
| PE_PGRS34 | -0.351 | 0.999 | -0.297 | 0.878 | 0.054 | 0.967 |
| Rv1841c | 0.039 | 0.999 | -0.060 | 0.939 | -0.099 | 0.875 |
| Rv1842c | 0.015 | 0.999 | 0.090 | 0.886 | 0.075 | 0.888 |
| guaB1 | -0.074 | 0.999 | 0.336 | 0.831 | 0.410 | 0.816 |
| gnd1 | -0.007 | 0.999 | -0.005 | 0.987 | 0.002 | 0.996 |
| blaR | -0.010 | 0.999 | 0.580 | 0.826 | 0.589 | 0.816 |
| blaI | -0.034 | 0.999 | -0.254 | 0.820 | -0.220 | 0.816 |
| Rv1847 | 0.075 | 0.999 | 0.138 | 0.912 | 0.063 | 0.947 |
| ureA | 0.274 | 0.999 | 0.424 | 0.826 | 0.150 | 0.884 |
| ureB | 0.042 | 0.999 | -0.236 | 0.902 | -0.278 | 0.866 |
| ureC | 0.134 | 0.999 | -0.241 | 0.832 | -0.375 | 0.816 |
| ureF | 0.095 | 0.999 | -0.333 | 0.716 | -0.428 | 0.816 |
| ureG | -0.023 | 0.999 | 0.071 | 0.946 | 0.095 | 0.906 |
| ureD | 0.023 | 0.999 | -0.020 | 0.978 | -0.042 | 0.956 |
| ndh | -0.272 | 0.999 | -0.191 | 0.831 | 0.081 | 0.879 |
| Rv1855c | -0.068 | 0.999 | 0.130 | 0.879 | 0.198 | 0.816 |
| Rv1856c | 0.271 | 0.999 | 0.523 | 0.659 | 0.252 | 0.816 |
| modA | 0.297 | 0.999 | 0.625 | 0.307 | 0.328 | 0.816 |
| modB | 0.300 | NA | 0.625 | 0.669 | 0.325 | 0.816 |
| modC | -0.073 | 0.999 | 0.266 | 0.826 | 0.339 | 0.816 |
| apa | 0.213 | 0.999 | 0.056 | 0.964 | -0.157 | 0.882 |
| Rv1861 | -0.062 | 0.999 | -0.059 | 0.974 | 0.002 | 0.999 |
| adhA | 0.130 | 0.999 | 0.284 | 0.826 | 0.155 | 0.835 |
| Rv1863c | 0.026 | 0.999 | -0.050 | 0.937 | -0.076 | 0.883 |
| Rv1864c | 0.078 | 0.999 | -0.161 | 0.877 | -0.239 | 0.816 |
| Rv1865c | 0.083 | 0.999 | 0.154 | 0.859 | 0.070 | 0.909 |
| Rv1866 | 0.069 | 0.999 | 0.167 | 0.831 | 0.098 | 0.843 |
| Rv1867 | 0.090 | 0.999 | 0.040 | 0.960 | -0.050 | 0.945 |
| Rv1868 | 0.008 | 0.999 | -0.231 | 0.826 | -0.239 | 0.816 |
| Rv1869c | -0.089 | 0.999 | -0.045 | 0.946 | 0.044 | 0.932 |
| Rv1870c | 0.090 | 0.999 | -0.192 | 0.879 | -0.282 | 0.817 |
| Rv1871c | 0.157 | 0.999 | -0.156 | 0.859 | -0.313 | 0.816 |
| lldD2 | 0.033 | 0.999 | 0.071 | 0.922 | 0.038 | 0.943 |
| Rv1873 | -0.069 | NA | -0.252 | 0.858 | -0.183 | 0.862 |
| Rv1874 | 0.034 | 0.999 | -0.053 | 0.951 | -0.087 | 0.900 |
| Rv1875 | 0.077 | 0.999 | 0.019 | 0.974 | -0.058 | 0.921 |
| bfrA | 0.185 | 0.999 | -0.162 | 0.907 | -0.347 | 0.816 |
| Rv1877 | 0.029 | 0.999 | 0.316 | 0.660 | 0.287 | 0.816 |
| glnA3 | -0.056 | 0.999 | -0.246 | 0.831 | -0.190 | 0.817 |
| Rv1879 | 0.029 | NA | 0.095 | 0.915 | 0.066 | 0.925 |
| cyp140 | 0.007 | 0.999 | -0.246 | 0.820 | -0.254 | 0.816 |
| lppE | -0.003 | 0.999 | -0.076 | 0.895 | -0.073 | 0.879 |
| Rv1882c | 0.006 | 0.999 | 0.058 | 0.913 | 0.052 | 0.897 |
| Rv1883c | -0.018 | 0.999 | -0.024 | 0.970 | -0.005 | 0.992 |
| rpfC | 0.054 | 0.999 | -0.005 | 0.989 | -0.059 | 0.892 |
| Rv1885c | 0.082 | 0.999 | -0.185 | 0.848 | -0.267 | 0.816 |
| fbpB | 0.154 | 0.999 | 0.339 | 0.826 | 0.185 | 0.837 |
| Rv1887 | 0.054 | 0.999 | 0.193 | 0.831 | 0.139 | 0.831 |
| Rv1890c | -0.002 | 0.999 | -0.270 | 0.831 | -0.269 | 0.816 |
| Rv1891 | 0.103 | 0.999 | 0.338 | 0.832 | 0.235 | 0.837 |
| Rv1892 | 0.311 | 0.999 | 0.244 | 0.829 | -0.066 | 0.917 |
| Rv1893 | 0.070 | 0.999 | -0.421 | 0.820 | -0.491 | 0.816 |
| Rv1894c | -0.120 | 0.999 | -0.198 | 0.820 | -0.077 | 0.859 |
| Rv1895 | -0.037 | 0.999 | 0.009 | 0.985 | 0.046 | 0.930 |
| Rv1896c | 0.030 | 0.999 | -0.109 | 0.881 | -0.139 | 0.837 |
| Rv1897c | -0.072 | NA | -0.017 | 0.981 | 0.055 | 0.942 |
| Rv1898 | -0.076 | 0.999 | -0.430 | 0.826 | -0.354 | 0.816 |
| lppD | -0.171 | 0.999 | 0.151 | 0.900 | 0.322 | 0.816 |
| lipJ | -0.005 | 0.999 | 0.130 | 0.895 | 0.135 | 0.868 |
| cinA | 0.066 | 0.999 | 0.141 | 0.831 | 0.075 | 0.852 |
| nanT | -0.040 | 0.999 | 0.087 | 0.915 | 0.127 | 0.855 |
| Rv1903 | -0.050 | 0.999 | -0.523 | 0.772 | -0.473 | 0.816 |
| Rv1904 | 0.181 | 0.999 | -0.189 | 0.834 | -0.370 | 0.816 |
| aao | -0.131 | 0.999 | 0.121 | 0.879 | 0.252 | 0.816 |
| Rv1906c | -0.073 | 0.999 | 0.002 | 0.995 | 0.075 | 0.868 |
| Rv1907c | -0.101 | 0.999 | -0.262 | 0.772 | -0.162 | 0.816 |
| katG | -0.180 | 0.999 | -0.208 | 0.767 | -0.028 | 0.940 |
| furA | -0.052 | 0.999 | -0.131 | 0.843 | -0.079 | 0.859 |
| Rv1910c | -0.089 | 0.999 | 0.029 | 0.963 | 0.118 | 0.838 |
| lppC | -0.131 | 0.999 | 0.266 | 0.831 | 0.397 | 0.816 |
| fadB5 | 0.182 | 0.999 | -0.032 | 0.970 | -0.214 | 0.819 |
| Rv1913 | -0.006 | 0.999 | 0.168 | 0.848 | 0.174 | 0.816 |
| Rv1914c | -0.073 | 0.999 | -0.052 | 0.940 | 0.021 | 0.969 |
| aceAa | -0.050 | 0.999 | -0.021 | 0.974 | 0.029 | 0.962 |
| aceAb | 0.078 | 0.999 | -0.032 | 0.967 | -0.110 | 0.866 |
| PPE34 | -0.150 | NA | -0.964 | NA | -0.814 | NA |
| PPE35 | 0.080 | 0.999 | -0.215 | 0.891 | -0.295 | 0.837 |
| Rv1919c | 0.109 | 0.999 | 0.035 | 0.975 | -0.074 | 0.947 |
| Rv1920 | 0.192 | 0.999 | 0.027 | 0.961 | -0.166 | 0.816 |
| lppF | -0.053 | NA | -0.447 | 0.832 | -0.395 | 0.816 |
| Rv1922 | -0.018 | 0.999 | -0.342 | 0.826 | -0.324 | 0.816 |
| lipD | 0.044 | 0.999 | -0.042 | 0.948 | -0.087 | 0.875 |
| Rv1924c | -0.004 | 0.999 | 0.329 | 0.832 | 0.333 | 0.816 |
| fadD31 | -0.006 | 0.999 | -0.037 | 0.931 | -0.031 | 0.930 |
| mpt63 | 0.038 | 0.999 | 0.077 | 0.922 | 0.039 | 0.947 |
| Rv1927 | -0.051 | 0.999 | -0.070 | 0.927 | -0.019 | 0.976 |
| Rv1928c | -0.084 | 0.999 | -0.540 | 0.678 | -0.457 | 0.816 |
| Rv1929c | -0.050 | 0.999 | 0.318 | 0.848 | 0.368 | 0.816 |
| Rv1930c | -0.064 | NA | 0.347 | 0.859 | 0.411 | 0.817 |
| Rv1931c | -0.008 | NA | 0.072 | 0.931 | 0.080 | 0.904 |
| tpx | 0.007 | 0.999 | 0.094 | 0.886 | 0.087 | 0.887 |
| fadE18 | -0.173 | NA | -0.305 | 0.895 | -0.133 | 0.944 |
| fadE17 | -0.015 | NA | -0.067 | 0.962 | -0.052 | 0.963 |
| echA13 | -0.069 | NA | -0.496 | 0.825 | -0.427 | 0.816 |
| Rv1936 | -0.220 | NA | -0.354 | 0.829 | -0.133 | 0.882 |
| Rv1937 | -0.092 | 0.999 | -0.276 | 0.826 | -0.184 | 0.816 |
| ephB | -0.292 | NA | -0.198 | 0.834 | 0.093 | 0.879 |
| Rv1939 | 0.011 | NA | -0.078 | 0.948 | -0.089 | 0.930 |
| ribA1 | -0.165 | NA | 0.198 | 0.859 | 0.363 | 0.816 |
| Rv1941 | 0.040 | 0.999 | -0.008 | 0.987 | -0.048 | 0.930 |
| mazF5 | -0.044 | 0.999 | -0.154 | 0.846 | -0.109 | 0.840 |
| mazE5 | 0.004 | 0.999 | -0.092 | 0.922 | -0.095 | 0.896 |
| Rv1944c | 0.008 | 0.999 | 0.046 | 0.945 | 0.038 | 0.942 |
| Rv1945 | -0.039 | 0.999 | -0.224 | 0.848 | -0.185 | 0.837 |
| lppG | -0.270 | NA | 0.084 | 0.962 | 0.354 | 0.836 |
| Rv1947 | -0.076 | 0.999 | 0.102 | 0.918 | 0.178 | 0.837 |
| Rv1948c | -0.110 | 0.999 | -0.388 | 0.834 | -0.278 | 0.836 |
| Rv1950c | 0.025 | NA | 0.019 | 0.982 | -0.006 | 0.996 |
| Rv1951c | -0.078 | NA | 0.241 | 0.848 | 0.319 | 0.816 |
| vapB14 | -0.246 | 0.999 | 0.527 | 0.832 | 0.773 | 0.816 |
| vapC14 | 0.141 | 0.999 | -0.052 | 0.947 | -0.193 | 0.854 |
| Rv1954c | -0.083 | NA | -0.474 | 0.843 | -0.391 | 0.827 |
| higB | -0.025 | 0.999 | -0.335 | 0.826 | -0.310 | 0.816 |
| higA | -0.024 | 0.999 | -0.448 | 0.652 | -0.424 | 0.816 |
| Rv1957 | 0.023 | 0.999 | -0.337 | 0.856 | -0.360 | 0.816 |
| Rv1958c | -0.158 | NA | -0.441 | 0.845 | -0.282 | 0.854 |
| parE1 | -0.035 | 0.999 | -0.461 | 0.825 | -0.426 | 0.816 |
| parD1 | -0.084 | NA | -0.025 | 0.982 | 0.059 | 0.958 |
| Rv1961 | -0.156 | 0.999 | -0.132 | 0.859 | 0.024 | 0.962 |
| vapB35 | 0.062 | NA | 0.167 | 0.848 | 0.105 | 0.860 |
| vapC35 | -0.020 | NA | -0.213 | 0.858 | -0.194 | 0.837 |
| mce3R | 0.041 | 0.999 | -0.126 | 0.879 | -0.167 | 0.820 |
| yrbE3A | -0.170 | NA | -0.342 | 0.831 | -0.172 | 0.862 |
| yrbE3B | -0.126 | NA | -0.333 | 0.848 | -0.208 | 0.868 |
| mce3A | -0.089 | NA | -0.395 | 0.843 | -0.306 | 0.835 |
| mce3B | -0.787 | NA | -0.871 | 0.659 | -0.084 | 0.947 |
| mce3C | -0.648 | NA | -0.909 | 0.555 | -0.261 | 0.854 |
| mce3D | -0.480 | NA | -1.532 | NA | -1.051 | NA |
| lprM | -0.503 | NA | -0.552 | 0.820 | -0.048 | 0.962 |
| mce3F | -0.141 | NA | -0.516 | 0.832 | -0.374 | 0.835 |
| Rv1972 | -0.072 | NA | -0.716 | 0.846 | -0.644 | 0.819 |
| Rv1973 | -0.188 | NA | -0.354 | 0.879 | -0.166 | 0.924 |
| Rv1974 | -0.680 | NA | -0.753 | 0.853 | -0.074 | 0.979 |
| tesB2 | 0.037 | 0.999 | 0.152 | 0.832 | 0.114 | 0.824 |
| snzP | -0.082 | 0.999 | 0.171 | 0.826 | 0.253 | 0.816 |
| pdxH | -0.175 | NA | -0.090 | 0.895 | 0.085 | 0.882 |
| PPE42 | 0.076 | 0.999 | 0.274 | 0.820 | 0.198 | 0.816 |
| Rv2609c | 0.132 | 0.999 | 0.320 | 0.826 | 0.188 | 0.821 |
| pimA | 0.030 | 0.999 | -0.115 | 0.879 | -0.145 | 0.836 |
| Rv2611c | 0.056 | 0.999 | 0.274 | 0.859 | 0.218 | 0.854 |
| pgsA1 | -0.085 | 0.999 | 0.065 | 0.936 | 0.150 | 0.837 |
| Rv2613c | 0.066 | 0.999 | -0.162 | 0.879 | -0.227 | 0.819 |
| Rv2614A | -0.493 | NA | 0.025 | 0.991 | 0.519 | 0.847 |
| thrS | 0.084 | 0.999 | 0.036 | 0.933 | -0.048 | 0.892 |
| PE_PGRS45 | -0.086 | NA | -0.416 | 0.859 | -0.330 | 0.862 |
| Rv2616 | 0.011 | NA | 0.459 | 0.669 | 0.448 | 0.816 |
| Rv2617c | -0.043 | 0.999 | 0.189 | 0.834 | 0.232 | 0.816 |
| Rv2618 | -0.165 | NA | -0.097 | 0.927 | 0.068 | 0.936 |
| Rv2619c | -0.044 | 0.999 | -0.221 | 0.826 | -0.177 | 0.816 |
| Rv2620c | -0.316 | NA | -0.079 | 0.959 | 0.237 | 0.857 |
| Rv2621c | -0.230 | 0.999 | 0.290 | 0.826 | 0.521 | 0.816 |
| Rv2622 | -0.116 | NA | -0.139 | 0.879 | -0.024 | 0.974 |
| TB31.7 | -0.760 | 0.005 | -0.931 | 0.000 | -0.171 | 0.819 |
| Rv2624c | -0.720 | 0.044 | -1.092 | 0.000 | -0.372 | 0.816 |
| Rv2625c | -0.682 | 0.999 | -0.637 | 0.669 | 0.045 | 0.963 |
| hrp1 | -0.667 | 0.073 | -1.048 | 0.000 | -0.381 | 0.816 |
| Rv2627c | -0.864 | 0.001 | -0.977 | 0.000 | -0.113 | 0.862 |
| Rv2628 | -0.816 | 0.033 | -1.298 | 0.000 | -0.482 | 0.816 |
| Rv2629 | -0.561 | 0.002 | -0.823 | 0.000 | -0.262 | 0.816 |
| Rv2630 | -0.520 | 0.374 | -0.779 | 0.002 | -0.259 | 0.816 |
| Rv2631 | -0.678 | 0.000 | -0.728 | 0.000 | -0.050 | 0.892 |
| Rv2632c | -0.097 | 0.999 | 0.312 | 0.769 | 0.408 | 0.816 |
| Rv2633c | 0.161 | 0.999 | -0.202 | 0.885 | -0.363 | 0.816 |
| PE_PGRS46 | -0.179 | 0.999 | -0.863 | 0.832 | -0.684 | 0.822 |
| Rv2635 | 0.125 | NA | -2.045 | NA | -2.170 | NA |
| Rv2636 | 0.186 | NA | 0.057 | 0.922 | -0.129 | 0.819 |
| dedA | 0.053 | 0.999 | 0.319 | 0.845 | 0.266 | 0.829 |
| Rv2638 | -0.049 | 0.999 | -0.050 | 0.948 | 0.000 | 0.999 |
| Rv2639c | -0.234 | 0.999 | 0.165 | 0.886 | 0.399 | 0.816 |
| Rv2640c | -0.335 | 0.999 | -0.274 | 0.832 | 0.061 | 0.936 |
| cadI | -0.162 | 0.999 | -0.036 | 0.962 | 0.126 | 0.857 |
| Rv2642 | 0.023 | NA | 0.422 | 0.826 | 0.398 | 0.816 |
| arsC | -0.130 | 0.999 | -0.019 | 0.973 | 0.112 | 0.840 |
| Rv2644c | 0.201 | NA | 0.198 | 0.879 | -0.003 | 0.997 |
| Rv2645 | 0.003 | 0.999 | -0.006 | 0.995 | -0.008 | 0.994 |
| Rv2646 | -0.040 | 0.999 | -0.047 | 0.962 | -0.007 | 0.994 |
| Rv2647 | -0.245 | 0.999 | 0.295 | 0.848 | 0.540 | 0.816 |
| Rv2650c | -0.132 | 0.999 | 0.051 | 0.951 | 0.184 | 0.831 |
| Rv2651c | -0.163 | 0.999 | 0.043 | 0.959 | 0.206 | 0.816 |
| Rv2652c | 0.026 | NA | 0.206 | 0.879 | 0.180 | 0.865 |
| Rv2653c | 0.188 | NA | 0.001 | 0.999 | -0.187 | 0.930 |
| Rv2654c | 0.142 | NA | 0.192 | 0.962 | 0.050 | 0.988 |
| Rv2655c | 0.112 | NA | -0.131 | 0.922 | -0.243 | 0.838 |
| Rv2656c | 0.149 | NA | 0.069 | 0.961 | -0.080 | 0.942 |
| Rv2657c | -0.218 | 0.999 | 0.601 | 0.820 | 0.819 | 0.816 |
| Rv2658c | -0.455 | NA | -0.526 | 0.750 | -0.072 | 0.937 |
| Rv2659c | -0.278 | NA | -1.523 | NA | -1.245 | NA |
| Rv2660c | 0.301 | NA | -0.587 | 0.859 | -0.888 | 0.816 |
| Rv2661c | -0.171 | NA | 0.196 | 0.848 | 0.368 | 0.816 |
| Rv2662 | -0.244 | NA | -0.568 | 0.829 | -0.324 | 0.837 |
| Rv2663 | -0.261 | 0.999 | -0.911 | 0.826 | -0.650 | 0.817 |
| Rv2664 | -0.152 | 0.999 | -0.127 | 0.935 | 0.025 | 0.986 |
| Rv2665 | -0.051 | 0.999 | 0.124 | 0.927 | 0.175 | 0.876 |
| clpC2 | -0.143 | NA | -0.561 | 0.843 | -0.418 | 0.836 |
| Rv2668 | 0.018 | 0.999 | 0.280 | 0.848 | 0.262 | 0.822 |
| Rv2669 | -0.016 | NA | -0.030 | 0.973 | -0.014 | 0.987 |
| Rv2670c | 0.002 | NA | -0.030 | 0.970 | -0.031 | 0.961 |
| ribD | -0.149 | 0.999 | -0.072 | 0.918 | 0.076 | 0.891 |
| Rv2672 | 0.147 | 0.999 | -0.121 | 0.879 | -0.268 | 0.816 |
| aftC | 0.169 | 0.999 | -0.087 | 0.879 | -0.256 | 0.816 |
| msrB | 0.083 | 0.999 | 0.011 | 0.978 | -0.071 | 0.879 |
| Rv2675c | -0.003 | 0.999 | 0.058 | 0.913 | 0.061 | 0.884 |
| Rv2676c | -0.029 | 0.999 | 0.062 | 0.962 | 0.091 | 0.932 |
| hemY | -0.115 | 0.999 | 0.432 | 0.848 | 0.546 | 0.816 |
| hemE | -0.056 | 0.999 | 0.246 | 0.834 | 0.302 | 0.816 |
| echA15 | 0.090 | 0.999 | 0.508 | 0.826 | 0.418 | 0.816 |
| Rv2680 | 0.002 | 0.999 | 0.111 | 0.866 | 0.109 | 0.837 |
| Rv2681 | 0.055 | 0.999 | 0.311 | 0.826 | 0.256 | 0.816 |
| dxs1 | 0.030 | 0.999 | 0.082 | 0.890 | 0.051 | 0.918 |
| Rv2683 | 0.070 | 0.999 | -0.052 | 0.933 | -0.122 | 0.836 |
| arsA | 0.057 | 0.999 | 0.083 | 0.940 | 0.025 | 0.977 |
| arsB1 | 0.050 | 0.999 | 0.014 | 0.977 | -0.037 | 0.942 |
| Rv2686c | 0.050 | 0.999 | 0.179 | 0.848 | 0.128 | 0.846 |
| Rv2687c | 0.066 | NA | 0.095 | 0.931 | 0.030 | 0.974 |
| Rv2688c | -0.064 | 0.999 | 0.106 | 0.903 | 0.170 | 0.836 |
| Rv2689c | -0.054 | 0.999 | 0.179 | 0.848 | 0.233 | 0.816 |
| Rv2690c | 0.077 | 0.999 | -0.152 | 0.886 | -0.230 | 0.831 |
| ceoB | 0.118 | 0.999 | 0.124 | 0.879 | 0.006 | 0.994 |
| ceoC | 0.063 | 0.999 | 0.096 | 0.922 | 0.032 | 0.966 |
| Rv2693c | -0.097 | 0.999 | 0.106 | 0.927 | 0.203 | 0.846 |
| Rv2694c | -0.011 | 0.999 | -0.338 | 0.367 | -0.328 | 0.816 |
| Rv2695 | -0.088 | NA | 0.108 | 0.915 | 0.197 | 0.835 |
| Rv2696c | -0.029 | 0.999 | 0.405 | 0.820 | 0.435 | 0.816 |
| dut | 0.051 | 0.999 | 0.620 | 0.820 | 0.569 | 0.816 |
| Rv2698 | -0.111 | 0.999 | 0.224 | 0.866 | 0.335 | 0.816 |
| Rv2699c | -0.014 | 0.999 | -0.069 | 0.931 | -0.055 | 0.932 |
| Rv2700 | -0.095 | 0.999 | 0.152 | 0.883 | 0.246 | 0.817 |
| suhB | -0.070 | 0.999 | 0.149 | 0.907 | 0.218 | 0.846 |
| ppgK | -0.089 | 0.999 | -0.027 | 0.969 | 0.062 | 0.905 |
| sigA | -0.035 | 0.999 | -0.044 | 0.931 | -0.009 | 0.984 |
| Rv2704 | -0.108 | 0.999 | -0.332 | 0.826 | -0.224 | 0.819 |
| Rv2705c | -0.339 | 0.999 | -0.241 | 0.848 | 0.098 | 0.899 |
| Rv2706c | -0.222 | NA | -0.881 | 0.769 | -0.658 | 0.816 |
| Rv2707 | -0.338 | 0.999 | -0.296 | 0.826 | 0.042 | 0.951 |
| Rv2708c | -0.113 | 0.999 | 0.147 | 0.859 | 0.261 | 0.816 |
| Rv2709 | -0.095 | 0.999 | -0.140 | 0.879 | -0.045 | 0.942 |
| sigB | -0.210 | 0.999 | -0.255 | 0.769 | -0.045 | 0.924 |
| ideR | -0.180 | 0.999 | -0.164 | 0.826 | 0.015 | 0.968 |
| Rv2712c | 0.138 | 0.999 | -0.039 | 0.970 | -0.176 | 0.842 |
| sthA | -0.003 | 0.999 | -0.019 | 0.958 | -0.016 | 0.954 |
| Rv2714 | 0.051 | 0.999 | -0.124 | 0.848 | -0.175 | 0.816 |
| Rv2715 | 0.043 | 0.999 | 0.123 | 0.907 | 0.080 | 0.925 |
| Rv2716 | 0.108 | 0.999 | -0.451 | 0.831 | -0.559 | 0.816 |
| Rv2717c | 0.043 | NA | 0.034 | 0.973 | -0.009 | 0.994 |
| nrdR | -0.090 | 0.999 | -0.048 | 0.970 | 0.042 | 0.967 |
| Rv2719c | 0.023 | 0.999 | 0.214 | 0.832 | 0.191 | 0.817 |
| lexA | 0.018 | 0.999 | 0.095 | 0.886 | 0.078 | 0.891 |
| Rv2721c | 0.085 | 0.999 | 0.251 | 0.826 | 0.166 | 0.816 |
| Rv2722 | 0.112 | NA | 0.001 | 0.999 | -0.112 | 0.936 |
| Rv2723 | 0.149 | 0.999 | -0.041 | 0.971 | -0.190 | 0.862 |
| fadE20 | 0.193 | 0.999 | 0.139 | 0.900 | -0.054 | 0.951 |
| hflX | 0.120 | 0.999 | 0.038 | 0.946 | -0.082 | 0.862 |
| dapF | 0.024 | 0.999 | 0.211 | 0.826 | 0.187 | 0.816 |
| miaA | 0.055 | 0.999 | -0.034 | 0.970 | -0.089 | 0.898 |
| Rv2728c | -0.197 | 0.999 | 0.069 | 0.953 | 0.265 | 0.819 |
| Rv2729c | -0.178 | 0.999 | -0.253 | 0.826 | -0.076 | 0.896 |
| Rv2730 | 0.081 | NA | -0.049 | 0.973 | -0.130 | 0.925 |
| Rv2731 | 0.032 | 0.999 | 0.236 | 0.831 | 0.204 | 0.816 |
| Rv2732c | -0.040 | 0.999 | 0.405 | 0.831 | 0.445 | 0.816 |
| Rv2733c | 0.006 | 0.999 | 0.063 | 0.931 | 0.057 | 0.924 |
| Rv2734 | -0.045 | 0.999 | -0.297 | 0.831 | -0.252 | 0.816 |
| Rv2735c | 0.082 | 0.999 | -0.410 | 0.848 | -0.492 | 0.816 |
| recX | 0.014 | 0.999 | 0.212 | 0.859 | 0.198 | 0.839 |
| recA | 0.021 | 0.999 | -0.089 | 0.846 | -0.109 | 0.816 |
| Rv2738c | -0.047 | 0.999 | 0.155 | 0.859 | 0.202 | 0.816 |
| Rv2739c | 0.193 | 0.999 | 0.505 | 0.820 | 0.311 | 0.816 |
| ephG | 0.117 | 0.999 | 0.153 | 0.895 | 0.036 | 0.969 |
| PE_PGRS47 | -0.079 | 0.999 | -0.060 | 0.970 | 0.019 | 0.988 |
| Rv2742c | -0.013 | 0.999 | -0.247 | 0.843 | -0.233 | 0.817 |
| Rv2743c | 0.021 | 0.999 | 0.086 | 0.914 | 0.065 | 0.918 |
| 35kd_ag | -0.105 | 0.999 | -0.160 | 0.848 | -0.054 | 0.925 |
| clgR | -0.123 | 0.999 | 0.010 | 0.989 | 0.133 | 0.868 |
| pgsA3 | 0.161 | 0.999 | 0.303 | 0.848 | 0.142 | 0.888 |
| argA | 0.111 | 0.999 | -0.038 | 0.970 | -0.149 | 0.862 |
| ftsK | 0.020 | 0.999 | 0.229 | 0.826 | 0.208 | 0.816 |
| Rv2749 | -0.055 | 0.999 | 0.092 | 0.922 | 0.147 | 0.859 |
| Rv2750 | -0.112 | 0.999 | 0.027 | 0.975 | 0.138 | 0.882 |
| Rv2751 | -0.100 | 0.999 | 0.259 | 0.854 | 0.359 | 0.816 |
| Rv2752c | 0.037 | 0.999 | 0.037 | 0.965 | 0.000 | 0.999 |
| dapA | 0.006 | 0.999 | 0.354 | 0.826 | 0.347 | 0.816 |
| thyX | 0.072 | 0.999 | -0.341 | 0.879 | -0.413 | 0.838 |
| hsdM | 0.085 | 0.999 | -0.105 | 0.948 | -0.190 | 0.882 |
| vapC21 | 0.182 | 0.999 | -0.103 | 0.895 | -0.285 | 0.816 |
| vapB21 | 0.230 | NA | 0.094 | 0.940 | -0.137 | 0.892 |
| vapC42 | 0.065 | NA | 0.007 | 0.988 | -0.057 | 0.906 |
| vapB42 | -0.054 | NA | -0.553 | 0.831 | -0.499 | 0.816 |
| hsdS | 0.061 | 0.999 | -0.038 | 0.948 | -0.100 | 0.852 |
| Rv2762c | 0.321 | 0.999 | 0.264 | 0.879 | -0.057 | 0.961 |
| dfrA | -0.012 | 0.999 | 0.459 | 0.829 | 0.471 | 0.816 |
| thyA | 0.061 | 0.999 | -0.159 | 0.848 | -0.220 | 0.816 |
| Rv2765 | -0.158 | NA | 0.061 | 0.958 | 0.219 | 0.836 |
| Rv2766c | 0.176 | 0.999 | -0.010 | 0.988 | -0.186 | 0.824 |
| Rv2767c | -0.126 | NA | -0.417 | 0.826 | -0.291 | 0.816 |
| PPE43 | 0.061 | 0.999 | 0.197 | 0.837 | 0.136 | 0.838 |
| PE27 | -0.029 | NA | 0.035 | 0.975 | 0.064 | 0.955 |
| PPE44 | -0.052 | 0.999 | 0.092 | 0.903 | 0.144 | 0.837 |
| Rv2771c | 0.136 | 0.999 | -0.179 | 0.871 | -0.316 | 0.816 |
| Rv2772c | 0.164 | 0.999 | 0.476 | 0.021 | 0.312 | 0.816 |
| dapB | 0.045 | 0.999 | -0.158 | 0.846 | -0.202 | 0.816 |
| Rv2774c | -0.069 | NA | -0.483 | 0.853 | -0.413 | 0.837 |
| Rv2775 | 0.023 | 0.999 | -0.081 | 0.879 | -0.105 | 0.835 |
| Rv2776c | -0.167 | 0.999 | -0.031 | 0.971 | 0.136 | 0.866 |
| Rv2777c | -0.109 | 0.999 | -0.156 | 0.826 | -0.047 | 0.899 |
| Rv2778c | 0.046 | 0.999 | -0.069 | 0.948 | -0.115 | 0.892 |
| Rv2779c | -0.071 | 0.999 | 0.365 | 0.660 | 0.436 | 0.816 |
| ald | 0.007 | 0.999 | 0.016 | 0.967 | 0.009 | 0.976 |
| Rv2781c | 0.023 | 0.999 | 0.209 | 0.859 | 0.185 | 0.846 |
| pepR | -0.026 | 0.999 | 0.398 | 0.832 | 0.424 | 0.816 |
| whiB3 | 0.205 | 0.999 | -0.093 | 0.931 | -0.298 | 0.816 |
| groEL1 | -0.189 | 0.999 | -0.234 | 0.728 | -0.045 | 0.909 |
| groES | -0.496 | 0.999 | -0.271 | 0.948 | 0.225 | 0.944 |
| gcp | -0.043 | 0.999 | 0.195 | 0.859 | 0.237 | 0.817 |
| rimI | -0.068 | 0.999 | 0.142 | 0.951 | 0.210 | 0.909 |
| Rv3421c | -0.064 | 0.999 | 0.266 | 0.877 | 0.330 | 0.819 |
| Rv3422c | 0.058 | 0.999 | -0.228 | 0.879 | -0.286 | 0.835 |
| alr | 0.038 | 0.999 | 0.166 | 0.848 | 0.128 | 0.846 |
| Rv3424c | 0.059 | 0.999 | -0.314 | 0.826 | -0.373 | 0.816 |
| PPE57 | -0.319 | 0.999 | -0.429 | 0.832 | -0.111 | 0.932 |
| PPE58 | -0.223 | 0.999 | -0.429 | 0.660 | -0.205 | 0.817 |
| Rv3427c | -0.105 | 0.999 | -0.602 | 0.848 | -0.497 | 0.836 |
| Rv3428c | -0.323 | NA | -0.764 | 0.826 | -0.441 | 0.835 |
| PPE59 | 0.068 | 0.999 | -0.161 | 0.919 | -0.229 | 0.862 |
| Rv3430a | 0.099 | 0.999 | 0.375 | 0.826 | 0.276 | 0.816 |
| Rv3430c | 0.070 | 0.999 | -0.172 | 0.855 | -0.242 | 0.816 |
| gadB | 0.007 | 0.999 | 0.134 | 0.848 | 0.127 | 0.819 |
| Rv3433c | -0.113 | 0.999 | 0.658 | 0.820 | 0.771 | 0.816 |
| Rv3434c | 0.159 | 0.999 | 0.069 | 0.899 | -0.091 | 0.855 |
| Rv3435c | -0.034 | 0.999 | 0.218 | 0.859 | 0.252 | 0.819 |
| glmS | 0.064 | 0.999 | 0.218 | 0.848 | 0.154 | 0.854 |
| Rv3437 | 0.061 | 0.999 | -0.008 | 0.984 | -0.069 | 0.868 |
| Rv3438 | -0.030 | 0.999 | 0.509 | 0.826 | 0.540 | 0.816 |
| Rv3439c | -0.017 | 0.999 | -0.152 | 0.826 | -0.135 | 0.816 |
| Rv3440c | -0.004 | NA | 0.171 | 0.879 | 0.175 | 0.838 |
| mrsA | 0.068 | 0.999 | 0.538 | 0.826 | 0.470 | 0.816 |
| rpsI | 0.023 | 0.999 | 0.214 | 0.879 | 0.191 | 0.866 |
| rplM | 0.095 | 0.999 | -0.094 | 0.848 | -0.189 | 0.816 |
| esxT | 0.053 | NA | 0.544 | 0.848 | 0.491 | 0.819 |
| esxU | 0.395 | NA | 0.130 | 0.927 | -0.266 | 0.837 |
| Rv3446c | -0.110 | NA | -0.238 | 0.922 | -0.128 | 0.942 |
| eccC4 | -0.039 | NA | -0.341 | 0.883 | -0.302 | 0.875 |
| eccD4 | -0.222 | NA | -0.516 | 0.848 | -0.295 | 0.878 |
| mycP4 | -0.219 | NA | -0.128 | 0.941 | 0.091 | 0.946 |
| eccB4 | 0.032 | 0.999 | 0.448 | 0.527 | 0.416 | 0.816 |
| cut3 | -0.076 | 0.999 | -0.057 | 0.879 | 0.018 | 0.946 |
| cut4 | 0.013 | 0.999 | 0.029 | 0.959 | 0.015 | 0.972 |
| truA | 0.023 | 0.999 | 0.277 | 0.825 | 0.254 | 0.816 |
| rplQ | -0.073 | 0.999 | 0.021 | 0.970 | 0.094 | 0.854 |
| rpoA | 0.061 | 0.999 | 0.081 | 0.895 | 0.020 | 0.967 |
| rpsD | 0.055 | 0.999 | -0.050 | 0.879 | -0.104 | 0.816 |
| rpsK | 0.037 | 0.999 | 0.066 | 0.895 | 0.029 | 0.942 |
| rpsM | 0.090 | 0.999 | -0.149 | 0.875 | -0.239 | 0.816 |
| rpmJ | 0.264 | 0.999 | -0.218 | 0.886 | -0.482 | 0.816 |
| infA | 0.116 | 0.999 | -0.364 | 0.918 | -0.480 | 0.866 |
| Rv3463 | -0.100 | 0.999 | 0.008 | 0.988 | 0.108 | 0.854 |
| rmlB | 0.147 | 0.999 | 0.043 | 0.951 | -0.103 | 0.866 |
| rmlC | 0.150 | 0.999 | 0.157 | 0.858 | 0.006 | 0.993 |
| Rv3466 | -0.058 | 0.999 | -0.491 | 0.846 | -0.433 | 0.819 |
| Rv3467 | -0.066 | 0.999 | -1.054 | 0.848 | -0.988 | 0.827 |
| Rv3468c | 0.016 | 0.999 | -0.184 | 0.859 | -0.201 | 0.825 |
| mhpE | -0.006 | 0.999 | -0.100 | 0.951 | -0.094 | 0.942 |
| ilvB2 | -0.077 | 0.999 | 0.154 | 0.848 | 0.231 | 0.816 |
| Rv3471c | -0.041 | NA | -0.626 | 0.848 | -0.586 | 0.819 |
| Rv3472 | 0.043 | 0.999 | 0.282 | 0.831 | 0.239 | 0.816 |
| bpoA | -0.050 | NA | -0.763 | 0.848 | -0.713 | 0.831 |
| kgtP | 0.150 | 0.999 | -0.018 | 0.973 | -0.168 | 0.816 |
| PE31 | 0.291 | 0.999 | -0.110 | 0.879 | -0.401 | 0.816 |
| PPE60 | 0.089 | 0.999 | 0.110 | 0.879 | 0.021 | 0.967 |
| Rv3479 | 0.070 | 0.999 | 0.061 | 0.885 | -0.009 | 0.981 |
| Rv3480c | -0.058 | 0.999 | 0.083 | 0.886 | 0.141 | 0.817 |
| Rv3481c | -0.120 | NA | 0.001 | 0.997 | 0.121 | 0.845 |
| Rv3482c | -0.181 | 0.999 | -0.200 | 0.832 | -0.019 | 0.972 |
| Rv3483c | -0.131 | 0.999 | 0.182 | 0.875 | 0.314 | 0.816 |
| Rv3484 | -0.261 | 0.999 | -0.484 | 0.016 | -0.223 | 0.816 |
| Rv3485c | 0.080 | 0.999 | 0.077 | 0.879 | -0.003 | 0.996 |
| Rv3486 | -0.240 | 0.999 | -0.303 | 0.820 | -0.063 | 0.918 |
| lipF | 0.230 | 0.999 | 0.107 | 0.895 | -0.123 | 0.862 |
| Rv3488 | -0.017 | 0.999 | 0.416 | 0.820 | 0.433 | 0.816 |
| Rv3489 | 0.067 | 0.999 | 0.169 | 0.859 | 0.101 | 0.884 |
| otsA | 0.077 | 0.999 | 0.106 | 0.848 | 0.029 | 0.941 |
| Rv3491 | -0.010 | 0.999 | -0.353 | 0.820 | -0.343 | 0.816 |
| Rv3492c | 0.088 | 0.999 | -0.026 | 0.971 | -0.114 | 0.864 |
| Rv3493c | 0.039 | 0.999 | 0.203 | 0.826 | 0.164 | 0.816 |
| mce4F | -0.068 | 0.999 | 0.146 | 0.883 | 0.214 | 0.824 |
| lprN | 0.065 | 0.999 | -0.149 | 0.879 | -0.214 | 0.817 |
| mce4D | 0.111 | 0.999 | 0.040 | 0.948 | -0.071 | 0.891 |
| mce4C | 0.011 | 0.999 | -0.070 | 0.927 | -0.081 | 0.896 |
| mce4B | 0.121 | 0.999 | -0.093 | 0.895 | -0.214 | 0.816 |
| mce4A | 0.097 | 0.999 | -0.107 | 0.894 | -0.204 | 0.816 |
| yrbE4B | 0.116 | 0.999 | -0.210 | 0.834 | -0.326 | 0.816 |
| yrbE4A | -0.080 | 0.999 | 0.096 | 0.893 | 0.175 | 0.817 |
| Rv3502c | -0.036 | 0.999 | -0.077 | 0.918 | -0.041 | 0.942 |
| fdxD | -0.123 | NA | -0.599 | 0.848 | -0.477 | 0.835 |
| fadE26 | -0.221 | NA | -0.226 | 0.848 | -0.005 | 0.996 |
| fadE27 | -0.088 | NA | 0.154 | 0.879 | 0.242 | 0.816 |
| fadD17 | -0.089 | 0.999 | 0.097 | 0.912 | 0.186 | 0.822 |
| PE_PGRS53 | -0.307 | 0.999 | -0.175 | 0.927 | 0.131 | 0.932 |
| PE_PGRS54 | 0.039 | 0.999 | -0.191 | 0.922 | -0.229 | 0.879 |
| ilvX | 0.083 | 0.999 | 0.329 | 0.826 | 0.246 | 0.816 |
| Rv3510c | 0.058 | 0.999 | 0.122 | 0.848 | 0.064 | 0.879 |
| PE_PGRS55 | -0.027 | 0.999 | -0.036 | 0.978 | -0.009 | 0.996 |
| PE_PGRS56 | -0.011 | NA | -0.746 | 0.848 | -0.735 | 0.819 |
| fadD18 | -0.168 | NA | -0.081 | 0.962 | 0.087 | 0.950 |
| PE_PGRS57 | -0.045 | 0.999 | -0.098 | 0.958 | -0.053 | 0.969 |
| fadD19 | -0.122 | 0.999 | -0.252 | 0.831 | -0.130 | 0.855 |
| echA19 | -0.049 | 0.999 | 0.106 | 0.879 | 0.155 | 0.822 |
| Rv3517 | 0.077 | NA | -0.008 | 0.994 | -0.085 | 0.936 |
| cyp142 | -0.036 | 0.999 | 0.132 | 0.895 | 0.168 | 0.854 |
| Rv3519 | -0.052 | 0.999 | 0.149 | 0.848 | 0.201 | 0.816 |
| Rv3520c | 0.032 | 0.999 | 0.169 | 0.834 | 0.137 | 0.822 |
| Rv3521 | 0.069 | 0.999 | -0.132 | 0.858 | -0.201 | 0.816 |
| ltp4 | -0.052 | 0.999 | 0.043 | 0.942 | 0.095 | 0.855 |
| ltp3 | -0.146 | 0.999 | 0.097 | 0.920 | 0.243 | 0.816 |
| Rv3524 | -0.226 | 0.999 | 0.025 | 0.973 | 0.251 | 0.816 |
| Rv3525c | 0.177 | NA | -0.066 | 0.962 | -0.244 | 0.852 |
| kshA | 0.039 | 0.999 | -0.120 | 0.922 | -0.159 | 0.875 |
| Rv3527 | 0.118 | 0.999 | -0.135 | 0.879 | -0.252 | 0.816 |
| Rv3528c | 0.160 | 0.999 | -0.382 | 0.848 | -0.543 | 0.816 |
| Rv3529c | -0.178 | 0.999 | -0.432 | 0.832 | -0.255 | 0.854 |
| Rv3530c | 0.094 | 0.999 | -0.066 | 0.962 | -0.160 | 0.891 |
| Rv3531c | -0.105 | 0.999 | -0.207 | 0.848 | -0.102 | 0.888 |
| PPE61 | -0.092 | 0.999 | -0.082 | 0.927 | 0.009 | 0.989 |
| PPE62 | -0.104 | 0.999 | -0.563 | 0.848 | -0.460 | 0.837 |
| hsaF | -0.117 | 0.999 | -0.068 | 0.912 | 0.049 | 0.920 |
| hsaG | -0.060 | 0.999 | -0.155 | 0.848 | -0.095 | 0.868 |
| hsaE | -0.172 | 0.999 | -0.171 | 0.906 | 0.001 | 0.999 |
| kstD | -0.074 | 0.999 | -0.264 | 0.832 | -0.190 | 0.831 |
| Rv3538 | 0.003 | NA | -0.310 | 0.848 | -0.312 | 0.817 |
| PPE63 | -0.239 | NA | -0.242 | 0.879 | -0.003 | 0.998 |
| ltp2 | -0.052 | 0.999 | -0.219 | 0.826 | -0.167 | 0.816 |
| Rv3541c | 0.075 | NA | -0.267 | 0.848 | -0.342 | 0.816 |
| Rv3542c | -0.020 | 0.999 | -0.103 | 0.901 | -0.084 | 0.900 |
| fadE29 | -0.084 | 0.999 | -0.271 | 0.716 | -0.187 | 0.816 |
| fadE28 | -0.059 | 0.999 | -0.018 | 0.974 | 0.041 | 0.942 |
| cyp125 | -0.069 | 0.999 | -0.090 | 0.879 | -0.022 | 0.958 |
| fadA5 | -0.135 | 0.999 | 0.190 | 0.832 | 0.326 | 0.816 |
| ddn | 0.001 | 0.999 | -0.070 | 0.962 | -0.071 | 0.954 |
| Rv3548c | -0.132 | 0.999 | -0.074 | 0.922 | 0.058 | 0.924 |
| Rv3549c | -0.254 | 0.999 | -0.250 | 0.873 | 0.004 | 0.997 |
| echA20 | -0.184 | 0.999 | 0.139 | 0.879 | 0.323 | 0.816 |
| Rv3551 | -0.289 | 0.999 | -0.180 | 0.837 | 0.110 | 0.855 |
| Rv3552 | -0.198 | 0.999 | -0.193 | 0.832 | 0.005 | 0.994 |
| Rv3553 | -0.257 | 0.999 | 0.118 | 0.895 | 0.375 | 0.816 |
| fdxB | -0.082 | 0.999 | 0.057 | 0.927 | 0.139 | 0.821 |
| Rv3555c | -0.127 | 0.999 | -0.031 | 0.970 | 0.096 | 0.892 |
| fadA6 | -0.152 | 0.999 | 0.015 | 0.974 | 0.167 | 0.816 |
| Rv3557c | -0.157 | 0.999 | -0.088 | 0.927 | 0.069 | 0.930 |
| PPE64 | -0.031 | 0.999 | -0.348 | 0.859 | -0.318 | 0.837 |
| Rv3559c | -0.151 | 0.999 | 0.040 | 0.971 | 0.191 | 0.854 |
| fadE30 | -0.098 | 0.999 | 0.224 | 0.826 | 0.321 | 0.816 |
| fadD3 | -0.111 | 0.999 | 0.019 | 0.978 | 0.130 | 0.868 |
| fadE31 | -0.113 | 0.999 | -0.042 | 0.962 | 0.071 | 0.926 |
| fadE32 | -0.253 | 0.999 | 0.113 | 0.848 | 0.366 | 0.816 |
| fadE33 | -0.219 | 0.999 | -0.400 | 0.831 | -0.181 | 0.868 |
| aspB | -0.245 | NA | 0.167 | 0.859 | 0.412 | 0.816 |
| Rv3566A | -0.107 | 0.999 | -0.277 | 0.831 | -0.171 | 0.838 |
| nat | -0.002 | 0.999 | -0.232 | 0.826 | -0.231 | 0.816 |
| hsaB | -0.009 | 0.999 | -0.004 | 0.995 | 0.006 | 0.996 |
| hsaC | -0.133 | 0.999 | 0.124 | 0.912 | 0.257 | 0.817 |
| hsaD | 0.043 | 0.999 | -0.201 | 0.858 | -0.245 | 0.816 |
| hsaA | -0.093 | 0.999 | 0.016 | 0.994 | 0.109 | 0.962 |
| kshB | 0.059 | 0.999 | 0.211 | 0.856 | 0.153 | 0.862 |
| Rv3572 | -0.010 | 0.999 | 0.222 | 0.879 | 0.232 | 0.852 |
| fadE34 | -0.188 | NA | -0.059 | 0.961 | 0.129 | 0.894 |
| kstR | -0.150 | 0.999 | -0.031 | 0.971 | 0.120 | 0.868 |
| Rv3575c | 0.083 | 0.999 | 0.277 | 0.820 | 0.193 | 0.816 |
| lppH | 0.050 | 0.999 | -0.247 | 0.728 | -0.297 | 0.816 |
| Rv3577 | 0.104 | 0.999 | 0.218 | 0.846 | 0.114 | 0.870 |
| arsB2 | -0.002 | 0.999 | 0.735 | 0.660 | 0.736 | 0.816 |
| Rv3579c | -0.055 | 0.999 | -0.109 | 0.831 | -0.054 | 0.862 |
| cysS1 | -0.090 | 0.999 | -0.141 | 0.854 | -0.051 | 0.925 |
| ispF | -0.149 | 0.999 | 0.156 | 0.903 | 0.306 | 0.817 |
| ispD | -0.080 | 0.999 | -0.028 | 0.970 | 0.052 | 0.936 |
| Rv3583c | -0.002 | 0.999 | -0.095 | 0.951 | -0.092 | 0.942 |
| lpqE | 0.057 | 0.999 | -0.108 | 0.879 | -0.165 | 0.816 |
| radA | 0.005 | 0.999 | -0.025 | 0.951 | -0.030 | 0.932 |
| Rv3586 | -0.016 | 0.999 | -0.325 | 0.826 | -0.310 | 0.816 |
| Rv3587c | 0.105 | 0.999 | 0.202 | 0.848 | 0.097 | 0.892 |
| canB | -0.007 | 0.999 | 0.192 | 0.848 | 0.200 | 0.816 |
| mutY | 0.042 | 0.999 | 0.474 | 0.820 | 0.432 | 0.816 |
| PE_PGRS58 | -0.119 | 0.999 | -0.238 | 0.895 | -0.119 | 0.936 |
| Rv3591c | 0.080 | NA | 0.154 | 0.906 | 0.073 | 0.942 |
| Rv3592 | -0.115 | 0.999 | -0.183 | 0.848 | -0.068 | 0.917 |
| lpqF | 0.082 | 0.999 | -0.107 | 0.879 | -0.189 | 0.816 |

| | | | | | | |
|---|---|---|---|---|---|---|
| iniC | -0.230 | 0.999 | -0.126 | 0.848 | 0.104 | 0.832 |
| lpqJ | 0.034 | 0.999 | -0.254 | 0.886 | -0.289 | 0.859 |
| Rv0345 | -0.089 | 0.999 | 0.033 | 0.970 | 0.122 | 0.868 |
| ansP2 | 0.293 | 0.999 | 0.103 | 0.879 | -0.190 | 0.816 |
| Rv0347 | 0.047 | 0.999 | -0.100 | 0.958 | -0.147 | 0.922 |
| Rv0348 | -0.024 | NA | -0.259 | 0.879 | -0.236 | 0.856 |
| Rv0349 | -0.097 | 0.999 | -0.444 | 0.728 | -0.347 | 0.816 |
| dnaK | -0.152 | 0.999 | -0.298 | 0.829 | -0.146 | 0.857 |
| grpE | -0.257 | 0.999 | -0.467 | 0.895 | -0.210 | 0.942 |
| dnaJ1 | -0.232 | 0.999 | -0.051 | 0.948 | 0.181 | 0.819 |
| hspR | -0.166 | 0.999 | -0.415 | 0.848 | -0.249 | 0.868 |
| PPE7 | 0.050 | NA | -0.564 | 0.772 | -0.614 | 0.816 |
| PPE8 | 0.102 | 0.999 | -0.351 | 0.879 | -0.453 | 0.831 |
| Rv0356c | -0.071 | NA | 0.335 | 0.820 | 0.406 | 0.816 |
| purA | 0.089 | 0.999 | -0.134 | 0.877 | -0.223 | 0.816 |
| Rv0358 | -0.051 | NA | -0.085 | 0.951 | -0.034 | 0.977 |
| Rv0359 | -0.045 | NA | 0.012 | 0.990 | 0.057 | 0.956 |
| Rv0360c | 0.456 | 0.999 | 0.311 | 0.831 | -0.146 | 0.868 |
| Rv0361 | -0.127 | 0.999 | -0.325 | 0.832 | -0.198 | 0.846 |
| mgtE | -0.099 | 0.999 | -0.097 | 0.832 | 0.002 | 0.994 |
| fba | 0.004 | 0.999 | -0.200 | 0.831 | -0.204 | 0.816 |
| Rv0364 | 0.158 | 0.999 | 0.343 | 0.820 | 0.185 | 0.819 |
| Rv0365c | -0.077 | 0.999 | -0.047 | 0.935 | 0.029 | 0.947 |
| Rv0366c | -0.059 | 0.999 | 0.107 | 0.918 | 0.165 | 0.852 |
| Rv0367c | -0.122 | 0.999 | 0.020 | 0.973 | 0.141 | 0.819 |
| Rv0368c | 0.014 | NA | 0.290 | 0.879 | 0.276 | 0.866 |
| Rv0369c | 0.110 | NA | -0.029 | 0.973 | -0.140 | 0.868 |
| Rv0370c | -0.079 | NA | -0.301 | 0.879 | -0.222 | 0.884 |
| Rv0371c | 0.269 | NA | 0.106 | 0.971 | -0.163 | 0.946 |
| Rv0372c | -0.013 | NA | 0.140 | 0.931 | 0.153 | 0.904 |
| Rv0373c | -0.010 | NA | -0.029 | 0.969 | -0.019 | 0.974 |
| Rv0374c | 0.005 | NA | -0.114 | 0.948 | -0.120 | 0.932 |
| Rv0375c | -0.064 | NA | 0.034 | 0.975 | 0.098 | 0.930 |
| Rv0376c | -0.084 | NA | 0.263 | 0.834 | 0.347 | 0.816 |
| Rv0377 | -0.085 | 0.999 | 0.096 | 0.918 | 0.181 | 0.832 |
| Rv0378 | 0.062 | NA | -1.459 | 0.826 | -1.522 | 0.816 |
| secE2 | -0.178 | NA | -1.268 | NA | -1.090 | NA |
| Rv0380c | 0.069 | 0.999 | 0.080 | 0.930 | 0.011 | 0.988 |
| Rv0381c | -0.062 | 0.999 | 0.265 | 0.859 | 0.327 | 0.817 |
| pyrE | -0.086 | 0.999 | 0.355 | 0.826 | 0.441 | 0.816 |
| Rv0383c | 0.033 | 0.999 | -0.025 | 0.974 | -0.057 | 0.942 |
| clpB | 0.026 | 0.999 | 0.078 | 0.919 | 0.051 | 0.932 |
| Rv0385 | -0.016 | 0.999 | 0.108 | 0.883 | 0.124 | 0.852 |
| Rv0386 | -0.066 | 0.999 | 0.040 | 0.958 | 0.105 | 0.866 |
| purT | -0.114 | 0.999 | 0.172 | 0.879 | 0.286 | 0.816 |
| Rv0390 | 0.061 | 0.999 | 0.259 | 0.785 | 0.197 | 0.816 |
| metZ | 0.144 | 0.999 | 0.132 | 0.832 | -0.012 | 0.976 |
| ndhA | -0.001 | 0.999 | -0.016 | 0.962 | -0.015 | 0.955 |
| Rv0393 | -0.322 | NA | -1.360 | NA | -1.038 | NA |
| Rv0394c | -0.199 | 0.999 | 0.050 | 0.963 | 0.248 | 0.819 |
| Rv0395 | -0.220 | NA | -0.067 | 0.974 | 0.153 | 0.939 |
| Rv0396 | 0.184 | NA | -0.807 | 0.848 | -0.991 | 0.816 |
| Rv0397 | 0.053 | NA | -1.239 | NA | -1.292 | NA |
| Rv0397A | -0.240 | 0.999 | 0.391 | 0.831 | 0.631 | 0.816 |
| Rv0398c | -0.023 | 0.999 | 0.178 | 0.872 | 0.201 | 0.831 |
| lpqK | 0.018 | 0.999 | -0.152 | 0.927 | -0.170 | 0.897 |
| fadE7 | -0.067 | 0.999 | 0.305 | 0.831 | 0.372 | 0.816 |
| Rv0401 | -0.123 | NA | 0.522 | 0.770 | 0.645 | 0.816 |
| mmpL1 | 0.198 | 0.999 | -0.242 | 0.898 | -0.440 | 0.819 |
| mmpS1 | -0.042 | 0.999 | -0.450 | 0.832 | -0.408 | 0.816 |
| fadD30 | 0.194 | 0.999 | -0.318 | 0.877 | -0.513 | 0.816 |
| pks6 | 0.040 | 0.999 | -0.330 | 0.859 | -0.370 | 0.819 |
| Rv0406c | 0.040 | 0.999 | 0.329 | 0.825 | 0.289 | 0.816 |
| fgd1 | 0.035 | 0.999 | 0.032 | 0.954 | -0.003 | 0.996 |
| pta | 0.142 | 0.999 | -0.175 | 0.826 | -0.317 | 0.816 |
| ackA | 0.083 | 0.999 | 0.290 | 0.820 | 0.207 | 0.816 |
| pknG | -0.192 | 0.999 | -0.283 | 0.527 | -0.091 | 0.838 |
| glnH | -0.107 | 0.999 | -0.250 | 0.439 | -0.143 | 0.816 |
| Rv0412c | -0.192 | 0.999 | -0.151 | 0.831 | 0.041 | 0.918 |
| mutT3 | -0.078 | 0.999 | 0.083 | 0.922 | 0.162 | 0.835 |
| thiE | -0.140 | NA | 0.352 | 0.826 | 0.492 | 0.816 |
| thiO | -0.066 | 0.999 | 0.492 | 0.820 | 0.558 | 0.816 |
| thiS | -0.012 | NA | -0.608 | 0.824 | -0.596 | 0.816 |
| thiG | 0.000 | 0.999 | -0.106 | 0.903 | -0.106 | 0.884 |
| lpqL | 0.007 | 0.999 | -0.111 | 0.843 | -0.118 | 0.816 |
| lpqM | -0.004 | 0.999 | -0.013 | 0.970 | -0.009 | 0.976 |
| Rv0420c | -0.125 | 0.999 | 0.055 | 0.958 | 0.180 | 0.839 |
| Rv0421c | -0.033 | 0.999 | 0.439 | 0.826 | 0.472 | 0.816 |
| thiD | -0.012 | 0.999 | 0.355 | 0.832 | 0.367 | 0.816 |
| thiC | 0.041 | 0.999 | -0.049 | 0.929 | -0.089 | 0.854 |
| Rv0424c | 0.069 | 0.999 | 0.086 | 0.879 | 0.016 | 0.967 |
| ctpH | 0.025 | 0.999 | 0.165 | 0.879 | 0.140 | 0.865 |
| Rv0426c | -0.125 | 0.999 | -0.093 | 0.958 | 0.033 | 0.983 |
| xthA | 0.030 | 0.999 | 0.072 | 0.915 | 0.041 | 0.937 |
| Rv0428c | 0.128 | 0.999 | 0.292 | 0.820 | 0.164 | 0.820 |
| def | 0.093 | 0.999 | 0.113 | 0.879 | 0.020 | 0.968 |
| Rv0430 | 0.051 | 0.999 | 0.311 | 0.831 | 0.260 | 0.816 |
| Rv0431 | 0.149 | 0.999 | 0.002 | 0.996 | -0.147 | 0.837 |
| sodC | 0.132 | 0.999 | 0.059 | 0.895 | -0.072 | 0.856 |
| Rv0433 | 0.104 | 0.999 | -0.093 | 0.879 | -0.197 | 0.816 |
| Rv0434 | 0.044 | 0.999 | 0.072 | 0.914 | 0.027 | 0.956 |
| Rv0435c | 0.032 | 0.999 | 0.123 | 0.879 | 0.091 | 0.891 |
| pssA | 0.111 | 0.999 | 0.197 | 0.825 | 0.086 | 0.852 |
| psd | 0.049 | 0.999 | 0.140 | 0.916 | 0.090 | 0.932 |
| moeA2 | 0.047 | 0.999 | 0.429 | 0.831 | 0.382 | 0.816 |
| Rv0439c | 0.146 | 0.999 | 0.037 | 0.936 | -0.109 | 0.817 |
| groEL2 | -0.177 | 0.999 | 0.049 | 0.942 | 0.225 | 0.816 |
| Rv0441c | 0.014 | 0.999 | 0.647 | 0.826 | 0.633 | 0.816 |
| PPE10 | -0.035 | 0.999 | -0.213 | 0.848 | -0.177 | 0.835 |
| Rv0443 | -0.047 | 0.999 | 0.140 | 0.846 | 0.187 | 0.816 |
| rskA | -0.095 | 0.999 | 0.159 | 0.859 | 0.254 | 0.816 |
| sigK | 0.005 | 0.999 | -0.081 | 0.931 | -0.086 | 0.909 |
| Rv0446c | 0.064 | 0.999 | 0.201 | 0.854 | 0.137 | 0.866 |
| ufaA1 | -0.010 | 0.999 | 0.094 | 0.922 | 0.104 | 0.891 |
| Rv0448c | -0.066 | NA | -0.269 | 0.831 | -0.203 | 0.817 |
| Rv0449c | -0.126 | 0.999 | -0.108 | 0.832 | 0.017 | 0.954 |
| mmpL4 | 0.048 | 0.999 | -0.430 | 0.695 | -0.478 | 0.816 |
| mmpS4 | 0.065 | 0.999 | -0.475 | 0.843 | -0.540 | 0.816 |
| Rv0452 | 0.000 | NA | -0.184 | 0.879 | -0.184 | 0.862 |
| PPE11 | -0.242 | NA | -0.283 | 0.886 | -0.042 | 0.981 |
| Rv0455c | 0.091 | 0.999 | -0.530 | 0.804 | -0.621 | 0.816 |
| mazF1 | 0.003 | NA | -0.047 | 0.973 | -0.050 | 0.969 |
| mazE1 | 0.139 | NA | -0.572 | 0.832 | -0.710 | 0.816 |
| echA2 | 0.093 | NA | -0.122 | 0.922 | -0.214 | 0.838 |
| Rv0457c | -0.196 | 0.999 | -0.310 | 0.826 | -0.114 | 0.867 |
| Rv0458 | 0.179 | 0.999 | 0.368 | 0.659 | 0.189 | 0.816 |
| Rv0459 | 0.107 | NA | 0.303 | 0.831 | 0.197 | 0.833 |
| Rv0460 | -0.054 | 0.999 | 0.398 | 0.826 | 0.452 | 0.816 |
| Rv0461 | 0.008 | 0.999 | 0.508 | 0.826 | 0.500 | 0.816 |
| lpdC | -0.011 | 0.999 | -0.170 | 0.848 | -0.160 | 0.830 |
| Rv0463 | -0.050 | NA | -0.244 | 0.848 | -0.194 | 0.838 |
| Rv0464c | 0.068 | 0.999 | -0.121 | 0.907 | -0.189 | 0.838 |
| Rv0465c | -0.028 | 0.999 | -0.090 | 0.947 | -0.061 | 0.951 |
| Rv0466 | -0.006 | 0.999 | 0.216 | 0.848 | 0.222 | 0.817 |
| icl1 | -0.467 | 0.100 | -0.531 | 0.011 | -0.065 | 0.891 |
| fadB2 | 0.007 | 0.999 | -0.248 | 0.659 | -0.255 | 0.816 |
| umaA | 0.090 | 0.999 | -0.223 | 0.886 | -0.313 | 0.836 |
| Rv0470A | -0.011 | 0.999 | 0.163 | 0.831 | 0.174 | 0.816 |
| pcaA | 0.107 | 0.999 | -0.301 | 0.826 | -0.409 | 0.816 |
| Rv0471c | 0.145 | 0.999 | -0.048 | 0.924 | -0.193 | 0.816 |
| Rv0472c | 0.004 | 0.999 | -0.264 | 0.832 | -0.268 | 0.816 |
| Rv0473 | -0.036 | 0.999 | -0.169 | 0.832 | -0.134 | 0.819 |
| Rv0474 | 0.068 | 0.999 | -0.132 | 0.922 | -0.200 | 0.865 |
| hbhA | -0.066 | 0.999 | 0.359 | 0.826 | 0.425 | 0.816 |
| Rv0476 | 0.036 | 0.999 | 0.099 | 0.951 | 0.062 | 0.960 |
| Rv0477 | -0.164 | 0.999 | 0.441 | 0.853 | 0.605 | 0.816 |
| deoC | 0.060 | 0.999 | 0.170 | 0.831 | 0.110 | 0.833 |
| Rv0479c | 0.104 | 0.999 | -0.215 | 0.848 | -0.320 | 0.816 |
| Rv0480c | 0.019 | 0.999 | 0.064 | 0.886 | 0.045 | 0.904 |
| Rv0481c | 0.107 | 0.999 | 0.057 | 0.962 | -0.050 | 0.955 |
| murB | -0.055 | 0.999 | 0.253 | 0.826 | 0.308 | 0.816 |
| lprQ | -0.384 | 0.999 | -0.037 | 0.975 | 0.348 | 0.819 |
| Rv0484c | 0.059 | 0.999 | 0.348 | 0.826 | 0.289 | 0.816 |
| Rv0485 | -0.191 | 0.999 | -0.128 | 0.832 | 0.063 | 0.868 |
| mshA | -0.068 | 0.999 | 0.106 | 0.922 | 0.174 | 0.855 |
| Rv0487 | -0.014 | 0.999 | -0.069 | 0.922 | -0.055 | 0.922 |
| Rv0488 | -0.074 | 0.999 | -0.301 | 0.843 | -0.226 | 0.836 |
| gpm1 | 0.024 | 0.999 | 0.142 | 0.895 | 0.118 | 0.896 |
| senX3 | -0.029 | 0.999 | -0.041 | 0.970 | -0.012 | 0.989 |
| regX3 | 0.002 | 0.999 | -0.070 | 0.879 | -0.072 | 0.842 |
| Rv0492A | 0.242 | NA | 0.552 | 0.831 | 0.309 | 0.852 |
| Rv0492c | -0.016 | 0.999 | 0.380 | 0.394 | 0.396 | 0.816 |
| Rv0493c | -0.103 | NA | -0.011 | 0.983 | 0.092 | 0.868 |
| Rv0494 | -0.575 | NA | -0.264 | 0.928 | 0.311 | 0.896 |
| Rv0495c | 0.199 | 0.999 | -0.062 | 0.940 | -0.261 | 0.816 |
| Rv0496 | -0.034 | 0.999 | 0.011 | 0.975 | 0.046 | 0.909 |
| Rv0497 | 0.045 | 0.999 | 0.120 | 0.855 | 0.076 | 0.869 |
| Rv0498 | 0.081 | 0.999 | 0.283 | 0.826 | 0.202 | 0.817 |
| Rv0499 | 0.091 | 0.999 | 0.622 | 0.510 | 0.531 | 0.816 |
| proC | -0.034 | 0.999 | 0.118 | 0.922 | 0.152 | 0.879 |
| Rv0500A | -0.011 | 0.999 | 0.611 | 0.826 | 0.622 | 0.816 |
| galE2 | 0.001 | 0.999 | 0.025 | 0.941 | 0.024 | 0.930 |
| Rv0502 | -0.011 | 0.999 | 0.042 | 0.962 | 0.053 | 0.940 |
| cmaA2 | 0.125 | 0.999 | -0.254 | 0.846 | -0.379 | 0.816 |
| Rv0504c | 0.239 | 0.999 | -0.051 | 0.958 | -0.289 | 0.816 |
| serB1 | 0.019 | 0.999 | 0.123 | 0.879 | 0.104 | 0.868 |
| mmpS2 | 0.080 | 0.999 | -0.034 | 0.952 | -0.114 | 0.835 |
| mmpL2 | 0.165 | 0.999 | -0.223 | 0.850 | -0.388 | 0.816 |
| Rv0508 | 0.118 | NA | -0.436 | 0.829 | -0.555 | 0.816 |
| hemA | -0.052 | 0.999 | 0.014 | 0.978 | 0.065 | 0.904 |
| hemC | 0.068 | 0.999 | 0.134 | 0.893 | 0.066 | 0.936 |
| hemD | -0.056 | 0.999 | 0.240 | 0.863 | 0.296 | 0.817 |
| hemB | -0.044 | 0.999 | 0.139 | 0.877 | 0.184 | 0.817 |
| Rv0513 | -0.002 | 0.999 | 0.334 | 0.831 | 0.335 | 0.816 |
| Rv0514 | 0.137 | 0.999 | 0.211 | 0.879 | 0.074 | 0.942 |

| | | | | | | |
|---|---|---|---|---|---|---|
| narG | -0.002 | 0.999 | -0.204 | 0.848 | -0.201 | 0.817 |
| narH | -0.004 | 0.999 | -0.268 | 0.659 | -0.264 | 0.816 |
| narJ | -0.161 | 0.999 | 0.178 | 0.909 | 0.339 | 0.819 |
| narI | -0.025 | 0.999 | 0.104 | 0.935 | 0.128 | 0.897 |
| typA | -0.002 | 0.999 | -0.032 | 0.956 | -0.030 | 0.946 |
| lpqW | -0.060 | 0.999 | -0.151 | 0.826 | -0.091 | 0.822 |
| Rv1167c | 0.050 | 0.999 | -0.018 | 0.970 | -0.068 | 0.870 |
| PPE17 | -0.127 | 0.999 | -0.257 | 0.728 | -0.130 | 0.817 |
| lipX | 0.183 | 0.999 | -0.481 | 0.820 | -0.664 | 0.816 |
| mshB | -0.190 | 0.999 | -0.176 | 0.879 | 0.014 | 0.988 |
| Rv1171 | -0.025 | 0.999 | 0.391 | 0.826 | 0.417 | 0.816 |
| PE12 | 0.175 | 0.999 | 0.032 | 0.958 | -0.143 | 0.817 |
| fbiC | 0.045 | 0.999 | 0.116 | 0.890 | 0.071 | 0.920 |
| TB8.4 | -0.011 | 0.999 | 0.217 | 0.826 | 0.228 | 0.816 |
| fadH | 0.057 | 0.999 | 0.146 | 0.832 | 0.090 | 0.852 |
| Rv1176c | 0.149 | 0.999 | 0.080 | 0.946 | -0.069 | 0.940 |
| fdxC | -0.100 | 0.999 | -0.231 | 0.865 | -0.132 | 0.895 |
| Rv1178 | 0.006 | 0.999 | 0.208 | 0.831 | 0.202 | 0.816 |
| Rv1179c | 0.086 | 0.999 | 0.194 | 0.820 | 0.108 | 0.819 |
| pks3 | 0.095 | 0.999 | 0.152 | 0.853 | 0.057 | 0.921 |
| pks4 | 0.076 | 0.999 | 0.108 | 0.879 | 0.032 | 0.950 |
| papA3 | 0.102 | 0.999 | -0.007 | 0.990 | -0.109 | 0.867 |
| mmpL10 | 0.064 | 0.999 | 0.041 | 0.935 | -0.023 | 0.954 |
| Rv1184c | 0.042 | 0.999 | 0.091 | 0.922 | 0.049 | 0.942 |
| fadD21 | 0.056 | 0.999 | -0.239 | 0.831 | -0.296 | 0.816 |
| Rv1186c | -0.196 | 0.999 | 0.104 | 0.895 | 0.301 | 0.816 |
| rocA | -0.060 | 0.999 | -0.090 | 0.879 | -0.030 | 0.942 |
| Rv1188 | -0.007 | 0.999 | 0.105 | 0.879 | 0.111 | 0.840 |
| sigI | -0.363 | NA | 0.061 | 0.962 | 0.424 | 0.816 |
| Rv1190 | -0.179 | NA | -0.187 | 0.879 | -0.008 | 0.994 |
| Rv1191 | -0.283 | 0.999 | 0.493 | 0.831 | 0.776 | 0.816 |
| Rv1192 | -0.225 | 0.999 | -0.007 | 0.994 | 0.218 | 0.837 |
| fadD36 | -0.204 | 0.999 | 0.219 | 0.879 | 0.423 | 0.816 |
| Rv1194c | -0.102 | 0.999 | -0.209 | 0.826 | -0.107 | 0.836 |
| PE13 | 0.227 | 0.999 | -0.292 | 0.883 | -0.518 | 0.816 |
| PPE18 | -0.111 | 0.999 | -0.218 | 0.831 | -0.107 | 0.866 |
| esxK | -0.005 | 0.999 | -0.406 | 0.885 | -0.400 | 0.866 |
| esxL | -0.047 | 0.999 | 0.108 | 0.931 | 0.155 | 0.879 |
| Rv1199c | -0.152 | 0.999 | -0.417 | 0.848 | -0.265 | 0.862 |
| Rv1200 | 0.014 | 0.999 | 0.129 | 0.859 | 0.115 | 0.842 |
| dapD | 0.029 | 0.999 | 0.155 | 0.854 | 0.125 | 0.846 |
| dapE | 0.077 | 0.999 | -0.032 | 0.970 | -0.109 | 0.870 |
| Rv1203c | -0.196 | NA | 0.097 | 0.922 | 0.292 | 0.816 |
| Rv1204c | -0.004 | 0.999 | 0.429 | 0.820 | 0.433 | 0.816 |
| Rv1205 | 0.124 | 0.999 | -0.152 | 0.941 | -0.275 | 0.868 |
| fadD6 | 0.050 | 0.999 | 0.067 | 0.913 | 0.017 | 0.970 |
| folP2 | 0.073 | 0.999 | 0.290 | 0.669 | 0.216 | 0.816 |
| gpgS | -0.010 | 0.999 | 0.332 | 0.831 | 0.342 | 0.816 |
| Rv1209 | 0.054 | 0.999 | 0.238 | 0.831 | 0.184 | 0.817 |
| tagA | 0.155 | 0.999 | 0.151 | 0.859 | -0.004 | 0.996 |
| Rv1211 | -0.147 | 0.999 | -0.014 | 0.991 | 0.133 | 0.925 |
| glgA | -0.082 | 0.999 | -0.103 | 0.895 | -0.021 | 0.975 |
| glgC | -0.029 | 0.999 | -0.158 | 0.848 | -0.130 | 0.835 |
| PE14 | -0.031 | NA | 0.214 | 0.848 | 0.245 | 0.816 |
| Rv1215c | 0.047 | 0.999 | 0.238 | 0.826 | 0.190 | 0.816 |
| Rv1216c | 0.070 | 0.999 | -0.017 | 0.974 | -0.087 | 0.868 |
| Rv1217c | -0.106 | 0.999 | 0.452 | 0.826 | 0.558 | 0.816 |
| Rv1218c | 0.076 | 0.999 | -0.295 | 0.826 | -0.371 | 0.816 |
| Rv1219c | 0.190 | 0.999 | -0.189 | 0.866 | -0.379 | 0.816 |
| Rv1220c | 0.211 | 0.999 | 0.486 | 0.055 | 0.275 | 0.816 |
| sigE | -0.118 | 0.999 | -0.190 | 0.846 | -0.073 | 0.902 |
| rseA | 0.092 | 0.999 | -0.065 | 0.918 | -0.157 | 0.816 |
| htrA | 0.029 | 0.999 | 0.249 | 0.659 | 0.220 | 0.816 |
| tatB | 0.038 | 0.999 | 0.070 | 0.879 | 0.032 | 0.932 |
| Rv1225c | -0.078 | 0.999 | 0.022 | 0.975 | 0.100 | 0.896 |
| Rv1226c | 0.057 | 0.999 | -0.011 | 0.983 | -0.069 | 0.900 |
| Rv1227c | 0.029 | 0.999 | -0.208 | 0.870 | -0.238 | 0.831 |
| lpqX | -0.155 | NA | -0.290 | 0.826 | -0.135 | 0.846 |
| Rv1228a | -0.470 | NA | -0.759 | 0.831 | -0.289 | 0.887 |
| mrp | 0.024 | 0.999 | 0.234 | 0.701 | 0.211 | 0.816 |
| Rv1230c | 0.157 | 0.999 | 0.060 | 0.920 | -0.096 | 0.852 |
| Rv1231c | 0.112 | 0.999 | -0.123 | 0.922 | -0.235 | 0.837 |
| Rv1232c | -0.086 | 0.999 | 0.339 | 0.846 | 0.425 | 0.816 |
| Rv1233c | -0.078 | 0.999 | 0.037 | 0.961 | 0.115 | 0.860 |
| Rv1234 | 0.003 | 0.999 | 0.031 | 0.962 | 0.028 | 0.958 |
| lpqY | 0.014 | 0.999 | -0.021 | 0.970 | -0.035 | 0.942 |
| sugA | 0.105 | 0.999 | -0.087 | 0.885 | -0.192 | 0.816 |
| sugB | 0.125 | 0.999 | 0.260 | 0.848 | 0.134 | 0.882 |
| sugC | 0.087 | 0.999 | -0.162 | 0.879 | -0.249 | 0.816 |
| corA | 0.046 | 0.999 | -0.213 | 0.826 | -0.258 | 0.816 |
| mdh | -0.087 | 0.999 | 0.111 | 0.879 | 0.198 | 0.816 |
| vapB33 | 0.086 | 0.999 | -0.151 | 0.879 | -0.238 | 0.816 |
| vapC33 | 0.074 | 0.999 | -0.256 | 0.826 | -0.330 | 0.816 |
| PE_PGRS23 | 0.026 | NA | -0.299 | 0.879 | -0.325 | 0.846 |
| lpqZ | -0.041 | 0.999 | 0.638 | 0.820 | 0.679 | 0.816 |
| Rv1245c | -0.046 | 0.999 | 0.321 | 0.361 | 0.367 | 0.816 |
| relE | 0.018 | NA | -0.655 | 0.669 | -0.673 | 0.816 |
| relB | 0.131 | 0.999 | -0.352 | 0.826 | -0.482 | 0.816 |
| Rv1248c | 0.101 | 0.999 | -0.024 | 0.962 | -0.125 | 0.816 |
| Rv1249c | 0.031 | 0.999 | 0.360 | 0.826 | 0.328 | 0.816 |
| Rv1250 | 0.048 | 0.999 | 0.253 | 0.826 | 0.206 | 0.816 |
| Rv1251c | 0.104 | 0.999 | 0.276 | 0.659 | 0.173 | 0.816 |
| lprE | 0.309 | 0.999 | -0.028 | 0.973 | -0.337 | 0.816 |
| deaD | -0.017 | 0.999 | -0.247 | 0.848 | -0.230 | 0.822 |
| Rv1254 | -0.042 | 0.999 | 0.218 | 0.856 | 0.260 | 0.816 |
| Rv1255c | 0.079 | 0.999 | -0.086 | 0.927 | -0.165 | 0.842 |
| cyp130 | 0.094 | 0.999 | -0.100 | 0.885 | -0.194 | 0.816 |
| Rv1257c | 0.152 | 0.999 | 0.229 | 0.854 | 0.078 | 0.930 |
| Rv1258c | 1.573 | 0.000 | 0.397 | 0.831 | -1.176 | 0.125 |
| udgB | 0.245 | NA | 0.429 | 0.826 | 0.184 | 0.860 |
| Rv1260 | 0.157 | 0.999 | 0.070 | 0.916 | -0.087 | 0.868 |
| Rv1261c | -0.020 | 0.999 | 0.043 | 0.958 | 0.062 | 0.922 |
| Rv1262c | 0.014 | 0.999 | -0.401 | 0.758 | -0.415 | 0.816 |
| amiB2 | 0.143 | 0.999 | 0.321 | 0.659 | 0.177 | 0.816 |
| Rv1264 | -0.003 | 0.999 | 0.329 | 0.826 | 0.333 | 0.816 |
| Rv1265 | -0.091 | 0.999 | 0.075 | 0.922 | 0.166 | 0.819 |
| pknH | -0.133 | 0.999 | -0.392 | 0.826 | -0.259 | 0.819 |
| embR | -0.048 | 0.999 | -0.014 | 0.983 | 0.034 | 0.958 |
| Rv1268c | -0.057 | NA | -0.959 | NA | -0.902 | NA |
| Rv1269c | 0.096 | NA | 0.083 | 0.948 | -0.013 | 0.989 |
| lprA | -0.005 | 0.999 | -0.405 | 0.716 | -0.400 | 0.816 |
| Rv1271c | 0.012 | 0.999 | -0.472 | 0.869 | -0.485 | 0.837 |
| Rv1272c | 0.101 | 0.999 | -0.009 | 0.975 | -0.110 | 0.816 |
| Rv1273c | 0.124 | 0.999 | -0.155 | 0.834 | -0.279 | 0.816 |
| lprB | -0.065 | 0.999 | -0.020 | 0.981 | 0.045 | 0.958 |
| lprC | 0.019 | 0.999 | 0.122 | 0.914 | 0.103 | 0.904 |
| Rv1276c | 0.005 | NA | 0.129 | 0.895 | 0.124 | 0.882 |
| Rv1277 | -0.020 | 0.999 | -0.038 | 0.948 | -0.018 | 0.970 |
| Rv1278 | -0.011 | 0.999 | -0.029 | 0.951 | -0.019 | 0.960 |
| Rv1279 | 0.090 | 0.999 | -0.143 | 0.826 | -0.233 | 0.816 |
| oppA | 0.121 | 0.999 | -0.030 | 0.970 | -0.151 | 0.838 |
| oppD | 0.147 | 0.999 | 0.084 | 0.879 | -0.062 | 0.879 |
| oppC | 0.178 | 0.999 | -0.158 | 0.879 | -0.336 | 0.816 |
| oppB | 0.096 | 0.999 | 0.085 | 0.879 | -0.011 | 0.977 |
| canA | 0.178 | NA | -0.093 | 0.958 | -0.271 | 0.855 |
| cysD | -0.143 | 0.999 | -0.070 | 0.924 | 0.073 | 0.898 |
| Rv1286 | -0.077 | 0.999 | -0.370 | 0.826 | -0.293 | 0.816 |
| Rv1287 | 0.036 | 0.999 | -0.096 | 0.895 | -0.132 | 0.846 |
| Rv1288 | -0.017 | 0.999 | -0.188 | 0.820 | -0.170 | 0.816 |
| Rv1289 | -0.021 | 0.999 | -0.142 | 0.848 | -0.121 | 0.835 |
| Rv1290A | -0.314 | NA | -1.409 | NA | -1.095 | NA |
| Rv1290c | -0.051 | 0.999 | -0.076 | 0.886 | -0.025 | 0.954 |
| Rv1291c | -0.089 | NA | -0.317 | 0.849 | -0.228 | 0.856 |
| argS | -0.048 | 0.999 | 0.017 | 0.970 | 0.066 | 0.865 |
| lysA | -0.059 | 0.999 | 0.017 | 0.975 | 0.076 | 0.896 |
| thrA | -0.006 | 0.999 | -0.115 | 0.859 | -0.109 | 0.837 |
| thrC | 0.088 | 0.999 | 0.002 | 0.996 | -0.086 | 0.891 |
| thrB | -0.058 | 0.999 | 0.069 | 0.895 | 0.127 | 0.819 |
| rho | 0.075 | 0.999 | 0.029 | 0.964 | -0.046 | 0.932 |
| rpmE | 0.049 | 0.999 | 0.027 | 0.985 | -0.022 | 0.988 |
| prfA | 0.123 | 0.999 | -0.064 | 0.933 | -0.187 | 0.817 |
| hemK | 0.014 | 0.999 | 0.149 | 0.879 | 0.135 | 0.864 |
| Rv1301 | 0.024 | 0.999 | -0.062 | 0.948 | -0.087 | 0.913 |
| rfe | -0.023 | 0.999 | 0.323 | 0.826 | 0.345 | 0.816 |
| Rv1303 | -0.083 | 0.999 | -0.097 | 0.931 | -0.014 | 0.988 |
| atpB | 0.035 | 0.999 | -0.257 | 0.826 | -0.292 | 0.816 |
| atpE | -0.212 | 0.999 | 0.205 | 0.895 | 0.418 | 0.816 |
| atpF | 0.039 | 0.999 | -0.285 | 0.767 | -0.323 | 0.816 |
| atpH | 0.007 | 0.999 | -0.224 | 0.826 | -0.231 | 0.816 |
| atpA | 0.021 | 0.999 | -0.059 | 0.879 | -0.080 | 0.819 |
| atpG | 0.012 | 0.999 | -0.325 | 0.571 | -0.337 | 0.816 |
| atpD | -0.008 | 0.999 | -0.144 | 0.848 | -0.137 | 0.819 |
| atpC | -0.034 | 0.999 | -0.355 | 0.826 | -0.321 | 0.816 |
| Rv1312 | -0.058 | 0.999 | -0.166 | 0.829 | -0.109 | 0.831 |
| Rv1313c | -0.223 | 0.999 | -0.008 | 0.993 | 0.215 | 0.835 |
| Rv1314c | 0.147 | 0.999 | 0.044 | 0.951 | -0.103 | 0.868 |
| murA | -0.083 | 0.999 | -0.031 | 0.973 | 0.052 | 0.946 |
| ogt | -0.016 | 0.999 | -0.238 | 0.832 | -0.223 | 0.816 |
| alkA | -0.074 | 0.999 | -0.046 | 0.951 | 0.029 | 0.961 |
| Rv1318c | 0.056 | 0.999 | 0.079 | 0.886 | 0.023 | 0.958 |
| Rv1319c | 0.087 | 0.999 | -0.093 | 0.848 | -0.180 | 0.816 |
| Rv1320c | -0.021 | 0.999 | -0.100 | 0.878 | -0.079 | 0.866 |
| Rv1321 | 0.022 | 0.999 | 0.042 | 0.948 | 0.021 | 0.969 |
| Rv1322 | -0.061 | 0.999 | -0.155 | 0.832 | -0.094 | 0.852 |
| Rv1322A | -0.103 | 0.999 | -0.121 | 0.879 | -0.019 | 0.975 |
| fadA4 | -0.055 | 0.999 | -0.044 | 0.927 | 0.011 | 0.979 |
| Rv1324 | -0.088 | 0.999 | -0.011 | 0.974 | 0.077 | 0.837 |
| PE_PGRS24 | -0.181 | 0.999 | -0.264 | 0.866 | -0.084 | 0.942 |
| glgB | -0.055 | 0.999 | -0.104 | 0.848 | -0.049 | 0.891 |
| glgE | -0.020 | 0.999 | -0.162 | 0.879 | -0.142 | 0.866 |
| glgP | 0.060 | 0.999 | 0.179 | 0.820 | 0.119 | 0.816 |
| dinG | -0.071 | 0.999 | 0.182 | 0.877 | 0.253 | 0.816 |
| pncB1 | 0.105 | 0.999 | -0.012 | 0.977 | -0.116 | 0.831 |
| Rv1331 | -0.014 | 0.999 | -0.017 | 0.983 | -0.003 | 0.997 |
| Rv1332 | 0.013 | 0.999 | -0.074 | 0.931 | -0.087 | 0.898 |

| | | | | | | |
|---|---|---|---|---|---|---|
| Rv1975 | -0.197 | NA | -0.637 | 0.832 | -0.440 | 0.835 |
| Rv1976c | -0.158 | 0.999 | -0.062 | 0.967 | 0.095 | 0.938 |
| Rv1977 | -0.007 | 0.999 | 0.192 | 0.831 | 0.199 | 0.816 |
| Rv1978 | 0.041 | 0.999 | -0.288 | 0.858 | -0.329 | 0.817 |
| Rv1979c | 0.036 | 0.999 | -0.112 | 0.879 | -0.149 | 0.819 |
| mpt64 | 0.123 | 0.999 | -0.153 | 0.935 | -0.276 | 0.865 |
| nrdF1 | 0.026 | 0.999 | -0.054 | 0.933 | -0.080 | 0.879 |
| vapB36 | 0.034 | 0.999 | -0.413 | 0.853 | -0.447 | 0.817 |
| vapC36 | -0.109 | 0.999 | -0.162 | 0.879 | -0.053 | 0.942 |
| PE_PGRS35 | 0.007 | 0.999 | -0.150 | 0.941 | -0.156 | 0.925 |
| RV1984a | -0.283 | 0.999 | 0.235 | 0.848 | 0.518 | 0.816 |
| cfp21 | 0.028 | 0.999 | 0.041 | 0.916 | 0.012 | 0.967 |
| Rv1985c | -0.032 | 0.999 | -0.075 | 0.922 | -0.043 | 0.942 |
| Rv1986 | 0.092 | 0.999 | 0.370 | 0.725 | 0.278 | 0.816 |
| Rv1987 | 0.037 | 0.999 | -0.148 | 0.850 | -0.185 | 0.816 |
| erm(37) | -0.024 | 0.999 | 0.048 | 0.948 | 0.073 | 0.897 |
| Rv1989c | -0.156 | 0.999 | -0.222 | 0.846 | -0.066 | 0.927 |
| Rv1990c | -0.073 | 0.999 | -0.574 | 0.576 | -0.501 | 0.816 |
| mazE6 | -0.010 | NA | -0.557 | 0.714 | -0.547 | 0.816 |
| mazF6 | -0.087 | 0.999 | -0.155 | 0.879 | -0.068 | 0.932 |
| ctpG | -0.359 | 0.096 | -0.356 | 0.052 | 0.004 | 0.993 |
| Rv1993c | -0.394 | NA | -0.639 | 0.826 | -0.245 | 0.866 |
| cmtR | -0.256 | 0.999 | -0.345 | 0.848 | -0.088 | 0.942 |
| Rv1995 | -0.219 | 0.999 | -0.756 | 0.728 | -0.537 | 0.816 |
| Rv1996 | -0.711 | 0.000 | -1.018 | 0.000 | -0.306 | 0.816 |
| ctpF | -0.630 | 0.008 | -1.081 | 0.000 | -0.451 | 0.816 |
| Rv1998c | -0.011 | 0.999 | -0.316 | 0.843 | -0.305 | 0.816 |
| Rv1999c | -0.473 | 0.999 | -0.260 | 0.848 | 0.213 | 0.835 |
| Rv2000 | 0.092 | 0.999 | -0.493 | 0.820 | -0.585 | 0.816 |
| Rv2001 | 0.260 | 0.999 | -0.332 | 0.859 | -0.592 | 0.816 |
| fabG3 | 0.015 | 0.999 | 0.012 | 0.988 | -0.003 | 0.997 |
| Rv2003c | -0.643 | 0.999 | -0.600 | 0.805 | 0.043 | 0.969 |
| Rv2004c | -0.451 | 0.999 | -0.873 | 0.029 | -0.422 | 0.816 |
| Rv2005c | -0.662 | 0.100 | -0.919 | 0.000 | -0.258 | 0.816 |
| otsB1 | -0.237 | 0.999 | -0.635 | 0.002 | -0.398 | 0.816 |
| fdxA | -0.418 | 0.999 | -0.625 | 0.163 | -0.207 | 0.817 |
| Rv2008c | -0.329 | NA | -0.506 | 0.834 | -0.177 | 0.905 |
| vapB15 | 0.122 | 0.999 | -0.364 | 0.831 | -0.486 | 0.816 |
| vapC15 | 0.066 | 0.999 | 0.045 | 0.936 | -0.021 | 0.961 |
| Rv2011c | -0.098 | NA | -0.286 | 0.859 | -0.188 | 0.870 |
| Rv2012 | -0.329 | NA | -0.482 | 0.826 | -0.153 | 0.892 |
| Rv2015c | -0.102 | 0.999 | -0.177 | 0.688 | -0.076 | 0.830 |
| Rv2016 | 0.033 | 0.999 | -0.235 | 0.858 | -0.268 | 0.817 |
| Rv2017 | 0.005 | 0.999 | -0.331 | 0.826 | -0.336 | 0.816 |
| Rv2018 | -0.030 | 0.999 | -0.252 | 0.806 | -0.222 | 0.816 |
| Rv2019 | 0.020 | NA | 0.203 | 0.846 | 0.183 | 0.819 |
| Rv2021c | -0.086 | 0.999 | -0.226 | 0.826 | -0.139 | 0.819 |
| Rv2022c | -0.009 | 0.999 | -0.398 | 0.848 | -0.388 | 0.822 |
| Rv2023c | -0.153 | 0.999 | 0.186 | 0.879 | 0.339 | 0.816 |
| Rv2024c | -0.058 | 0.999 | -0.174 | 0.879 | -0.116 | 0.884 |
| Rv2025c | -0.173 | NA | -0.395 | 0.846 | -0.222 | 0.866 |
| Rv2026c | -0.249 | NA | -0.355 | 0.826 | -0.106 | 0.889 |
| dosT | -0.306 | 0.999 | -0.674 | 0.016 | -0.367 | 0.816 |
| Rv2028c | -0.445 | 0.922 | -0.480 | 0.361 | -0.035 | 0.951 |
| pfkB | -0.597 | 0.999 | -0.539 | 0.826 | 0.058 | 0.958 |
| Rv2030c | -0.478 | 0.999 | -0.644 | 0.660 | -0.166 | 0.870 |
| hspX | -0.392 | 0.587 | -0.765 | 0.000 | -0.373 | 0.816 |
| acg | -0.413 | 0.260 | -0.584 | 0.002 | -0.171 | 0.816 |
| Rv2033c | -0.123 | 0.999 | 0.378 | 0.820 | 0.500 | 0.816 |
| Rv2034 | -0.426 | 0.999 | -0.352 | 0.858 | 0.074 | 0.955 |
| Rv2035 | -0.138 | 0.999 | -0.441 | 0.826 | -0.303 | 0.816 |
| Rv2036 | -0.246 | 0.999 | -0.235 | 0.834 | 0.011 | 0.988 |
| Rv2037c | -0.180 | 0.999 | -0.302 | 0.831 | -0.123 | 0.882 |
| Rv2038c | -0.015 | 0.999 | -0.254 | 0.848 | -0.239 | 0.817 |
| Rv2039c | -0.187 | NA | -0.284 | 0.848 | -0.097 | 0.921 |
| Rv2040c | 0.049 | 0.999 | -0.127 | 0.859 | -0.176 | 0.816 |
| Rv2041c | -0.083 | 0.999 | -0.026 | 0.951 | 0.056 | 0.875 |
| Rv2042c | -0.003 | 0.999 | -0.039 | 0.951 | -0.036 | 0.944 |
| pncA | 0.011 | 0.999 | 0.003 | 0.995 | -0.008 | 0.989 |
| Rv2044c | -0.049 | NA | -0.366 | 0.873 | -0.317 | 0.857 |
| lipT | 0.114 | 0.999 | -0.109 | 0.859 | -0.223 | 0.816 |
| lppI | -0.068 | 0.999 | 0.252 | 0.846 | 0.320 | 0.816 |
| Rv2047c | 0.006 | 0.999 | 0.396 | 0.829 | 0.390 | 0.816 |
| pks12 | 0.111 | 0.999 | 0.091 | 0.922 | -0.020 | 0.979 |
| Rv2049c | 0.056 | 0.999 | 0.422 | 0.826 | 0.365 | 0.816 |
| Rv2050 | -0.107 | 0.999 | -0.046 | 0.948 | 0.061 | 0.911 |
| ppm1 | 0.012 | 0.999 | 0.075 | 0.918 | 0.063 | 0.909 |
| Rv2052c | -0.061 | 0.999 | 0.379 | 0.832 | 0.440 | 0.816 |
| fxsA | -0.170 | 0.999 | 0.356 | 0.852 | 0.525 | 0.816 |
| Rv2054 | 0.085 | 0.999 | 0.157 | 0.848 | 0.073 | 0.892 |
| rpsR2 | 0.028 | NA | 0.075 | 0.906 | 0.047 | 0.930 |
| rpsN2 | -0.209 | NA | -0.372 | 0.826 | -0.163 | 0.854 |
| rpmG1 | -0.048 | NA | -0.561 | 0.865 | -0.513 | 0.846 |
| rpmB2 | -0.134 | NA | -0.061 | 0.966 | 0.073 | 0.947 |
| Rv2059 | 0.129 | 0.999 | -0.065 | 0.951 | -0.194 | 0.838 |
| Rv2060 | NA | NA | NA | NA | NA | NA |
| Rv2061c | 0.041 | 0.999 | 0.059 | 0.954 | 0.018 | 0.985 |
| cobN | 0.099 | 0.999 | 0.208 | 0.826 | 0.109 | 0.846 |
| mazE7 | -0.021 | NA | 0.234 | 0.878 | 0.255 | 0.836 |
| mazF7 | 0.072 | NA | -0.185 | 0.863 | -0.257 | 0.816 |
| cobG | -0.018 | 0.999 | 0.782 | 0.820 | 0.800 | 0.816 |
| cobH | 0.073 | 0.999 | 0.700 | 0.804 | 0.627 | 0.816 |
| cobIJ | -0.089 | 0.999 | 0.183 | 0.879 | 0.272 | 0.816 |
| Rv2067c | 0.148 | 0.999 | -0.090 | 0.885 | -0.238 | 0.816 |
| blaC | 0.073 | 0.999 | 0.347 | 0.829 | 0.274 | 0.816 |
| sigC | 0.080 | 0.999 | -0.090 | 0.879 | -0.170 | 0.816 |
| cobK | 0.068 | NA | 0.483 | 0.660 | 0.415 | 0.816 |
| cobM | 0.047 | NA | 0.042 | 0.970 | -0.006 | 0.996 |
| cobL | 0.110 | NA | 0.155 | 0.886 | 0.045 | 0.958 |
| Rv2073c | 0.128 | NA | 0.379 | 0.826 | 0.251 | 0.817 |
| Rv2074 | 0.180 | 0.999 | 0.265 | 0.848 | 0.085 | 0.930 |
| Rv2075c | 0.046 | 0.999 | 0.377 | 0.684 | 0.330 | 0.816 |
| Rv2076c | 0.030 | NA | 0.075 | 0.930 | 0.045 | 0.946 |
| Rv2077A | -0.157 | NA | -0.045 | 0.969 | 0.111 | 0.899 |
| Rv2077c | -0.338 | 0.999 | 0.112 | 0.922 | 0.450 | 0.816 |
| Rv2078 | -0.008 | NA | 0.169 | 0.895 | 0.177 | 0.868 |
| Rv2079 | -0.006 | 0.999 | -0.069 | 0.948 | -0.063 | 0.942 |
| lppJ | 0.105 | 0.999 | -0.262 | 0.886 | -0.367 | 0.835 |
| Rv2081c | -0.115 | 0.999 | 0.370 | 0.826 | 0.485 | 0.816 |
| Rv2082 | 0.249 | 0.999 | -0.320 | 0.886 | -0.569 | 0.816 |
| Rv2083 | 0.164 | 0.999 | 0.154 | 0.879 | -0.010 | 0.989 |
| Rv2084 | 0.268 | 0.999 | 0.114 | 0.879 | -0.154 | 0.819 |
| Rv2085 | 0.055 | NA | 0.538 | 0.826 | 0.483 | 0.816 |
| Rv2086 | 0.106 | NA | -0.351 | 0.848 | -0.457 | 0.816 |
| pknJ | 0.075 | 0.999 | -0.136 | 0.879 | -0.211 | 0.816 |
| pepE | -0.092 | NA | -0.010 | 0.988 | 0.082 | 0.904 |
| Rv2090 | -0.163 | NA | -0.078 | 0.934 | 0.085 | 0.909 |
| Rv2091c | -0.051 | 0.999 | -0.282 | 0.834 | -0.231 | 0.819 |
| helY | 0.054 | 0.999 | 0.093 | 0.886 | 0.040 | 0.942 |
| tatC | -0.104 | 0.999 | -0.177 | 0.853 | -0.074 | 0.909 |
| tatA | -0.142 | 0.999 | -0.523 | 0.805 | -0.381 | 0.816 |
| pafC | -0.009 | 0.999 | 0.429 | 0.826 | 0.438 | 0.816 |
| pafB | -0.125 | 0.999 | -0.075 | 0.948 | 0.050 | 0.955 |
| pafA | 0.169 | 0.999 | -0.140 | 0.879 | -0.309 | 0.816 |
| Rv2100 | 0.141 | 0.999 | -0.475 | 0.832 | -0.616 | 0.816 |
| helZ | 0.123 | 0.999 | 0.320 | 0.510 | 0.197 | 0.816 |
| Rv2102 | 0.088 | 0.999 | 0.109 | 0.850 | 0.021 | 0.956 |
| vapC37 | 0.112 | 0.999 | 0.384 | 0.659 | 0.272 | 0.816 |
| Rv2104a | -0.036 | 0.999 | 0.063 | 0.885 | 0.099 | 0.819 |
| vapB37 | -0.115 | 0.999 | 0.075 | 0.943 | 0.190 | 0.837 |
| PE22 | -0.373 | NA | -0.827 | 0.767 | -0.454 | 0.816 |
| PPE36 | -0.140 | 0.999 | -0.810 | 0.831 | -0.669 | 0.816 |
| prcA | -0.051 | 0.999 | -0.118 | 0.831 | -0.067 | 0.852 |
| prcB | 0.011 | 0.999 | 0.233 | 0.826 | 0.222 | 0.816 |
| pup | -0.001 | 0.999 | -0.211 | 0.843 | -0.209 | 0.816 |
| dop | 0.039 | 0.999 | -0.119 | 0.832 | -0.158 | 0.816 |
| Rv2113 | 0.040 | 0.999 | 0.370 | 0.732 | 0.330 | 0.816 |
| Rv2114 | 0.108 | 0.999 | 0.059 | 0.886 | -0.049 | 0.885 |
| mpa | 0.101 | 0.999 | -0.135 | 0.839 | -0.236 | 0.816 |
| lppK | -0.174 | 0.999 | 0.084 | 0.952 | 0.257 | 0.840 |
| Rv2117 | 0.084 | NA | 0.068 | 0.962 | -0.016 | 0.989 |
| Rv2118c | 0.069 | 0.999 | 0.458 | 0.767 | 0.389 | 0.816 |
| Rv2119 | 0.011 | 0.999 | -0.033 | 0.948 | -0.044 | 0.911 |
| Rv2120c | 0.222 | 0.999 | 0.225 | 0.660 | 0.002 | 0.996 |
| hisG | 0.128 | 0.999 | 0.303 | 0.831 | 0.175 | 0.849 |
| hisE | 0.068 | NA | 0.049 | 0.973 | -0.019 | 0.988 |
| PPE37 | 0.100 | 0.999 | -0.186 | 0.831 | -0.286 | 0.816 |
| metH | 0.118 | 0.999 | 0.054 | 0.922 | -0.064 | 0.882 |
| Rv2125 | -0.028 | 0.999 | 0.204 | 0.826 | 0.231 | 0.816 |
| PE_PGRS37 | -0.011 | NA | -0.013 | 0.994 | -0.002 | 0.999 |
| ansP1 | 0.067 | 0.999 | 0.119 | 0.879 | 0.052 | 0.926 |
| Rv2128 | 0.120 | 0.999 | -0.122 | 0.906 | -0.242 | 0.817 |
| Rv2129c | -0.063 | 0.999 | -0.002 | 0.996 | 0.061 | 0.936 |
| mshC | 0.018 | 0.999 | -0.131 | 0.848 | -0.149 | 0.816 |
| cysQ | -0.062 | 0.999 | 0.108 | 0.879 | 0.170 | 0.816 |
| Rv2132 | -0.071 | 0.999 | 0.033 | 0.971 | 0.104 | 0.896 |
| Rv2133c | 0.019 | 0.999 | -0.005 | 0.994 | -0.024 | 0.975 |
| Rv2134c | -0.002 | 0.999 | 0.003 | 0.995 | 0.005 | 0.994 |
| Rv2135c | 0.067 | 0.999 | -0.042 | 0.962 | -0.108 | 0.882 |
| Rv2136c | 0.011 | 0.999 | 0.110 | 0.891 | 0.098 | 0.883 |
| Rv2137c | 0.084 | 0.999 | 0.334 | 0.853 | 0.250 | 0.855 |
| lppL | -0.065 | 0.999 | -0.077 | 0.922 | -0.012 | 0.985 |
| pyrD | 0.163 | 0.999 | 0.193 | 0.826 | 0.031 | 0.942 |
| TB18.6 | 0.102 | NA | 0.114 | 0.883 | 0.013 | 0.985 |
| Rv2141c | 0.029 | 0.999 | 0.090 | 0.859 | 0.061 | 0.870 |
| parD2 | -0.215 | 0.999 | -0.056 | 0.963 | 0.159 | 0.879 |
| parE2 | -0.076 | 0.999 | -0.537 | 0.659 | -0.461 | 0.816 |
| Rv2143 | -0.030 | 0.999 | 0.274 | 0.831 | 0.303 | 0.816 |
| Rv2144c | 0.120 | 0.999 | -0.002 | 0.997 | -0.122 | 0.897 |
| wag31 | 0.108 | 0.999 | 0.010 | 0.986 | -0.098 | 0.877 |
| Rv2146c | 0.178 | 0.999 | 0.060 | 0.931 | -0.118 | 0.850 |
| Rv2147c | 0.212 | 0.999 | 0.072 | 0.879 | -0.140 | 0.816 |
| Rv2148c | 0.183 | 0.999 | -0.053 | 0.964 | -0.236 | 0.835 |
| yfiH | -0.181 | 0.999 | 0.113 | 0.929 | 0.293 | 0.819 |
| ftsZ | 0.022 | 0.999 | 0.295 | 0.820 | 0.273 | 0.816 |

| | | | | | | |
|---|---|---|---|---|---|---|
| gpsI | 0.042 | 0.999 | 0.019 | 0.970 | -0.024 | 0.951 |
| lppU | -0.137 | 0.999 | -0.174 | 0.879 | -0.037 | 0.964 |
| rpsO | -0.045 | 0.999 | -0.626 | 0.785 | -0.581 | 0.816 |
| ribF | -0.028 | 0.999 | -0.043 | 0.951 | -0.015 | 0.980 |
| Rv2787 | -0.089 | 0.999 | -0.078 | 0.947 | 0.011 | 0.989 |
| sirR | -0.086 | 0.999 | 0.617 | 0.741 | 0.703 | 0.816 |
| fadE21 | -0.024 | 0.999 | 0.127 | 0.859 | 0.151 | 0.817 |
| ltp1 | 0.060 | 0.999 | -0.357 | 0.831 | -0.417 | 0.816 |
| Rv2791c | 0.004 | 0.999 | 0.071 | 0.879 | 0.067 | 0.862 |
| Rv2792c | 0.080 | 0.999 | 0.006 | 0.989 | -0.073 | 0.891 |
| truB | 0.100 | 0.999 | 0.434 | 0.826 | 0.334 | 0.816 |
| pptT | 0.069 | 0.999 | 0.015 | 0.988 | -0.054 | 0.958 |
| Rv2795c | 0.068 | 0.999 | -0.114 | 0.859 | -0.181 | 0.816 |
| lppV | -0.103 | 0.999 | 0.108 | 0.862 | 0.211 | 0.816 |
| Rv2797c | -0.065 | 0.999 | 0.103 | 0.938 | 0.169 | 0.877 |
| Rv2798c | 0.032 | 0.999 | -0.145 | 0.927 | -0.176 | 0.892 |
| Rv2799 | -0.066 | 0.999 | -0.137 | 0.831 | -0.071 | 0.862 |
| Rv2800 | -0.044 | 0.999 | 0.372 | 0.831 | 0.415 | 0.816 |
| mazE9 | 0.062 | NA | -0.588 | 0.826 | -0.651 | 0.816 |
| mazF9 | 0.014 | NA | 0.320 | 0.843 | 0.306 | 0.817 |
| Rv2802c | -0.035 | 0.999 | 0.351 | 0.831 | 0.386 | 0.816 |
| Rv2803 | 0.136 | 0.999 | -0.363 | 0.879 | -0.499 | 0.817 |
| Rv2804c | -0.649 | NA | -0.884 | 0.843 | -0.235 | 0.932 |
| Rv2805 | -0.131 | NA | 0.132 | 0.895 | 0.263 | 0.816 |
| Rv2806 | -0.018 | NA | -0.054 | 0.970 | -0.036 | 0.977 |
| Rv2807 | -0.344 | NA | -0.900 | 0.826 | -0.556 | 0.817 |
| Rv2808 | 0.055 | 0.999 | -0.352 | 0.832 | -0.407 | 0.816 |
| Rv2809 | 0.080 | 0.999 | 0.079 | 0.973 | -0.001 | 0.999 |
| Rv2811 | -0.184 | NA | 0.175 | 0.933 | 0.359 | 0.846 |
| Rv2812 | 0.123 | NA | -0.368 | 0.879 | -0.490 | 0.832 |
| Rv2813 | 0.294 | NA | -0.895 | NA | -1.189 | NA |
| Rv2816c | -0.132 | NA | -0.396 | 0.848 | -0.265 | 0.859 |
| Rv2817c | -0.073 | NA | -0.792 | 0.820 | -0.719 | 0.816 |
| Rv2818c | 0.005 | 0.999 | -0.371 | 0.836 | -0.376 | 0.816 |
| Rv2819c | -0.040 | 0.999 | -0.586 | 0.767 | -0.546 | 0.816 |
| Rv2820c | -0.145 | 0.999 | -0.398 | 0.770 | -0.253 | 0.816 |
| Rv2821c | -0.039 | 0.999 | -0.341 | 0.826 | -0.302 | 0.816 |
| Rv2822c | 0.053 | NA | -0.536 | 0.832 | -0.589 | 0.816 |
| Rv2823c | -0.074 | 0.999 | -0.379 | 0.826 | -0.305 | 0.816 |
| Rv2824c | 0.120 | 0.999 | -0.073 | 0.922 | -0.193 | 0.816 |
| Rv2825c | -0.170 | 0.999 | 0.195 | 0.826 | 0.364 | 0.816 |
| Rv2826c | -0.104 | 0.999 | -0.118 | 0.834 | -0.015 | 0.967 |
| Rv2827c | -0.202 | 0.999 | 0.204 | 0.879 | 0.406 | 0.816 |
| Rv2828A | -0.479 | NA | 0.723 | 0.826 | 1.202 | 0.816 |
| Rv2828c | -0.013 | 0.999 | 0.319 | 0.826 | 0.332 | 0.816 |
| vapC22 | -0.301 | 0.999 | 0.176 | 0.920 | 0.477 | 0.816 |
| vapB22 | -0.067 | NA | -0.112 | 0.899 | -0.045 | 0.950 |
| echA16 | 0.085 | 0.999 | 0.393 | 0.843 | 0.307 | 0.835 |
| ugpC | -0.116 | 0.999 | 0.179 | 0.832 | 0.295 | 0.816 |
| ugpB | -0.124 | 0.999 | -0.260 | 0.820 | -0.136 | 0.824 |
| ugpE | -0.002 | 0.999 | -0.287 | 0.820 | -0.285 | 0.816 |
| ugpA | 0.029 | 0.999 | -0.144 | 0.856 | -0.173 | 0.816 |
| dinF | 0.029 | 0.999 | 0.185 | 0.879 | 0.157 | 0.866 |
| Rv2837c | 0.056 | 0.999 | 0.142 | 0.859 | 0.086 | 0.884 |
| rbfA | -0.056 | 0.999 | -0.256 | 0.848 | -0.200 | 0.846 |
| infB | -0.058 | 0.999 | -0.058 | 0.913 | 0.000 | 0.999 |
| Rv2840c | 0.013 | 0.999 | -0.310 | 0.879 | -0.322 | 0.855 |
| nusA | -0.076 | 0.999 | 0.039 | 0.958 | 0.115 | 0.855 |
| Rv2842c | 0.004 | 0.999 | -0.029 | 0.961 | -0.033 | 0.942 |
| Rv2843 | -0.026 | 0.999 | -0.053 | 0.962 | -0.027 | 0.977 |
| Rv2844 | 0.106 | 0.999 | 0.140 | 0.843 | 0.034 | 0.939 |
| proS | -0.059 | 0.999 | 0.116 | 0.854 | 0.175 | 0.816 |
| efpA | 0.081 | 0.999 | 0.174 | 0.879 | 0.093 | 0.907 |
| cysG | -0.040 | 0.999 | 0.254 | 0.801 | 0.294 | 0.816 |
| cobB | 0.036 | 0.999 | 0.327 | 0.834 | 0.291 | 0.817 |
| cobO | -0.045 | 0.999 | -0.047 | 0.962 | -0.002 | 0.998 |
| Rv2850c | 0.017 | 0.999 | 0.238 | 0.832 | 0.221 | 0.816 |
| Rv2851c | 0.005 | 0.999 | -0.081 | 0.890 | -0.086 | 0.866 |
| mqo | 0.088 | 0.999 | -0.042 | 0.918 | -0.129 | 0.816 |
| PE_PGRS48 | 0.068 | 0.999 | -0.170 | 0.923 | -0.238 | 0.868 |
| Rv2854 | 0.147 | 0.999 | -0.020 | 0.974 | -0.168 | 0.831 |
| mtr | 0.244 | 0.999 | -0.080 | 0.895 | -0.324 | 0.816 |
| nicT | -0.007 | 0.999 | -0.288 | 0.826 | -0.281 | 0.816 |
| Rv2857c | -0.034 | 0.999 | -0.027 | 0.955 | 0.007 | 0.986 |
| aldC | -0.112 | 0.999 | 0.054 | 0.970 | 0.165 | 0.889 |
| Rv2859c | 0.071 | 0.999 | 0.335 | 0.826 | 0.264 | 0.816 |
| glnA4 | -0.066 | 0.999 | -0.055 | 0.940 | 0.012 | 0.985 |
| mapB | 0.070 | 0.999 | -0.167 | 0.879 | -0.237 | 0.817 |
| vapB23 | -0.546 | NA | -1.591 | 0.848 | -1.046 | 0.865 |
| Rv2862c | -0.192 | 0.999 | 0.128 | 0.926 | 0.321 | 0.817 |
| vapC23 | 0.090 | NA | -0.130 | 0.970 | -0.220 | 0.939 |
| Rv2864c | 0.044 | 0.999 | 0.207 | 0.834 | 0.163 | 0.831 |
| relF | -0.111 | 0.999 | -0.373 | 0.820 | -0.261 | 0.816 |
| relG | -0.163 | NA | -0.311 | 0.848 | -0.148 | 0.896 |
| Rv2867c | -0.105 | 0.999 | 0.328 | 0.859 | 0.433 | 0.816 |
| gcpE | 0.029 | 0.999 | 0.003 | 0.995 | -0.027 | 0.947 |
| rip | -0.020 | 0.999 | 0.184 | 0.885 | 0.204 | 0.858 |
| dxr | -0.058 | 0.999 | 0.112 | 0.927 | 0.171 | 0.866 |
| vapB43 | -0.111 | NA | 0.178 | 0.919 | 0.288 | 0.846 |
| vapC43 | -0.015 | NA | -0.173 | 0.879 | -0.158 | 0.855 |
| mpt83 | -0.163 | 0.999 | -0.260 | 0.831 | -0.097 | 0.892 |
| dipZ | -0.035 | 0.999 | -0.237 | 0.866 | -0.202 | 0.856 |
| mpt70 | -0.255 | 0.999 | -0.267 | 0.795 | -0.012 | 0.983 |
| Rv2876 | 0.046 | 0.999 | -0.253 | 0.879 | -0.299 | 0.835 |
| Rv2877c | -0.148 | 0.999 | 0.266 | 0.853 | 0.413 | 0.816 |
| mpt53 | -0.010 | 0.999 | -0.358 | 0.801 | -0.348 | 0.816 |
| cdsA | 0.060 | 0.999 | 0.187 | 0.848 | 0.128 | 0.857 |
| frr | 0.088 | 0.999 | -0.381 | 0.843 | -0.468 | 0.816 |
| pyrH | -0.010 | 0.999 | -0.059 | 0.970 | -0.049 | 0.967 |
| Rv2884 | 0.125 | 0.999 | 0.103 | 0.879 | -0.022 | 0.962 |
| Rv2885c | -0.079 | 0.999 | -0.047 | 0.888 | 0.032 | 0.904 |
| Rv2886c | -0.038 | 0.999 | -0.423 | 0.848 | -0.384 | 0.819 |
| Rv2887 | -0.173 | 0.999 | 0.678 | 0.772 | 0.850 | 0.816 |
| amiC | 0.033 | 0.999 | 0.282 | 0.831 | 0.248 | 0.816 |
| tsf | 0.094 | 0.999 | -0.024 | 0.971 | -0.118 | 0.854 |
| rpsB | 0.090 | 0.999 | -0.077 | 0.962 | -0.167 | 0.896 |
| Rv2891 | -0.124 | NA | -0.093 | 0.962 | 0.030 | 0.986 |
| PPE45 | -0.246 | 0.999 | 0.390 | 0.659 | 0.636 | 0.264 |
| Rv2893 | -0.323 | NA | 0.084 | 0.946 | 0.407 | 0.816 |
| xerC | 0.112 | 0.999 | 0.455 | 0.832 | 0.343 | 0.824 |
| viuB | -0.015 | 0.999 | 0.110 | 0.909 | 0.125 | 0.870 |
| Rv2896c | 0.042 | 0.999 | 0.598 | 0.676 | 0.556 | 0.816 |
| Rv2897c | 0.055 | 0.999 | 0.226 | 0.832 | 0.171 | 0.832 |
| Rv2898c | 0.055 | 0.999 | 0.067 | 0.934 | 0.012 | 0.986 |
| fdhD | 0.081 | 0.999 | -0.027 | 0.971 | -0.108 | 0.868 |
| fdhF | -0.028 | 0.999 | -0.066 | 0.879 | -0.038 | 0.900 |
| Rv2901c | -0.044 | 0.999 | -0.053 | 0.981 | -0.009 | 0.997 |
| rnhB | 0.079 | 0.999 | 0.226 | 0.848 | 0.147 | 0.855 |
| lepB | 0.018 | 0.999 | 0.162 | 0.879 | 0.143 | 0.868 |
| rplS | 0.140 | 0.999 | -0.036 | 0.973 | -0.176 | 0.868 |
| lppW | 0.050 | 0.999 | 0.143 | 0.848 | 0.093 | 0.862 |
| trmD | 0.106 | 0.999 | -0.168 | 0.859 | -0.274 | 0.816 |
| rimM | -0.023 | 0.999 | -0.201 | 0.832 | -0.178 | 0.817 |
| Rv2908c | 0.178 | 0.999 | -0.561 | 0.820 | -0.739 | 0.816 |
| rpsP | 0.026 | 0.999 | -0.019 | 0.970 | -0.045 | 0.904 |
| Rv2910c | -0.077 | NA | -0.041 | 0.962 | 0.036 | 0.956 |
| dacB2 | 0.115 | 0.999 | 0.145 | 0.848 | 0.030 | 0.951 |
| Rv2912c | -0.133 | 0.999 | -0.172 | 0.879 | -0.038 | 0.961 |
| Rv2913c | -0.192 | 0.999 | -0.335 | 0.610 | -0.142 | 0.819 |
| pknI | 0.114 | 0.999 | 0.069 | 0.928 | -0.045 | 0.942 |
| Rv2915c | 0.123 | 0.999 | -0.126 | 0.868 | -0.249 | 0.816 |
| ffh | -0.034 | 0.999 | 0.142 | 0.848 | 0.175 | 0.816 |
| Rv2917 | 0.042 | NA | -0.604 | 0.826 | -0.646 | 0.816 |
| glnD | -0.053 | 0.999 | -0.055 | 0.931 | -0.002 | 0.997 |
| glnB | -0.087 | 0.999 | -0.536 | 0.848 | -0.449 | 0.835 |
| amt | -0.096 | 0.999 | -0.450 | 0.832 | -0.353 | 0.821 |
| ftsY | -0.064 | 0.999 | 0.135 | 0.848 | 0.199 | 0.816 |
| acyP | 0.104 | NA | 0.376 | 0.858 | 0.273 | 0.862 |
| smc | 0.060 | 0.999 | 0.072 | 0.848 | 0.012 | 0.958 |
| Rv2923c | 0.001 | NA | 0.242 | 0.837 | 0.241 | 0.816 |
| fpg | -0.016 | 0.999 | 0.122 | 0.879 | 0.138 | 0.852 |
| rnc | -0.014 | 0.999 | 0.083 | 0.895 | 0.096 | 0.865 |
| Rv2926c | -0.094 | 0.999 | 0.121 | 0.914 | 0.215 | 0.835 |
| Rv2927c | 0.113 | 0.999 | -0.041 | 0.923 | -0.154 | 0.816 |
| tesA | 0.191 | 0.999 | -0.096 | 0.907 | -0.287 | 0.816 |
| Rv2929 | 0.073 | 0.999 | -0.081 | 0.926 | -0.154 | 0.842 |
| fadD26 | -0.204 | 0.999 | -0.544 | 0.043 | -0.340 | 0.816 |
| ppsA | -0.289 | 0.293 | -0.358 | 0.016 | -0.069 | 0.849 |
| ppsB | -0.224 | 0.999 | -0.363 | 0.684 | -0.139 | 0.837 |
| ppsC | -0.227 | 0.999 | -0.279 | 0.293 | -0.052 | 0.881 |
| ppsD | -0.021 | 0.999 | -0.018 | 0.973 | 0.003 | 0.996 |
| ppsE | 0.080 | 0.999 | 0.101 | 0.895 | 0.021 | 0.973 |
| drrA | 0.037 | 0.999 | -0.212 | 0.831 | -0.249 | 0.816 |
| drrB | 0.058 | 0.999 | -0.143 | 0.879 | -0.202 | 0.819 |
| drrC | 0.117 | 0.999 | -0.023 | 0.977 | -0.139 | 0.879 |
| papA5 | 0.229 | 0.999 | 0.002 | 0.997 | -0.227 | 0.837 |
| mas | -0.006 | 0.999 | -0.154 | 0.719 | -0.148 | 0.816 |
| fadD28 | 0.067 | 0.999 | -0.176 | 0.848 | -0.243 | 0.816 |
| mmpL7 | 0.036 | 0.999 | 0.049 | 0.948 | 0.013 | 0.985 |
| Rv2943 | -0.079 | 0.999 | -0.056 | 0.944 | 0.023 | 0.970 |
| Rv2943A | -0.531 | NA | -0.229 | 0.931 | 0.302 | 0.888 |
| Rv2944 | 0.007 | 0.999 | -0.105 | 0.934 | -0.112 | 0.912 |
| lppX | 0.034 | 0.999 | -0.296 | 0.826 | -0.330 | 0.816 |
| pks1 | 0.067 | 0.999 | 0.261 | 0.831 | 0.194 | 0.819 |
| pks15 | 0.040 | 0.999 | 0.069 | 0.970 | 0.030 | 0.986 |
| fadD22 | 0.087 | 0.999 | -0.151 | 0.879 | -0.237 | 0.817 |
| Rv2949c | 0.128 | 0.999 | -0.141 | 0.914 | -0.269 | 0.826 |
| fadD29 | 0.044 | 0.999 | -0.155 | 0.879 | -0.199 | 0.836 |
| Rv2951c | 0.031 | 0.999 | -0.010 | 0.989 | -0.041 | 0.955 |
| Rv2952 | 0.176 | 0.999 | 0.022 | 0.974 | -0.154 | 0.836 |
| Rv2953 | 0.144 | 0.999 | 0.083 | 0.879 | -0.061 | 0.875 |
| Rv2954c | 0.145 | 0.999 | -0.145 | 0.848 | -0.290 | 0.816 |
| Rv2955c | -0.022 | 0.999 | 0.396 | 0.795 | 0.418 | 0.816 |
| Rv2956 | 0.024 | 0.999 | -0.123 | 0.879 | -0.147 | 0.835 |
| Rv2957 | -0.022 | 0.999 | -0.207 | 0.826 | -0.185 | 0.816 |

| | | | | | | |
|---|---|---|---|---|---|---|
| Rv3594 | -0.036 | 0.999 | 0.244 | 0.831 | 0.280 | 0.816 |
| PE_PGRS59 | -0.095 | 0.999 | -0.183 | 0.859 | -0.088 | 0.904 |
| clpC1 | 0.007 | 0.999 | -0.054 | 0.917 | -0.061 | 0.881 |
| lsr2 | -0.159 | 0.999 | -0.140 | 0.942 | 0.019 | 0.990 |
| lysS | -0.178 | 0.999 | -0.121 | 0.854 | 0.057 | 0.897 |
| Rv3600c | -0.483 | 0.999 | -3.580 | 0.000 | -3.097 | 0.000 |
| panD | 1.296 | 0.000 | 0.331 | 0.396 | -0.965 | 0.000 |
| panC | 0.216 | 0.999 | 0.272 | 0.832 | 0.055 | 0.942 |
| Rv3603c | 0.257 | 0.999 | 0.483 | 0.701 | 0.227 | 0.819 |
| Rv3604c | 0.023 | 0.999 | -0.150 | 0.879 | -0.172 | 0.837 |
| Rv3605c | -0.041 | 0.999 | 0.219 | 0.883 | 0.260 | 0.846 |
| folK | 0.118 | 0.999 | -0.521 | 0.795 | -0.639 | 0.816 |
| folB | -0.021 | 0.999 | -0.105 | 0.922 | -0.085 | 0.924 |
| folP1 | -0.104 | 0.999 | -0.051 | 0.948 | 0.053 | 0.936 |
| folE | 0.038 | 0.999 | -0.130 | 0.879 | -0.168 | 0.825 |
| ftsH | -0.073 | 0.999 | -0.071 | 0.879 | 0.002 | 0.996 |
| Rv3612c | 0.098 | 0.999 | 0.121 | 0.879 | 0.024 | 0.969 |
| Rv3613c | 0.095 | 0.999 | -0.229 | 0.873 | -0.324 | 0.816 |
| espD | 0.065 | 0.999 | -0.283 | 0.659 | -0.347 | 0.816 |
| espC | 0.125 | 0.999 | -0.182 | 0.848 | -0.308 | 0.816 |
| espA | 0.105 | 0.999 | 0.066 | 0.921 | -0.039 | 0.939 |
| ephA | 0.046 | 0.999 | 0.065 | 0.918 | 0.019 | 0.967 |
| Rv3618 | 0.217 | 0.999 | -0.024 | 0.959 | -0.242 | 0.816 |
| esxV | -0.182 | NA | -0.022 | 0.974 | 0.160 | 0.837 |
| esxW | -0.029 | 0.999 | -0.627 | 0.831 | -0.599 | 0.816 |
| PPE65 | 0.025 | NA | 0.112 | 0.895 | 0.087 | 0.902 |
| PE32 | -0.026 | NA | -0.817 | 0.832 | -0.792 | 0.816 |
| lpqG | 0.113 | 0.999 | 0.019 | 0.960 | -0.094 | 0.816 |
| hpt | 0.199 | 0.999 | 0.110 | 0.879 | -0.089 | 0.879 |
| mesJ | -0.099 | 0.999 | 0.282 | 0.832 | 0.382 | 0.816 |
| Rv3626c | -0.043 | 0.999 | -0.079 | 0.923 | -0.035 | 0.955 |
| Rv3627c | -0.065 | 0.999 | 0.009 | 0.988 | 0.074 | 0.906 |
| ppa | 0.019 | 0.999 | -0.044 | 0.970 | -0.063 | 0.944 |
| Rv3629c | 0.092 | 0.999 | 0.109 | 0.883 | 0.017 | 0.977 |
| Rv3630 | -0.068 | 0.999 | 0.368 | 0.826 | 0.435 | 0.816 |
| Rv3631 | 0.061 | 0.999 | 0.304 | 0.826 | 0.243 | 0.816 |
| Rv3632 | 0.148 | 0.999 | 0.273 | 0.826 | 0.125 | 0.854 |
| Rv3633 | 0.126 | 0.999 | 0.118 | 0.924 | -0.008 | 0.994 |
| galE1 | -0.082 | 0.999 | 0.055 | 0.948 | 0.137 | 0.855 |
| Rv3635 | 0.028 | 0.999 | 0.112 | 0.853 | 0.085 | 0.854 |
| Rv3638 | -0.156 | NA | -1.745 | NA | -1.589 | NA |
| Rv3639c | -0.194 | NA | -0.443 | 0.854 | -0.249 | 0.883 |
| Rv3640c | -0.335 | NA | -0.429 | 0.825 | -0.094 | 0.917 |
| fic | -0.076 | 0.999 | -0.301 | 0.820 | -0.226 | 0.816 |
| Rv3642c | -0.092 | 0.999 | -0.398 | 0.895 | -0.306 | 0.898 |
| Rv3643 | 0.115 | NA | -0.331 | 0.948 | -0.446 | 0.912 |
| Rv3644c | -0.007 | 0.999 | 0.219 | 0.854 | 0.226 | 0.820 |
| Rv3645 | -0.243 | 0.999 | -0.062 | 0.961 | 0.180 | 0.866 |
| topA | 0.111 | 0.999 | 0.074 | 0.879 | -0.036 | 0.916 |
| Rv3647c | 0.074 | 0.999 | 0.797 | 0.660 | 0.723 | 0.816 |
| cspA | 0.139 | 0.999 | 0.027 | 0.977 | -0.112 | 0.916 |
| Rv3649 | 0.174 | 0.999 | 0.356 | 0.660 | 0.182 | 0.816 |
| PE33 | -0.169 | NA | -0.237 | 0.859 | -0.069 | 0.942 |
| Rv3651 | -0.050 | 0.999 | -0.358 | 0.832 | -0.308 | 0.817 |
| PE_PGRS60 | 0.098 | NA | 0.226 | 0.832 | 0.128 | 0.859 |
| PE_PGRS61 | -0.323 | NA | 0.072 | 0.970 | 0.395 | 0.835 |
| Rv3654c | 0.004 | NA | 0.573 | 0.831 | 0.569 | 0.816 |
| Rv3655c | 0.213 | NA | 0.625 | 0.826 | 0.412 | 0.819 |
| Rv3656c | 0.164 | NA | -0.251 | 0.886 | -0.415 | 0.819 |
| Rv3657c | -0.017 | NA | 0.379 | 0.848 | 0.395 | 0.819 |
| Rv3658c | -0.019 | 0.999 | 0.446 | 0.848 | 0.466 | 0.817 |
| Rv3659c | 0.095 | 0.999 | -0.091 | 0.912 | -0.187 | 0.819 |
| Rv3660c | -0.054 | 0.999 | 0.330 | 0.848 | 0.384 | 0.816 |
| Rv3661 | 0.088 | 0.999 | -0.155 | 0.832 | -0.243 | 0.816 |
| Rv3662c | 0.291 | NA | -0.003 | 0.995 | -0.294 | 0.816 |
| dppD | 0.069 | 0.999 | -0.020 | 0.971 | -0.089 | 0.862 |
| dppC | 0.091 | NA | -0.124 | 0.909 | -0.214 | 0.835 |
| dppB | 0.110 | NA | -0.143 | 0.879 | -0.253 | 0.816 |
| dppA | 0.039 | 0.999 | -0.106 | 0.848 | -0.145 | 0.816 |
| acs | -0.006 | 0.999 | -0.144 | 0.831 | -0.138 | 0.816 |
| Rv3668c | 0.100 | 0.999 | 0.227 | 0.831 | 0.127 | 0.843 |
| Rv3669 | 0.099 | 0.999 | -0.133 | 0.885 | -0.233 | 0.816 |
| ephE | 0.023 | 0.999 | 0.336 | 0.832 | 0.313 | 0.816 |
| Rv3671c | 0.018 | 0.999 | 0.355 | 0.832 | 0.338 | 0.816 |
| Rv3672c | -0.073 | 0.999 | 0.393 | 0.826 | 0.466 | 0.816 |
| Rv3673c | 0.183 | 0.999 | 0.338 | 0.769 | 0.156 | 0.831 |
| nth | 0.059 | 0.999 | -0.093 | 0.859 | -0.152 | 0.816 |
| Rv3675 | -0.161 | 0.999 | -0.148 | 0.875 | 0.013 | 0.985 |
| crp | 0.133 | 0.999 | 0.120 | 0.871 | -0.014 | 0.980 |
| Rv3677c | 0.015 | 0.999 | -0.141 | 0.853 | -0.156 | 0.817 |
| Rv3678A | 0.103 | 0.999 | 0.144 | 0.887 | 0.040 | 0.960 |
| Rv3678c | 0.071 | 0.999 | 0.180 | 0.879 | 0.109 | 0.896 |
| Rv3679 | 0.050 | 0.999 | -0.077 | 0.895 | -0.127 | 0.831 |
| Rv3680 | 0.100 | 0.999 | -0.014 | 0.973 | -0.114 | 0.819 |
| whiB4 | 0.058 | 0.999 | -0.061 | 0.931 | -0.119 | 0.846 |
| ponA2 | -0.054 | 0.999 | -0.276 | 0.701 | -0.222 | 0.816 |
| Rv3683 | 0.079 | 0.999 | -0.128 | 0.826 | -0.207 | 0.816 |
| Rv3684 | -0.008 | 0.999 | 0.045 | 0.959 | 0.054 | 0.939 |
| cyp137 | 0.097 | 0.999 | 0.102 | 0.834 | 0.005 | 0.987 |
| Rv3686c | -0.114 | 0.999 | 0.282 | 0.859 | 0.396 | 0.816 |
| rsfB | 0.071 | 0.999 | -0.303 | 0.848 | -0.374 | 0.816 |
| Rv3688c | -0.008 | 0.999 | 0.136 | 0.906 | 0.145 | 0.879 |
| Rv3689 | 0.114 | 0.999 | 0.298 | 0.820 | 0.184 | 0.816 |
| Rv3690 | 0.023 | 0.999 | 0.036 | 0.951 | 0.013 | 0.979 |
| Rv3691 | 0.074 | 0.999 | -0.057 | 0.922 | -0.131 | 0.817 |
| moxR2 | -0.031 | 0.999 | -0.097 | 0.879 | -0.067 | 0.896 |
| Rv3693 | 0.105 | 0.999 | 0.498 | 0.660 | 0.393 | 0.816 |
| Rv3694c | 0.012 | 0.999 | 0.057 | 0.919 | 0.045 | 0.920 |
| Rv3695 | -0.151 | 0.999 | -0.011 | 0.985 | 0.140 | 0.836 |
| glpK | 0.055 | 0.999 | -0.017 | 0.971 | -0.072 | 0.867 |
| vapB48 | 0.069 | 0.999 | -0.152 | 0.859 | -0.221 | 0.816 |
| vapC48 | 0.054 | 0.999 | -0.496 | 0.463 | -0.550 | 0.816 |
| Rv3698 | 0.038 | 0.999 | -0.183 | 0.859 | -0.221 | 0.817 |
| Rv3699 | -0.229 | 0.999 | 0.050 | 0.958 | 0.279 | 0.816 |
| egtE | 0.033 | 0.999 | 0.275 | 0.848 | 0.242 | 0.827 |
| egtD | 0.123 | 0.999 | 0.149 | 0.832 | 0.025 | 0.951 |
| egtC | 0.001 | 0.999 | -0.122 | 0.879 | -0.124 | 0.854 |
| etgB | -0.073 | 0.999 | 0.112 | 0.916 | 0.185 | 0.838 |
| gshA | 0.069 | 0.999 | 0.126 | 0.879 | 0.058 | 0.924 |
| Rv3705A | 0.197 | 0.999 | -0.872 | 0.826 | -1.070 | 0.816 |
| Rv3705c | 0.185 | 0.999 | -0.090 | 0.879 | -0.275 | 0.816 |
| Rv3706c | 0.284 | 0.999 | 0.232 | 0.919 | -0.051 | 0.977 |
| Rv3707c | -0.198 | 0.999 | 0.188 | 0.879 | 0.386 | 0.816 |
| asd | -0.008 | 0.999 | 0.251 | 0.858 | 0.259 | 0.825 |
| ask | 0.016 | 0.999 | -0.190 | 0.826 | -0.206 | 0.816 |
| leuA | -0.115 | 0.999 | -0.048 | 0.907 | 0.068 | 0.854 |
| dnaQ | 0.062 | 0.999 | 0.214 | 0.832 | 0.153 | 0.835 |
| Rv3712 | 0.083 | 0.999 | 0.414 | 0.825 | 0.331 | 0.816 |
| cobQ2 | 0.207 | NA | 0.404 | 0.588 | 0.197 | 0.816 |
| Rv3714c | 0.093 | 0.999 | -0.004 | 0.995 | -0.097 | 0.891 |
| recR | 0.106 | 0.999 | 0.159 | 0.826 | 0.053 | 0.878 |
| Rv3716c | 0.127 | 0.999 | 0.053 | 0.912 | -0.074 | 0.855 |
| Rv3717 | -0.023 | 0.999 | 0.246 | 0.806 | 0.269 | 0.816 |
| Rv3718c | -0.029 | 0.999 | 0.082 | 0.944 | 0.111 | 0.900 |
| Rv3719 | 0.092 | 0.999 | 0.098 | 0.927 | 0.006 | 0.996 |
| Rv3720 | 0.132 | 0.999 | -0.004 | 0.990 | -0.136 | 0.816 |
| dnaZX | 0.065 | 0.999 | 0.039 | 0.879 | -0.026 | 0.902 |
| Rv3722c | 0.083 | 0.999 | -0.013 | 0.977 | -0.096 | 0.860 |
| Rv3723 | -0.098 | 0.999 | 0.144 | 0.848 | 0.242 | 0.816 |
| Rv3725 | 0.051 | 0.999 | 0.371 | 0.820 | 0.320 | 0.816 |
| Rv3726 | 0.194 | 0.999 | 0.021 | 0.973 | -0.173 | 0.819 |
| Rv3727 | 0.141 | 0.999 | -0.223 | 0.886 | -0.364 | 0.819 |
| Rv3728 | 0.143 | 0.999 | 0.197 | 0.848 | 0.054 | 0.937 |
| Rv3729 | 0.116 | 0.999 | -0.108 | 0.895 | -0.224 | 0.816 |
| Rv3730c | 0.052 | 0.999 | 0.395 | 0.826 | 0.344 | 0.816 |
| ligC | 0.090 | 0.999 | 0.028 | 0.958 | -0.062 | 0.884 |
| Rv3732 | -0.011 | 0.999 | -0.280 | 0.820 | -0.269 | 0.816 |
| Rv3733c | 0.114 | 0.999 | 0.224 | 0.826 | 0.110 | 0.854 |
| tgs2 | 0.080 | 0.999 | 0.029 | 0.959 | -0.052 | 0.904 |
| Rv3735 | 0.074 | 0.999 | 0.139 | 0.853 | 0.065 | 0.897 |
| Rv3736 | 0.077 | 0.999 | -0.065 | 0.922 | -0.142 | 0.822 |
| Rv3737 | -0.084 | 0.999 | 0.125 | 0.897 | 0.209 | 0.831 |
| PPE66 | -0.177 | 0.999 | -0.074 | 0.922 | 0.102 | 0.866 |
| PPE67 | 0.080 | NA | 0.040 | 0.970 | -0.041 | 0.961 |
| Rv3740c | -0.044 | 0.999 | -0.174 | 0.859 | -0.130 | 0.862 |
| Rv3741c | 0.079 | 0.999 | -0.305 | 0.848 | -0.384 | 0.816 |
| Rv3742c | 0.179 | 0.999 | -0.207 | 0.878 | -0.385 | 0.816 |
| ctpJ | -0.211 | NA | -0.422 | 0.831 | -0.211 | 0.856 |
| nmtR | -0.078 | 0.999 | -0.131 | 0.879 | -0.053 | 0.932 |
| Rv3745c | -0.613 | NA | -0.264 | 0.918 | 0.349 | 0.866 |
| PE34 | -0.128 | NA | -0.042 | 0.962 | 0.086 | 0.899 |
| Rv3747 | -0.113 | 0.999 | -0.360 | 0.820 | -0.247 | 0.816 |
| Rv3748 | -0.119 | 0.999 | 0.191 | 0.885 | 0.310 | 0.819 |
| Rv3749c | 0.058 | 0.999 | -0.488 | 0.447 | -0.547 | 0.816 |
| Rv3750c | 0.192 | 0.999 | -0.418 | 0.895 | -0.610 | 0.836 |
| Rv3752c | -0.013 | NA | 0.415 | 0.820 | 0.428 | 0.816 |
| Rv3753c | 0.138 | 0.999 | 0.428 | 0.520 | 0.291 | 0.816 |
| tyrA | -0.088 | 0.999 | 0.182 | 0.832 | 0.270 | 0.816 |
| Rv3755c | 0.082 | 0.999 | -0.162 | 0.879 | -0.244 | 0.816 |
| proZ | 0.122 | 0.999 | 0.260 | 0.848 | 0.138 | 0.882 |
| proW | 0.164 | 0.999 | 0.029 | 0.974 | -0.135 | 0.882 |
| proV | -0.120 | 0.999 | 0.142 | 0.896 | 0.261 | 0.819 |
| proX | -0.025 | 0.999 | 0.105 | 0.893 | 0.130 | 0.854 |
| Rv3760 | 0.024 | 0.999 | -0.006 | 0.994 | -0.029 | 0.963 |
| fadE36 | 0.051 | 0.999 | 0.006 | 0.994 | -0.045 | 0.957 |
| Rv3762c | 0.080 | 0.999 | 0.088 | 0.892 | 0.008 | 0.988 |
| lpqH | -0.014 | 0.999 | 0.162 | 0.843 | 0.176 | 0.816 |
| tcrY | 0.004 | 0.999 | 0.193 | 0.859 | 0.189 | 0.836 |
| tcrX | 0.039 | 0.999 | -0.009 | 0.985 | -0.048 | 0.924 |
| Rv3766 | -0.132 | 0.999 | -0.476 | 0.826 | -0.344 | 0.816 |
| Rv3767c | -0.177 | 0.999 | -0.074 | 0.922 | 0.102 | 0.868 |
| Rv3768 | 0.024 | NA | -0.396 | 0.820 | -0.420 | 0.816 |
| Rv3769 | 0.022 | 0.999 | 0.040 | 0.958 | 0.019 | 0.975 |
| Rv3770c | -0.031 | 0.999 | 0.048 | 0.946 | 0.079 | 0.891 |
| Rv3771c | -0.063 | 0.999 | 0.169 | 0.885 | 0.232 | 0.835 |

| | | | | | | |
|---|---|---|---|---|---|---|
| Rv0515 | -0.028 | 0.999 | -0.061 | 0.930 | -0.033 | 0.951 |
| Rv0516c | 0.053 | 0.999 | 0.325 | 0.831 | 0.272 | 0.816 |
| Rv0517 | 0.015 | 0.999 | 0.253 | 0.826 | 0.238 | 0.816 |
| Rv0518 | 0.038 | 0.999 | 0.142 | 0.865 | 0.103 | 0.868 |
| Rv0519c | -0.067 | 0.999 | 0.104 | 0.915 | 0.171 | 0.838 |
| gabP | -0.025 | 0.999 | 0.160 | 0.869 | 0.185 | 0.827 |
| Rv0523c | 0.002 | 0.999 | 0.035 | 0.974 | 0.033 | 0.976 |
| hemL | -0.172 | 0.999 | 0.315 | 0.848 | 0.487 | 0.816 |
| Rv0525 | -0.161 | 0.999 | 0.181 | 0.885 | 0.342 | 0.816 |
| Rv0526 | 0.018 | 0.999 | 0.029 | 0.970 | 0.010 | 0.986 |
| ccdA | 0.003 | 0.999 | 0.265 | 0.854 | 0.261 | 0.831 |
| Rv0528 | 0.026 | 0.999 | 0.043 | 0.962 | 0.017 | 0.983 |
| ccsA | 0.080 | 0.999 | 0.167 | 0.848 | 0.086 | 0.884 |
| Rv0530 | -0.012 | 0.999 | 0.322 | 0.812 | 0.333 | 0.816 |
| Rv0530A | -0.174 | 0.999 | 0.207 | 0.868 | 0.380 | 0.816 |
| Rv0531 | -0.029 | 0.999 | 0.271 | 0.832 | 0.300 | 0.816 |
| PE_PGRS6 | -0.079 | 0.999 | -0.450 | 0.846 | -0.372 | 0.831 |
| fabH | 0.048 | 0.999 | 0.302 | 0.826 | 0.253 | 0.816 |
| menA | 0.204 | 0.999 | -0.098 | 0.927 | -0.301 | 0.816 |
| Rv0535 | -0.004 | 0.999 | -0.038 | 0.962 | -0.034 | 0.958 |
| galE3 | 0.057 | 0.999 | 0.349 | 0.826 | 0.292 | 0.816 |
| Rv0537c | -0.018 | 0.999 | 0.405 | 0.848 | 0.422 | 0.816 |
| Rv0538 | -0.002 | 0.999 | -0.063 | 0.940 | -0.061 | 0.930 |
| Rv0539 | -0.024 | 0.999 | 0.366 | 0.848 | 0.389 | 0.816 |
| Rv0540 | 0.080 | NA | 0.338 | 0.826 | 0.258 | 0.816 |
| Rv0541c | -0.023 | 0.999 | 0.358 | 0.826 | 0.382 | 0.816 |
| menE | -0.022 | 0.999 | 0.275 | 0.859 | 0.296 | 0.822 |
| Rv0543c | 0.033 | 0.999 | -0.079 | 0.949 | -0.112 | 0.913 |
| Rv0544c | 0.149 | 0.999 | 0.084 | 0.936 | -0.064 | 0.939 |
| pitA | 0.137 | 0.999 | 0.166 | 0.856 | 0.029 | 0.962 |
| Rv0546c | 0.012 | 0.999 | 0.066 | 0.895 | 0.054 | 0.896 |
| Rv0547c | -0.041 | 0.999 | 0.110 | 0.897 | 0.152 | 0.852 |
| menB | -0.047 | 0.999 | 0.090 | 0.854 | 0.138 | 0.816 |
| vapC3 | -0.049 | 0.999 | -0.202 | 0.832 | -0.153 | 0.830 |
| vapB3 | -0.210 | NA | -0.177 | 0.877 | 0.033 | 0.966 |
| fadD8 | -0.004 | 0.999 | -0.135 | 0.831 | -0.131 | 0.816 |
| Rv0552 | -0.011 | 0.999 | 0.269 | 0.820 | 0.280 | 0.816 |
| menC | -0.119 | NA | 0.403 | 0.826 | 0.522 | 0.816 |
| bpoC | 0.085 | 0.999 | 0.063 | 0.918 | -0.022 | 0.961 |
| menD | 0.088 | 0.999 | 0.326 | 0.820 | 0.239 | 0.816 |
| Rv0556 | 0.159 | 0.999 | -0.187 | 0.879 | -0.347 | 0.816 |
| mgtA | -0.005 | 0.999 | 0.167 | 0.895 | 0.172 | 0.868 |
| menH | -0.015 | 0.999 | 0.482 | 0.826 | 0.497 | 0.816 |
| Rv0559c | -0.099 | 0.999 | -0.386 | 0.659 | -0.287 | 0.816 |
| Rv0560c | 0.043 | 0.999 | 0.135 | 0.863 | 0.092 | 0.875 |
| Rv0561c | -0.004 | 0.999 | 0.116 | 0.859 | 0.120 | 0.831 |
| grcC1 | 0.095 | 0.999 | 0.214 | 0.831 | 0.119 | 0.854 |
| htpX | 0.061 | 0.999 | -0.081 | 0.913 | -0.142 | 0.835 |
| gpdA1 | -0.048 | 0.999 | -0.084 | 0.927 | -0.037 | 0.958 |
| Rv0565c | 0.041 | 0.999 | -0.484 | 0.831 | -0.525 | 0.816 |
| Rv0566c | -0.014 | 0.999 | -0.142 | 0.879 | -0.128 | 0.855 |
| Rv0567 | 0.038 | 0.999 | 0.029 | 0.967 | -0.009 | 0.988 |
| cyp135B1 | 0.027 | 0.999 | 0.244 | 0.659 | 0.217 | 0.816 |
| Rv0569 | -0.450 | 0.999 | -1.017 | 0.527 | -0.566 | 0.816 |
| nrdZ | -0.428 | 0.999 | -0.667 | 0.023 | -0.239 | 0.816 |
| Rv0571c | -0.450 | 0.906 | -0.457 | 0.442 | -0.008 | 0.989 |
| Rv0572c | -0.342 | 0.999 | -0.818 | 0.072 | -0.477 | 0.816 |
| pncB2 | -0.323 | 0.999 | -0.296 | 0.785 | 0.027 | 0.958 |
| Rv0574c | -0.408 | 0.999 | -0.494 | 0.361 | -0.086 | 0.892 |
| Rv0575c | -0.400 | NA | -0.278 | 0.848 | 0.122 | 0.893 |
| Rv0576 | -0.079 | 0.999 | 0.180 | 0.886 | 0.259 | 0.835 |
| TB27.3 | -0.146 | 0.999 | 0.057 | 0.924 | 0.204 | 0.816 |
| PE_PGRS7 | -0.074 | 0.999 | -0.139 | 0.941 | -0.065 | 0.964 |
| Rv0579 | -0.095 | 0.999 | 0.295 | 0.848 | 0.390 | 0.816 |
| Rv0580c | 0.176 | 0.999 | 0.464 | 0.826 | 0.288 | 0.817 |
| vapB26 | 0.009 | NA | 0.257 | 0.859 | 0.248 | 0.836 |
| vapC26 | 0.151 | NA | 0.041 | 0.961 | -0.110 | 0.868 |
| lpqN | 0.294 | 0.999 | -0.158 | 0.927 | -0.452 | 0.816 |
| Rv0584 | -0.071 | 0.999 | -0.059 | 0.942 | 0.013 | 0.986 |
| Rv0585c | 0.136 | 0.999 | -0.081 | 0.936 | -0.217 | 0.829 |
| mce2R | -0.063 | 0.999 | 0.120 | 0.848 | 0.183 | 0.816 |
| yrbE2A | -0.093 | 0.999 | 0.197 | 0.879 | 0.291 | 0.819 |
| Rv0588 | -0.035 | 0.999 | -0.272 | 0.820 | -0.238 | 0.816 |
| mce2A | -0.094 | 0.999 | -0.347 | 0.832 | -0.253 | 0.835 |
| mce2C | -0.155 | 0.999 | -0.475 | 0.699 | -0.320 | 0.816 |
| mce2D | -0.146 | 0.999 | -0.547 | 0.826 | -0.401 | 0.817 |
| lprL | -0.006 | 0.999 | -0.243 | 0.879 | -0.237 | 0.854 |
| mce2F | -0.098 | 0.999 | -0.128 | 0.848 | -0.029 | 0.943 |
| vapC4 | 0.033 | 0.999 | -0.283 | 0.832 | -0.316 | 0.816 |
| vapB4 | 0.110 | NA | 0.218 | 0.848 | 0.108 | 0.891 |
| Rv0597c | 0.033 | 0.999 | 0.202 | 0.879 | 0.169 | 0.866 |
| vapC27 | 0.160 | 0.999 | -0.224 | 0.832 | -0.384 | 0.816 |
| vapB27 | -0.201 | NA | -0.198 | 0.879 | 0.003 | 0.997 |
| Rv0600c | -0.455 | NA | -2.189 | NA | -1.733 | NA |
| Rv0601c | 0.119 | NA | -1.389 | 0.826 | -1.508 | 0.816 |
| tcrA | -0.143 | NA | -0.201 | 0.915 | -0.059 | 0.967 |
| Rv0603 | -0.149 | NA | -0.620 | 0.859 | -0.471 | 0.862 |
| lpqO | 0.082 | 0.999 | 0.273 | 0.826 | 0.191 | 0.816 |
| Rv0605 | 0.019 | 0.999 | 0.056 | 0.942 | 0.037 | 0.950 |
| Rv0607 | 0.003 | 0.999 | 0.122 | 0.934 | 0.119 | 0.922 |
| vapB28 | -0.015 | 0.999 | -0.282 | 0.826 | -0.267 | 0.816 |
| vapC28 | -0.076 | 0.999 | 0.189 | 0.826 | 0.265 | 0.816 |
| Rv0609A | -0.086 | 0.999 | -0.241 | 0.879 | -0.154 | 0.896 |
| Rv0610c | 0.077 | 0.999 | 0.113 | 0.895 | 0.036 | 0.958 |
| Rv0611c | -0.052 | 0.999 | 0.456 | 0.848 | 0.508 | 0.816 |
| Rv0612 | -0.138 | 0.999 | 0.187 | 0.879 | 0.326 | 0.816 |
| Rv0613c | 0.063 | 0.999 | 0.183 | 0.831 | 0.120 | 0.835 |
| Rv0614 | -0.046 | NA | -0.434 | 0.848 | -0.388 | 0.834 |
| Rv0615 | -0.151 | 0.999 | 0.128 | 0.883 | 0.278 | 0.816 |
| vapB29 | 0.046 | 0.999 | 0.123 | 0.879 | 0.077 | 0.899 |
| vapC29 | 0.045 | 0.999 | -0.040 | 0.962 | -0.086 | 0.896 |
| galK | -0.102 | NA | 0.003 | 0.997 | 0.105 | 0.939 |
| Rv0621 | -0.319 | 0.999 | 0.436 | 0.848 | 0.755 | 0.816 |
| Rv0622 | -0.269 | NA | -0.345 | 0.831 | -0.076 | 0.936 |
| vapB30 | 0.064 | NA | 0.676 | 0.826 | 0.612 | 0.816 |
| vapC30 | 0.058 | 0.999 | -0.081 | 0.886 | -0.139 | 0.817 |
| Rv0625c | -0.068 | 0.999 | 0.750 | 0.701 | 0.817 | 0.816 |
| vapB5 | -0.074 | 0.999 | 0.488 | 0.826 | 0.563 | 0.816 |
| vapC5 | -0.149 | 0.999 | 0.448 | 0.834 | 0.597 | 0.816 |
| Rv0628c | -0.014 | 0.999 | 0.253 | 0.848 | 0.267 | 0.816 |
| recD | 0.092 | 0.999 | 0.284 | 0.826 | 0.193 | 0.816 |
| recB | 0.144 | 0.999 | 0.260 | 0.769 | 0.116 | 0.835 |
| recC | 0.039 | 0.999 | -0.005 | 0.994 | -0.044 | 0.945 |
| echA3 | 0.073 | 0.999 | 0.075 | 0.931 | 0.002 | 0.997 |
| Rv0633c | -0.021 | 0.999 | -0.002 | 0.995 | 0.018 | 0.961 |
| Rv0634A | 0.063 | 0.999 | -0.068 | 0.941 | -0.131 | 0.866 |
| rpmG2 | 0.101 | 0.999 | -0.423 | 0.832 | -0.524 | 0.816 |
| Rv0634c | 0.049 | 0.999 | 0.131 | 0.848 | 0.082 | 0.865 |
| hadA | 0.093 | 0.999 | -0.055 | 0.973 | -0.148 | 0.922 |
| hadB | 0.150 | 0.999 | 0.032 | 0.970 | -0.117 | 0.866 |
| hadC | 0.128 | 0.999 | -0.083 | 0.884 | -0.211 | 0.816 |
| secE1 | -0.060 | 0.999 | 0.120 | 0.886 | 0.180 | 0.831 |
| nusG | -0.030 | 0.999 | -0.147 | 0.855 | -0.117 | 0.852 |
| rplK | 0.092 | 0.999 | -0.307 | 0.834 | -0.399 | 0.816 |
| rplA | -0.140 | 0.999 | 0.147 | 0.879 | 0.287 | 0.816 |
| mmaA4 | 0.048 | 0.999 | -0.130 | 0.848 | -0.178 | 0.816 |
| mmaA3 | 0.126 | 0.999 | -0.204 | 0.832 | -0.330 | 0.816 |
| mmaA2 | 0.071 | 0.999 | -0.210 | 0.826 | -0.281 | 0.816 |
| mmaA1 | 0.005 | 0.999 | -0.348 | 0.826 | -0.353 | 0.816 |
| lipG | -0.041 | 0.999 | -0.103 | 0.873 | -0.062 | 0.892 |
| Rv0647c | -0.136 | 0.999 | 0.150 | 0.905 | 0.286 | 0.819 |
| Rv0648 | -0.096 | 0.999 | -0.130 | 0.859 | -0.034 | 0.947 |
| fabD2 | -0.317 | NA | 0.054 | 0.975 | 0.370 | 0.852 |
| Rv0650 | -0.231 | NA | -0.482 | 0.859 | -0.251 | 0.896 |
| rplJ | -0.084 | 0.999 | 0.420 | 0.826 | 0.504 | 0.816 |
| rplL | -0.061 | 0.999 | 0.223 | 0.879 | 0.284 | 0.819 |
| Rv0653c | -0.090 | NA | -0.094 | 0.925 | -0.004 | 0.996 |
| Rv0654 | 0.123 | 0.999 | 0.040 | 0.948 | -0.083 | 0.868 |
| mkl | 0.239 | 0.999 | 0.014 | 0.973 | -0.225 | 0.816 |
| vapC6 | -0.106 | 0.999 | -0.188 | 0.826 | -0.082 | 0.863 |
| vapB6 | -0.208 | NA | -0.104 | 0.962 | 0.104 | 0.951 |
| Rv0658c | -0.059 | 0.999 | 0.209 | 0.830 | 0.268 | 0.816 |
| mazF2 | -0.094 | 0.999 | 0.157 | 0.895 | 0.251 | 0.830 |
| mazE2 | -0.065 | 0.999 | 0.000 | 1.000 | 0.065 | 0.951 |
| vapC7 | -0.128 | NA | 0.089 | 0.958 | 0.218 | 0.872 |
| vapB7 | 0.120 | NA | 0.393 | 0.832 | 0.273 | 0.837 |
| atsD | 0.017 | 0.999 | -0.049 | 0.948 | -0.067 | 0.909 |
| vapB8 | 0.131 | NA | -0.620 | 0.716 | -0.750 | 0.816 |
| vapC8 | -0.066 | NA | 0.065 | 0.959 | 0.131 | 0.896 |
| Rv0666 | -0.274 | NA | -0.582 | 0.772 | -0.308 | 0.817 |
| rpoB | -0.028 | 0.999 | -0.060 | 0.951 | -0.031 | 0.968 |
| rpoC | -0.001 | 0.999 | -0.070 | 0.938 | -0.069 | 0.925 |
| Rv0669c | 0.047 | 0.999 | -0.157 | 0.848 | -0.204 | 0.816 |
| end | -0.085 | 0.999 | -0.020 | 0.974 | 0.065 | 0.918 |
| lpqP | -0.138 | 0.999 | -0.174 | 0.879 | -0.036 | 0.962 |
| fadE8 | 0.040 | 0.999 | 0.124 | 0.879 | 0.084 | 0.896 |
| echA4 | 0.034 | 0.999 | 0.093 | 0.883 | 0.059 | 0.905 |
| Rv0674 | -0.028 | 0.999 | -0.105 | 0.885 | -0.077 | 0.896 |
| echA5 | -0.114 | 0.999 | 0.031 | 0.970 | 0.145 | 0.849 |
| mmpL5 | -0.145 | 0.999 | -0.312 | 0.445 | -0.167 | 0.816 |
| mmpS5 | -0.204 | 0.999 | -0.393 | 0.659 | -0.189 | 0.816 |
| Rv0678 | -0.079 | 0.999 | -0.051 | 0.958 | 0.027 | 0.970 |
| Rv0679c | 0.055 | 0.999 | -0.370 | 0.831 | -0.424 | 0.816 |
| Rv0680c | -0.010 | 0.999 | -0.281 | 0.826 | -0.270 | 0.816 |
| Rv0681 | 0.020 | 0.999 | 0.281 | 0.826 | 0.261 | 0.816 |
| rpsL | 0.040 | 0.999 | -0.148 | 0.859 | -0.187 | 0.816 |
| rpsG | 0.168 | 0.999 | 0.164 | 0.846 | -0.003 | 0.996 |
| fusA1 | 0.095 | 0.999 | -0.036 | 0.951 | -0.131 | 0.829 |
| tuf | 0.074 | 0.999 | 0.042 | 0.951 | -0.032 | 0.953 |
| Rv0686 | 0.046 | 0.999 | 0.043 | 0.946 | -0.003 | 0.996 |
| Rv0687 | 0.009 | 0.999 | -0.073 | 0.922 | -0.082 | 0.893 |
| Rv0688 | -0.096 | 0.999 | -0.079 | 0.946 | 0.017 | 0.986 |
| Rv0689c | -0.572 | NA | -0.019 | 0.994 | 0.553 | 0.846 |
| Rv0690c | -0.161 | 0.999 | 0.073 | 0.941 | 0.234 | 0.817 |
| Rv0691A | 0.071 | NA | -0.045 | 0.973 | -0.116 | 0.924 |

| | | | | | | |
|---|---|---|---|---|---|---|
| Rv1333 | -0.111 | 0.999 | 0.220 | 0.857 | 0.330 | 0.816 |
| mec | -0.053 | 0.999 | -0.451 | 0.843 | -0.398 | 0.819 |
| cysO | -0.205 | 0.999 | -0.387 | 0.820 | -0.182 | 0.837 |
| cysM | -0.041 | 0.999 | -0.074 | 0.927 | -0.033 | 0.957 |
| Rv1337 | -0.076 | 0.999 | 0.223 | 0.832 | 0.299 | 0.816 |
| murI | 0.016 | 0.999 | -0.377 | 0.663 | -0.393 | 0.816 |
| Rv1339 | -0.098 | 0.999 | -0.203 | 0.826 | -0.105 | 0.851 |
| rphA | 0.105 | 0.999 | -0.007 | 0.990 | -0.112 | 0.858 |
| Rv1341 | 0.063 | 0.999 | 0.091 | 0.880 | 0.027 | 0.954 |
| Rv1342c | -0.374 | 0.999 | -0.047 | 0.970 | 0.326 | 0.817 |
| lprD | -0.092 | 0.999 | -0.370 | 0.728 | -0.278 | 0.816 |
| mbtL | -0.100 | 0.999 | -0.125 | 0.940 | -0.026 | 0.986 |
| mbtM | -0.134 | 0.999 | 0.234 | 0.892 | 0.367 | 0.831 |
| mbtN | 0.096 | 0.999 | -0.041 | 0.956 | -0.137 | 0.837 |
| mbtK | 0.227 | 0.999 | 0.025 | 0.975 | -0.202 | 0.835 |
| irtA | 0.183 | 0.999 | 0.087 | 0.940 | -0.096 | 0.918 |
| irtB | 0.197 | 0.999 | 0.458 | 0.826 | 0.260 | 0.836 |
| fabG2 | 0.123 | 0.999 | -0.134 | 0.879 | -0.257 | 0.816 |
| Rv1351 | 0.128 | 0.999 | 0.006 | 0.995 | -0.122 | 0.904 |
| Rv1352 | -0.022 | 0.999 | -0.198 | 0.848 | -0.176 | 0.835 |
| Rv1353c | -0.145 | NA | -0.342 | 0.879 | -0.196 | 0.906 |
| Rv1354c | -0.132 | 0.999 | -0.537 | 0.767 | -0.405 | 0.816 |
| moeY | -0.164 | 0.999 | -0.329 | 0.659 | -0.165 | 0.816 |
| Rv1356c | -0.260 | NA | -1.433 | NA | -1.173 | NA |
| Rv1357c | -0.159 | NA | -1.617 | NA | -1.457 | NA |
| Rv1358 | -0.079 | 0.999 | -0.462 | 0.859 | -0.384 | 0.854 |
| Rv1359 | 0.059 | NA | -0.054 | 0.960 | -0.114 | 0.896 |
| Rv1360 | 0.082 | 0.999 | 0.027 | 0.951 | -0.055 | 0.879 |
| PPE19 | 0.032 | 0.999 | -0.197 | 0.826 | -0.229 | 0.816 |
| Rv1362c | 0.008 | 0.999 | -0.058 | 0.951 | -0.066 | 0.936 |
| Rv1363c | 0.024 | 0.999 | -0.411 | 0.832 | -0.436 | 0.816 |
| Rv1364c | 0.011 | 0.999 | -0.122 | 0.857 | -0.133 | 0.819 |
| rsfA | 0.052 | 0.999 | 0.036 | 0.972 | -0.016 | 0.986 |
| Rv1366 | -0.030 | 0.999 | -0.385 | 0.854 | -0.355 | 0.835 |
| Rv1366A | -0.022 | 0.999 | -0.063 | 0.948 | -0.041 | 0.955 |
| Rv1367c | 0.090 | 0.999 | -0.123 | 0.883 | -0.213 | 0.816 |
| lprF | 0.127 | 0.999 | 0.112 | 0.831 | -0.016 | 0.956 |
| Rv1371 | -0.259 | NA | -1.634 | NA | -1.375 | NA |
| Rv1372 | -0.100 | 0.999 | -0.261 | 0.883 | -0.161 | 0.909 |
| Rv1373 | -0.024 | 0.999 | -0.262 | 0.826 | -0.238 | 0.816 |
| Rv1374c | -0.060 | NA | -0.452 | 0.831 | -0.392 | 0.816 |
| Rv1375 | -0.274 | 0.999 | -0.046 | 0.946 | 0.228 | 0.816 |
| Rv1376 | -0.043 | 0.999 | 0.137 | 0.820 | 0.179 | 0.816 |
| Rv1377c | -0.004 | 0.999 | -0.019 | 0.978 | -0.016 | 0.985 |
| Rv1378c | -0.151 | 0.999 | -0.083 | 0.948 | 0.068 | 0.946 |
| pyrR | 0.045 | 0.999 | 0.259 | 0.848 | 0.214 | 0.836 |
| pyrB | 0.108 | 0.999 | -0.128 | 0.883 | -0.236 | 0.816 |
| pyrC | 0.129 | 0.999 | 0.209 | 0.858 | 0.080 | 0.924 |
| Rv1382 | 0.216 | NA | 0.241 | 0.879 | 0.025 | 0.984 |
| carA | 0.115 | 0.999 | 0.289 | 0.812 | 0.174 | 0.816 |
| carB | 0.111 | 0.999 | 0.075 | 0.895 | -0.035 | 0.942 |
| pyrF | 0.246 | NA | 0.646 | 0.410 | 0.399 | 0.816 |
| PE15 | -0.309 | 0.999 | 0.386 | 0.854 | 0.695 | 0.816 |
| PPE20 | -0.166 | 0.999 | -0.319 | 0.564 | -0.153 | 0.816 |
| mihF | -0.174 | 0.999 | -0.197 | 0.826 | -0.023 | 0.955 |
| gmk | -0.236 | 0.999 | 0.421 | 0.832 | 0.657 | 0.816 |
| rpoZ | 0.096 | 0.999 | -0.431 | 0.820 | -0.528 | 0.816 |
| dfp | -0.058 | 0.999 | 0.035 | 0.962 | 0.094 | 0.884 |
| metK | -0.126 | 0.999 | 0.002 | 0.995 | 0.128 | 0.820 |
| Rv1393c | 0.020 | 0.999 | -0.108 | 0.848 | -0.128 | 0.816 |
| cyp132 | 0.036 | 0.999 | -0.229 | 0.848 | -0.265 | 0.816 |
| Rv1395 | -0.067 | NA | -0.003 | 0.995 | 0.063 | 0.936 |
| PE_PGRS25 | -0.073 | 0.999 | -0.009 | 0.993 | 0.064 | 0.951 |
| vapC10 | 0.040 | 0.999 | -0.349 | 0.659 | -0.389 | 0.816 |
| vapB10 | -0.063 | 0.999 | 0.147 | 0.887 | 0.210 | 0.835 |
| nlhH | 0.099 | 0.999 | 0.057 | 0.948 | -0.042 | 0.953 |
| lipI | -0.139 | 0.999 | 0.475 | 0.831 | 0.614 | 0.816 |
| Rv1401 | 0.172 | 0.999 | 0.777 | 0.483 | 0.604 | 0.816 |
| priA | 0.037 | 0.999 | 0.455 | 0.831 | 0.417 | 0.816 |
| Rv1403c | -0.510 | NA | -0.668 | 0.826 | -0.158 | 0.924 |
| Rv1404 | -0.028 | 0.999 | -0.173 | 0.843 | -0.145 | 0.822 |
| Rv1405c | -0.790 | 0.001 | -0.833 | 0.000 | -0.043 | 0.941 |
| fmt | -0.151 | 0.999 | 0.052 | 0.962 | 0.203 | 0.837 |
| fmu | -0.221 | 0.999 | 0.311 | 0.859 | 0.532 | 0.816 |
| rpe | -0.036 | 0.999 | 0.124 | 0.862 | 0.160 | 0.817 |
| ribG | 0.014 | 0.999 | 0.232 | 0.853 | 0.218 | 0.834 |
| Rv1410c | 0.009 | 0.999 | 0.121 | 0.913 | 0.112 | 0.896 |
| lprG | 0.094 | 0.999 | 0.003 | 0.995 | -0.091 | 0.913 |
| ribC | -0.093 | 0.999 | -0.024 | 0.982 | 0.069 | 0.950 |
| Rv1413 | 0.026 | 0.999 | 0.082 | 0.931 | 0.056 | 0.942 |
| Rv1414 | -0.044 | NA | 0.059 | 0.959 | 0.103 | 0.909 |
| ribA2 | -0.101 | 0.999 | 0.077 | 0.922 | 0.178 | 0.819 |
| ribH | 0.089 | 0.999 | 0.017 | 0.975 | -0.072 | 0.896 |
| Rv1417 | 0.004 | 0.999 | 0.008 | 0.987 | 0.004 | 0.994 |
| lprH | -0.123 | 0.999 | 0.174 | 0.879 | 0.297 | 0.816 |
| Rv1419 | -0.167 | 0.999 | -0.261 | 0.701 | -0.094 | 0.846 |
| uvrC | -0.054 | 0.999 | -0.046 | 0.934 | 0.007 | 0.988 |
| Rv1421 | -0.034 | 0.999 | -0.067 | 0.942 | -0.033 | 0.962 |
| Rv1422 | -0.053 | 0.999 | 0.205 | 0.860 | 0.258 | 0.817 |
| whiA | -0.006 | 0.999 | 0.078 | 0.895 | 0.084 | 0.868 |
| Rv1424c | -0.024 | 0.999 | -0.110 | 0.879 | -0.086 | 0.875 |
| Rv1425 | -0.043 | 0.999 | 0.071 | 0.881 | 0.114 | 0.817 |
| lipO | 0.154 | 0.999 | 0.096 | 0.859 | -0.059 | 0.884 |
| fadD12 | 0.162 | 0.999 | -0.009 | 0.987 | -0.171 | 0.816 |
| Rv1428c | -0.038 | 0.999 | -0.134 | 0.868 | -0.096 | 0.868 |
| Rv1429 | -0.047 | 0.999 | -0.219 | 0.879 | -0.173 | 0.877 |
| PE16 | 0.113 | 0.999 | -0.045 | 0.962 | -0.158 | 0.852 |
| Rv1431 | -0.006 | 0.999 | -0.148 | 0.831 | -0.142 | 0.816 |
| Rv1432 | -0.157 | NA | -0.043 | 0.966 | 0.114 | 0.891 |
| Rv1433 | 0.024 | 0.999 | -0.078 | 0.895 | -0.102 | 0.852 |
| Rv1434 | -0.007 | NA | 0.073 | 0.948 | 0.080 | 0.930 |
| Rv1435c | 0.083 | 0.999 | 0.740 | 0.826 | 0.657 | 0.816 |
| gap | 0.073 | 0.999 | -0.303 | 0.820 | -0.376 | 0.816 |
| pgk | 0.159 | 0.999 | 0.071 | 0.927 | -0.088 | 0.891 |
| tpi | 0.133 | 0.999 | -0.099 | 0.895 | -0.232 | 0.816 |
| Rv1439c | -0.076 | 0.999 | -0.199 | 0.883 | -0.123 | 0.909 |
| secG | -0.150 | 0.999 | -0.170 | 0.848 | -0.020 | 0.973 |
| PE_PGRS26 | -0.049 | 0.999 | -0.150 | 0.859 | -0.101 | 0.868 |
| bisC | -0.077 | 0.999 | 0.121 | 0.913 | 0.198 | 0.837 |
| Rv1443c | -0.010 | 0.999 | -0.183 | 0.831 | -0.173 | 0.816 |
| Rv1444c | 0.071 | 0.999 | -0.129 | 0.879 | -0.200 | 0.816 |
| devB | -0.047 | 0.999 | 0.362 | 0.826 | 0.409 | 0.816 |
| opcA | 0.042 | 0.999 | 0.116 | 0.922 | 0.074 | 0.936 |
| zwf2 | 0.091 | 0.999 | -0.047 | 0.961 | -0.138 | 0.866 |
| tal | 0.122 | 0.999 | 0.027 | 0.962 | -0.095 | 0.846 |
| tkt | 0.062 | 0.999 | 0.036 | 0.938 | -0.026 | 0.942 |
| PE_PGRS27 | -0.049 | NA | -0.455 | 0.866 | -0.406 | 0.852 |
| Rv1451 | 0.004 | 0.999 | 0.238 | 0.831 | 0.233 | 0.816 |
| PE_PGRS28 | 0.015 | NA | -0.293 | 0.879 | -0.307 | 0.852 |
| Rv1453 | 0.075 | 0.999 | -0.348 | 0.866 | -0.423 | 0.819 |
| qor | -0.096 | 0.999 | 0.418 | 0.831 | 0.515 | 0.816 |
| Rv1455 | 0.061 | NA | 0.470 | 0.701 | 0.410 | 0.816 |
| Rv1456c | -0.144 | 0.999 | 0.521 | 0.826 | 0.665 | 0.816 |
| Rv1457c | 0.017 | 0.999 | 0.209 | 0.879 | 0.192 | 0.868 |
| Rv1458c | 0.159 | 0.999 | -0.132 | 0.848 | -0.291 | 0.816 |
| Rv1459c | -0.011 | 0.999 | 0.249 | 0.859 | 0.260 | 0.825 |
| Rv1460 | -0.173 | 0.999 | -0.321 | 0.716 | -0.149 | 0.819 |
| Rv1461 | -0.083 | 0.999 | -0.316 | 0.772 | -0.234 | 0.816 |
| Rv1462 | -0.160 | 0.999 | -0.065 | 0.925 | 0.095 | 0.868 |
| Rv1463 | -0.236 | 0.999 | -0.114 | 0.922 | 0.122 | 0.896 |
| csd | -0.140 | 0.999 | -0.191 | 0.678 | -0.052 | 0.868 |
| Rv1465 | -0.224 | 0.999 | -0.284 | 0.848 | -0.060 | 0.951 |
| Rv1466 | -0.112 | 0.999 | -0.249 | 0.820 | -0.138 | 0.819 |
| fadE15 | -0.091 | 0.999 | -0.288 | 0.826 | -0.197 | 0.817 |
| PE_PGRS29 | -0.232 | 0.999 | 0.065 | 0.949 | 0.297 | 0.816 |
| ctpD | 0.067 | 0.999 | 0.106 | 0.915 | 0.039 | 0.958 |
| trxA | -0.070 | 0.999 | -0.099 | 0.925 | -0.029 | 0.972 |
| trxB1 | 0.062 | 0.999 | 0.043 | 0.960 | -0.019 | 0.977 |
| echA12 | 0.123 | 0.999 | -0.358 | 0.804 | -0.481 | 0.816 |
| Rv1473 | 0.015 | 0.999 | 0.275 | 0.856 | 0.260 | 0.835 |
| Rv1473A | 0.468 | NA | -0.215 | 0.908 | -0.683 | 0.816 |
| Rv1474c | 0.038 | 0.999 | -0.042 | 0.970 | -0.081 | 0.932 |
| acn | 0.030 | 0.999 | -0.105 | 0.883 | -0.135 | 0.837 |
| Rv1476 | 0.110 | 0.999 | 0.089 | 0.922 | -0.021 | 0.977 |
| ripA | 0.201 | 0.999 | 0.202 | 0.826 | 0.001 | 0.998 |
| Rv1478 | 0.124 | 0.999 | 0.369 | 0.660 | 0.245 | 0.816 |
| moxR1 | 0.029 | 0.999 | 0.023 | 0.951 | -0.006 | 0.984 |
| Rv1480 | 0.045 | 0.999 | -0.075 | 0.961 | -0.120 | 0.922 |
| Rv1481 | 0.001 | 0.999 | 0.075 | 0.862 | 0.073 | 0.837 |
| Rv1482c | 0.050 | NA | -0.434 | 0.843 | -0.484 | 0.816 |
| fabG1 | 0.006 | 0.999 | -0.200 | 0.927 | -0.206 | 0.905 |
| inhA | -0.127 | 0.999 | 0.319 | 0.854 | 0.446 | 0.816 |
| hemZ | -0.006 | 0.999 | 0.027 | 0.971 | 0.033 | 0.957 |
| Rv1486c | -0.022 | 0.999 | 0.030 | 0.973 | 0.053 | 0.944 |
| Rv1487 | -0.065 | 0.999 | -0.043 | 0.974 | 0.022 | 0.988 |
| Rv1488 | -0.043 | 0.999 | -0.095 | 0.922 | -0.052 | 0.942 |
| Rv1489 | 0.013 | 0.999 | 0.070 | 0.923 | 0.057 | 0.924 |
| Rv1489A | -0.128 | 0.999 | -0.150 | 0.879 | -0.022 | 0.975 |
| Rv1490 | -0.087 | 0.999 | -0.288 | 0.826 | -0.201 | 0.819 |
| Rv1491c | 0.053 | 0.999 | -0.315 | 0.832 | -0.368 | 0.816 |
| mutA | -0.073 | 0.999 | -0.074 | 0.922 | -0.001 | 0.999 |
| mutB | 0.013 | 0.999 | -0.108 | 0.832 | -0.121 | 0.816 |
| mazE4 | 0.015 | 0.999 | -0.187 | 0.854 | -0.202 | 0.819 |
| mazF4 | 0.032 | NA | -0.349 | 0.826 | -0.380 | 0.816 |
| Rv1496 | -0.059 | 0.999 | 0.068 | 0.931 | 0.126 | 0.854 |
| lipL | 0.011 | 0.999 | 0.278 | 0.820 | 0.267 | 0.816 |
| Rv1498A | -0.054 | 0.999 | -0.302 | 0.717 | -0.248 | 0.816 |
| Rv1498c | 0.089 | 0.999 | -0.206 | 0.834 | -0.295 | 0.816 |
| Rv1499 | 0.242 | NA | 0.059 | 0.972 | -0.183 | 0.899 |
| Rv1500 | 0.106 | 0.999 | -0.275 | 0.879 | -0.381 | 0.819 |
| Rv1501 | 0.118 | 0.999 | -0.220 | 0.879 | -0.337 | 0.816 |
| Rv1502 | 0.193 | 0.999 | -0.050 | 0.975 | -0.244 | 0.884 |
| Rv1505c | 0.123 | 0.999 | -0.145 | 0.895 | -0.268 | 0.819 |
| Rv1506c | 0.082 | 0.999 | -0.349 | 0.848 | -0.431 | 0.816 |
| Rv1507A | -0.111 | NA | -0.561 | 0.832 | -0.450 | 0.821 |

| | | | | | | |
|---|---|---|---|---|---|---|
| ftsQ | 0.158 | 0.999 | -0.032 | 0.958 | -0.189 | 0.816 |
| murC | 0.096 | 0.999 | 0.040 | 0.958 | -0.055 | 0.930 |
| murG | -0.112 | 0.999 | 0.707 | 0.831 | 0.818 | 0.816 |
| ftsW | -0.046 | 0.999 | 0.210 | 0.879 | 0.255 | 0.836 |
| murD | 0.012 | 0.999 | 0.164 | 0.879 | 0.152 | 0.857 |
| murX | -0.142 | 0.999 | -0.038 | 0.974 | 0.104 | 0.932 |
| murF | -0.081 | 0.999 | -0.189 | 0.843 | -0.107 | 0.865 |
| murE | -0.194 | 0.999 | 0.219 | 0.879 | 0.413 | 0.816 |
| Rv2159c | -0.204 | 0.999 | -0.155 | 0.859 | 0.049 | 0.936 |
| Rv2161c | -0.060 | 0.999 | 0.294 | 0.866 | 0.354 | 0.819 |
| PE_PGRS38 | -0.153 | 0.999 | -0.155 | 0.879 | -0.002 | 0.998 |
| pbpB | 0.087 | 0.999 | -0.148 | 0.843 | -0.235 | 0.816 |
| Rv2164c | -0.180 | 0.999 | -0.021 | 0.970 | 0.160 | 0.816 |
| Rv2165c | 0.024 | 0.999 | 0.088 | 0.848 | 0.064 | 0.854 |
| Rv2166c | 0.019 | 0.999 | -0.038 | 0.973 | -0.058 | 0.955 |
| Rv2169c | -0.049 | 0.999 | 0.256 | 0.832 | 0.305 | 0.816 |
| Rv2170 | -0.104 | NA | 0.212 | 0.859 | 0.316 | 0.816 |
| lppM | -0.011 | 0.999 | -0.028 | 0.973 | -0.017 | 0.985 |
| Rv2172c | -0.028 | 0.999 | 0.073 | 0.832 | 0.101 | 0.816 |
| idsA2 | 0.017 | 0.999 | 0.052 | 0.922 | 0.035 | 0.932 |
| mptA | 0.089 | 0.999 | 0.228 | 0.831 | 0.139 | 0.836 |
| Rv2175c | 0.145 | 0.999 | 0.217 | 0.832 | 0.071 | 0.909 |
| pknL | 0.049 | 0.999 | 0.191 | 0.843 | 0.143 | 0.836 |
| Rv2177c | -0.001 | 0.999 | -0.208 | 0.820 | -0.207 | 0.816 |
| aroG | 0.040 | 0.999 | 0.133 | 0.859 | 0.093 | 0.868 |
| Rv2179c | 0.038 | NA | -0.074 | 0.927 | -0.112 | 0.868 |
| Rv2180c | -0.029 | NA | -0.117 | 0.879 | -0.089 | 0.891 |
| Rv2181 | -0.044 | 0.999 | 0.305 | 0.848 | 0.349 | 0.816 |
| Rv2182c | -0.065 | 0.999 | -0.102 | 0.879 | -0.037 | 0.942 |
| Rv2183c | -0.023 | 0.999 | -0.026 | 0.975 | -0.003 | 0.997 |
| Rv2184c | -0.011 | 0.999 | 0.340 | 0.660 | 0.351 | 0.816 |
| TB16.3 | 0.135 | 0.999 | -0.181 | 0.848 | -0.316 | 0.816 |
| Rv2186c | -0.081 | 0.999 | 0.075 | 0.936 | 0.156 | 0.852 |
| fadD15 | 0.054 | 0.999 | 0.052 | 0.883 | -0.002 | 0.994 |
| pimB | -0.173 | 0.999 | 0.093 | 0.926 | 0.266 | 0.816 |
| Rv2189c | 0.167 | 0.999 | 0.517 | 0.820 | 0.350 | 0.816 |
| Rv2190c | 0.140 | 0.999 | 0.017 | 0.977 | -0.123 | 0.862 |
| Rv2191 | 0.047 | 0.999 | 0.066 | 0.927 | 0.020 | 0.973 |
| trpD | 0.055 | 0.999 | 0.257 | 0.826 | 0.202 | 0.816 |
| ctaE | 0.061 | 0.999 | -0.022 | 0.970 | -0.084 | 0.865 |
| qcrC | 0.017 | 0.999 | -0.280 | 0.822 | -0.297 | 0.816 |
| qcrA | 0.061 | 0.999 | -0.072 | 0.895 | -0.133 | 0.819 |
| qcrB | 0.058 | 0.999 | 0.079 | 0.946 | 0.020 | 0.984 |
| Rv2197c | -0.029 | 0.999 | 0.203 | 0.848 | 0.232 | 0.816 |
| mmpS3 | -0.005 | 0.999 | -0.132 | 0.879 | -0.127 | 0.866 |
| Rv2199c | -0.053 | 0.999 | 0.298 | 0.832 | 0.351 | 0.816 |
| ctaC | 0.081 | 0.999 | -0.137 | 0.856 | -0.218 | 0.816 |
| asnB | -0.004 | 0.999 | 0.125 | 0.832 | 0.130 | 0.816 |
| adoK | -0.025 | 0.999 | 0.134 | 0.879 | 0.159 | 0.837 |
| Rv2203 | 0.216 | 0.999 | -0.003 | 0.995 | -0.219 | 0.835 |
| Rv2204c | -0.052 | 0.999 | -0.433 | 0.859 | -0.382 | 0.838 |
| Rv2205c | -0.143 | 0.999 | 0.361 | 0.832 | 0.505 | 0.816 |
| Rv2206 | 0.007 | 0.999 | 0.578 | 0.826 | 0.572 | 0.816 |
| cobT | 0.088 | 0.999 | 0.443 | 0.831 | 0.355 | 0.817 |
| cobS | 0.059 | 0.999 | 0.671 | 0.826 | 0.612 | 0.816 |
| Rv2209 | -0.178 | 0.999 | 0.404 | 0.832 | 0.582 | 0.816 |
| ilvE | 0.066 | 0.999 | 0.251 | 0.826 | 0.185 | 0.816 |
| gcvT | 0.085 | 0.999 | 0.154 | 0.829 | 0.069 | 0.866 |
| Rv2212 | 0.050 | 0.999 | -0.005 | 0.988 | -0.054 | 0.879 |
| pepB | 0.076 | 0.999 | 0.238 | 0.820 | 0.162 | 0.816 |
| ephD | 0.099 | 0.999 | 0.145 | 0.826 | 0.046 | 0.882 |
| dlaT | -0.046 | 0.999 | 0.077 | 0.879 | 0.123 | 0.816 |
| Rv2216 | 0.006 | 0.999 | 0.251 | 0.834 | 0.245 | 0.816 |
| lipB | -0.147 | 0.999 | 0.186 | 0.848 | 0.333 | 0.816 |
| lipA | -0.060 | 0.999 | -0.020 | 0.961 | 0.040 | 0.899 |
| Rv2219 | 0.033 | 0.999 | -0.180 | 0.826 | -0.213 | 0.816 |
| Rv2219A | -0.115 | 0.999 | 0.340 | 0.826 | 0.455 | 0.816 |
| glnA1 | 0.056 | 0.999 | -0.162 | 0.886 | -0.218 | 0.837 |
| glnE | 0.068 | 0.999 | 0.300 | 0.826 | 0.232 | 0.816 |
| glnA2 | 0.031 | 0.999 | 0.230 | 0.829 | 0.199 | 0.816 |
| Rv2223c | 0.073 | 0.999 | 0.187 | 0.826 | 0.113 | 0.819 |
| caeA | -0.019 | 0.999 | 0.065 | 0.895 | 0.083 | 0.852 |
| Rv2224d | -0.025 | NA | 0.390 | 0.826 | 0.415 | 0.816 |
| panB | 0.365 | 0.999 | 0.683 | 0.022 | 0.318 | 0.816 |
| Rv2226 | 0.466 | 0.058 | 0.533 | 0.004 | 0.067 | 0.881 |
| Rv2227 | 0.098 | 0.999 | 0.027 | 0.974 | -0.071 | 0.932 |
| Rv2228c | 0.102 | 0.999 | 0.063 | 0.895 | -0.039 | 0.922 |
| Rv2229c | 0.202 | 0.999 | 0.024 | 0.974 | -0.178 | 0.835 |
| Rv2230c | 0.105 | 0.999 | 0.113 | 0.895 | 0.008 | 0.991 |
| vapC16 | 0.028 | NA | -0.270 | 0.885 | -0.298 | 0.859 |
| vapB16 | -0.176 | NA | 0.267 | 0.879 | 0.443 | 0.816 |
| cobC | 0.121 | 0.999 | -0.050 | 0.962 | -0.171 | 0.862 |
| ptkA | 0.043 | 0.999 | 0.177 | 0.859 | 0.133 | 0.865 |
| ptpA | -0.162 | 0.999 | -0.250 | 0.848 | -0.087 | 0.924 |
| Rv2235 | -0.081 | 0.999 | 0.136 | 0.879 | 0.217 | 0.816 |
| cobD | -0.109 | NA | 0.564 | 0.767 | 0.673 | 0.816 |
| Rv2237 | -0.023 | 0.999 | 0.179 | 0.848 | 0.203 | 0.816 |
| Rv2237A | 0.033 | 0.999 | 0.114 | 0.886 | 0.081 | 0.901 |
| ahpE | 0.117 | 0.999 | -0.014 | 0.982 | -0.131 | 0.854 |
| Rv2239c | -0.249 | 0.999 | -0.132 | 0.926 | 0.118 | 0.918 |
| Rv2240c | 0.075 | 0.999 | 0.415 | 0.527 | 0.340 | 0.816 |
| aceE | -0.007 | 0.999 | 0.026 | 0.948 | 0.033 | 0.918 |
| Rv2242 | 0.101 | 0.999 | -0.165 | 0.879 | -0.266 | 0.816 |
| fabD | 0.077 | 0.999 | 0.194 | 0.848 | 0.117 | 0.865 |
| acpM | 0.206 | 0.999 | -0.185 | 0.879 | -0.391 | 0.816 |
| kasA | 0.018 | 0.999 | 0.143 | 0.895 | 0.125 | 0.891 |
| kasB | 0.060 | 0.999 | 0.002 | 0.995 | -0.057 | 0.904 |
| accD6 | 0.156 | 0.999 | 0.089 | 0.912 | -0.068 | 0.916 |
| Rv2248 | 0.027 | 0.999 | 0.434 | 0.176 | 0.408 | 0.816 |
| glpD1 | -0.042 | NA | 0.231 | 0.854 | 0.274 | 0.816 |
| Rv2250c | 0.227 | NA | 0.017 | 0.987 | -0.211 | 0.855 |
| Rv2251 | 0.015 | 0.999 | 0.219 | 0.826 | 0.204 | 0.816 |
| Rv2252 | -0.012 | NA | -0.199 | 0.848 | -0.186 | 0.819 |
| Rv2253 | 0.081 | NA | -0.551 | 0.831 | -0.632 | 0.816 |
| Rv2254c | 0.077 | NA | 0.239 | 0.848 | 0.161 | 0.854 |
| Rv2256a | 0.044 | 0.999 | 0.089 | 0.962 | 0.045 | 0.977 |
| Rv2256c | 0.224 | 0.999 | 0.268 | 0.826 | 0.044 | 0.942 |
| Rv2257c | 0.156 | 0.999 | 0.152 | 0.826 | -0.003 | 0.994 |
| Rv2258c | 0.016 | 0.999 | 0.068 | 0.879 | 0.052 | 0.886 |
| mscR | -0.018 | 0.999 | 0.061 | 0.920 | 0.078 | 0.868 |
| Rv2260 | 0.147 | 0.999 | -0.043 | 0.946 | -0.191 | 0.816 |
| Rv2263 | -0.146 | 0.999 | 0.206 | 0.866 | 0.352 | 0.816 |
| Rv2264c | -0.119 | 0.999 | -0.058 | 0.922 | 0.061 | 0.896 |
| Rv2265 | -0.070 | NA | -0.030 | 0.975 | 0.041 | 0.967 |
| cyp124 | -0.159 | 0.999 | -0.015 | 0.974 | 0.144 | 0.816 |
| Rv2267c | -0.055 | NA | -0.875 | 0.772 | -0.820 | 0.816 |
| cyp128 | -0.046 | NA | 0.003 | 0.996 | 0.049 | 0.962 |
| Rv2269c | -0.402 | NA | -1.020 | 0.848 | -0.618 | 0.862 |
| lppN | 0.010 | NA | -0.015 | 0.988 | -0.026 | 0.983 |
| Rv2271 | 0.153 | 0.999 | -0.262 | 0.883 | -0.414 | 0.819 |
| Rv2272 | 0.061 | 0.999 | 0.153 | 0.879 | 0.092 | 0.904 |
| Rv2273 | -0.081 | NA | 0.193 | 0.848 | 0.274 | 0.816 |
| mazE8 | -0.012 | NA | -0.338 | 0.848 | -0.326 | 0.822 |
| mazF8 | -0.021 | NA | -0.085 | 0.940 | -0.064 | 0.942 |
| Rv2275 | -0.099 | 0.999 | -0.207 | 0.826 | -0.108 | 0.851 |
| cyp121 | -0.007 | 0.999 | -0.367 | 0.859 | -0.361 | 0.835 |
| Rv2277c | -0.194 | NA | -1.131 | 0.826 | -0.937 | 0.816 |
| Rv2280 | -0.097 | 0.999 | 0.060 | 0.922 | 0.157 | 0.816 |
| pitB | -0.104 | 0.999 | -0.371 | 0.820 | -0.267 | 0.816 |
| Rv2282c | -0.187 | NA | 0.000 | 1.000 | 0.187 | 0.835 |
| Rv2283 | -1.183 | NA | -2.098 | 0.613 | -0.915 | 0.817 |
| lipM | -0.096 | 0.999 | 0.064 | 0.931 | 0.160 | 0.824 |
| Rv2285 | -0.024 | 0.999 | 0.076 | 0.879 | 0.101 | 0.835 |
| Rv2286c | 0.076 | 0.999 | 0.215 | 0.832 | 0.139 | 0.838 |
| yjcE | 0.011 | 0.999 | 0.190 | 0.832 | 0.179 | 0.816 |
| Rv2288 | -0.117 | 0.999 | -0.020 | 0.976 | 0.097 | 0.896 |
| cdh | 0.041 | 0.999 | -0.110 | 0.879 | -0.151 | 0.817 |
| lppO | 0.003 | 0.999 | -0.039 | 0.967 | -0.042 | 0.954 |
| sseB | -0.154 | 0.999 | 0.208 | 0.820 | 0.361 | 0.816 |
| Rv2292c | 0.101 | NA | -0.059 | 0.951 | -0.159 | 0.856 |
| Rv2293c | 0.164 | 0.999 | 0.012 | 0.985 | -0.152 | 0.837 |
| Rv2294 | -0.049 | 0.999 | 0.029 | 0.970 | 0.077 | 0.897 |
| Rv2295 | 0.039 | 0.999 | 0.102 | 0.879 | 0.062 | 0.897 |
| Rv2296 | -0.005 | 0.999 | -0.061 | 0.948 | -0.056 | 0.942 |
| Rv2297 | 0.050 | 0.999 | -0.215 | 0.917 | -0.265 | 0.868 |
| Rv2298 | 0.089 | 0.999 | -0.046 | 0.971 | -0.135 | 0.899 |
| htpG | 0.098 | 0.999 | -0.152 | 0.848 | -0.250 | 0.816 |
| Rv2300c | 0.051 | 0.999 | -0.129 | 0.859 | -0.180 | 0.816 |
| cut2 | 0.014 | 0.999 | 0.211 | 0.859 | 0.197 | 0.838 |
| Rv2302 | 0.139 | 0.999 | 0.362 | 0.820 | 0.223 | 0.816 |
| Rv2303c | 0.067 | 0.999 | -0.157 | 0.853 | -0.224 | 0.816 |
| Rv2304c | -0.017 | NA | -0.422 | 0.832 | -0.405 | 0.816 |
| Rv2305 | -0.009 | 0.999 | 0.188 | 0.832 | 0.197 | 0.816 |
| Rv2307A | 0.013 | NA | 0.117 | 0.918 | 0.104 | 0.904 |
| Rv2307B | -0.022 | NA | -0.160 | 0.886 | -0.138 | 0.884 |
| Rv2307c | 0.003 | 0.999 | 0.099 | 0.912 | 0.097 | 0.892 |
| Rv2307D | -0.192 | 0.999 | 0.315 | 0.820 | 0.507 | 0.816 |
| Rv2308 | -0.147 | 0.999 | -0.198 | 0.848 | -0.051 | 0.936 |
| Rv2308a | -0.065 | NA | -0.394 | 0.832 | -0.329 | 0.817 |
| Rv2309A | -0.241 | NA | -0.553 | 0.866 | -0.312 | 0.896 |
| Rv2310 | 0.242 | NA | 0.286 | 0.922 | 0.044 | 0.986 |
| Rv2311 | -0.084 | NA | -0.260 | 0.879 | -0.175 | 0.896 |
| Rv2312 | -0.044 | NA | -0.547 | 0.831 | -0.503 | 0.816 |
| Rv2313c | -0.003 | 0.999 | 0.094 | 0.879 | 0.097 | 0.857 |
| Rv2314c | -0.016 | 0.999 | 0.094 | 0.922 | 0.109 | 0.884 |
| Rv2315c | 0.010 | 0.999 | -0.005 | 0.991 | -0.015 | 0.977 |
| uspA | -0.208 | NA | 0.090 | 0.948 | 0.297 | 0.830 |
| uspB | 0.042 | NA | -0.555 | 0.832 | -0.597 | 0.816 |
| uspC | -0.126 | 0.999 | 0.069 | 0.922 | 0.194 | 0.816 |
| Rv2319c | -0.065 | NA | -0.181 | 0.854 | -0.116 | 0.868 |
| rocE | -0.064 | 0.999 | 0.068 | 0.938 | 0.131 | 0.862 |
| Rv2323c | 0.058 | 0.999 | -0.105 | 0.879 | -0.163 | 0.817 |
| Rv2324 | -0.149 | 0.999 | -0.053 | 0.941 | 0.096 | 0.868 |
| Rv2325c | -0.018 | 0.999 | -0.032 | 0.958 | -0.014 | 0.977 |
| Rv2326c | 0.024 | 0.999 | 0.017 | 0.973 | -0.007 | 0.988 |

| | | | | | | |
|---|---|---|---|---|---|---|
| Rv2958c | 0.122 | 0.999 | -0.048 | 0.948 | -0.170 | 0.819 |
| Rv2959c | 0.265 | 0.999 | -0.124 | 0.931 | -0.388 | 0.816 |
| Rv2960c | 0.178 | 0.999 | -0.356 | 0.848 | -0.534 | 0.816 |
| Rv2961 | -0.252 | NA | -0.892 | 0.826 | -0.641 | 0.816 |
| Rv2962c | -0.146 | 0.999 | -0.114 | 0.868 | 0.032 | 0.947 |
| Rv2963 | -0.492 | 0.119 | -0.588 | 0.007 | -0.096 | 0.862 |
| purU | -0.229 | 0.999 | -0.662 | 0.820 | -0.433 | 0.816 |
| kdtB | 0.032 | 0.999 | -0.215 | 0.848 | -0.247 | 0.816 |
| Rv2966c | -0.003 | 0.999 | 0.206 | 0.848 | 0.208 | 0.819 |
| pca | 0.051 | 0.999 | 0.020 | 0.962 | -0.030 | 0.935 |
| Rv2968c | -0.057 | 0.999 | -0.136 | 0.879 | -0.078 | 0.897 |
| Rv2969c | -0.058 | 0.999 | 0.039 | 0.951 | 0.096 | 0.862 |
| Rv2970A | -0.003 | 0.999 | -0.098 | 0.879 | -0.095 | 0.865 |
| lipN | 0.071 | 0.999 | -0.167 | 0.834 | -0.237 | 0.816 |
| Rv2971 | 0.071 | 0.999 | -0.144 | 0.895 | -0.215 | 0.835 |
| Rv2972c | 0.099 | NA | -0.269 | 0.820 | -0.368 | 0.816 |
| recG | 0.049 | 0.999 | -0.086 | 0.886 | -0.135 | 0.819 |
| Rv2974c | 0.192 | 0.999 | 0.242 | 0.831 | 0.050 | 0.939 |
| Rv2975a | -0.039 | 0.999 | -0.103 | 0.908 | -0.064 | 0.930 |
| Rv2975c | -0.099 | 0.999 | -0.234 | 0.857 | -0.136 | 0.882 |
| ung | -0.085 | 0.999 | 0.083 | 0.951 | 0.167 | 0.882 |
| thiL | 0.114 | 0.999 | -0.034 | 0.970 | -0.148 | 0.855 |
| Rv2978c | -0.019 | 0.999 | -0.047 | 0.962 | -0.028 | 0.972 |
| Rv2979c | -0.149 | 0.999 | -0.386 | 0.728 | -0.237 | 0.816 |
| Rv2980 | 0.038 | 0.999 | 0.337 | 0.825 | 0.299 | 0.816 |
| ddlA | 0.041 | 0.999 | 0.107 | 0.878 | 0.066 | 0.892 |
| gpdA2 | 0.083 | 0.999 | 0.164 | 0.866 | 0.081 | 0.906 |
| Rv2983 | 0.021 | 0.999 | 0.216 | 0.859 | 0.195 | 0.838 |
| ppk1 | 0.099 | 0.999 | -0.218 | 0.826 | -0.317 | 0.816 |
| mutT1 | -0.030 | 0.999 | -0.220 | 0.879 | -0.190 | 0.868 |
| hupB | -0.182 | 0.999 | -0.046 | 0.947 | 0.136 | 0.835 |
| leuD | -0.031 | 0.999 | -0.067 | 0.895 | -0.036 | 0.932 |
| leuC | -0.049 | 0.999 | 0.058 | 0.927 | 0.107 | 0.846 |
| Rv2989 | -0.235 | 0.999 | 0.116 | 0.922 | 0.350 | 0.816 |
| Rv2990c | -0.034 | 0.999 | 0.209 | 0.832 | 0.243 | 0.816 |
| Rv2991 | -0.032 | 0.999 | -0.063 | 0.922 | -0.031 | 0.947 |
| gltS | 0.079 | 0.999 | -0.069 | 0.914 | -0.149 | 0.817 |
| Rv2993c | 0.015 | 0.999 | 0.013 | 0.982 | -0.002 | 0.997 |
| Rv2994 | -0.001 | 0.999 | 0.476 | 0.831 | 0.477 | 0.816 |
| leuB | 0.098 | 0.999 | 0.564 | 0.686 | 0.466 | 0.816 |
| serA1 | 0.085 | 0.999 | 0.290 | 0.831 | 0.205 | 0.819 |
| Rv2997 | 0.084 | NA | 0.452 | 0.820 | 0.368 | 0.816 |
| Rv2998 | 0.101 | 0.999 | -0.201 | 0.914 | -0.303 | 0.852 |
| lppY | 0.021 | NA | -0.010 | 0.989 | -0.031 | 0.970 |
| Rv3000 | -0.021 | NA | 0.311 | 0.848 | 0.332 | 0.817 |
| ilvC | -0.001 | 0.999 | 0.126 | 0.879 | 0.127 | 0.852 |
| ilvN | 0.053 | 0.999 | -0.074 | 0.947 | -0.127 | 0.884 |
| ilvB1 | -0.057 | 0.999 | 0.075 | 0.879 | 0.133 | 0.816 |
| cfp6 | 0.101 | 0.999 | 0.128 | 0.890 | 0.027 | 0.970 |
| Rv3005c | 0.080 | 0.999 | 0.360 | 0.802 | 0.280 | 0.816 |
| lppZ | 0.078 | 0.999 | 0.225 | 0.826 | 0.147 | 0.817 |
| Rv3007c | -0.176 | NA | -0.109 | 0.895 | 0.067 | 0.924 |
| Rv3008 | 0.037 | 0.999 | -0.081 | 0.879 | -0.118 | 0.816 |
| gatB | -0.065 | 0.999 | -0.069 | 0.897 | -0.004 | 0.994 |
| pfkA | -0.006 | 0.999 | -0.024 | 0.962 | -0.018 | 0.964 |
| gatA | -0.001 | 0.999 | 0.268 | 0.832 | 0.269 | 0.816 |
| gatC | -0.029 | 0.999 | -0.277 | 0.879 | -0.248 | 0.865 |
| Rv3013 | -0.091 | NA | -0.014 | 0.987 | 0.077 | 0.933 |
| ligA | 0.028 | 0.999 | 0.159 | 0.859 | 0.131 | 0.852 |
| Rv3015c | -0.098 | 0.999 | 0.373 | 0.832 | 0.470 | 0.816 |
| lpqA | -0.373 | 0.999 | -0.386 | 0.562 | -0.013 | 0.984 |
| esxQ | 0.110 | NA | -0.018 | 0.985 | -0.128 | 0.896 |
| PPE46 | -0.122 | NA | -0.947 | NA | -0.825 | NA |
| esxR | -0.060 | NA | -0.510 | 0.879 | -0.450 | 0.868 |
| esxS | -0.079 | NA | -1.084 | 0.854 | -1.005 | 0.835 |
| PE29 | -0.152 | NA | -0.868 | 0.848 | -0.716 | 0.837 |
| Rv3023c | -0.069 | NA | -0.329 | 0.879 | -0.260 | 0.871 |
| trmU | 0.025 | 0.999 | -0.106 | 0.879 | -0.131 | 0.836 |
| iscS | 0.010 | 0.999 | -0.053 | 0.931 | -0.063 | 0.897 |
| Rv3026c | 0.181 | NA | 0.386 | 0.588 | 0.205 | 0.816 |
| Rv3027c | 0.118 | 0.999 | -0.278 | 0.834 | -0.396 | 0.816 |
| fixB | 0.055 | 0.999 | 0.516 | 0.825 | 0.461 | 0.816 |
| fixA | 0.207 | 0.999 | 0.177 | 0.853 | -0.030 | 0.961 |
| Rv3030 | 0.240 | 0.999 | 0.043 | 0.946 | -0.197 | 0.816 |
| Rv3031 | 0.078 | 0.999 | 0.395 | 0.826 | 0.317 | 0.816 |
| Rv3032 | 0.224 | 0.999 | 0.391 | 0.359 | 0.167 | 0.816 |
| Rv3032A | 0.095 | NA | 0.088 | 0.922 | -0.008 | 0.992 |
| Rv3033 | 0.102 | 0.999 | -0.223 | 0.879 | -0.325 | 0.816 |
| Rv3034c | 0.026 | 0.999 | 0.099 | 0.886 | 0.073 | 0.896 |
| Rv3035 | -0.043 | 0.999 | 0.159 | 0.848 | 0.203 | 0.816 |
| TB22.2 | -0.022 | 0.999 | 0.014 | 0.964 | 0.036 | 0.892 |
| Rv3037c | -0.156 | 0.999 | 0.167 | 0.895 | 0.323 | 0.817 |
| Rv3038c | -0.057 | 0.999 | 0.143 | 0.879 | 0.200 | 0.817 |
| echA17 | -0.028 | 0.999 | 0.265 | 0.848 | 0.292 | 0.816 |
| Rv3040c | -0.019 | 0.999 | 0.112 | 0.935 | 0.131 | 0.903 |
| Rv3041c | 0.139 | 0.999 | 0.340 | 0.820 | 0.201 | 0.817 |
| serB2 | 0.053 | 0.999 | 0.227 | 0.832 | 0.174 | 0.822 |
| ctaD | 0.032 | 0.999 | -0.041 | 0.962 | -0.073 | 0.919 |
| fecB | 0.110 | 0.999 | 0.077 | 0.895 | -0.032 | 0.946 |
| adhC | 0.097 | 0.999 | -0.075 | 0.886 | -0.171 | 0.816 |
| Rv3046c | -0.009 | 0.999 | 0.449 | 0.826 | 0.458 | 0.816 |
| Rv3047c | 0.155 | 0.999 | -0.151 | 0.859 | -0.305 | 0.816 |
| nrdF2 | 0.132 | 0.999 | 0.037 | 0.951 | -0.095 | 0.857 |
| Rv3049c | 0.227 | 0.999 | 0.166 | 0.826 | -0.061 | 0.876 |
| Rv3050c | 0.059 | 0.999 | -0.114 | 0.879 | -0.173 | 0.819 |
| nrdE | 0.127 | 0.999 | -0.201 | 0.861 | -0.328 | 0.816 |
| nrdI | 0.138 | 0.999 | -0.285 | 0.826 | -0.424 | 0.816 |
| nrdH | 0.200 | 0.999 | 0.100 | 0.927 | -0.100 | 0.909 |
| Rv3054c | -0.546 | NA | -0.419 | 0.879 | 0.127 | 0.944 |
| Rv3055 | 0.049 | 0.999 | 0.112 | 0.895 | 0.062 | 0.931 |
| dinP | 0.030 | 0.999 | -0.228 | 0.859 | -0.258 | 0.819 |
| Rv3057c | 0.110 | 0.999 | 0.043 | 0.951 | -0.067 | 0.905 |
| Rv3058c | 0.266 | 0.999 | -0.231 | 0.858 | -0.496 | 0.816 |
| cyp136 | 0.057 | 0.999 | -0.128 | 0.886 | -0.185 | 0.835 |
| Rv3060c | -0.007 | 0.999 | -0.198 | 0.856 | -0.190 | 0.835 |
| fadE22 | -0.057 | 0.999 | -0.137 | 0.925 | -0.080 | 0.942 |
| ligB | 0.040 | 0.999 | 0.183 | 0.826 | 0.142 | 0.816 |
| cstA | 0.062 | 0.999 | -0.101 | 0.850 | -0.163 | 0.816 |
| Rv3064c | -0.107 | 0.999 | -0.074 | 0.922 | 0.032 | 0.955 |
| mmr | -0.142 | NA | -0.391 | 0.728 | -0.249 | 0.816 |
| Rv3066 | 0.004 | NA | -0.135 | 0.879 | -0.139 | 0.852 |
| Rv3067 | -0.058 | NA | -0.513 | 0.826 | -0.455 | 0.816 |
| pgmA | 0.007 | 0.999 | -0.080 | 0.922 | -0.087 | 0.892 |
| Rv3069 | -0.347 | NA | 0.005 | 0.995 | 0.352 | 0.830 |
| Rv3070 | 0.030 | NA | -0.335 | 0.832 | -0.365 | 0.816 |
| Rv3071 | 0.013 | 0.999 | -0.159 | 0.879 | -0.172 | 0.837 |
| Rv3072c | -0.118 | 0.999 | 0.072 | 0.942 | 0.190 | 0.836 |
| Rv3073c | 0.036 | 0.999 | -0.170 | 0.879 | -0.206 | 0.835 |
| Rv3074 | 0.070 | 0.999 | 0.115 | 0.866 | 0.045 | 0.930 |
| Rv3075c | -0.117 | 0.999 | 0.179 | 0.863 | 0.296 | 0.816 |
| Rv3076 | 0.098 | NA | -0.107 | 0.885 | -0.205 | 0.816 |
| Rv3077 | 0.133 | 0.999 | 0.006 | 0.987 | -0.128 | 0.816 |
| hab | 0.047 | 0.999 | -0.117 | 0.886 | -0.163 | 0.836 |
| Rv3079c | 0.035 | NA | -0.262 | 0.862 | -0.297 | 0.824 |
| pknK | -0.124 | 0.999 | 0.049 | 0.946 | 0.173 | 0.816 |
| Rv3081 | -0.041 | 0.999 | -0.087 | 0.912 | -0.046 | 0.942 |
| virS | -0.204 | NA | -0.156 | 0.931 | 0.048 | 0.973 |
| Rv3083 | 0.082 | 0.999 | -0.136 | 0.879 | -0.219 | 0.817 |
| lipR | 0.105 | 0.999 | 0.051 | 0.962 | -0.054 | 0.950 |
| Rv3085 | 0.137 | 0.999 | -0.250 | 0.848 | -0.387 | 0.816 |
| adhD | 0.018 | 0.999 | -0.043 | 0.958 | -0.061 | 0.925 |
| Rv3087 | -0.052 | 0.999 | 0.075 | 0.933 | 0.126 | 0.866 |
| tgs4 | -0.002 | 0.999 | -0.274 | 0.831 | -0.272 | 0.816 |
| fadD13 | -0.016 | 0.999 | -0.287 | 0.820 | -0.271 | 0.816 |
| Rv3090 | -0.046 | NA | -0.485 | 0.728 | -0.439 | 0.816 |
| Rv3091 | -0.127 | 0.999 | 0.186 | 0.848 | 0.312 | 0.816 |
| Rv3092c | 0.085 | 0.999 | 0.563 | 0.701 | 0.478 | 0.816 |
| Rv3093c | 0.255 | 0.999 | 0.826 | 0.054 | 0.571 | 0.816 |
| Rv3094c | 0.251 | 0.504 | 0.440 | 0.000 | 0.189 | 0.816 |
| Rv3095 | 0.226 | 0.999 | 0.369 | 0.826 | 0.143 | 0.871 |
| Rv3096 | 0.136 | 0.999 | 0.245 | 0.168 | 0.109 | 0.816 |
| lipY | -0.023 | NA | -0.093 | 0.927 | -0.070 | 0.932 |
| Rv3098A | -0.022 | 0.999 | -0.091 | 0.891 | -0.069 | 0.897 |
| Rv3098c | 0.148 | NA | 0.118 | 0.895 | -0.029 | 0.967 |
| Rv3099c | 0.078 | 0.999 | 0.061 | 0.965 | -0.017 | 0.988 |
| smpB | 0.007 | 0.999 | -0.145 | 0.879 | -0.153 | 0.854 |
| ftsX | 0.125 | 0.999 | 0.006 | 0.989 | -0.119 | 0.835 |
| ftsE | 0.031 | 0.999 | -0.176 | 0.858 | -0.206 | 0.817 |
| Rv3103c | 0.081 | 0.999 | -0.041 | 0.970 | -0.122 | 0.892 |
| Rv3104c | 0.048 | 0.999 | 0.296 | 0.772 | 0.247 | 0.816 |
| prfB | 0.047 | 0.999 | -0.223 | 0.826 | -0.270 | 0.816 |
| fprA | 0.109 | 0.999 | 0.227 | 0.695 | 0.118 | 0.816 |
| agpS | -0.040 | 0.999 | -0.018 | 0.970 | 0.022 | 0.951 |
| Rv3108 | 0.000 | NA | -0.434 | 0.886 | -0.434 | 0.868 |
| moaA1 | 0.132 | 0.999 | -0.544 | 0.854 | -0.675 | 0.816 |
| moaB1 | 0.138 | NA | -0.443 | 0.848 | -0.581 | 0.816 |
| moaC1 | 0.145 | NA | -0.704 | 0.831 | -0.849 | 0.816 |
| moaD1 | -0.100 | NA | -1.245 | 0.831 | -1.145 | 0.816 |
| Rv3113 | 0.072 | NA | -0.586 | 0.832 | -0.657 | 0.816 |
| Rv3114 | -0.045 | NA | -1.177 | 0.826 | -1.132 | 0.816 |
| Rv3115 | -0.041 | NA | -0.299 | 0.879 | -0.258 | 0.862 |
| moeB2 | 0.021 | 0.999 | -0.331 | 0.801 | -0.352 | 0.816 |
| cysA3 | 0.025 | 0.999 | -0.234 | 0.820 | -0.259 | 0.816 |
| Rv3118 | 0.096 | 0.999 | 0.615 | 0.826 | 0.519 | 0.816 |
| moaE1 | 0.270 | 0.999 | -0.054 | 0.959 | -0.324 | 0.816 |
| Rv3120 | -0.007 | 0.999 | -0.101 | 0.869 | -0.094 | 0.846 |
| cyp141 | 0.096 | 0.999 | -0.258 | 0.820 | -0.355 | 0.816 |
| Rv3122 | -0.010 | 0.999 | -0.274 | 0.918 | -0.264 | 0.896 |
| Rv3123 | -0.058 | NA | 0.402 | 0.826 | 0.459 | 0.816 |
| moaR1 | 0.103 | 0.999 | -0.620 | 0.848 | -0.723 | 0.816 |
| PPE49 | -0.218 | 0.999 | -0.592 | 0.846 | -0.374 | 0.855 |
| Rv3126c | -0.595 | NA | -1.182 | 0.029 | -0.587 | 0.816 |
| Rv3127 | -0.647 | 0.298 | -0.959 | 0.001 | -0.312 | 0.816 |
| tgs1 | -0.380 | 0.999 | -0.731 | 0.001 | -0.352 | 0.816 |
| Rv3131 | -0.638 | 0.306 | -0.786 | 0.022 | -0.148 | 0.854 |

| | | | | | | |
|---|---|---|---|---|---|---|
| hisC2 | 0.001 | 0.999 | 0.108 | 0.879 | 0.107 | 0.846 |
| Rv3773c | -0.028 | 0.999 | 0.223 | 0.832 | 0.251 | 0.816 |
| echA21 | -0.074 | 0.999 | 0.197 | 0.848 | 0.270 | 0.816 |
| lipE | 0.046 | 0.999 | -0.042 | 0.920 | -0.088 | 0.819 |
| Rv3776 | -0.010 | 0.999 | -0.350 | 0.855 | -0.340 | 0.831 |
| Rv3777 | -0.004 | 0.999 | 0.136 | 0.836 | 0.139 | 0.816 |
| Rv3778c | 0.100 | 0.999 | 0.304 | 0.829 | 0.204 | 0.825 |
| Rv3779 | -0.099 | 0.999 | -0.013 | 0.973 | 0.086 | 0.831 |
| Rv3780 | 0.035 | 0.999 | -0.223 | 0.849 | -0.258 | 0.816 |
| rfbE | 0.018 | 0.999 | -0.209 | 0.832 | -0.227 | 0.816 |
| glfT1 | -0.033 | 0.999 | -0.226 | 0.826 | -0.193 | 0.816 |
| rfbD | -0.041 | 0.999 | -0.125 | 0.885 | -0.084 | 0.904 |
| Rv3784 | -0.038 | 0.999 | -0.222 | 0.846 | -0.184 | 0.831 |
| Rv3785 | 0.096 | 0.999 | 0.014 | 0.974 | -0.082 | 0.865 |
| Rv3786c | 0.136 | 0.999 | 0.132 | 0.831 | -0.004 | 0.993 |
| Rv3787c | 0.016 | NA | 0.160 | 0.848 | 0.144 | 0.822 |
| Rv3788 | 0.090 | 0.999 | 0.018 | 0.971 | -0.073 | 0.868 |
| Rv3789 | 0.100 | 0.999 | -0.123 | 0.890 | -0.223 | 0.817 |
| dprE1 | 0.138 | 0.999 | -0.013 | 0.973 | -0.151 | 0.816 |
| dprE2 | 0.026 | 0.999 | -0.015 | 0.982 | -0.041 | 0.951 |
| aftA | 0.040 | 0.999 | 0.332 | 0.820 | 0.292 | 0.816 |
| embC | 0.106 | 0.999 | 0.308 | 0.831 | 0.203 | 0.836 |
| embA | 0.139 | 0.999 | 0.188 | 0.848 | 0.049 | 0.937 |
| embB | 0.100 | 0.999 | 0.322 | 0.820 | 0.222 | 0.816 |
| Rv3796 | 0.191 | 0.999 | -0.229 | 0.879 | -0.420 | 0.816 |
| fadE35 | -0.071 | 0.999 | -0.064 | 0.897 | 0.008 | 0.986 |
| Rv3798 | -0.205 | 0.999 | -0.014 | 0.986 | 0.190 | 0.840 |
| accD4 | -0.072 | 0.999 | -0.098 | 0.885 | -0.025 | 0.961 |
| pks13 | -0.103 | 0.999 | -0.023 | 0.971 | 0.079 | 0.891 |
| fadD32 | -0.007 | 0.999 | -0.122 | 0.883 | -0.115 | 0.868 |
| Rv3802c | 0.011 | 0.999 | -0.233 | 0.741 | -0.243 | 0.816 |
| fbpD | -0.055 | 0.999 | 0.284 | 0.826 | 0.338 | 0.816 |
| fbpA | -0.030 | 0.999 | 0.246 | 0.831 | 0.276 | 0.816 |
| aftB | 0.030 | 0.999 | 0.216 | 0.843 | 0.186 | 0.819 |
| ubiA | 0.048 | 0.999 | 0.470 | 0.826 | 0.423 | 0.816 |
| Rv3807c | 0.043 | 0.999 | 0.213 | 0.848 | 0.169 | 0.842 |
| glfT2 | 0.158 | 0.999 | 0.159 | 0.843 | 0.001 | 0.998 |
| glf | 0.193 | 0.999 | -0.011 | 0.985 | -0.203 | 0.816 |
| pirG | 0.052 | 0.999 | 0.287 | 0.831 | 0.235 | 0.816 |
| Rv3811 | 0.127 | 0.999 | 0.076 | 0.878 | -0.051 | 0.882 |
| PE_PGRS62 | 0.101 | 0.999 | -0.144 | 0.895 | -0.244 | 0.819 |
| Rv3813c | -0.030 | 0.999 | 0.070 | 0.936 | 0.099 | 0.890 |
| Rv3814c | 0.050 | 0.999 | -0.224 | 0.848 | -0.274 | 0.816 |
| Rv3815c | 0.105 | 0.999 | 0.012 | 0.986 | -0.092 | 0.896 |
| Rv3816c | 0.041 | 0.999 | -0.172 | 0.826 | -0.213 | 0.816 |
| Rv3817 | 0.079 | 0.999 | 0.308 | 0.831 | 0.229 | 0.822 |
| Rv3818 | 0.103 | 0.999 | -0.126 | 0.879 | -0.229 | 0.816 |
| Rv3819 | 0.045 | 0.999 | -0.128 | 0.875 | -0.173 | 0.817 |
| papA2 | 0.152 | 0.999 | -0.380 | 0.831 | -0.532 | 0.816 |
| Rv3821 | 0.024 | 0.999 | 0.176 | 0.853 | 0.152 | 0.837 |
| Rv3822 | 0.211 | 0.999 | -0.125 | 0.931 | -0.336 | 0.819 |
| mmpL8 | 0.106 | 0.999 | 0.010 | 0.980 | -0.096 | 0.846 |
| papA1 | 0.208 | 0.999 | -0.117 | 0.879 | -0.324 | 0.816 |
| pks2 | 0.144 | 0.999 | 0.078 | 0.879 | -0.066 | 0.869 |
| fadD23 | 0.093 | 0.999 | -0.207 | 0.840 | -0.299 | 0.816 |
| Rv3827c | -0.097 | 0.999 | 0.096 | 0.848 | 0.193 | 0.816 |
| Rv3828c | -0.177 | 0.999 | -0.276 | 0.879 | -0.099 | 0.942 |
| Rv3829c | 0.128 | 0.999 | 0.045 | 0.951 | -0.083 | 0.892 |
| Rv3830c | 0.133 | NA | 0.339 | 0.831 | 0.206 | 0.837 |
| Rv3831 | 0.076 | 0.999 | 0.047 | 0.963 | -0.029 | 0.973 |
| Rv3832c | -0.007 | NA | 0.309 | 0.831 | 0.316 | 0.816 |
| Rv3833 | -0.064 | 0.999 | -0.084 | 0.922 | -0.020 | 0.976 |
| serS | -0.095 | NA | 0.073 | 0.922 | 0.168 | 0.817 |
| Rv3835 | 0.135 | 0.999 | -0.191 | 0.843 | -0.326 | 0.816 |
| Rv3836 | 0.005 | 0.999 | -0.091 | 0.922 | -0.096 | 0.896 |
| Rv3837c | 0.076 | 0.999 | 0.086 | 0.922 | 0.010 | 0.988 |
| pheA | 0.005 | 0.999 | 0.224 | 0.859 | 0.219 | 0.836 |
| Rv3839 | 0.111 | 0.999 | -0.130 | 0.856 | -0.241 | 0.816 |
| Rv3840 | 0.223 | 0.999 | 0.246 | 0.826 | 0.023 | 0.969 |
| bfrB | -0.294 | 0.999 | -0.422 | 0.439 | -0.129 | 0.837 |
| glpQ1 | 0.069 | 0.999 | 0.081 | 0.895 | 0.012 | 0.983 |
| Rv3843c | 0.001 | 0.999 | 0.337 | 0.830 | 0.336 | 0.816 |
| Rv3844 | 0.058 | 0.999 | -0.393 | 0.781 | -0.451 | 0.816 |
| Rv3845 | 0.146 | NA | -0.514 | 0.859 | -0.660 | 0.816 |
| sodA | 0.122 | 0.999 | -0.293 | 0.840 | -0.415 | 0.816 |
| Rv3847 | 0.052 | 0.999 | -0.003 | 0.993 | -0.055 | 0.868 |
| Rv3848 | -0.180 | 0.999 | -0.087 | 0.879 | 0.093 | 0.855 |
| espR | 0.263 | 0.999 | -0.041 | 0.959 | -0.304 | 0.816 |
| Rv3850 | 0.100 | 0.999 | 0.029 | 0.948 | -0.071 | 0.860 |
| Rv3851 | 0.047 | NA | 0.142 | 0.859 | 0.095 | 0.871 |
| hns | -0.105 | 0.999 | -0.362 | 0.695 | -0.257 | 0.816 |
| rraA | 0.106 | 0.999 | 0.184 | 0.848 | 0.078 | 0.898 |
| ethA | 0.322 | 0.999 | 0.083 | 0.895 | -0.238 | 0.816 |
| ethR | 0.227 | 0.999 | 0.232 | 0.795 | 0.006 | 0.989 |
| Rv3856c | 0.128 | 0.999 | 0.106 | 0.878 | -0.021 | 0.962 |
| Rv3857c | 0.231 | 0.999 | 0.241 | 0.795 | 0.010 | 0.984 |
| gltD | 0.057 | 0.999 | -0.129 | 0.834 | -0.186 | 0.816 |
| gltB | 0.057 | 0.999 | -0.040 | 0.938 | -0.096 | 0.837 |
| Rv3860 | -0.158 | 0.999 | -0.367 | 0.826 | -0.209 | 0.834 |
| Rv3861 | -0.747 | NA | 0.033 | 0.982 | 0.781 | 0.816 |
| whiB6 | -0.548 | NA | -0.054 | 0.969 | 0.494 | 0.816 |
| Rv3863 | -0.052 | 0.999 | -0.046 | 0.937 | 0.006 | 0.989 |
| espE | -0.059 | 0.999 | -0.273 | 0.848 | -0.214 | 0.842 |
| espF | 0.106 | 0.999 | -0.553 | 0.820 | -0.659 | 0.816 |
| espG1 | 0.016 | 0.999 | -0.248 | 0.659 | -0.264 | 0.816 |
| espH | -0.060 | 0.999 | -0.383 | 0.732 | -0.323 | 0.816 |
| eccA1 | -0.005 | 0.999 | -0.126 | 0.856 | -0.120 | 0.835 |
| eccB1 | 0.031 | 0.999 | -0.101 | 0.879 | -0.132 | 0.824 |
| eccCa1 | -0.033 | 0.999 | -0.168 | 0.831 | -0.135 | 0.817 |
| eccCb1 | -0.070 | 0.999 | -0.183 | 0.659 | -0.114 | 0.816 |
| PE35 | 0.010 | 0.999 | 0.096 | 0.879 | 0.086 | 0.868 |
| PPE68 | -0.084 | 0.999 | -0.161 | 0.820 | -0.077 | 0.835 |
| esxB | -0.350 | 0.999 | -0.917 | 0.877 | -0.567 | 0.892 |
| esxA | 0.129 | 0.999 | -0.024 | 0.971 | -0.153 | 0.831 |
| espI | 0.154 | 0.999 | -0.187 | 0.922 | -0.342 | 0.842 |
| eccD1 | -0.014 | 0.999 | 0.118 | 0.879 | 0.131 | 0.852 |
| espJ | -0.067 | 0.999 | -0.296 | 0.660 | -0.229 | 0.816 |
| espK | 0.079 | 0.999 | -0.077 | 0.926 | -0.156 | 0.836 |
| espL | 0.138 | 0.999 | 0.171 | 0.834 | 0.034 | 0.946 |
| espB | 0.082 | 0.999 | 0.112 | 0.890 | 0.030 | 0.962 |
| eccE1 | 0.007 | 0.999 | 0.129 | 0.856 | 0.122 | 0.835 |
| mycP1 | 0.013 | 0.999 | -0.137 | 0.879 | -0.150 | 0.854 |
| eccA2 | 0.148 | 0.999 | -0.064 | 0.946 | -0.212 | 0.819 |
| eccE2 | 0.163 | 0.999 | 0.000 | 0.999 | -0.163 | 0.816 |
| mycP2 | 0.078 | 0.999 | -0.123 | 0.853 | -0.202 | 0.816 |
| eccD2 | 0.014 | 0.999 | -0.265 | 0.831 | -0.279 | 0.816 |
| Rv3888c | 0.234 | 0.999 | -0.064 | 0.971 | -0.298 | 0.857 |
| espG2 | 0.238 | 0.999 | -0.086 | 0.922 | -0.324 | 0.816 |
| esxC | 0.317 | 0.999 | 0.128 | 0.886 | -0.189 | 0.832 |
| esxD | 0.094 | 0.999 | -0.563 | 0.879 | -0.657 | 0.838 |
| PPE69 | 0.100 | NA | 0.153 | 0.859 | 0.053 | 0.932 |
| PE36 | 0.025 | NA | 0.005 | 0.995 | -0.021 | 0.984 |
| eccC2 | 0.047 | 0.999 | -0.015 | 0.974 | -0.062 | 0.896 |
| eccB2 | 0.112 | 0.999 | 0.289 | 0.826 | 0.177 | 0.835 |
| Rv3896c | 0.009 | 0.999 | -0.022 | 0.977 | -0.031 | 0.969 |
| Rv3897c | -0.138 | 0.999 | -0.027 | 0.973 | 0.112 | 0.881 |
| Rv3898c | 0.073 | 0.999 | 0.081 | 0.879 | 0.008 | 0.986 |
| Rv3899c | 0.026 | 0.999 | 0.298 | 0.820 | 0.272 | 0.816 |
| Rv3900c | 0.127 | 0.999 | -0.062 | 0.961 | -0.189 | 0.862 |
| Rv3901c | 0.024 | 0.999 | -0.163 | 0.834 | -0.187 | 0.816 |
| Rv3902c | 0.180 | 0.999 | -0.392 | 0.820 | -0.572 | 0.816 |
| Rv3903c | 0.047 | 0.999 | -0.281 | 0.879 | -0.328 | 0.839 |
| esxE | 0.018 | NA | -0.768 | 0.831 | -0.786 | 0.816 |
| esxF | 0.078 | NA | -0.641 | 0.848 | -0.719 | 0.817 |
| Rv3906c | -0.118 | 0.999 | -0.053 | 0.946 | 0.065 | 0.917 |
| pcnA | 0.036 | 0.999 | 0.161 | 0.684 | 0.124 | 0.816 |
| mutT4 | 0.052 | 0.999 | 0.092 | 0.862 | 0.040 | 0.920 |
| Rv3909 | 0.050 | 0.999 | 0.261 | 0.832 | 0.210 | 0.819 |
| Rv3910 | -0.003 | 0.999 | 0.368 | 0.826 | 0.371 | 0.816 |
| sigM | -0.056 | 0.999 | 0.102 | 0.909 | 0.158 | 0.840 |
| Rv3912 | -0.008 | NA | -0.099 | 0.946 | -0.091 | 0.936 |
| trxB2 | 0.065 | 0.999 | 0.019 | 0.973 | -0.046 | 0.927 |
| trxC | 0.117 | 0.999 | -0.211 | 0.868 | -0.328 | 0.816 |
| Rv3915 | -0.040 | 0.999 | 0.068 | 0.951 | 0.108 | 0.904 |
| Rv3916c | -0.003 | 0.999 | -0.121 | 0.883 | -0.118 | 0.866 |
| parB | 0.062 | 0.999 | -0.083 | 0.872 | -0.145 | 0.816 |
| parA | 0.200 | 0.999 | -0.166 | 0.848 | -0.365 | 0.816 |
| gid | 0.103 | 0.999 | -0.125 | 0.948 | -0.228 | 0.882 |
| Rv3920c | 0.086 | 0.999 | -0.022 | 0.985 | -0.108 | 0.930 |
| Rv3921c | 0.130 | 0.999 | -0.012 | 0.989 | -0.142 | 0.879 |
| Rv3922c | 0.088 | 0.999 | 0.163 | 0.870 | 0.075 | 0.918 |
| rnpA | 0.198 | 0.999 | -0.092 | 0.859 | -0.290 | 0.816 |
| rpmH | 0.028 | 0.999 | -0.020 | 0.973 | -0.049 | 0.931 |
| socA | -0.460 | 0.999 | -0.805 | 0.374 | -0.345 | 0.816 |
| socB | -0.622 | NA | -0.934 | 0.010 | -0.313 | 0.816 |
| AS1890 | 0.128 | NA | -0.099 | 0.940 | -0.227 | 0.846 |
| AS1890 | 0.128 | NA | -0.099 | 0.940 | -0.227 | 0.846 |
| ASdes | 0.150 | NA | -0.816 | 0.795 | -0.965 | 0.816 |
| ASdes | 0.150 | NA | -0.816 | 0.795 | -0.965 | 0.816 |
| ASpks | 0.072 | NA | -2.799 | NA | -2.871 | NA |
| ASpks | 0.072 | NA | -2.799 | NA | -2.871 | NA |
| B11 | -0.549 | 0.999 | 0.874 | 0.850 | 1.424 | 0.816 |
| B11 | -0.549 | 0.999 | 0.874 | 0.850 | 1.424 | 0.816 |
| B55 | -0.357 | NA | 0.811 | 0.848 | 1.167 | 0.816 |
| B55 | -0.357 | NA | 0.811 | 0.848 | 1.167 | 0.816 |
| C8 | -0.244 | NA | -0.680 | 0.825 | -0.437 | 0.816 |
| C8 | -0.244 | NA | -0.680 | 0.825 | -0.437 | 0.816 |
| F6 | 0.029 | NA | 0.809 | 0.732 | 0.780 | 0.816 |
| F6 | 0.029 | NA | 0.809 | 0.732 | 0.780 | 0.816 |
| G2 | -0.422 | NA | -0.758 | 0.832 | -0.336 | 0.879 |
| G2 | -0.422 | NA | -0.758 | 0.832 | -0.336 | 0.879 |
| mcr10 | 0.119 | 0.999 | 0.025 | 0.974 | -0.093 | 0.897 |
| mcr10 | 0.119 | 0.999 | 0.025 | 0.974 | -0.093 | 0.897 |
| mcr11 | -0.502 | 0.999 | 0.828 | 0.826 | 1.329 | 0.816 |
| mcr11 | -0.502 | 0.999 | 0.828 | 0.826 | 1.329 | 0.816 |

| | | | | | | |
|---|---|---|---|---|---|---|
| Rv0691c | -0.204 | 0.999 | -0.290 | 0.861 | -0.086 | 0.942 |
| Rv0692 | 0.211 | NA | 0.247 | 0.826 | 0.037 | 0.950 |
| Rv0693 | 0.120 | 0.999 | 0.193 | 0.826 | 0.073 | 0.866 |
| Rv0694 | 0.010 | 0.999 | 0.274 | 0.826 | 0.263 | 0.816 |
| Rv0695 | 0.040 | 0.999 | 0.139 | 0.848 | 0.100 | 0.855 |
| Rv0696 | 0.113 | 0.999 | 0.001 | 0.997 | -0.112 | 0.843 |
| Rv0697 | -0.020 | 0.999 | -0.049 | 0.926 | -0.029 | 0.942 |
| Rv0698 | 0.044 | 0.999 | 0.238 | 0.832 | 0.193 | 0.819 |
| Rv0699 | -0.043 | NA | -0.224 | 0.868 | -0.181 | 0.862 |
| rpsJ | 0.094 | 0.999 | -0.016 | 0.973 | -0.111 | 0.831 |
| rplC | 0.049 | 0.999 | 0.171 | 0.879 | 0.122 | 0.883 |
| rplD | 0.097 | 0.999 | -0.123 | 0.895 | -0.220 | 0.819 |
| rplW | 0.008 | 0.999 | -0.403 | 0.848 | -0.412 | 0.816 |
| rplB | 0.160 | 0.999 | -0.015 | 0.987 | -0.175 | 0.862 |
| rpsS | 0.090 | 0.999 | -0.123 | 0.886 | -0.214 | 0.817 |
| rplV | 0.042 | 0.999 | 0.267 | 0.831 | 0.224 | 0.816 |
| rpsC | 0.119 | 0.999 | 0.149 | 0.826 | 0.030 | 0.932 |
| rplP | 0.145 | 0.999 | 0.157 | 0.859 | 0.012 | 0.986 |
| rpmC | 0.226 | 0.999 | -0.017 | 0.990 | -0.243 | 0.868 |
| rpsQ | 0.041 | 0.999 | -0.060 | 0.970 | -0.101 | 0.936 |
| atsA | 0.040 | 0.999 | -0.176 | 0.848 | -0.216 | 0.816 |
| Rv0712 | -0.014 | 0.999 | 0.039 | 0.918 | 0.053 | 0.865 |
| Rv0713 | 0.077 | 0.999 | -0.031 | 0.948 | -0.108 | 0.822 |
| rplN | 0.064 | 0.999 | -0.030 | 0.973 | -0.095 | 0.911 |
| rplX | 0.024 | 0.999 | -0.147 | 0.859 | -0.171 | 0.817 |
| rplE | 0.030 | 0.999 | -0.349 | 0.717 | -0.379 | 0.816 |
| rpsN1 | -0.267 | 0.999 | -0.399 | 0.931 | -0.132 | 0.971 |
| rpsH | 0.001 | 0.999 | 0.078 | 0.866 | 0.077 | 0.837 |
| rplF | 0.031 | 0.999 | -0.161 | 0.841 | -0.192 | 0.816 |
| rplR | 0.074 | 0.999 | 0.234 | 0.895 | 0.160 | 0.917 |
| rpsE | 0.102 | 0.999 | -0.016 | 0.974 | -0.118 | 0.838 |
| rpmD | 0.158 | 0.999 | -0.333 | 0.848 | -0.490 | 0.816 |
| rplO | -0.044 | 0.999 | 0.078 | 0.883 | 0.122 | 0.819 |
| sppA | -0.071 | 0.999 | 0.018 | 0.973 | 0.089 | 0.865 |
| Rv0725c | 0.123 | 0.999 | -0.023 | 0.977 | -0.146 | 0.873 |
| Rv0726c | 0.022 | 0.999 | -0.165 | 0.832 | -0.187 | 0.816 |
| fucA | 0.042 | NA | -0.020 | 0.989 | -0.062 | 0.967 |
| serA2 | -0.245 | NA | -0.048 | 0.967 | 0.197 | 0.846 |
| xylB | 0.079 | NA | 0.199 | 0.852 | 0.120 | 0.870 |
| Rv0730 | -0.017 | 0.999 | 0.022 | 0.973 | 0.039 | 0.946 |
| Rv0731c | -0.036 | NA | -0.006 | 0.994 | 0.030 | 0.969 |
| secY | 0.185 | 0.999 | -0.061 | 0.886 | -0.247 | 0.816 |
| adk | 0.019 | 0.999 | -0.084 | 0.916 | -0.103 | 0.868 |
| mapA | 0.045 | 0.999 | 0.347 | 0.831 | 0.302 | 0.816 |
| sigL | 0.040 | NA | -0.204 | 0.848 | -0.244 | 0.816 |
| rslA | -0.108 | NA | 0.218 | 0.866 | 0.326 | 0.816 |
| Rv0737 | 0.047 | NA | 0.009 | 0.989 | -0.038 | 0.957 |
| Rv0738 | -0.127 | 0.999 | 0.801 | 0.829 | 0.927 | 0.816 |
| Rv0739 | 0.117 | 0.999 | -0.128 | 0.922 | -0.245 | 0.836 |
| Rv0740 | -0.003 | 0.999 | 0.135 | 0.832 | 0.137 | 0.816 |
| Rv0742 | -0.049 | 0.999 | 0.168 | 0.848 | 0.217 | 0.816 |
| Rv0743c | -0.178 | 0.999 | -0.264 | 0.845 | -0.085 | 0.921 |
| Rv0744c | 0.008 | 0.999 | -0.277 | 0.820 | -0.284 | 0.816 |
| Rv0745 | 0.013 | NA | 0.244 | 0.886 | 0.231 | 0.871 |
| PE_PGRS9 | -0.129 | 0.999 | -0.166 | 0.924 | -0.038 | 0.979 |
| PE_PGRS10 | -0.467 | 0.999 | -0.450 | 0.832 | 0.017 | 0.989 |
| vapB31 | 0.003 | 0.999 | 0.284 | 0.829 | 0.282 | 0.816 |
| vapC31 | 0.093 | 0.999 | -0.198 | 0.848 | -0.291 | 0.816 |
| Rv0749A | -0.024 | 0.999 | -0.360 | 0.845 | -0.336 | 0.817 |
| Rv0750 | -0.280 | 0.999 | 0.054 | 0.952 | 0.333 | 0.816 |
| mmsB | -0.121 | 0.999 | 0.012 | 0.987 | 0.134 | 0.868 |
| fadE9 | -0.108 | 0.999 | -0.015 | 0.973 | 0.093 | 0.835 |
| mmsA | -0.045 | 0.999 | 0.018 | 0.973 | 0.062 | 0.895 |
| PE_PGRS11 | -0.218 | 0.999 | -0.338 | 0.866 | -0.119 | 0.936 |
| PPE12 | 0.043 | 0.999 | -0.075 | 0.977 | -0.118 | 0.967 |
| Rv0756c | -0.078 | 0.999 | 0.417 | 0.795 | 0.495 | 0.816 |
| phoP | 0.011 | 0.999 | -0.244 | 0.848 | -0.255 | 0.817 |
| phoR | 0.041 | 0.999 | 0.081 | 0.895 | 0.040 | 0.939 |
| Rv0759c | 0.028 | 0.999 | -0.182 | 0.879 | -0.210 | 0.835 |
| Rv0760c | 0.073 | 0.999 | 0.093 | 0.886 | 0.021 | 0.969 |
| adhB | -0.049 | 0.999 | 0.026 | 0.969 | 0.074 | 0.884 |
| Rv0762c | -0.039 | 0.999 | -0.248 | 0.843 | -0.210 | 0.820 |
| Rv0763c | 0.141 | NA | -0.112 | 0.970 | -0.253 | 0.921 |
| cyp51 | -0.103 | 0.999 | -0.515 | 0.699 | -0.412 | 0.816 |
| Rv0765c | -0.132 | NA | 0.130 | 0.859 | 0.262 | 0.816 |
| cyp123 | -0.077 | 0.999 | -0.150 | 0.848 | -0.074 | 0.891 |
| Rv0767c | -0.214 | NA | -0.376 | 0.831 | -0.162 | 0.877 |
| aldA | -0.062 | 0.999 | -0.281 | 0.832 | -0.218 | 0.827 |
| Rv0769 | -0.182 | NA | -0.339 | 0.826 | -0.157 | 0.842 |
| Rv0770 | 0.078 | NA | -0.397 | 0.785 | -0.475 | 0.816 |
| Rv0771 | 0.148 | NA | -0.373 | 0.831 | -0.521 | 0.816 |
| purD | -0.103 | 0.999 | 0.044 | 0.958 | 0.147 | 0.837 |
| ggtA | 0.102 | 0.999 | 0.037 | 0.958 | -0.065 | 0.904 |
| Rv0774c | -0.058 | 0.999 | 0.357 | 0.832 | 0.415 | 0.816 |
| Rv0775 | 0.157 | 0.999 | 0.218 | 0.832 | 0.061 | 0.924 |
| Rv0776c | -0.236 | NA | 0.210 | 0.848 | 0.446 | 0.816 |
| purB | 0.023 | 0.999 | 0.145 | 0.834 | 0.122 | 0.819 |
| cyp126 | -0.038 | 0.999 | 0.116 | 0.865 | 0.154 | 0.816 |
| Rv0779c | -0.101 | NA | 0.128 | 0.940 | 0.230 | 0.868 |
| purC | 0.113 | 0.999 | -0.209 | 0.831 | -0.322 | 0.816 |
| emrB | -0.038 | 0.999 | 0.025 | 0.964 | 0.063 | 0.893 |
| Rv0784 | 0.041 | 0.999 | 0.300 | 0.832 | 0.258 | 0.817 |
| Rv0785 | 0.037 | 0.999 | -0.033 | 0.964 | -0.070 | 0.908 |
| Rv0786c | -0.005 | 0.999 | -0.169 | 0.843 | -0.164 | 0.816 |
| Rv0787 | -0.057 | NA | 0.079 | 0.948 | 0.136 | 0.892 |
| Rv0787A | -0.367 | 0.999 | -0.787 | 0.826 | -0.420 | 0.837 |
| purQ | -0.169 | 0.999 | 0.128 | 0.879 | 0.298 | 0.816 |
| Rv0789c | 0.063 | 0.999 | -0.080 | 0.922 | -0.143 | 0.838 |
| Rv0790c | -0.210 | 0.999 | -0.697 | 0.374 | -0.486 | 0.816 |
| Rv0791c | -0.202 | 0.999 | -0.212 | 0.832 | -0.010 | 0.988 |
| Rv0792c | -0.165 | 0.999 | -0.031 | 0.970 | 0.134 | 0.848 |
| Rv0793 | -0.093 | NA | 0.039 | 0.975 | 0.132 | 0.924 |
| Rv0794c | 0.033 | 0.999 | -0.034 | 0.938 | -0.067 | 0.862 |
| Rv0797 | -0.296 | NA | -0.558 | 0.820 | -0.262 | 0.837 |
| cfp29 | 0.094 | 0.999 | 0.019 | 0.973 | -0.075 | 0.891 |
| Rv0799c | 0.144 | 0.999 | -0.165 | 0.859 | -0.309 | 0.816 |
| pepC | -0.041 | 0.999 | -0.032 | 0.948 | 0.009 | 0.983 |
| Rv0801 | -0.125 | 0.999 | 0.234 | 0.859 | 0.359 | 0.816 |
| Rv0802c | 0.088 | 0.999 | 0.178 | 0.879 | 0.090 | 0.926 |
| purL | 0.102 | 0.999 | 0.158 | 0.866 | 0.056 | 0.936 |
| Rv1507c | 0.174 | 0.999 | -0.282 | 0.879 | -0.456 | 0.816 |
| Rv1508A | -0.097 | 0.999 | -0.057 | 0.942 | 0.040 | 0.947 |
| Rv1508c | 0.054 | 0.999 | -0.102 | 0.859 | -0.156 | 0.816 |
| Rv1509 | 0.011 | 0.999 | -0.398 | 0.859 | -0.409 | 0.832 |
| Rv1510 | 0.142 | 0.999 | 0.126 | 0.879 | -0.016 | 0.983 |
| gmdA | 0.035 | 0.999 | -0.132 | 0.848 | -0.166 | 0.816 |
| epiA | 0.042 | 0.999 | 0.145 | 0.879 | 0.103 | 0.886 |
| Rv1513 | 0.181 | 0.999 | -0.131 | 0.895 | -0.312 | 0.816 |
| Rv1514c | -0.042 | NA | 0.014 | 0.977 | 0.056 | 0.920 |
| Rv1515c | 0.091 | 0.999 | -0.158 | 0.879 | -0.249 | 0.816 |
| Rv1516c | 0.139 | 0.999 | 0.022 | 0.971 | -0.117 | 0.842 |
| Rv1517 | 0.295 | NA | 0.135 | 0.961 | -0.160 | 0.942 |
| Rv1518 | -0.114 | NA | -0.544 | 0.732 | -0.430 | 0.816 |
| Rv1519 | 0.077 | 0.999 | -0.121 | 0.927 | -0.197 | 0.865 |
| Rv1520 | 0.093 | 0.999 | -0.192 | 0.848 | -0.285 | 0.816 |
| fadD25 | 0.130 | 0.999 | -0.153 | 0.831 | -0.283 | 0.816 |
| mmpL12 | 0.100 | 0.999 | -0.093 | 0.879 | -0.193 | 0.816 |
| Rv1523 | -0.057 | 0.999 | 0.080 | 0.935 | 0.137 | 0.868 |
| Rv1524 | 0.227 | 0.999 | 0.140 | 0.831 | -0.087 | 0.837 |
| wbbL2 | 0.078 | 0.999 | 0.151 | 0.879 | 0.073 | 0.930 |
| Rv1526c | -0.035 | 0.999 | -0.077 | 0.942 | -0.042 | 0.958 |
| pks5 | 0.135 | 0.999 | -0.043 | 0.937 | -0.178 | 0.816 |
| papA4 | -0.287 | NA | -0.915 | NA | -0.628 | NA |
| fadD24 | 0.023 | 0.999 | -0.205 | 0.848 | -0.228 | 0.816 |
| adh | -0.228 | 0.999 | 0.169 | 0.848 | 0.398 | 0.816 |
| Rv1531 | -0.012 | 0.999 | 0.158 | 0.843 | 0.170 | 0.816 |
| Rv1532c | 0.113 | 0.999 | -0.110 | 0.879 | -0.223 | 0.816 |
| Rv1533 | 0.046 | 0.999 | 0.294 | 0.826 | 0.248 | 0.816 |
| Rv1534 | 0.184 | 0.999 | 0.218 | 0.592 | 0.034 | 0.917 |
| Rv1535 | -0.021 | 0.999 | 0.176 | 0.895 | 0.197 | 0.866 |
| ileS | -0.062 | 0.999 | -0.132 | 0.848 | -0.069 | 0.884 |
| dinX | -0.113 | 0.999 | -0.005 | 0.988 | 0.108 | 0.819 |
| ansA | 0.061 | 0.999 | 0.289 | 0.826 | 0.229 | 0.816 |
| lspA | -0.045 | 0.999 | -0.120 | 0.848 | -0.076 | 0.868 |
| Rv1540 | -0.007 | 0.999 | -0.005 | 0.994 | 0.002 | 0.998 |
| lprI | 0.028 | NA | -0.220 | 0.859 | -0.247 | 0.819 |
| glbN | -0.406 | NA | -0.730 | 0.832 | -0.324 | 0.879 |
| Rv1543 | 0.098 | 0.999 | -0.289 | 0.859 | -0.387 | 0.816 |
| Rv1544 | 0.071 | 0.999 | -0.152 | 0.826 | -0.223 | 0.816 |
| Rv1545 | 0.070 | 0.999 | -0.441 | 0.831 | -0.510 | 0.816 |
| Rv1546 | -0.019 | 0.999 | 0.220 | 0.862 | 0.239 | 0.831 |
| dnaE1 | 0.101 | 0.999 | -0.059 | 0.951 | -0.160 | 0.854 |
| PPE21 | 0.001 | 0.999 | -0.214 | 0.913 | -0.215 | 0.891 |
| fadD11 | 0.240 | 0.999 | 0.196 | 0.848 | -0.045 | 0.947 |
| plsB1 | 0.237 | 0.999 | 0.129 | 0.879 | -0.108 | 0.879 |
| frdA | 0.450 | 0.999 | 0.263 | 0.850 | -0.187 | 0.859 |
| frdB | 0.331 | NA | -0.525 | 0.855 | -0.856 | 0.816 |
| frdC | 0.106 | NA | -0.221 | 0.922 | -0.328 | 0.866 |
| frdD | 0.380 | NA | 0.020 | 0.988 | -0.359 | 0.835 |
| Rv1556 | 0.092 | 0.999 | 0.166 | 0.848 | 0.074 | 0.896 |
| mmpL6 | -0.130 | 0.999 | 0.074 | 0.940 | 0.205 | 0.830 |
| Rv1558 | -0.118 | 0.999 | 0.402 | 0.826 | 0.519 | 0.816 |
| ilvA | -0.021 | 0.999 | 0.080 | 0.918 | 0.101 | 0.868 |
| vapB11 | -0.127 | NA | 0.082 | 0.962 | 0.209 | 0.884 |
| vapC11 | -0.097 | 0.999 | -0.073 | 0.948 | 0.023 | 0.979 |
| treZ | 0.013 | 0.999 | -0.087 | 0.885 | -0.100 | 0.854 |
| treY | -0.018 | 0.999 | 0.122 | 0.850 | 0.140 | 0.816 |
| treX | 0.011 | 0.999 | -0.160 | 0.879 | -0.171 | 0.837 |
| Rv1565c | 0.001 | 0.999 | 0.057 | 0.941 | 0.055 | 0.930 |
| Rv1566c | 0.101 | 0.999 | 0.093 | 0.863 | -0.008 | 0.985 |
| Rv1567c | -0.045 | 0.999 | -0.210 | 0.831 | -0.164 | 0.817 |
| bioA | -0.190 | 0.999 | 0.344 | 0.831 | 0.534 | 0.816 |
| bioF1 | 0.004 | 0.999 | 0.749 | 0.795 | 0.746 | 0.816 |
| bioD | 0.180 | 0.999 | 0.525 | 0.820 | 0.346 | 0.816 |
| Rv1571 | 0.061 | NA | 0.676 | 0.820 | 0.616 | 0.816 |
| Rv1573 | -0.281 | NA | -0.094 | 0.922 | 0.187 | 0.835 |
| Rv1574 | 0.013 | NA | 0.152 | 0.927 | 0.139 | 0.920 |
| Rv1575 | -0.017 | NA | -0.184 | 0.879 | -0.167 | 0.855 |
| Rv1576c | -0.243 | 0.999 | -0.174 | 0.832 | 0.068 | 0.891 |
| Rv1577c | -0.130 | 0.999 | -0.269 | 0.854 | -0.138 | 0.892 |
| Rv1578c | 0.015 | NA | 0.488 | 0.826 | 0.473 | 0.816 |
| Rv1579c | -0.212 | NA | -0.210 | 0.853 | 0.002 | 0.998 |
| Rv1580c | -0.066 | NA | -0.406 | 0.826 | -0.340 | 0.816 |
| Rv1581c | -0.137 | NA | -0.556 | 0.842 | -0.420 | 0.835 |
| Rv1582c | -0.193 | 0.999 | -0.350 | 0.832 | -0.157 | 0.875 |
| Rv1583c | -0.147 | NA | 0.389 | 0.826 | 0.535 | 0.816 |
| Rv1584c | -0.004 | NA | 0.244 | 0.869 | 0.248 | 0.837 |
| Rv1585c | -0.026 | NA | -0.337 | 0.831 | -0.311 | 0.816 |
| Rv1586c | -0.225 | 0.999 | -0.121 | 0.879 | 0.104 | 0.868 |
| Rv1587c | -0.173 | NA | -0.833 | 0.845 | -0.660 | 0.835 |
| Rv1588c | -0.016 | 0.999 | -0.501 | 0.843 | -0.484 | 0.816 |
| bioB | 0.123 | 0.999 | 0.089 | 0.922 | -0.035 | 0.958 |
| Rv1590 | -0.034 | 0.999 | 0.311 | 0.859 | 0.344 | 0.827 |
| Rv1591 | 0.113 | 0.999 | 0.400 | 0.684 | 0.287 | 0.816 |
| Rv1592c | -0.159 | 0.999 | -0.119 | 0.832 | 0.040 | 0.904 |
| Rv1593c | -0.030 | 0.999 | 0.114 | 0.879 | 0.144 | 0.835 |
| nadA | 0.096 | 0.999 | 0.112 | 0.826 | 0.015 | 0.954 |
| nadB | 0.084 | 0.999 | 0.109 | 0.844 | 0.025 | 0.942 |
| nadC | 0.121 | 0.999 | 0.222 | 0.848 | 0.102 | 0.896 |
| Rv1597 | -0.117 | 0.999 | 0.179 | 0.883 | 0.296 | 0.817 |
| Rv1598c | 0.078 | 0.999 | 0.159 | 0.879 | 0.081 | 0.909 |
| hisD | -0.138 | 0.999 | -0.082 | 0.936 | 0.057 | 0.942 |
| hisC1 | -0.115 | 0.999 | -0.337 | 0.701 | -0.222 | 0.816 |
| hisB | -0.090 | 0.999 | -0.481 | 0.659 | -0.392 | 0.816 |
| hisH | -0.113 | 0.999 | -0.078 | 0.924 | 0.035 | 0.956 |
| hisA | -0.155 | 0.999 | 0.066 | 0.948 | 0.221 | 0.828 |
| impA | -0.021 | 0.999 | -0.185 | 0.831 | -0.165 | 0.816 |
| hisF | -0.229 | 0.999 | 0.191 | 0.879 | 0.420 | 0.816 |
| hisI | -0.181 | 0.999 | -0.080 | 0.910 | 0.101 | 0.865 |
| chaA | -0.006 | 0.999 | -0.074 | 0.934 | -0.068 | 0.926 |
| bcpB | -0.053 | 0.999 | 0.017 | 0.972 | 0.071 | 0.870 |
| trpE | 0.049 | 0.999 | -0.047 | 0.930 | -0.096 | 0.840 |
| Rv1610 | 0.087 | 0.999 | 0.442 | 0.820 | 0.354 | 0.816 |
| trpC | 0.041 | 0.999 | 0.108 | 0.895 | 0.067 | 0.924 |
| trpB | 0.146 | 0.999 | 0.190 | 0.879 | 0.044 | 0.962 |
| trpA | 0.219 | 0.999 | 0.009 | 0.992 | -0.209 | 0.846 |
| lgt | 0.088 | 0.999 | 0.048 | 0.938 | -0.040 | 0.936 |
| Rv1615 | -0.017 | 0.999 | 0.331 | 0.832 | 0.348 | 0.816 |
| Rv1616 | 0.065 | 0.999 | 0.183 | 0.895 | 0.117 | 0.920 |
| pykA | -0.055 | 0.999 | 0.136 | 0.879 | 0.191 | 0.827 |
| Rv2327 | -0.093 | 0.999 | 0.190 | 0.834 | 0.283 | 0.816 |
| PE23 | -0.066 | 0.999 | -0.102 | 0.907 | -0.036 | 0.957 |
| narK1 | 0.103 | 0.999 | 0.031 | 0.963 | -0.072 | 0.896 |
| lppP | 0.047 | NA | -0.366 | 0.831 | -0.413 | 0.816 |
| Rv2331 | 0.114 | 0.999 | 0.064 | 0.970 | -0.051 | 0.971 |
| Rv2331A | 0.012 | 0.999 | 0.135 | 0.903 | 0.123 | 0.892 |
| mez | 0.054 | 0.999 | -0.124 | 0.866 | -0.178 | 0.816 |
| stp | 0.019 | 0.999 | -0.067 | 0.922 | -0.086 | 0.877 |
| cysK1 | 0.065 | 0.999 | 0.026 | 0.962 | -0.039 | 0.932 |
| cysE | -0.069 | 0.999 | 0.195 | 0.855 | 0.264 | 0.816 |
| Rv2336 | 0.085 | 0.999 | -0.218 | 0.829 | -0.303 | 0.816 |
| Rv2337c | 0.253 | 0.999 | 0.116 | 0.848 | -0.137 | 0.816 |
| moeW | 0.276 | 0.999 | -0.552 | 0.826 | -0.827 | 0.816 |
| mmpL9 | 0.193 | 0.999 | -0.109 | 0.879 | -0.302 | 0.816 |
| PE_PGRS39 | -0.199 | 0.999 | -0.014 | 0.991 | 0.185 | 0.892 |
| lppQ | 0.019 | NA | -0.356 | 0.826 | -0.375 | 0.816 |
| Rv2342 | -0.117 | 0.999 | 0.166 | 0.877 | 0.284 | 0.816 |
| dnaG | -0.011 | 0.999 | 0.039 | 0.912 | 0.050 | 0.865 |
| dgt | 0.114 | 0.999 | 0.255 | 0.568 | 0.141 | 0.816 |
| Rv2345 | 0.051 | 0.999 | -0.035 | 0.922 | -0.085 | 0.817 |
| esxO | 0.070 | 0.999 | -0.019 | 0.973 | -0.088 | 0.868 |
| esxP | 0.054 | 0.999 | -0.499 | 0.854 | -0.554 | 0.817 |
| Rv2348c | 0.067 | 0.999 | -0.195 | 0.958 | -0.262 | 0.930 |
| plcC | 0.066 | 0.999 | 0.044 | 0.948 | -0.022 | 0.967 |
| plcB | 0.105 | 0.999 | -0.141 | 0.863 | -0.246 | 0.816 |
| plcA | 0.268 | 0.999 | 0.004 | 0.995 | -0.265 | 0.832 |
| PPE38 | -0.174 | 0.999 | -0.983 | 0.866 | -0.809 | 0.860 |
| PPE39 | -0.027 | NA | -0.963 | 0.852 | -0.936 | 0.827 |
| glyS | 0.003 | 0.999 | -0.152 | 0.879 | -0.155 | 0.840 |
| smtB | 0.051 | 0.999 | -0.022 | 0.979 | -0.073 | 0.937 |
| zur | -0.056 | 0.999 | -0.343 | 0.409 | -0.287 | 0.816 |
| Rv2360c | -0.158 | 0.999 | 0.043 | 0.970 | 0.201 | 0.852 |
| Rv2361c | 0.074 | 0.999 | -0.197 | 0.859 | -0.271 | 0.816 |
| recO | -0.112 | 0.999 | 0.179 | 0.879 | 0.291 | 0.816 |
| amiA2 | -0.187 | 0.999 | -0.105 | 0.879 | 0.082 | 0.868 |
| era | -0.157 | 0.999 | 0.006 | 0.993 | 0.163 | 0.835 |
| Rv2365c | -0.242 | 0.999 | 0.088 | 0.961 | 0.330 | 0.837 |
| Rv2366c | -0.078 | 0.999 | -0.079 | 0.905 | 0.000 | 0.999 |
| Rv2367c | 0.042 | 0.999 | -0.067 | 0.931 | -0.109 | 0.866 |
| phoH1 | -0.019 | 0.999 | -0.245 | 0.825 | -0.226 | 0.816 |
| Rv2369c | -0.304 | NA | -0.831 | 0.829 | -0.527 | 0.832 |
| Rv2370c | 0.102 | 0.999 | -0.970 | 0.848 | -1.071 | 0.816 |
| PE_PGRS40 | -0.460 | NA | -0.660 | 0.832 | -0.200 | 0.917 |
| Rv2372c | -0.138 | 0.999 | 0.072 | 0.946 | 0.210 | 0.832 |
| dnaJ2 | -0.078 | 0.999 | -0.025 | 0.960 | 0.053 | 0.894 |
| hrcA | 0.019 | 0.999 | 0.031 | 0.948 | 0.012 | 0.976 |
| Rv2375 | -0.033 | 0.999 | 0.005 | 0.994 | 0.038 | 0.955 |
| cfp2 | -0.070 | 0.999 | 1.005 | 0.803 | 1.075 | 0.816 |
| Rv2377c | 0.407 | 0.999 | 0.014 | 0.990 | -0.393 | 0.816 |
| mbtG | 0.224 | 0.999 | 0.234 | 0.795 | 0.009 | 0.984 |
| mbtF | 0.149 | 0.999 | 0.307 | 0.831 | 0.158 | 0.855 |
| mbtE | 0.079 | 0.999 | 0.126 | 0.879 | 0.047 | 0.939 |
| mbtD | 0.093 | 0.999 | 0.072 | 0.930 | -0.021 | 0.974 |
| mbtC | 0.058 | 0.999 | -0.110 | 0.848 | -0.168 | 0.816 |
| mbtB | -0.076 | 0.999 | 0.151 | 0.895 | 0.226 | 0.837 |
| mbtA | 0.134 | 0.999 | 0.059 | 0.941 | -0.075 | 0.904 |
| mbtJ | -0.054 | 0.999 | -0.026 | 0.970 | 0.028 | 0.958 |
| Rv2386a | -0.174 | 0.999 | 0.257 | 0.879 | 0.431 | 0.816 |
| mbtI | 0.025 | 0.999 | 0.110 | 0.920 | 0.086 | 0.921 |
| Rv2387 | 0.112 | 0.999 | -0.135 | 0.848 | -0.247 | 0.816 |
| hemN | -0.006 | 0.999 | 0.258 | 0.826 | 0.264 | 0.816 |
| rpfD | -0.018 | 0.999 | -0.083 | 0.916 | -0.065 | 0.917 |
| Rv2390c | -0.229 | 0.999 | -0.141 | 0.886 | 0.088 | 0.916 |
| sirA | 0.054 | 0.999 | -0.116 | 0.886 | -0.170 | 0.835 |
| cysH | 0.052 | 0.999 | 0.056 | 0.901 | 0.003 | 0.994 |
| che1 | 0.162 | 0.999 | 0.090 | 0.886 | -0.072 | 0.892 |
| ggtB | 0.077 | 0.999 | 0.079 | 0.903 | 0.001 | 0.998 |
| Rv2395 | 0.086 | 0.999 | 0.016 | 0.973 | -0.070 | 0.870 |
| aprA | NA | NA | NA | NA | NA | NA |
| aprB | -0.317 | 0.999 | -0.661 | 0.659 | -0.344 | 0.816 |
| PE_PGRS41 | -0.194 | 0.999 | -0.103 | 0.890 | 0.092 | 0.882 |
| cysA1 | -0.118 | 0.999 | 0.225 | 0.848 | 0.343 | 0.816 |
| cysW | 0.000 | 0.999 | -0.158 | 0.879 | -0.159 | 0.862 |
| cysT | 0.146 | 0.999 | 0.095 | 0.897 | -0.051 | 0.936 |
| subI | 0.079 | 0.999 | 0.313 | 0.361 | 0.234 | 0.816 |
| Rv2401 | -0.005 | NA | -0.746 | 0.732 | -0.740 | 0.816 |
| Rv2401A | -0.100 | NA | 0.317 | 0.848 | 0.417 | 0.816 |
| Rv2402 | 0.133 | 0.999 | -0.143 | 0.879 | -0.277 | 0.816 |
| lppR | 0.202 | 0.999 | 0.032 | 0.970 | -0.170 | 0.836 |
| lepA | 0.014 | 0.999 | -0.150 | 0.831 | -0.164 | 0.816 |
| Rv2405 | -0.069 | 0.999 | 0.120 | 0.879 | 0.189 | 0.817 |
| Rv2406c | 0.195 | 0.999 | 0.455 | 0.361 | 0.260 | 0.816 |
| Rv2407 | -0.076 | NA | -0.226 | 0.826 | -0.150 | 0.819 |
| PE24 | -0.325 | NA | -0.709 | 0.848 | -0.384 | 0.868 |
| Rv2409c | 0.190 | 0.999 | -0.074 | 0.935 | -0.265 | 0.816 |
| Rv2410c | 0.266 | 0.999 | -0.125 | 0.895 | -0.391 | 0.816 |
| Rv2411c | 0.116 | 0.999 | 0.135 | 0.845 | 0.019 | 0.962 |
| rpsT | 0.029 | 0.999 | 0.051 | 0.964 | 0.021 | 0.984 |
| Rv2413c | -0.238 | 0.999 | 0.809 | 0.809 | 1.047 | 0.816 |
| Rv2414c | -0.039 | 0.999 | 0.500 | 0.820 | 0.539 | 0.816 |
| Rv2415c | -0.038 | NA | 0.399 | 0.826 | 0.436 | 0.816 |
| eis | 0.101 | 0.999 | 0.123 | 0.848 | 0.022 | 0.955 |
| Rv2417c | 0.111 | 0.999 | 0.228 | 0.848 | 0.118 | 0.878 |
| Rv2418c | 0.135 | 0.999 | 0.069 | 0.901 | -0.066 | 0.889 |
| gpgP | -0.035 | 0.999 | 0.348 | 0.843 | 0.383 | 0.816 |
| Rv2420c | 0.186 | 0.999 | -0.214 | 0.866 | -0.400 | 0.816 |
| nadD | 0.123 | 0.999 | 0.023 | 0.963 | -0.100 | 0.837 |
| Rv2422 | 0.193 | NA | -0.019 | 0.994 | -0.212 | 0.930 |
| Rv2423 | 0.067 | 0.999 | -0.052 | 0.947 | -0.119 | 0.860 |
| Rv2424c | 0.095 | 0.999 | -0.199 | 0.826 | -0.294 | 0.816 |
| Rv2425c | 0.026 | 0.999 | 0.063 | 0.914 | 0.037 | 0.936 |
| Rv2426c | 0.128 | 0.999 | -0.225 | 0.848 | -0.353 | 0.816 |
| proA | 0.004 | 0.999 | 0.011 | 0.975 | 0.007 | 0.986 |
| ahpC | -0.487 | 0.999 | -0.957 | 0.054 | -0.470 | 0.816 |
| ahpD | -0.390 | 0.999 | -0.636 | 0.064 | -0.246 | 0.816 |
| PPE41 | -0.341 | 0.999 | -0.181 | 0.859 | 0.160 | 0.840 |
| PE25 | -0.123 | 0.999 | -0.522 | 0.826 | -0.398 | 0.816 |
| Rv2432c | -0.144 | 0.999 | -0.121 | 0.922 | 0.024 | 0.981 |
| Rv2433c | 0.096 | NA | -0.165 | 0.922 | -0.262 | 0.862 |
| Rv2434c | -0.030 | 0.999 | -0.358 | 0.820 | -0.328 | 0.816 |
| devS | -0.272 | 0.999 | -0.552 | 0.001 | -0.280 | 0.816 |
| devR | -0.490 | 0.757 | -0.905 | 0.000 | -0.415 | 0.816 |
| Rv3134c | -0.504 | 0.467 | -0.618 | 0.050 | -0.114 | 0.862 |
| PPE50 | -0.158 | 0.999 | -0.291 | 0.832 | -0.133 | 0.870 |
| PPE51 | -0.229 | 0.999 | 0.063 | 0.940 | 0.292 | 0.816 |
| Rv3136A | -0.196 | NA | 0.177 | 0.879 | 0.373 | 0.816 |
| Rv3137 | -0.177 | 0.999 | -0.055 | 0.879 | 0.122 | 0.816 |
| pflA | -0.070 | 0.999 | -0.157 | 0.826 | -0.086 | 0.832 |
| fadE24 | -0.178 | 0.999 | -0.114 | 0.848 | 0.064 | 0.879 |
| fadE23 | -0.226 | 0.999 | -0.262 | 0.769 | -0.036 | 0.939 |
| fadB4 | -0.104 | 0.999 | 0.086 | 0.879 | 0.190 | 0.816 |
| Rv3142c | 0.082 | 0.999 | -0.304 | 0.826 | -0.387 | 0.816 |
| Rv3143 | -0.043 | NA | -0.024 | 0.976 | 0.019 | 0.985 |
| PPE52 | 0.188 | 0.999 | 0.263 | 0.879 | 0.075 | 0.951 |
| nuoA | 0.221 | 0.999 | 0.191 | 0.826 | -0.031 | 0.944 |
| nuoB | 0.227 | 0.999 | -0.028 | 0.967 | -0.254 | 0.816 |
| nuoC | 0.236 | 0.999 | -0.023 | 0.975 | -0.260 | 0.816 |
| nuoD | 0.288 | 0.999 | -0.117 | 0.879 | -0.406 | 0.816 |
| nuoE | 0.206 | 0.999 | 0.141 | 0.848 | -0.065 | 0.896 |
| nuoF | 0.173 | 0.999 | 0.150 | 0.859 | -0.023 | 0.969 |
| nuoG | 0.211 | 0.999 | 0.188 | 0.848 | -0.024 | 0.970 |
| nuoH | 0.194 | 0.999 | 0.079 | 0.922 | -0.115 | 0.862 |
| nuoI | 0.291 | 0.999 | -0.005 | 0.995 | -0.296 | 0.835 |
| nuoJ | 0.166 | 0.999 | 0.136 | 0.873 | -0.030 | 0.958 |
| nuoK | 0.204 | 0.999 | -0.267 | 0.885 | -0.470 | 0.816 |
| nuoL | 0.207 | 0.999 | 0.309 | 0.831 | 0.103 | 0.899 |
| nuoM | 0.124 | 0.999 | 0.049 | 0.950 | -0.075 | 0.904 |
| nuoN | 0.131 | 0.999 | 0.225 | 0.848 | 0.094 | 0.899 |
| PPE53 | -0.125 | 0.999 | -0.345 | 0.927 | -0.220 | 0.942 |
| Rv3160c | 0.047 | 0.999 | -0.168 | 0.832 | -0.214 | 0.816 |
| Rv3161c | -0.022 | 0.999 | 0.044 | 0.934 | 0.066 | 0.879 |
| Rv3162c | -0.052 | NA | -0.459 | 0.831 | -0.407 | 0.816 |
| Rv3163c | -0.013 | 0.999 | 0.025 | 0.970 | 0.038 | 0.945 |
| moxR3 | -0.041 | 0.999 | 0.034 | 0.948 | 0.074 | 0.865 |
| Rv3165c | 0.153 | NA | -0.059 | 0.970 | -0.212 | 0.868 |
| Rv3166c | -0.014 | 0.999 | 0.516 | 0.701 | 0.530 | 0.816 |
| Rv3167c | -0.099 | NA | 0.057 | 0.961 | 0.156 | 0.868 |
| Rv3168 | 0.219 | 0.999 | -0.098 | 0.952 | -0.317 | 0.837 |
| Rv3169 | -0.042 | 0.999 | -0.156 | 0.879 | -0.114 | 0.887 |
| aofH | 0.008 | 0.999 | 0.298 | 0.829 | 0.290 | 0.816 |
| hpx | -0.145 | 0.999 | 0.549 | 0.826 | 0.694 | 0.816 |
| Rv3172c | 0.034 | 0.999 | -0.327 | 0.854 | -0.361 | 0.817 |
| Rv3173c | -0.037 | 0.999 | 0.049 | 0.951 | 0.086 | 0.896 |
| Rv3174 | 0.112 | NA | 0.082 | 0.942 | -0.031 | 0.973 |
| Rv3175 | 0.097 | NA | -0.561 | 0.846 | -0.658 | 0.816 |
| mesT | 0.015 | 0.999 | 0.044 | 0.964 | 0.029 | 0.970 |
| Rv3177 | 0.333 | NA | -0.481 | 0.848 | -0.814 | 0.816 |
| Rv3178 | 0.067 | NA | -0.895 | 0.834 | -0.963 | 0.816 |
| Rv3178a | -0.811 | NA | -2.081 | 0.829 | -1.270 | 0.835 |
| Rv3179 | -0.139 | NA | -0.310 | 0.847 | -0.171 | 0.868 |
| Rv3180c | -0.111 | NA | -0.022 | 0.978 | 0.090 | 0.920 |
| Rv3181c | -0.137 | NA | -0.057 | 0.967 | 0.080 | 0.942 |
| Rv3182 | -0.067 | 0.999 | -0.114 | 0.948 | -0.047 | 0.973 |
| Rv3183 | -0.168 | NA | -0.128 | 0.894 | 0.040 | 0.957 |
| Rv3188 | -0.164 | NA | -0.323 | 0.826 | -0.160 | 0.854 |
| Rv3189 | -0.027 | NA | -0.587 | 0.826 | -0.560 | 0.816 |
| Rv3190A | 0.028 | 0.999 | -0.264 | 0.879 | -0.291 | 0.840 |
| Rv3190c | 0.120 | 0.999 | -0.042 | 0.930 | -0.163 | 0.816 |
| Rv3191c | -0.039 | NA | -0.483 | NA | -0.444 | NA |
| Rv3192 | -0.238 | NA | -0.228 | 0.930 | 0.010 | 0.997 |
| Rv3193c | 0.074 | 0.999 | -0.083 | 0.886 | -0.157 | 0.816 |
| Rv3194c | 0.074 | 0.999 | -0.111 | 0.854 | -0.185 | 0.816 |
| Rv3195 | -0.236 | 0.999 | -0.031 | 0.942 | 0.205 | 0.816 |
| Rv3196 | 0.320 | NA | -0.173 | 0.922 | -0.493 | 0.816 |
| Rv3196A | 0.081 | 0.999 | -0.134 | 0.895 | -0.215 | 0.831 |
| Rv3197 | 0.021 | 0.999 | -0.149 | 0.728 | -0.170 | 0.816 |
| whiB7 | -0.003 | 0.999 | 0.405 | 0.826 | 0.407 | 0.816 |
| Rv3198A | 0.185 | 0.999 | -0.201 | 0.859 | -0.386 | 0.816 |
| uvrD2 | 0.025 | 0.999 | 0.086 | 0.879 | 0.061 | 0.884 |
| nudC | -0.084 | 0.999 | 0.220 | 0.848 | 0.304 | 0.816 |
| Rv3200c | 0.039 | 0.999 | -0.071 | 0.931 | -0.110 | 0.868 |
| Rv3201c | 0.015 | 0.999 | 0.176 | 0.865 | 0.161 | 0.846 |
| Rv3202a | -0.107 | NA | -0.149 | 0.879 | -0.042 | 0.954 |
| Rv3202c | -0.010 | 0.999 | 0.121 | 0.892 | 0.131 | 0.866 |
| lipV | -0.088 | 0.999 | 0.071 | 0.964 | 0.160 | 0.902 |
| Rv3204 | -0.017 | NA | 0.036 | 0.970 | 0.054 | 0.942 |
| Rv3205c | -0.076 | 0.999 | 0.065 | 0.927 | 0.142 | 0.835 |
| moeB1 | 0.019 | 0.999 | 0.086 | 0.886 | 0.066 | 0.896 |
| Rv3207c | -0.024 | 0.999 | 0.124 | 0.859 | 0.149 | 0.817 |
| Rv3208 | 0.032 | 0.999 | 0.007 | 0.986 | -0.025 | 0.951 |
| TB9.4 | 0.254 | 0.999 | 0.022 | 0.975 | -0.231 | 0.816 |
| Rv3209 | 0.097 | 0.999 | -0.102 | 0.879 | -0.199 | 0.816 |
| Rv3210c | 0.066 | 0.999 | 0.125 | 0.879 | 0.060 | 0.930 |
| rhlE | -0.009 | 0.999 | -0.041 | 0.933 | -0.033 | 0.936 |
| Rv3212 | -0.075 | 0.999 | 0.404 | 0.829 | 0.479 | 0.816 |
| Rv3213c | -0.071 | 0.999 | 0.044 | 0.948 | 0.115 | 0.854 |
| gpm2 | -0.150 | 0.999 | 0.018 | 0.982 | 0.168 | 0.858 |
| entC | 0.068 | 0.999 | -0.264 | 0.859 | -0.332 | 0.816 |
| Rv3217c | 0.041 | 0.999 | -0.097 | 0.848 | -0.137 | 0.816 |
| Rv3218 | 0.120 | 0.999 | 0.004 | 0.993 | -0.116 | 0.831 |
| whiB1 | -0.173 | 0.999 | -0.005 | 0.990 | 0.168 | 0.816 |
| Rv3220c | 0.036 | 0.999 | 0.230 | 0.831 | 0.194 | 0.816 |
| rshA | -0.058 | 0.999 | 0.126 | 0.885 | 0.184 | 0.831 |
| TB7.3 | -0.119 | 0.999 | -0.096 | 0.948 | 0.023 | 0.986 |
| Rv3222c | -0.192 | 0.999 | 0.095 | 0.922 | 0.288 | 0.816 |
| sigH | 0.068 | 0.999 | -0.254 | 0.848 | -0.322 | 0.816 |
| Rv3224 | 0.022 | 0.999 | 0.108 | 0.878 | 0.087 | 0.866 |
| Rv3225c | -0.018 | 0.999 | -0.060 | 0.946 | -0.042 | 0.950 |
| Rv3226c | -0.095 | 0.999 | 0.080 | 0.928 | 0.175 | 0.835 |
| aroA | 0.129 | 0.999 | 0.367 | 0.826 | 0.239 | 0.830 |
| Rv3228 | -0.014 | 0.999 | 0.208 | 0.879 | 0.221 | 0.842 |
| desA3 | 0.171 | 0.999 | -0.113 | 0.879 | -0.283 | 0.816 |
| Rv3230c | -0.225 | 0.999 | 0.073 | 0.950 | 0.298 | 0.816 |
| Rv3231c | 0.055 | 0.999 | 0.080 | 0.936 | 0.025 | 0.975 |
| ppk2 | -0.017 | 0.999 | -0.099 | 0.922 | -0.081 | 0.920 |
| tgs3 | 0.042 | 0.999 | 0.070 | 0.905 | 0.028 | 0.951 |
| Rv3235 | -0.054 | 0.999 | -0.065 | 0.936 | -0.011 | 0.987 |
| Rv3236c | -0.083 | 0.999 | 0.268 | 0.831 | 0.351 | 0.816 |
| Rv3237c | 0.001 | 0.999 | -0.181 | 0.879 | -0.183 | 0.846 |
| Rv3238c | 0.116 | 0.999 | 0.079 | 0.895 | -0.037 | 0.942 |
| mcr15 | 0.170 | 0.999 | -0.377 | 0.879 | -0.547 | 0.819 |
| mcr15 | 0.170 | 0.999 | -0.377 | 0.879 | -0.547 | 0.819 |
| mcr16 | -0.259 | NA | 0.202 | 0.948 | 0.461 | 0.865 |
| mcr16 | -0.259 | NA | 0.202 | 0.948 | 0.461 | 0.865 |
| mcr3 | -0.481 | 0.999 | 0.767 | 0.855 | 1.248 | 0.816 |
| mcr3 | -0.481 | 0.999 | 0.767 | 0.855 | 1.248 | 0.816 |
| mcr7 | -0.244 | 0.999 | 0.744 | 0.826 | 0.989 | 0.816 |
| mcr7 | -0.244 | 0.999 | 0.744 | 0.826 | 0.989 | 0.816 |
| MTS0858 | 0.125 | NA | 0.232 | 0.931 | 0.107 | 0.958 |
| MTS0858 | 0.125 | NA | 0.232 | 0.931 | 0.107 | 0.958 |
| MTS1082 | -0.015 | 0.999 | -0.104 | 0.883 | -0.088 | 0.879 |
| MTS1082 | -0.015 | 0.999 | -0.104 | 0.883 | -0.088 | 0.879 |
| MTS1338 | -0.377 | 0.999 | 0.558 | 0.879 | 0.934 | 0.816 |
| MTS1338 | -0.377 | 0.999 | 0.558 | 0.879 | 0.934 | 0.816 |
| MTS2823 | -0.034 | 0.999 | 0.578 | 0.831 | 0.612 | 0.816 |
| MTS2823 | -0.034 | 0.999 | 0.578 | 0.831 | 0.612 | 0.816 |
| MTS2975 | 0.115 | 0.999 | 0.724 | 0.522 | 0.609 | 0.816 |
| MTS2975 | 0.115 | 0.999 | 0.724 | 0.522 | 0.609 | 0.816 |
| ssr | -0.056 | 0.999 | 0.046 | 0.962 | 0.101 | 0.896 |
| ssr | -0.056 | 0.999 | 0.046 | 0.962 | 0.101 | 0.896 |
| AS1726 | -0.144 | NA | -0.659 | 0.589 | -0.514 | 0.816 |
| AS1726 | -0.144 | NA | -0.659 | 0.589 | -0.514 | 0.816 |
| rrs | -0.840 | NA | -4.472 | NA | -3.632 | NA |
| rrs | -0.840 | NA | -4.472 | NA | -3.632 | NA |
| rrl | -0.110 | 0.999 | -1.109 | 0.458 | -0.999 | 0.816 |
| rrl | -0.110 | 0.999 | -1.109 | 0.458 | -0.999 | 0.816 |
| rrf | -0.258 | 0.999 | -0.330 | 0.885 | -0.072 | 0.967 |
| rrf | -0.258 | 0.999 | -0.330 | 0.885 | -0.072 | 0.967 |
| rnpB | -0.273 | 0.999 | 0.194 | 0.899 | 0.467 | 0.816 |
| rnpB | -0.273 | 0.999 | 0.194 | 0.899 | 0.467 | 0.816 |
| ileT | 0.127 | NA | -0.006 | 0.995 | -0.133 | 0.909 |
| ileT | 0.127 | NA | -0.006 | 0.995 | -0.133 | 0.909 |
| alaT | 0.003 | NA | -0.225 | 0.848 | -0.227 | 0.819 |
| alaT | 0.003 | NA | -0.225 | 0.848 | -0.227 | 0.819 |
| leuT | -0.079 | 0.999 | 0.233 | 0.880 | 0.312 | 0.835 |
| leuT | -0.079 | 0.999 | 0.233 | 0.880 | 0.312 | 0.835 |
| glyU | 0.210 | NA | 0.342 | 0.832 | 0.131 | 0.896 |
| glyU | 0.210 | NA | 0.342 | 0.832 | 0.131 | 0.896 |
| tyrT | -0.142 | NA | 0.767 | 0.834 | 0.909 | 0.816 |
| tyrT | -0.142 | NA | 0.767 | 0.834 | 0.909 | 0.816 |
| thrT | -0.100 | NA | 0.548 | 0.843 | 0.648 | 0.816 |
| thrT | -0.100 | NA | 0.548 | 0.843 | 0.648 | 0.816 |
| metT | 0.232 | 0.999 | -0.229 | 0.859 | -0.461 | 0.816 |
| metT | 0.232 | 0.999 | -0.229 | 0.859 | -0.461 | 0.816 |
| trpT | -0.108 | 0.999 | -0.117 | 0.879 | -0.009 | 0.988 |
| trpT | -0.108 | 0.999 | -0.117 | 0.879 | -0.009 | 0.988 |
| thrV | -0.228 | NA | 0.056 | 0.971 | 0.284 | 0.852 |
| thrV | -0.228 | NA | 0.056 | 0.971 | 0.284 | 0.852 |
| lysT | 0.329 | NA | -0.792 | 0.848 | -1.120 | 0.816 |
| lysT | 0.329 | NA | -0.792 | 0.848 | -1.120 | 0.816 |
| gluT | -0.030 | NA | 0.482 | 0.859 | 0.512 | 0.822 |
| gluT | -0.030 | NA | 0.482 | 0.859 | 0.512 | 0.822 |
| aspT | -0.406 | NA | 0.848 | 0.826 | 1.254 | 0.816 |
| aspT | -0.406 | NA | 0.848 | 0.826 | 1.254 | 0.816 |
| pheU | -0.090 | NA | 0.090 | 0.947 | 0.180 | 0.868 |
| pheU | -0.090 | NA | 0.090 | 0.947 | 0.180 | 0.868 |
| argT | 0.223 | NA | -0.078 | 0.960 | -0.301 | 0.835 |
| argT | 0.223 | NA | -0.078 | 0.960 | -0.301 | 0.835 |
| alaV | -0.021 | NA | 0.214 | 0.879 | 0.235 | 0.842 |
| alaV | -0.021 | NA | 0.214 | 0.879 | 0.235 | 0.842 |
| glnT | -0.609 | NA | -1.143 | 0.843 | -0.534 | 0.882 |
| glnT | -0.609 | NA | -1.143 | 0.843 | -0.534 | 0.882 |
| leuX | -0.209 | NA | -0.218 | 0.860 | -0.009 | 0.994 |
| leuX | -0.209 | NA | -0.218 | 0.860 | -0.009 | 0.994 |
| argV | -0.205 | NA | 0.735 | 0.826 | 0.940 | 0.816 |
| argV | -0.205 | NA | 0.735 | 0.826 | 0.940 | 0.816 |
| leuW | -0.256 | NA | -0.268 | 0.883 | -0.012 | 0.994 |
| leuW | -0.256 | NA | -0.268 | 0.883 | -0.012 | 0.994 |
| leuV | -0.730 | NA | -0.957 | 0.854 | -0.227 | 0.947 |
| leuV | -0.730 | NA | -0.957 | 0.854 | -0.227 | 0.947 |
| proT | 0.040 | NA | 0.433 | 0.826 | 0.393 | 0.816 |
| proT | 0.040 | NA | 0.433 | 0.826 | 0.393 | 0.816 |
| leuU | -0.805 | NA | -1.645 | 0.826 | -0.840 | 0.836 |
| leuU | -0.805 | NA | -1.645 | 0.826 | -0.840 | 0.836 |
| valV | 0.172 | NA | 0.071 | 0.977 | -0.101 | 0.970 |
| valV | 0.172 | NA | 0.071 | 0.977 | -0.101 | 0.970 |
| metV | -0.119 | NA | 0.137 | 0.885 | 0.256 | 0.816 |
| metV | -0.119 | NA | 0.137 | 0.885 | 0.256 | 0.816 |
| asnT | -0.114 | NA | -0.151 | 0.890 | -0.037 | 0.967 |
| asnT | -0.114 | NA | -0.151 | 0.890 | -0.037 | 0.967 |
| proU | -0.088 | NA | -0.577 | 0.858 | -0.489 | 0.846 |
| proU | -0.088 | NA | -0.577 | 0.858 | -0.489 | 0.846 |
| glyV | -0.168 | NA | 1.453 | 0.155 | 1.621 | 0.464 |
| glyV | -0.168 | NA | 1.453 | 0.155 | 1.621 | 0.464 |
| argW | 0.144 | NA | 0.022 | 0.990 | -0.122 | 0.951 |
| argW | 0.144 | NA | 0.022 | 0.990 | -0.122 | 0.951 |
| hisT | -0.034 | NA | 0.446 | 0.831 | 0.480 | 0.816 |
| hisT | -0.034 | NA | 0.446 | 0.831 | 0.480 | 0.816 |
| lysU | 0.051 | 0.999 | -0.150 | 0.914 | -0.201 | 0.862 |
| lysU | 0.051 | 0.999 | -0.150 | 0.914 | -0.201 | 0.862 |
| valT | -0.091 | 0.999 | -0.059 | 0.962 | 0.032 | 0.975 |
| valT | -0.091 | 0.999 | -0.059 | 0.962 | 0.032 | 0.975 |
| glyT | -0.246 | NA | 0.257 | 0.879 | 0.503 | 0.816 |
| glyT | -0.246 | NA | 0.257 | 0.879 | 0.503 | 0.816 |
| cysU | -0.366 | NA | 0.891 | 0.826 | 1.256 | 0.816 |
| cysU | -0.366 | NA | 0.891 | 0.826 | 1.256 | 0.816 |
| valU | 0.042 | NA | -0.323 | 0.879 | -0.365 | 0.837 |
| valU | 0.042 | NA | -0.323 | 0.879 | -0.365 | 0.837 |
| gluU | -0.586 | NA | 0.229 | 0.927 | 0.815 | 0.816 |
| gluU | -0.586 | NA | 0.229 | 0.927 | 0.815 | 0.816 |
| glnU | 0.039 | NA | 0.235 | 0.922 | 0.196 | 0.921 |
| glnU | 0.039 | NA | 0.235 | 0.922 | 0.196 | 0.921 |
| alaU | 0.159 | NA | -0.008 | 0.994 | -0.166 | 0.891 |
| alaU | 0.159 | NA | -0.008 | 0.994 | -0.166 | 0.891 |
| metU | -0.445 | NA | -0.248 | 0.903 | 0.198 | 0.904 |
| metU | -0.445 | NA | -0.248 | 0.903 | 0.198 | 0.904 |
| thrU | -0.204 | NA | 0.544 | 0.831 | 0.748 | 0.816 |
| thrU | -0.204 | NA | 0.544 | 0.831 | 0.748 | 0.816 |
| proY | -0.034 | NA | -0.077 | 0.948 | -0.043 | 0.961 |
| proY | -0.034 | NA | -0.077 | 0.948 | -0.043 | 0.961 |
| serV | 0.072 | NA | 0.281 | 0.886 | 0.209 | 0.896 |
| serV | 0.072 | NA | 0.281 | 0.886 | 0.209 | 0.896 |
| serX | -0.109 | NA | 1.051 | 0.671 | 1.160 | 0.816 |
| serX | -0.109 | NA | 1.051 | 0.671 | 1.160 | 0.816 |
| argU | 0.119 | NA | -0.134 | 0.935 | -0.253 | 0.862 |
| argU | 0.119 | NA | -0.134 | 0.935 | -0.253 | 0.862 |
| serT | -0.063 | NA | 0.222 | 0.895 | 0.285 | 0.855 |
| serT | -0.063 | NA | 0.222 | 0.895 | 0.285 | 0.855 |
| serU | -0.551 | NA | 0.151 | 0.941 | 0.702 | 0.816 |
| serU | -0.551 | NA | 0.151 | 0.941 | 0.702 | 0.816 |
